# Kidney collecting duct cell type composition is regulated by Notch signaling via modulation of mTORC1

**DOI:** 10.1101/2024.04.09.587573

**Authors:** Jennifer deRiso, Malini Mukherjee, Madhusudhana Janga, Alicia Simmons, Michael Kareta, Jianning Tao, Indra Chandrasekar, Kameswaran Surendran

## Abstract

The plasticity and diversity of cell types with specialized functions likely defines the capacity of multicellular organisms to adapt to physiologic stressors. The kidney collecting ducts contribute to water, electrolyte, and pH homeostasis and are composed of mature intermingled epithelial cell types that are susceptible to transdifferentiate. The conversion of kidney collecting duct principal cells to intercalated cells is actively inhibited by Notch signaling to ensure urine concentrating capability. Here we identify Hes1, a target of Notch signaling, allows for maintenance of functionally distinct epithelial cell types within the same microenvironment by regulating mechanistic target of rapamycin complex 1 (mTORC1) activity. Hes1 directly represses the expression of *insulin receptor substrate 1* (*Irs1*), an upstream component of mTOR pathway and suppresses mTORC1 activity in principal cells. Genetic inactivation of *tuberous sclerosis complex 2* (*Tsc2*) to increase mTORC1 activity in mature principal cells is sufficient to promote acquisition of intercalated cell properties, while inhibition of mTORC1 in adult kidney epithelia suppresses intercalated cell properties. Considering that mTORC1 integrates environmental cues, the linkage of functionally distinct epithelial cell types to mTORC1 activity levels likely allows for cell plasticity to be regulated by physiologic and metabolic signals and the ability to sense/transduce these signals.

## Introduction

Multicellular organisms rely on the diversity of cell types with specialized functions for the maintenance of their internal milieu. An example, multiple different kidney epithelial cell types are required to ensure a stable internal environment by removing unwanted metabolic products while maintaining water and ion balance. Remodeling the nephron and collecting duct segments of the kidney by altering the ratios of the constituent epithelial cell types is likely a mechanism that enables the adaptive capacity necessary to sustain water, electrolyte and pH homeostasis when exposed to physiologic stressors ^1–4^. The distal nephron and collecting duct (CD) segments consist of intercalated cell (IC) types intermingled among principal cell (PC) types and can undergo changes in cell type composition, some of which involves direct conversion of PCs to ICs ^1,5–7^. Consistent with plasticity of collecting duct cells, single cell transcriptomic analysis has revealed cells in transition between cell types^8^. The plasticity of the collecting duct cell types has been further established by observations of direct conversion of PCs to ICs and the existence of transitional cells expressing both PC and IC markers ^7–9^, as well as altered ratios of PCs to ICs in cystic kidney diseases including in models of tuberous sclerosis complex (tsc) 1 and polycystic kidney disease (pkd) 2 deficiency^10,11^.

Drastic alterations in ratios of principal and intercalated cell types correlate with the occurrence of water and electrolyte imbalances. For example, lithium used to treat bipolar disorder can trigger an acquired form of nephrogenic diabetes insipidus (NDI), involving an inability to sufficiently reabsorb water, along with a decrease in ratio of PCs to ICs which is in part driven by increased PC to IC conversion as determined by PC-lineage tracing experiments ^7,12,13^. Additionally, potassium (K^+^) deficient diets increase the ratio of ICs to PCs in the outer medullary (OM) CD segments and can result in NDI ^5,6^. However, the molecular mechanisms that mediate alterations in the cell type composition of distal tubules and collecting ducts remain largely unknown. Also, it is unclear how neighboring epithelial cells within a tubule retain different identities and functional properties while subjected to the same microenvironment.

One insight into the regulation of kidney collecting duct cell type composition comes from studies in rodents that establish an essential role for Notch signaling in both the development and maintenance of mature PC types in the distal nephron and collecting duct segments of the kidney ^7,14–16^. Along with an increase in IC types and a decrease in PC types, an NDI-like phenotype occurs in mice with conditional inactivation of Notch signaling in the adult kidney epithelial cells ^7^. Within the collecting ducts the ICs express the Notch ligands Jagged1 (Jag1) and Delta-like 1 (Dll1) and the receptors Notch1 and Notch2 are activated in PCs and in turn induce the expression of hairy enhancer of split-1 (Hes1) to maintain cells in a mature PC state in adult kidney ^7,14,17,18^. This is consistent with Notch signaling occurring between neighboring cells to regulate cell fate and function ^19–21^, without which the PCs stop expressing proteins critical for PC function such as Aquaporin 2 and Aquaporin 4, and convert directly into ICs by expressing genes critical for IC functions ^7^. Hes1 is a direct target of Notch signaling that functions as a transcriptional repressor to mediate cell fate choices ^22,23^. While it is likely that the primary function of Hes1 in PCs is to suppress the expression of Foxi1 ^17^, a master transcriptional regulator of IC types ^24–26^, the mechanisms by which Hes1 inhibits Foxi1 expression to ensure PC state are unknown. One hypothesis is that local intercalated cell promoting signals instruct CD segment epithelial cell types to maintain an IC identity by ensuring Foxi1 expression, while Notch signaling actively interferes with the reception and/or transduction of these local signals and in effect prevents the IC state. Since we currently do not know the signals that promote IC properties, we sought to identify IC promoting signaling pathways by determining the genes directly repressed by Hes1 in adult kidney PC types that code for signal transduction components.

Our study reveals that Notch signaling regulates mature kidney collecting duct cell type composition by suppressing the activity of mechanistic target of rapamycin complex 1 (mTORC1) within PCs. The mTORC1 consists of six protein components including the regulatory-associated protein of mechanistic target of rapamycin (Raptor) and alters cellular state in response to environmental cues by modulating processes such as protein synthesis and autophagy ^27^. Interestingly, we find mTORC1 activity within the adult kidney collecting ducts is enriched in ICs and increasing the levels of mTORC1 activity in mature PCs promotes the IC state. We find that an upstream regulator of mTORC1, insulin receptor substrate 1 (Irs1), is specifically expressed in ICs and not PCs, and this selective expression is regulated by Notch signaling. Within the adult kidney collecting ducts, Notch signaling suppresses mTORC1 activity in PCs and this ensures that PCs do not gain IC properties.

## Methods

### Mice

All experiments with mice were approved by the Sanford Research IACUC. All mouse lines used in this study have been previously described and are listed in Supplementary Table 1. Mice used in this study were maintained on mixed backgrounds and were genotyped following a universal PCR genotyping protocol ^28^. Primer sequences are available upon request. Mice were fed normal chow ad libitum. Wild-type control groups consisted of age and sex-matched littermates housed along with mice with conditional inactivation of gene(s) of interest. Male and female mice were included in each group when comparing wild type versus mutant groups. For conditional inactivation we used *Pax8->rtTA* ^29^ and *TRE->Cre* ^30^ as previously described ^7^ to recombine *Hes1* or *Rptor* floxed alleles ^31,32^. Mice were given doxycycline (1 mg/mL; Sigma-Aldrich) in drinking water containing 5% sucrose for two weeks starting at 3 to 4 weeks of age.

The *Pax8->rtTA*; *TRE->Cre*; *Rptor f/f* mice (*Raptor* cKO) and control littermate mice were analyzed between 2 and 3 weeks after completion of doxycycline treatment. For early time points *Pax8->rtTA*; *TRE->Cre*; *Hes1 f/f* mice (*Hes1* cKO) and control littermate mice were given 1 mg/ml doxycycline for 4 or 7 days before analysis. Twenty-four-hour mouse urine samples were collected by housing mice individually in metabolic cages. The urine osmolality was determined using a VAPRO Vapor Pressure Osmometer (ELITechGroup) and urine pH determined using Mettler Toledo micro-pH electrode (01-913-998, Fisher). Age and sex matched offspring from *Cdh16->Cre*; TSC2*^+/f^* mice bred with *TSC2^f/f^* mice were analyzed. For the lineage tracing experiments *Elf5->rtTA; TRE->Cre* mice were bred with *Rosa^tdtomato/tdtomato^* to generate *Elf5->rtTA; TRE->Cre: Rosa^+/tdtomato^*and were compared with *Elf5->rtTA;TRE->Cre; Tsc1^+/*^; Tsc2^f/f^; Rosa^+/tdtomato^* (* denotes + or f) mice which were generated by breeding *Elf5->rtTA;TRE->Cre; Tsc2^f/f^* mice with *Tsc1^+/f^; Tsc2^f/f^; Rosa^tdtomato/tdtomato^*.

### Lineage tracing

*Elf5->rtTA; TRE->Cre; Rosa^+/tdtomato^* (wild type control group) and *Elf5->rtTA;TRE->Cre; Tsc1^+/*^; Tsc2^f/f^; Rosa^+/tdtomato^* (* denotes + or f; experimental group) were administered 1 mg/ml dox in the drinking water for two weeks from approximately 3 to 5 weeks of age, to turn on tdtomato expression in Elf5-expressing PCs within the collecting duct cells while also inactivating *Tsc1* and *Tsc2* floxed alleles in the experimental group. These mice were perfusion-fixed using cold 4% PFA/ 0.1 M cacodylate at 6 to 8 weeks of age and kidneys were isolated and processed for immunohistochemistry. A minimum of 10 cortical collecting duct and 20 medullary collecting duct segments per mouse kidney were analyzed. The number of cells expressing tdtomato, or c-kit and tdtomato were counted to calculate the percentage of principal lineage cells expressing IC markers. The presence of glomeruli in the region served to distinguish the cortex from the medulla.

### RNA-sequencing and RT-qPCR

Whole mouse kidneys were homogenized in lysis buffer (Qiagen) and total RNA was extracted from a portion of the lysate using RNeasy Mini kit. Following validation of RNA quality using a bioanalyzer, 150 bp paired-end sequencing was performed on the Illumina NovaSeq platform at Novogene. Sequencing results were processed utilizing trimmomatic software ^33^ and the quality of the processed reads verified using FASTQC. Filtered reads were then aligned to the mouse genome using HISAT2 ^34^. The resulting output from the HISAT2 program was then analyzed to quantify the reads per gene using the HTSeq count tool ^35^. The differentially expressed genes were identified using DESeq2 ^36^. An adjusted p-value <0.05 was used to determine differentially expressed genes. RNA-seq results were submitted to GEO (GSE260474). For RT-qPCR four micrograms of the extracted RNA from each sample were reverse transcribed with gene specific primers using Goscript reverse transcription kit (Promega). Quantitative PCR was performed using Power SYBR Green (Life Technologies), gene specific forward and reverse primers designed from different exons, and BioRad CFX-96 real-time PCR machine. Standard curves were generated by analyzing serially diluted cDNA reverse transcribed from mouse kidneys to determine the efficiency of each primer pair. Primer sequences are available upon request. Each sample was measured either in duplicate or triplicate and relative gene expression levels were normalized to that of *GAPDH* or *beta-2 microglobulin*.

### Chromatin immunoprecipitation (ChIP)

The ChIP experiments were done with mpkCCDc14 cells transfected with FLAG-tagged mouse Hes1 (Hes1-FLAG) under a Tet-responsive element and reverse tetracycline transactivator (rtTA). Cells were treated with 1µg/mL doxycycline for 72hrs and then crosslinked with 1% formaldehyde (Fisher Scientific, BP531-25) for 10 minutes at room temperature with rocking. The formaldehyde was quenched with 125mM glycine for 5 minutes at room temperature. Cells were then rinsed twice with ice cold PBS, scraped and lysed in buffer containing 10 mM PIPES pH 8.0, 85 mM KCl, 0.5% NP-40, 0.5 mM PMSF and protease inhibitors. The lysate was centrifuged at 2000 rpm for 10 minutes at 4°C and the pellet was resuspended in 120 µl nuclei lysis buffer (NLB) consisting of 50 mM Tris pH 8.1, 10 mM EDTA, 0.5 mM PMSF, and protease inhibitors and incubated on ice for 20 minutes. SDS was added to the final concentration of 1% and released chromatin solution was then transferred to microtubes with AFA fiber (Covaris, 520045) and sonicated for 14 cycles of 130 second each with peak incident power 70 and duty factor 20 in a Covaris ME220 focused ultrasonicator. Sonicated chromatin was transferred to a fresh microcentrifuge containing 500 µl NLB and centrifuged at 17000 g for 10 min at 4°C. A portion (50 µl) of the sonicated chromatin was de-crosslinked at 65°C with 300 mM NaCl with overnight to check shearing to 200-500 bp. For immunoprecipitation, the chromatin was precleared for 1hr at 4°C with pre-blocked ChIP-grade Protein A/G magnetic beads (Millipore Sigma, 16-663). The precleared lysate was divided into equal parts and immunoprecipitated using either anti-FLAG antibody (Sigma, F1804) or anti-mouse IgG (Jackson Immunoresearch, 715-005-150) overnight at 4°C with rotation. An aliquot equivalent to 10% of the volume of chromatin used for immunoprecipitation was kept as input control. After the overnight immunoprecipitation, pre-blocked Protein A/G magnetic beads were added to each reaction for 1 hr at 4°C. After bead incubation, the beads were washed twice each with low salt buffer (20 mM Tris-HCl pH 8.1,150 mM NaCl, 0.1% SDS, 2 mM EDTA, 1% Triton X-100), high salt buffer (20 mM Tris-HCl pH 8.1, 500 mM NaCl, 0.1% SDS, 2 mM EDTA, 1% Triton X-100) and lithium chloride buffer (10 mM Tris-HCl pH 8.1, 250 mM LiCl, 1% NP-40, 1 mM EDTA, 1% sodium deoxycholate) followed by a wash with TE buffer. After washes, the chromatin was eluted from the beads by two incubations of 100µl freshly prepared elution buffer (0.1 M NaHCO_3_, 1% SDS) for 15 minutes. The eluted chromatin was de-crosslinked at 65°C with 300mM NaCl for 4 hours, then 2 µl RNAse A (10 mg/mL) was added and incubated at 37°C for 30 minutes, followed by 2 µl Proteinase K (20 mg/mL) for 30 minutes at 55°C. The chromatin was purified with Qiaquik PCR purification kit (Qiagen, 28106) and used for next generation sequencing or for quantitative PCR. The ChIP-qPCR results were calculated as percent of input.

### ChIP-sequencing analysis

The precipitated genomic regions using anti-FLAG antibody from three independent repeats and two separate inputs were sequenced. The quality of the processed reads verified using FASTQC. Filtered reads were then aligned to the mouse genome (mm10) using bowtie2 (2.4.0) and peaks called using MACS2 (v2.2.6). Analysis of the Hes1::FLAG ChIP-seq from three repeats revealed 3039 common peaks of which ∼96% mapped to within 1 kb from transcriptional start sites (TSS). ChIP-seq results were deposited in GEO (GSE260474).

### Histology and Immunohistochemistry

For immunohistochemistry kidneys were cut in half longitudinally and one half fixed in Bouin’s fixative and the other half in 4% PFA fixative overnight at 4^0^C, rinsed in 70% ethanol prior to paraffin embedding and sectioning at 8 to 12 µm thickness. For immunohistochemistry requiring direct visualization of tdtomato, or Foxi1, bisected kidneys were fixed in 4% PFA for 20 minutes to 1 hour, rinsed in PBS, and incubated in 15% sucrose for 24 hours, and then another 24 hours in 30% sucrose before embedding in OCT on dry ice. Frozen tissue was then sectioned at a thickness of 12 µm thickness. Staining was done as previously described ^7^. In brief, paraffin embedded sections were deparaffinized and boiled in Trilogy (Cell Marque) for antigen unmasking and placed in PBS before blocking. OCT sections were placed directly in PBS before blocking. Sections were blocked in PBS containing 1% BSA, 0.2% powdered skim milk, and 0.3% Triton X-100 for at least 15 minutes at room temperature before incubation with primary antibodies overnight at 4°C. Primary antibody details are listed in Supplementary Table 2. Secondary antibodies conjugated with DyLight405, Alexa Fluor 488, Cy3, and Cy5 (Jackson ImmunoResearch) were used at 1:500 dilution to visualize the primary antibodies. For OCT embedded tissue, coverslips were mounted with VECTASHIELD fluorescence mounting media with or without DAPI (Vector Laboratories). For paraffin embedded tissue, Hoechst dye was used with the secondary antibody and coverslips were mounted with VECTASHEILD without DAPI. Coverslips were sealed with nail polish. Stained tissue sections were imaged on a Nikon A1 confocal microscope using NIS elements software for image acquisition.

### Statistical Analyses

Two-tailed unpaired t-tests were performed after testing for equal variance between groups using the F-test unless stated otherwise using Microsoft Excel. The resulting p-values are stated in the text and/or figure legends. Based on our prior studies three wild type versus three Notch-signaling deficient adult mice were sufficient to detect significant differences between the two groups. Here we continued with n = 3 to 9 per group with the exact numbers depending on the size of and genotype within the litters. For adult mice (3 weeks or older) male and female mice were analyzed separately to control for sex dependent variability. In the graphs, the height of each bar represents the mean, and the error bars represent one standard deviation.

## Results

### Identification of signaling pathways suppressed by Hes1 in the adult kidneys

The collecting duct segments of adult mammalian kidneys consist of IC types that are interspersed among PC types. Previous studies have established a critical requirement for Notch signaling via Hes1 in the maintenance of PC types, without which the PCs begin expressing Foxi1 ^7^, a critical transcriptional regulator of genes necessary for IC specific functions ^24^. Many collecting duct cells were observed to be in an intermediate state expressing both PC and IC markers within one week after initiation of *Hes1* inactivation in adult kidney epithelia and likely in the process of conversion from PC to IC ^7^. In order to identify the molecular mechanisms by which Hes1 maintains PCs in a mature state and prevents them from converting into ICs we determined the genes differentially expressed in the whole adult mouse kidneys one week after inactivation of *Hes1* in the kidney epithelial cells using *Pax8->rtTA;TRE-Cre;Hes1f/f* (Hes1cKO) mice. We compared 9 control littermates with 7 Hes1cKO adult mice, all of which were given doxycycline in the drinking water for one week before analysis. Comparison of RNA-sequencing results revealed 290 differentially expressed genes (DEG) with adjusted p-value <0.05, of which 229 genes are up regulated in Hes1cKO kidneys (Supplementary Table 3). Among the up regulated DEGs with highest log fold change and with the lowest adjusted p-values are genes critical for IC functions and not expressed in PCs such as *Foxi1*, *Slc26a4* (pendrin), *Slc4a9* (anion exchanger 4), *Hmx2*, and *Dll1* (Figure 1A). This is consistent with an increase in ICs following *Hes1* inactivation. With the intent of identifying signaling pathways that are modulated in response to loss of Hes1 we noticed that mTOR, Notch, Wnt, Hedgehog and TGF-beta signaling pathway components were also among the DEGs (Figure 1B). Of these signaling pathways, only the components of Notch and mTOR signaling pathways are significantly over-represented in the up regulated DEG list (Fig. 1C). The Notch signaling pathway ligands *Dll1* and *Jag1* are known to be expressed in ICs and are among the up regulated DEGs. While it may not be surprising that *Atp6v1g3*, *Atp6v1b1* and *Atp6v1c2* which code for vacuolar H^+^- ATPase subunits essential for IC functions are upregulated, several of the other up regulated mTOR signaling pathway component genes, such as *insulin receptor substrate1* (*Irs1*) and *growth factor receptor-bound protein10* (*Grb10)*, are expressed in ICs and not in PCs based on previously reported mouse adult kidney single cell RNA sequencing results (Fig.1D) ^37^.

**Figure 1.**
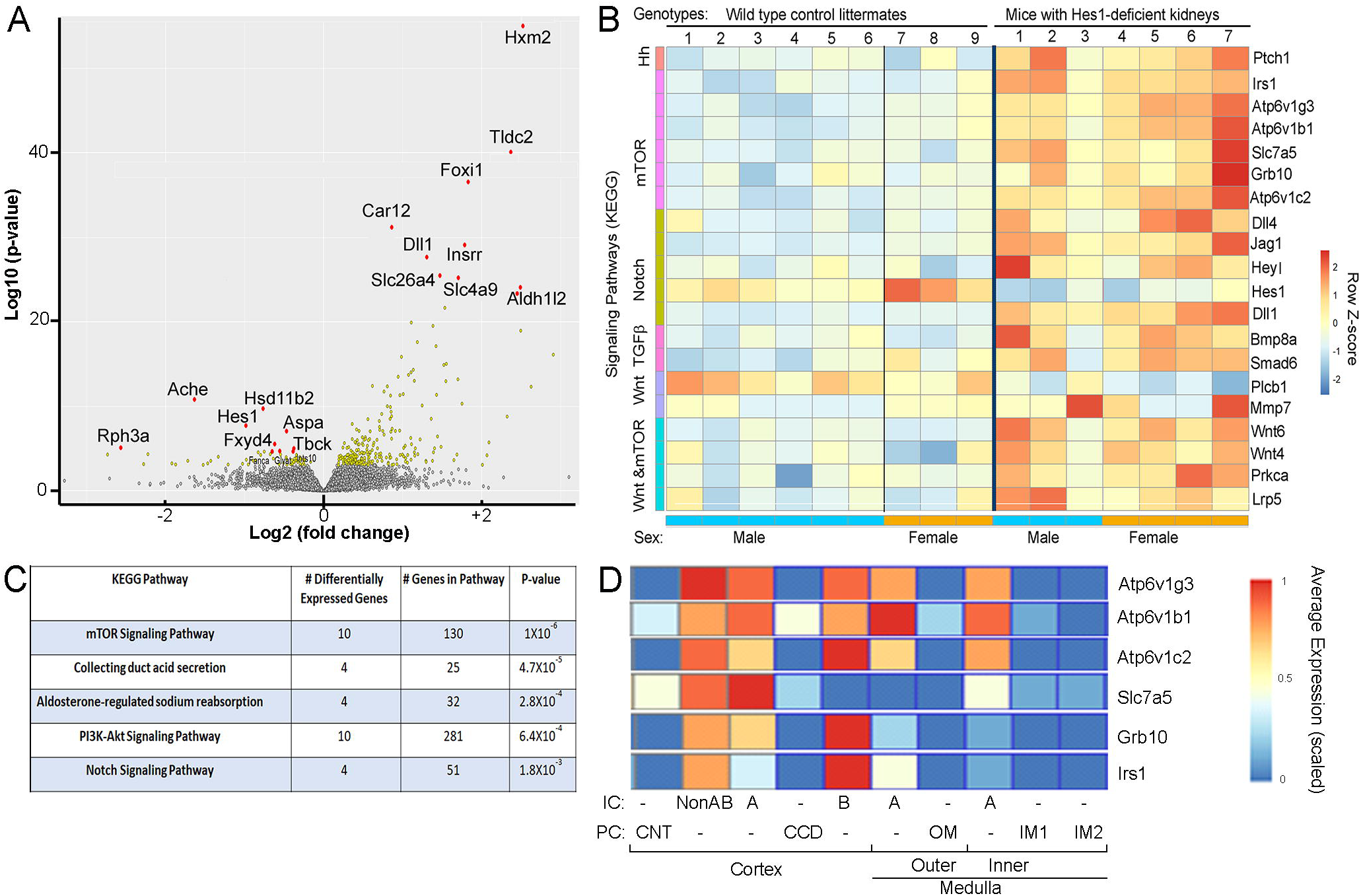
mTOR signaling pathway components are significantly enriched among the genes differentially expressed following inactivation of Hes1 in the adult mouse kidneys. **A.** Volcano plot highlighting the differentially expressed genes (DEGs) in yellow and red when comparing the kidneys of wild type control littermates (n=9) versus kidneys of mice one week after initiation of *Hes1* inactivation in the kidney epithelial cells (n=7). **B.** Heat map of DEGs that code for components of interesting signaling pathways reveals that Hes1-deficiency results in the up regulation of several components associated with mTOR signaling. **C.** Enrichment analysis revealed a significant over representation of mTOR/ PI3K-ATK signaling pathway components among the DEGs following *Hes1* inactivation. **D.** Heat map showing that several of the up regulated mTOR signaling pathway components in the Hes1-deficient kidneys are normally expressed in ICs and with low or no expression in PCs of wild type adult mouse kidneys. The heatmap is adapted from https://cello.shinyapps.io/kidneycellexplorer/.

### Insulin receptor substrate 1 (Irs1), a component upstream of mTORC1, is repressed by Hes1 in principal cells

To determine which if any of the DEGs are direct targets of Hes1 in PCs we performed sequencing of chromatin regions immunoprecipitated using anti-FLAG antibody from a mouse principal cortical collecting duct cell line (mpkCCDc14) ^38^ with ectopic expression of Hes1 fused with FLAG epitope (Figure 2A). Hes1-FLAG chromatin immunoprecipitation (ChIP) followed by sequencing revealed 3039 genomic regions that were consistently enriched in three different Hes1-FLAG ChIP samples. Mapping of the identified ChIP-seq peaks revealed that a majority (∼96%) of them are within 1 kb of a transcriptional start site (TSS). Of note, ingenuity pathway analysis revealed that several ChIP-seq peaks occurred near genes coding for PI3K-Akt-mTOR signaling pathway components, annotated as insulin signaling pathway (Supplementary Fig. S1). Among these regions visited by Hes1-FLAG in PCs is a region near the TSS of *insulin receptor substrate 1* (*Irs1)*. The ChIP-seq peak near *Irs1* TSS contains two E-box and one N-box DNA elements which are potential Hes1 binding sites. We validated by qPCR that chromatin fragments containing the N-box element near the *Irs1* TSS are enriched in the Hes1-FLAG ChIP (Fig. 2B &C).

**Figure 2.**
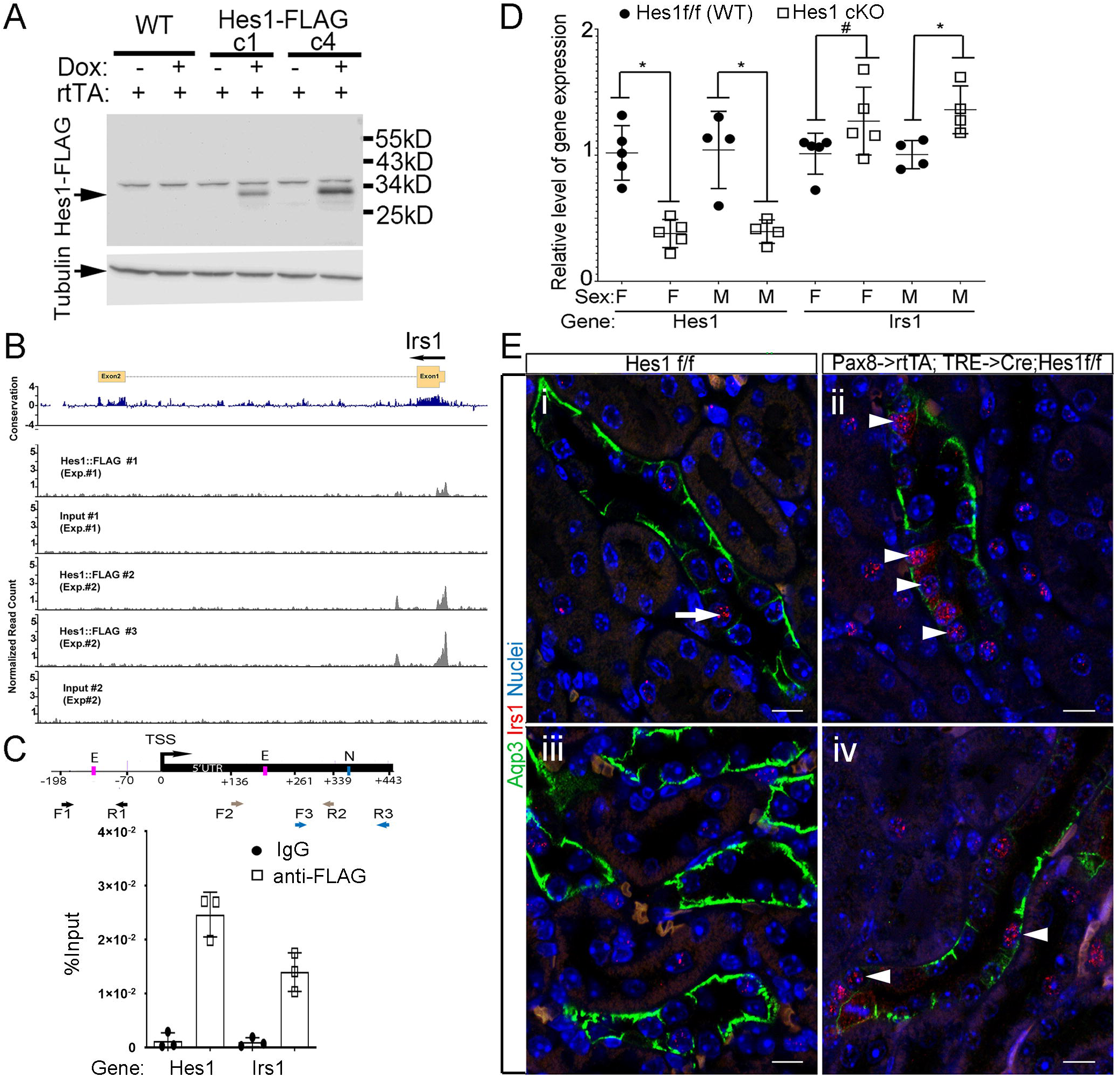
Irs1 is a direct target of Hes1 that is upregulated in principal cells following Hes1 inactivation. **A.** Western blot confirming doxycycline induced expression of Hes1 fused with 3xFLAG in two clones of mpkCCD_c14_ (top blot). Tubulin was detected as a loading control (bottom blot). **B.** Hes1::FLAG ChIP-seq analysis identified a peak near the transcriptional start site of Irs1 in three replicates when compared to inputs compiled from two experiments (Exp.#1 and Exp.#2). **C.** Schematic of the proximal regions flanking the transcription start site (TSS) of Irs1 revealing two E-box (pink) and one N-box (blue) DNA elements. Colored arrows depict three primer pairs used to perform ChIP-qPCR. ChIP-qPCR reveals significant enrichment of proximal promoter region of Hes1 and region +261 to +443 relative to the TSS of *Irs1* when using anti-FLAG versus IgG for ChIP. * denotes p<0.05, n=3 per condition, student t-test. **D.** RT-qPCR comparing gene expression levels in whole kidneys of *Pax8->rtTA; TRE->Cre; Hes1^f/f^*(Hes1cKO) versus control littermates (WT) that were given dox for 7 days. *Irs1* is up-regulated following inactivation of *Hes1* in the adult renal epithelial cell types. * p<0.05, # p=0.1, student t-test. **E.** Staining of Irs1 (red) and Aqp3 (green) in WT (i,iii) and Hes1cKO (ii,iv) mice reveals Irs1 protein is detectable in rare non-Aqp3+ cortical collecting duct cells in WT kidneys (arrow in i) and not in the outer medulla (iii), while Irs1 expression is detected in principal cells with reduced levels of Aqp3 expression in Hes1cKO kidneys (arrowheads) in the cortex (ii) and outer medulla (iv). Scale bars-10µm.

We next confirmed that renal *Irs1* mRNA levels increase in the kidneys one week after initiation of *Hes1* inactivation compared with control littermates by RT-qPCR (Fig. 2D). This increase is significant in the males and trending upwards in females with Hes1-deficiency. To determine the cell types in which *Irs1* expression increases we performed both RNAScope (data not shown) and Irs1 immunohistochemistry 4 and 7 days after initiation of *Hes1* inactivation. In wild type kidneys Irs1 is expressed in rare ICs in the cortical collecting ducts and not in PCs (Fig. 2E), consistent with type B ICs having highest level of Irs1 expression as previously determined by scRNA-seq (Fig. 1D) ^37^. In contrast, the Irs1 protein is detectable in PCs in cortical and medullary regions of Hes1cKO kidneys within 7 days of initiation of *Hes1* inactivation (Fig. 2E). Some of the PCs expressing Irs1 in the Hes1cKO kidneys have barely detectable expression of Aqp3 (Fig. 2E). These results support the idea that Hes1 directly represses *Irs1* expression in PCs of the adult mouse kidneys.

### mTORC1 activity increases in principal cells following inactivation of *Hes1*

Considering that components upstream of mTORC1, such as Irs1, are upregulated following *Hes1* inactivation and that Hes1 visits regions near TSS of some of these upregulated genes, we reasoned that the mTORC1 activity could be regulated by Notch signaling within the collecting ducts to regulate cell type composition. Consistent with this hypothesis phosphorylated S6 (pS6), a readout of mTORC1 activity, is predominantly absent in all PCs located in the different cortical and medullary collecting duct segments of mouse adult kidneys (Fig.3A,). While many nephron segments of the wild type kidneys have high levels of pS6, only Aqp2-negative cells have detectable pS6 within the collecting duct segments (Fig. 3A&B). However, within one week following initiation of *Hes1* inactivation the number of cells with pS6 is increased in collecting duct segments including in Aqp2 expressing cells (Fig. 3C&D). The detection of pS6 within PCs one week after *Hes1* inactivation, at a time point when PCs are converting to ICs ^7^, is consistent with Hes1 suppressing mTORC1 activity to maintain mature PC state.

**Figure 3.**
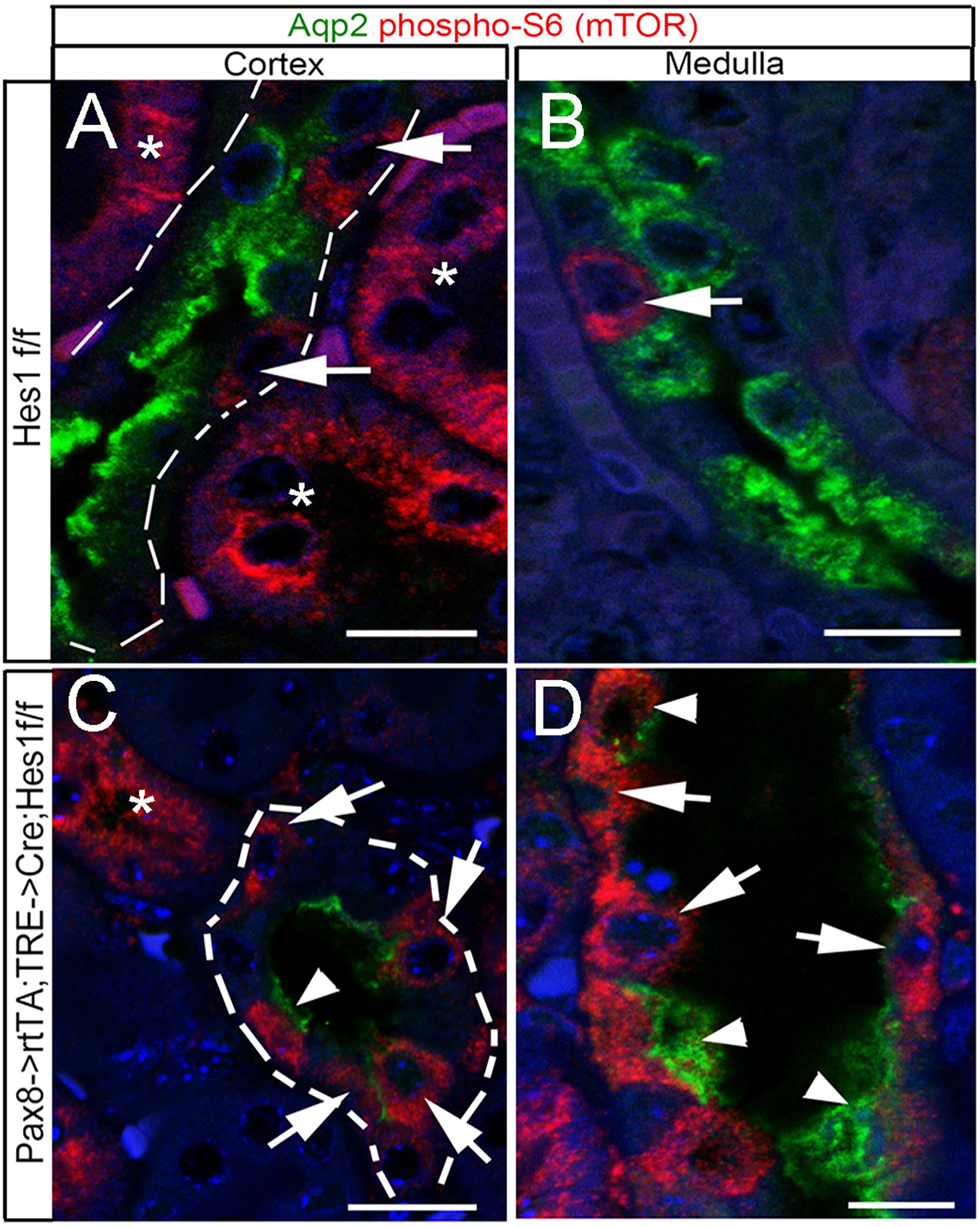
Staining for phospho-S6 reveals an increase in mTOR complex1 activity in principal cells after *Hes1* inactivation. **A & B.** Staining for Aqp2 and p-S6 in wild type cortical and medullary kidney sections reveals phospo-S6 (p-S6) staining in non-Aqp2-expressing ICs (arrows) within collecting duct segments and in cells of non-collecting duct segments (*). **C & D.** In Hes1-deficient (*Pax8-rtTA; TRE->Cre; Hes1^f/f^*) mouse kidneys, p-S6 is detected in many collecting duct cells (arrows) including Aqp2-expressing cells (arrowheads). Scale bars-10µm.

### *Tsc2* inactivation in the developing collecting ducts increases mTORC1 activity in principal cells and results in an increase in intercalated cells

To test whether increasing mTORC1 is sufficient to alter PC state towards an IC state, floxed alleles of *tuberous sclerosis complex 2* (*Tsc2*) were inactivated in the developing collecting ducts and distal nephron segments using the Cdh16-Cre transgenic mouse line. We confirmed that Tsc2 inhibits mTORC1 activity in the collecting ducts as we observed an increase in pS6 in the kidney collecting duct segments of 3-week-old *Cdh16-Cre; Tsc2f/f* mice (Fig. 4). While pS6 is not detectable in PCs of wild type littermate kidneys it is readily detected in PCs of *Cdh16-Cre; Tsc2f/f* mouse kidneys at 3-weeks of age (Fig. 4B&D). Along with increased mTORC1 activity in collecting duct segments there is a decrease in the number of Aqp2 and Aqp3 expressing PCs and an increase in the number of Foxi1 and Anion Exchanger 1 (AE1) expressing ICs in *Cdh16-Cre; Tsc2f/f* compared with wild type control littermate kidneys (Fig. 4E to H). Interestingly, in the cortical region where pendrin expressing type B and AE1 expressing type A ICs are present in wild type kidneys, there is only an increase in type A and not type B ICs in *Cdh16-Cre; Tsc2f/f* kidneys. Similarly, while there is increased mRNA levels of type A IC specific gene *Slc4a1* which codes for AE1 along with *Foxi1* which is expressed in all ICs there is a decrease in the mRNA levels of type B specific genes *Slc26a4* which codes for pendrin and *Slc4a9* which codes for AE4 in the *Tsc2*-deficient kidneys (Fig.4I & J). Consistent with reduced number of PCs, the mRNA levels of *Aqp2* and *Aqp3* are reduced in kidneys of 3-week-old *Cdh16-Cre; Tsc2f/f* mice compared with that of wild type littermates in female (Fig.4I), while there is a reduction in *Aqp3* mRNA levels and variable levels of *Aqp2* mRNA levels in the kidneys of male *Cdh16-Cre; Tsc2f/f* mice (Fig.4J). While the inactivation of *Tsc2* in the developing collecting ducts does increase pS6 in some PCs by post-natal day 0 (Supplementary Fig. S2) it does not prevent the selection of the PC fate as evidenced by no reduction in the number of Aqp2 expressing cells and no decrease in *Aqp2* and *Aqp4* gene expression when compared with that of wild type littermates (Supplementary Fig. S3). Additionally, there is not an increase in the number of Foxi1 expressing cells or the level of *Foxi1* gene expression in P0 *Cdh16-Cre; Tsc2f/f* kidneys compared with wild type littermates (Supplementary Fig. S3). Overall, these observations are consistent with mTORC1 activity promoting type A ICs at the expense of PCs and type B ICs in post-developmental stage mouse kidneys.

**Figure 4.**
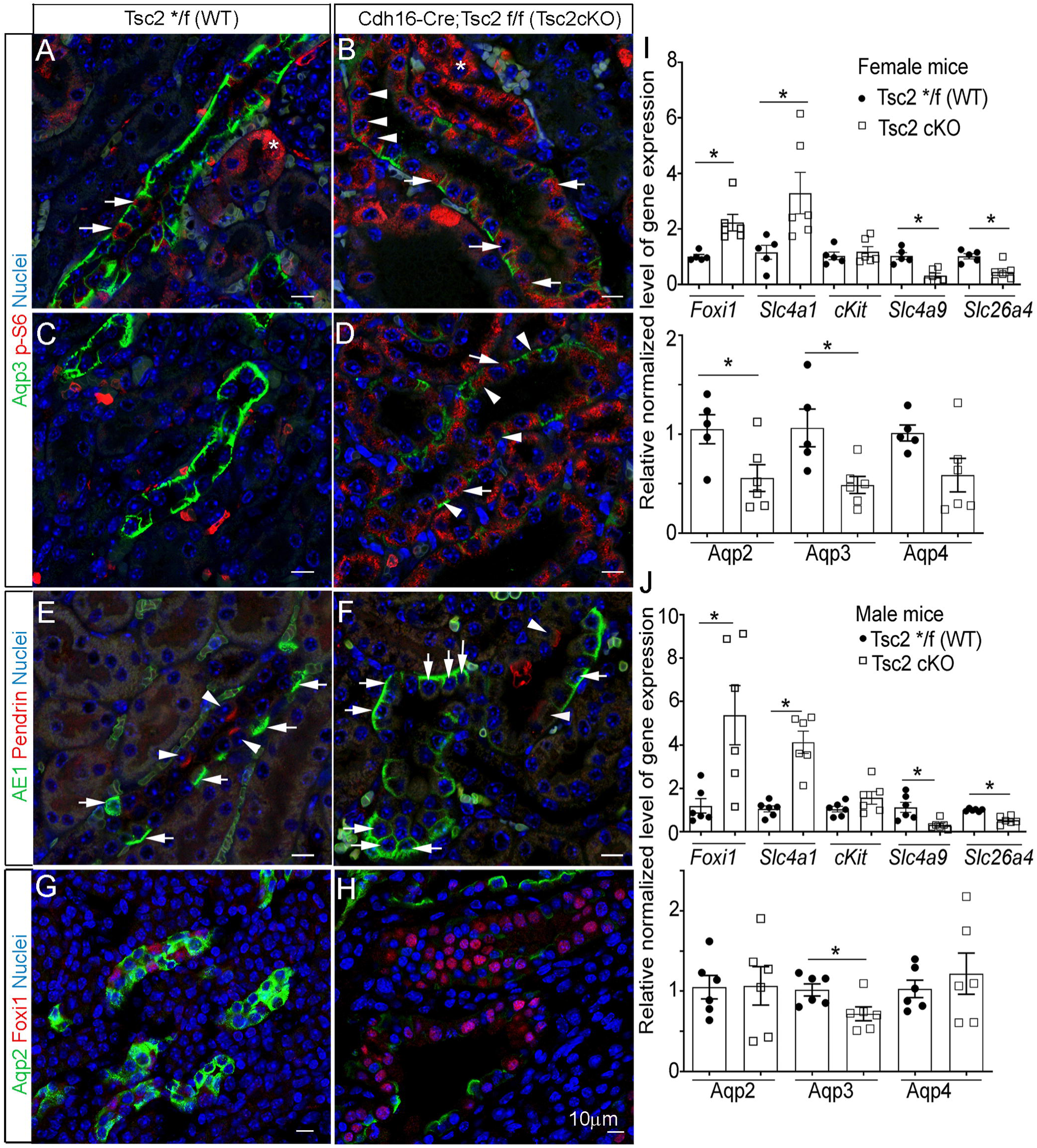
*Tsc2* is required for restricting the number of type A intercalated cells. **A-H.** Staining for IC, PC and mTORC1 activity markers in adult mouse kidneys of wild type mice (A, C, E, G) and Tsc2 cKO littermates (B, D, F, H). **A-D.** A marker of mTORC1 activity, p-S6 (red), is restricted to non-Aqp3 expressing cells within the cortical and medullary collecting duct segments (arrows) and in cells of non-collecting duct segments (asterisks) of wild type mice (A, C). Inactivation of *Tsc2* results in expression of p-S6 in Aqp3-expressing PCs (B, D arrowheads). **E, F.** Staining for AE1 a type A IC marker (arrows) and Pendrin a type B IC marker (arrowheads) reveals an increase in type A and a decrease in type B ICs in the Tsc2 cKO mouse cortical collecting ducts. **G, H.** Staining for Foxi1 an IC marker and Aqp2 a PC marker reveals an increase in the number of ICs in Tsc2 cKO mouse kidneys. Scale bars-10µm. **I, J.** RT-qPCR using RNA extracted from 3-week-old (I) female and (J) male mouse kidneys, Wt (n= 5 or 6 per sex) versus Tsc2 cKO (n = 6 per sex). There is an increase in *Foxi1* and *Slc4a1* (type A IC marker) expression in*Tsc2 c*KO mice, consistent with an increase in the number of type A ICs. Consistent with a decrease in type B ICs and in PCs there is a decrease in expression of *Slc4a9* and *Slc26a4* (Type B IC) and in some PC genes. Asterisks denote p<0.05, student t-test.

### Induction of mTORC1 in PCs of adult mice triggers PCs to express IC specific genes

To further determine if maintenance of mature PC state requires suppression of mTORC1 activity we initiated inactivation of *Tsc1* and *Tsc2* using *Elf5->rtTA;TRE->Cre* transgenic mice which targets the PC lineage. We have previously determined that doxycycline treatment of *Elf5->rtTA; TRE->Cre; Rosa^+/tdtomato^* adult mice from 3 to 5 weeks of age and analysis at 6 weeks of age revealed that a majority of the genetically labeled medullary CD cells, visualized by detection of tdtomato, also expressed PC markers while less than 0.2% expressed IC markers ^7^. We repeated similar lineage tracing of tdtomato-labeled PCs with wild type alleles of *Tsc1* and *Tsc2*. Analysis of *Elf5->rtTA;TRE->Cre; Tsc1^+/+^; Tsc2^+/+^; Rosa^+/tdtomato^* mice treated with doxycycline from 3 to 5 weeks of age and analyzed between 6 & 8 weeks of age revealed that none of the tdtomato labeled medullary CD cells express IC markers (Fig. 5), which is consistent with previous findings that Elf5->rtTA faithfully labels PCs and that a majority of labeled-PCs in the medulla do not gain IC properties within a short duration. In stark contrast, lineage tracing of tdtomato-labeled-PCs deficient in Tsc1 and Tsc2 using *Elf5->rtTA; TRE->Cre; Tsc1 ^+/*^; Tsc2 ^f/f^; Rosa^+/tdtomato^* mice (* denotes + or f) that were given doxycycline from 3 to 5 weeks of age revealed that on average 14% of tdtomato-labeled-medullary CD cells begin expressing IC markers, such as c-kit and pendrin between 6 & 8 weeks of age (Fig. 5). These observations are consistent with Tsc1 and Tsc2 restricting the level of mTORC1 activity in PCs, and that increasing mTORC1 activity drives PCs towards an IC state.

**Figure 5.**
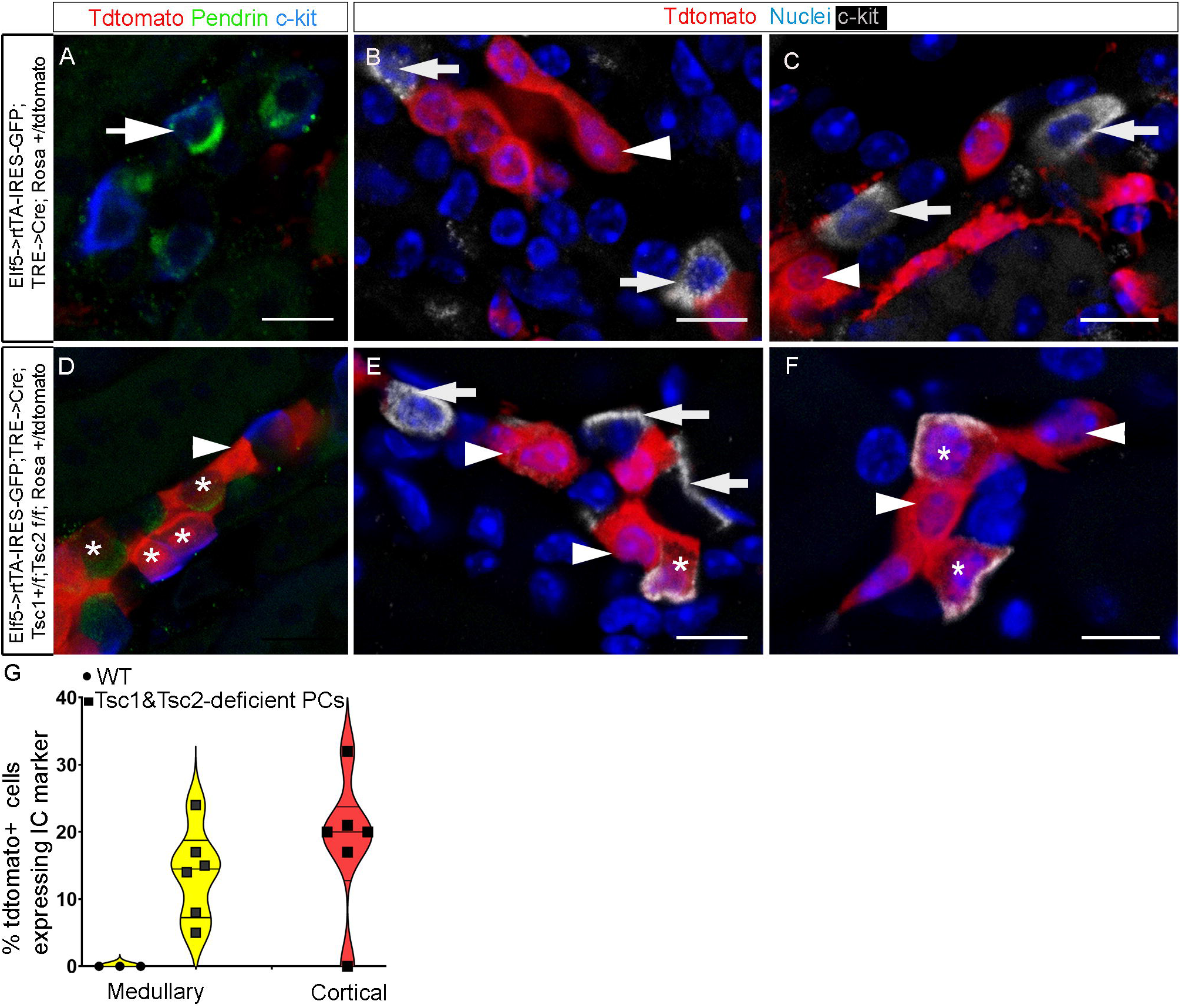
Lineage tracing of wild type versus Tsc1 and Tsc2 deficient PCs reveals that *Tsc1* and *Tsc2* suppress the expression of IC specific genes in mature PCs. A-F. Wild type (Wt) *Elf5->rtTA-IRES-GFP; TRE->Cre; Tsc1 +/+; Tsc2 +/+; Rosa +/tdtomato (A-C)* and *Elf5->rtTA-IRES-GFP; TRE->cre; Tsc1 +/*; Tsc2 f/f; Rosa +/tdtomato* mice (D-F, * denotes f or +) were given doxycycline from 3 to 5 weeks of age to genetically label Elf5 expressing principal cells (PCs) in order to compare the fate of PCs with or without inactivation of alleles of *Tsc1* and *Tsc2*. Staining for c-Kit and Pendrin, makers of ICs, revealed that genetically labeled (tdtomato+) wild type PCs which were detected in the medullary collecting ducts (arrowheads in B and C) and not in the cortical collecting ducts (A) never expressed ICs markers. IC marker expressing cells in wild type kidneys (arrows in A, B and C) were never from the genetically labeled PC lineage. In contrast genetically labeled (tdtomato+) PCs deficient for Tsc1 and Tsc2 begin expressing ICs markers (asterisks in D, E and F) both in cortical (D) and medullary (E and F) collecting duct segments. **G.** Violin plot of the percentage of tdtomato+ cells expressing IC markers in the cortex and medulla of wild type versus Tsc1 and Tsc2-deficient PCs. Approximately 18% of labeled-PCs in the cortex and 14% of labeled-PCs in the medulla that are deficient for Tsc1 and Tsc2 begin expressing IC markers. A minimum of 10 cortical collecting duct segments and 20 medullary collecting duct segments per mouse were analyzed for cells expressing tdtomato^+^ and c-Kit. A total of three mice with wild type PCs were compared with six mice with Tsc1 and Tsc2-deficient PCs. Scale bars-10µm.

### Raptor, a component of the mTORC1, is necessary for maintenance of intercalated cells

Considering that increasing mTORC1 activity by inactivation of *Tsc2* is sufficient to induce PCs towards an IC state we next tested the requirement of mTORC1 in maintenance of ICs in adult kidneys. A previous study reported that while Raptor is critical for mitochondrial biogenesis during kidney development, conditional inactivation of Raptor in the adult kidney epithelia resulted in no obvious phenotype in the short-term but altered the response to acute kidney injury ^39^. We conditionally inactivated Raptor, a component of the mTORC1, in the adult kidney epithelial cells by providing *Pax8->rtTA;TRE-Cre; Rptor f/f* (Raptor cKO) mice with doxycycline from 3 to 5 weeks of age and analyzed the kidney at 8 weeks of age. Along with reduced mTORC1 activity as determined by pS6 staining (Fig. 6 A&B), the Raptor cKO kidneys had significantly reduced mRNA levels of IC specific genes including *Foxi1, Slc4a1*, *Slc4a9* and *Slc26a4* when compared with wild type controls as determined by RT-qPCR (Fig. 6I & J). Among the IC specific genes examined the *c-Kit* mRNA level is not significantly different between Raptor cKO and control kidneys. However, *c-Kit* mRNA levels trend downwards in the Raptor cKO kidneys, and this is further supported by absence or reduced staining of c-Kit in the ICs of medullary collecting duct segments of Raptor cKO kidneys (Fig.6D). Additional IC markers Foxi1 and Pendrin have severely reduced expression in ICs of Raptor cKO mouse kidneys (Fig. 6E-H and Supplementary Fig.S4). In contrast to IC gene expression, the PC gene expression is not altered in female Raptor cKO kidneys but is reduced in male Raptor cKO kidney compared with sex matched control kidneys. (Fig. 6I & J). However, by staining we were not able to detect a change in protein levels of PC makers in male or female Raptor cKO kidneys compared with control kidneys (Fig. 6A-H). Consistent with intact PCs the urine osmolality was not significantly different between Raptor cKO and controls (Fig.6K). Interestingly, the urine pH was lower in Raptor cKO mice versus control littermates (Fig. 6K). These observations reveal a critical requirement of mTORC1 in maintenance of ICs in a mature state by ensuring expression of IC specific genes.

**Figure 6.**
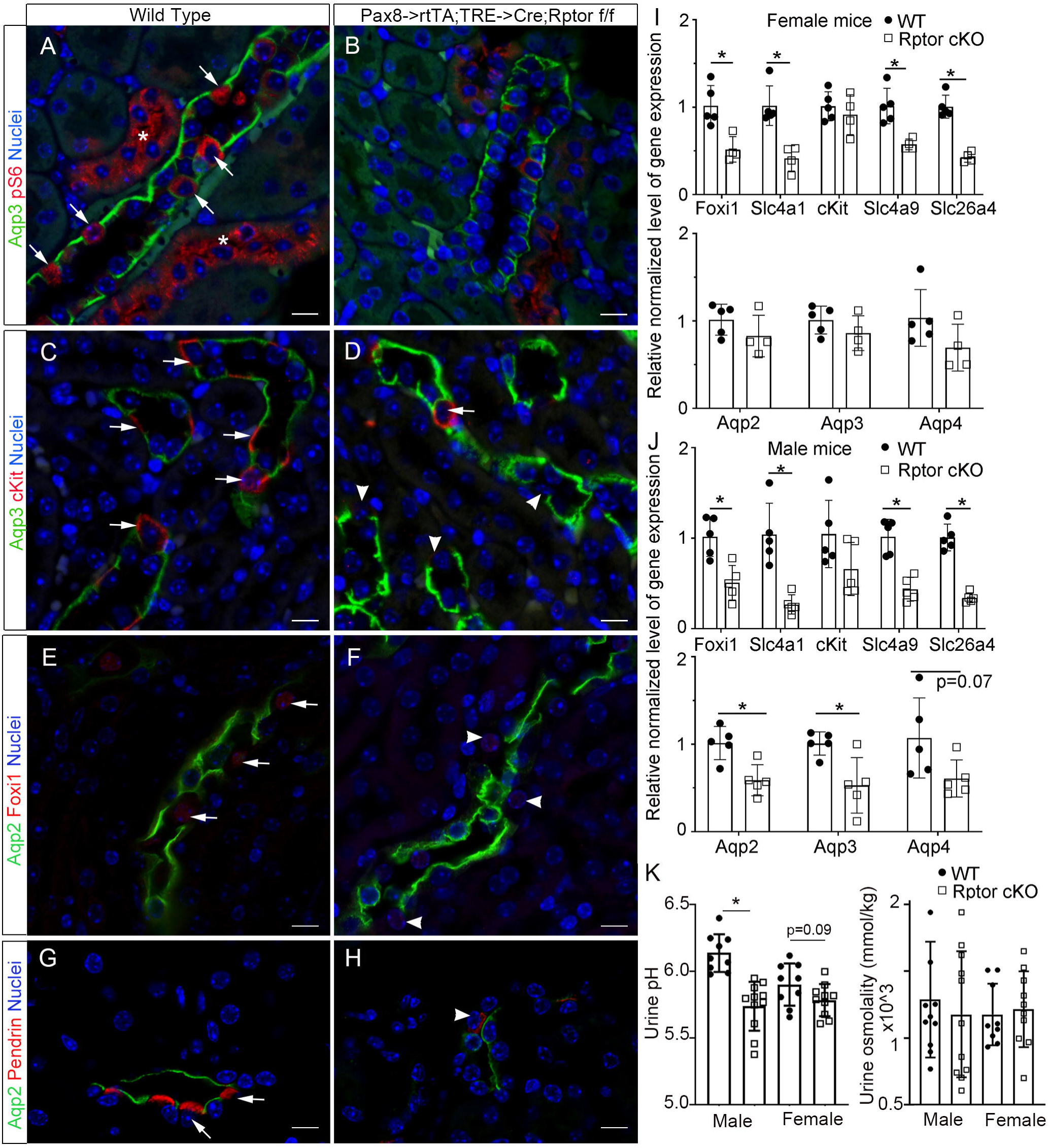
Raptor, a component of mTORC1, is required for maintenance of mature intercalated cell types. **A-H.** Staining for IC, PC and mTORC1 activity markers in adult mouse kidneys of wild type mice (A, C, E, G) and Raptor cKO mice (B, D, F, H). **A&B.** A marker of mTORC1 activity, p-S6 (red), is restricted to non-Aqp3 expressing cells within the collecting duct segments (arrows) and in cells of non-collecting duct segments (asterisks) of wild type mice (A). Inactivation of *Rptor* results in reduced p-S6 in epithelial cells (B). **C& D.** c-Kit expression is reduced or absent in ICs of the Raptor cKO mouse medullary collecting ducts (arrowheads in D). **E&F.** Foxi1 expression is reduced in many ICs in Raptor cKO mouse kidneys (arrowheads in F). **G& H**. Pendrin expression is reduced in ICs of Raptor cKO mouse kidneys (arrowhead in H). Scale bars-10µm. **I, J.** RT-qPCR using RNA extracted from 8-week-old (I) female and (J) male mouse kidneys, Wt (n= 5 per sex) versus Raptor cKO (n = 4 or 5 per sex). **K.** Urine pH is reduced in Raptor cKO mice while urine osmolality is not reduced. Asterisks denote p<0.05, student t-test.

## Discussion

Our observations reveal a role for Notch signaling via Hes1 in regulating the level of mTORC1 activity within the collecting duct segments. Specifically, inactivation of *Hes1* in the adult mouse kidney epithelia regulates the expression of Irs1 in PCs. Irs1 is a direct transcriptional target of Hes1 in cultured mature PCs, and Hes1 repression of Irs1 restricts Irs1 expression to ICs within the adult mouse kidney collecting duct segments. The observed increase in mTORC1 activity in PCs along with Irs1 within a week after initiation of *Hes1* inactivation in adult kidney epithelia is consistent with Irs1 being an upstream mediator of mTORC1 activity ^40^. These observations along with the known requirement of Hes1 in preventing the conversion of mature PCs into ICs raised the hypothesis that mTORC1 activity is suppressed by Notch signaling in PCs to maintain mature PC state.

We firmly establish a critical role for mTORC1 activity in regulating the cell type composition within the adult kidney collecting duct cells. Interestingly, neonatal *Cdh16-Cre; Tsc2f/f* mouse kidneys contained normal composition of PC and IC types and did not show a pervasive increase in mTORC1 activity. However, by 3 weeks of age increased mTORC1 activity was observed along with a reduction in number of PCs and an increase in ICs. This is suggestive that while initial differentiation of PCs is insensitive to *Tsc2* inactivation the maintenance of mature PCs requires suppression of mTORC1 activity. These results are consistent with previous reports that inactivation of *Tsc1* in mature PCs using *Aqp2-Cre* results in increased ICs ^11^. Here we further determined that increasing mTORC1 in genetically labeled PCs in the adult kidney using the *Elf5->rtTA;TRE-Cre*; Rosa^+/tdtomato^ PC lineage mice is sufficient to induce expression of mature IC markers in PCs. This observation reveals that PC state is incompatible with high levels of mTORC1 activity which promotes IC state. Increasing mTORC1 activity forced PCs to begin converting towards IC state including type B and type A ICs. While increasing mTORC1 in PCs forces them to turn on type A or type B IC markers in *Elf5->rtTA;TRE-Cre*; Tsc2^f/f^; Tsc1^+/f^; Rosa^+/tdtomato^ kidneys, having a sustained increase in mTORC1 as in the *Cdh16-Cre; Tsc2f/f* kidneys resulted in decreased type B ICs along with PCs and favored an increase in type A ICs.

Inactivation of *Rptor* in adult kidney epithelia revealed the requirement of mTORC1 for sustaining the expression of IC specific genes such as *Foxi1*, *Slc4a1* (AE1) and *Slc26a4* (Pendrin). The requirement of mTORC1 is not unique to ICs considering that expression of PC genes such as *Aqp2*, and *Aqp3* are reduced at the mRNA level following inactivation of *Rptor* in the adult kidney epithelia of male mice. These observations are consistent with the idea that the level of mTORC1 activity dictates the identity of the collecting duct cells. Considering that mTORC1 integrates environmental inputs to coordinate cell metabolism ^41^, this study reveals that neighboring epithelial cells within a tubular segment can be regulated to take on different identities by differential endowment to transduce metabolic cues. The expression of Irs1 in ICs and not in PCs in wild type kidneys and the induction of Irs1 in PCs following inactivation of *Hes1* raises the possibility that receptors whose activities are transduced by Irs1 are in part responsible for higher levels of mTORC1 activity in ICs and may mediate PC to IC conversions.

In the context of collecting ducts the insulin receptor-related receptor (Insrr), which belongs to the receptor tyrosine kinase (RTK) family and is expressed predominantly in type B ICs, could be one receptor that activates mTORC1 in an Irs1 dependent manner (Fig. 7). Interestingly, Insrr can be activated by alkaline pH and is required for normal response to an alkaline diet in rodents ^42^. Recent transcriptomic analysis of Insrr-null mouse kidneys revealed decreased expression of pendrin along with differential expression of genes associated with ATP metabolism and electron transport chain ^43^, suggestive that Insrr modulates metabolism to maintain type B ICs in a mature functional state. Although our study clearly identifies Irs1 as directly repressed by Hes1 in PCs, it remains to be determined if up regulation of Irs1 in PCs is sufficient to drive them towards an IC state. It is likely that Insrr, which is up regulated following *Hes1* inactivation, or other receptors up stream of mTORC1 are also required to be turned on in PCs for them convert to ICs.

**Figure 7.**
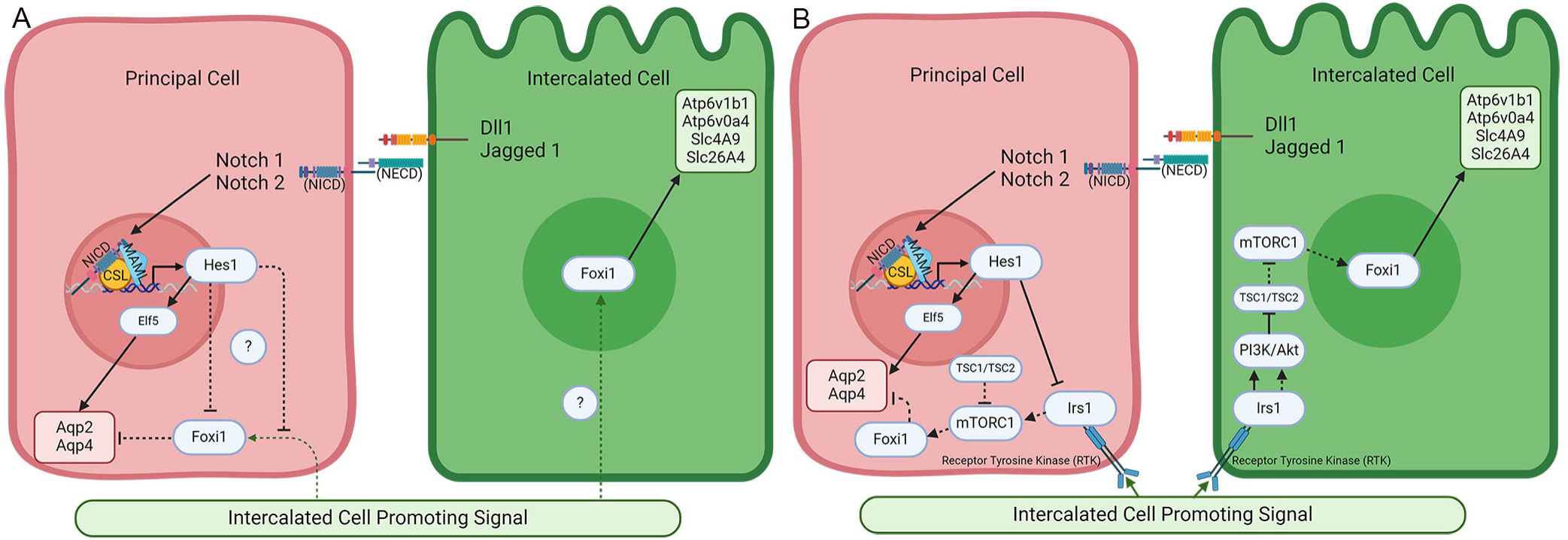
The objective of this study is to understand how Notch signaling via Hes1 suppresses the intercalated cell program from turning on in principal cells of adult kidney collecting ducts. **A.** Activation of Notch1 and Notch2 receptors in principal cells turns on Hes1 to suppress the expression of Foxi1, a critical intercalated cell specific transcription factor. Although, the primary function of Notch signaling via Hes1 is to suppress the expression of Foxi1, the mechanism mediating suppression of Foxi1 is unknown. A possibility is that Notch signaling prevents the reception and/or transduction of a hypothetical intercalated cell promoting signal. **B.** We identified that Hes1 suppresses expression of Irs1 and mTORC1 activity in principal cells and increasing mTORC1 activity by genetic inactivation of *Tsc2* promotes the intercalated program in principal cells. Additionally, inactivation of a component in mTORC1 suppressed the intercalated cell state.

While our study focused on the mechanisms by which Notch regulates plasticity of mature epithelial cell types in the kidney collecting ducts our findings may be of relevance in other adult tissues such as the lung where inhibition of Notch signaling triggers the direct conversion of club cells to multiciliated cells^44^. In both the airway epithelium and kidney collecting duct epithelium Notch signaling regulates cell fate specification during development and maintenance of distinct mature cell types in the adult. It remains to be determined if Notch regulation of mTORC1 is a conserved mechanism that governs cell type plasticity. This linking of the level mTORC1 activity to distinct functional cell types may endow adult tissues with the capacity to remodel their constituent cell types as part of adaptive responses to sustained physiologic stressors.

## Acknowledgements

Research reported here was supported by grants from NIDDK of NIH under award numbers R01DK106135, and R56DK106135. K.S. was supported by R01DK123180. We thank the Imaging and Histology Core, and Functional Genomics and Bioinformatics cores at Sanford Research for technical assistance. These Cores at Sanford Research were supported by NIH grants P30GM145398 and P20GM103620.

## Author Contributions

J. DeRiso, M. Mukherjee, M. Janga, A. Simmons, performed experiments, and analyzed the results; J. DeRiso A. Simmons, M. Mukherjee and M. Janga contributed to writing the methods sections; M. Kareta assisted with analysis of ChIP sequencing results; J. Tao contributed reagents for mTORC1 genetic modulation and critical discussion of results; I. Chandrasekar contributed to imaging experiments and critical discussion of results; K. Surendran designed experiments, helped analyze the results and wrote the manuscript; and all authors contributed to manuscript editing and revisions.

