## Supplementary Fig. S1 to S4 and Table1 and Table 2 for "Kidney collecting duct cell type composition is regulated by Notch signaling via modulation of mTORC1"

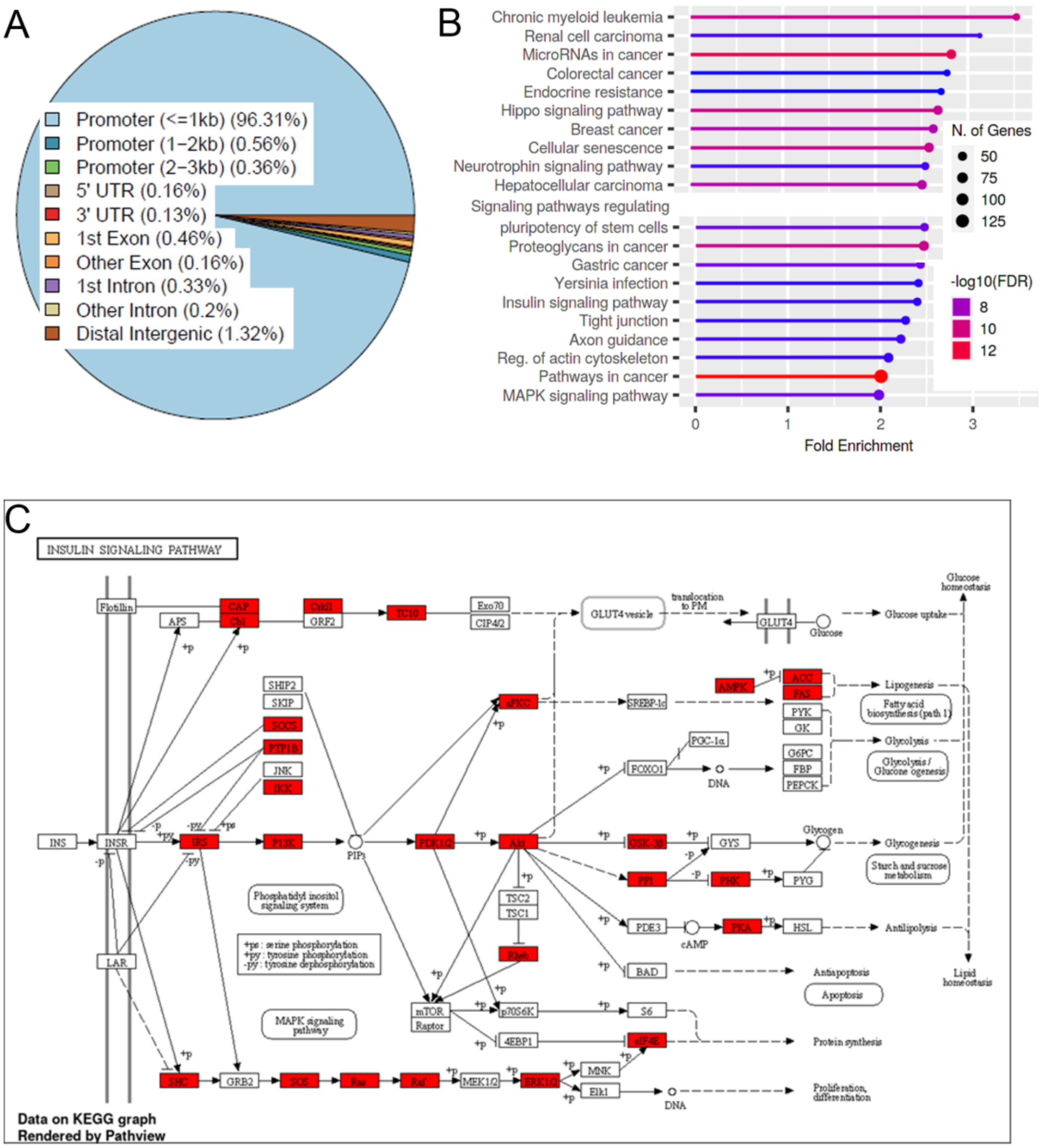


**Supplementary Figure S1. Hes1-FLAG ChIP-seq in mpkCCD$_{c14}$, a mature principal cell line, reveals enrichment of genes coding for components in the PI3K-Akt/insulin signaling pathway upstream of mTORC1.** A. Pie chart illustrating that most of the genomic locations visited by Hes1 are within 1kb of transcriptional start sites. B. Ingenuity Pathway Analysis of genes associated with ChIP-seq peaks reveals significantly enriched pathways, diseases and cellular functions that may be potential targets of Hes1 regulation. C. Insulin signaling pathway genes that were associated with the Hes1 ChIP-seq peaks are boxed in red and include irs1 and other components upstream of mTORC1.

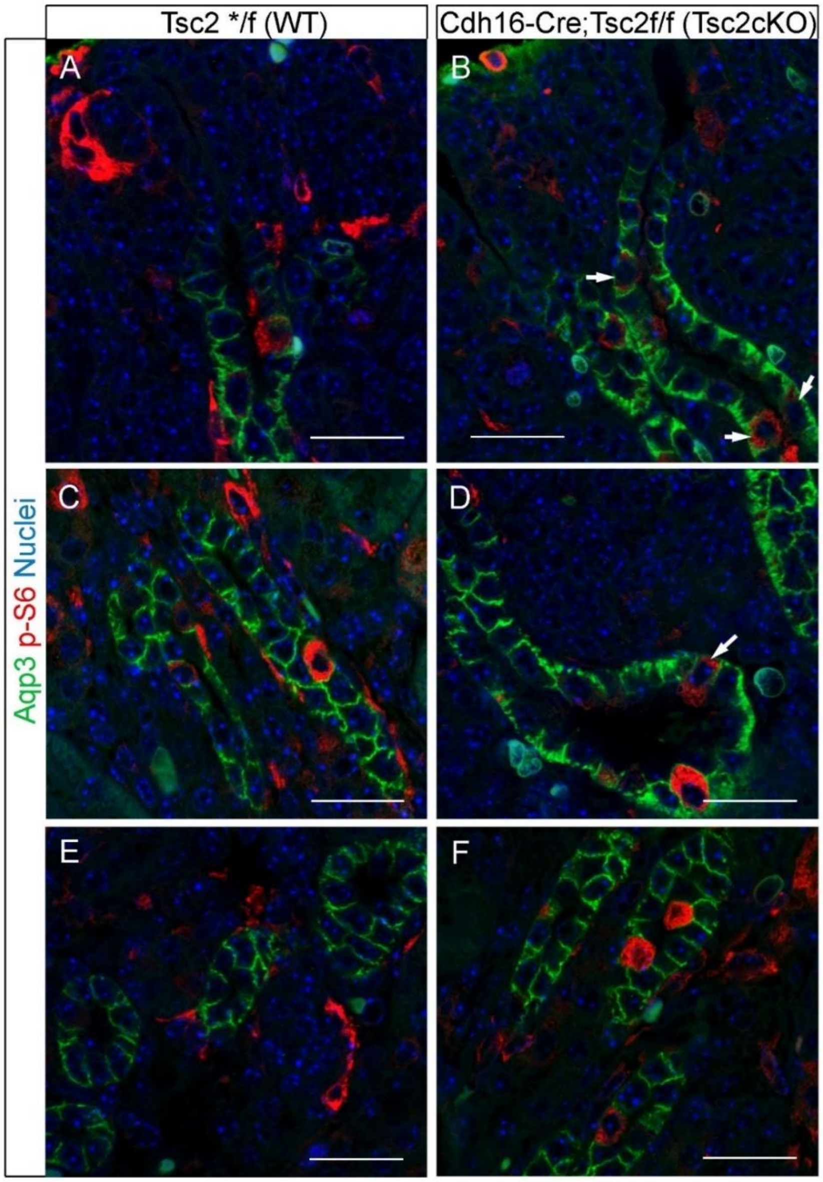


**Supplementary Figure S2. Inactivation of *Tsc2* in the developing kidney collecting ducts increases phospho-S6 in some PCs by post-natal day 0 but does not prevent the selection of the PC fate.** Wild type (WT) control littermates kidney sections (A, C & E) and *Cdh16-Cre;Tsc2f/*f (Tsc2ckO) kidneys sections (B, E & F) were stained with Aqp3 (green) and phospho-S6 (red). Nephrogenic zone containing collecting duct tips and stalk (A &B) reveal rare pS6+ duct cells in WT sections while there are more duct cells (arrows) that have pS6 in Tsc2cKO sections. Cortical collecting ducts (C & D) and medullary collecting ducts (E & F) also contain rare pS6+ cells. Scale bars = 25 μm

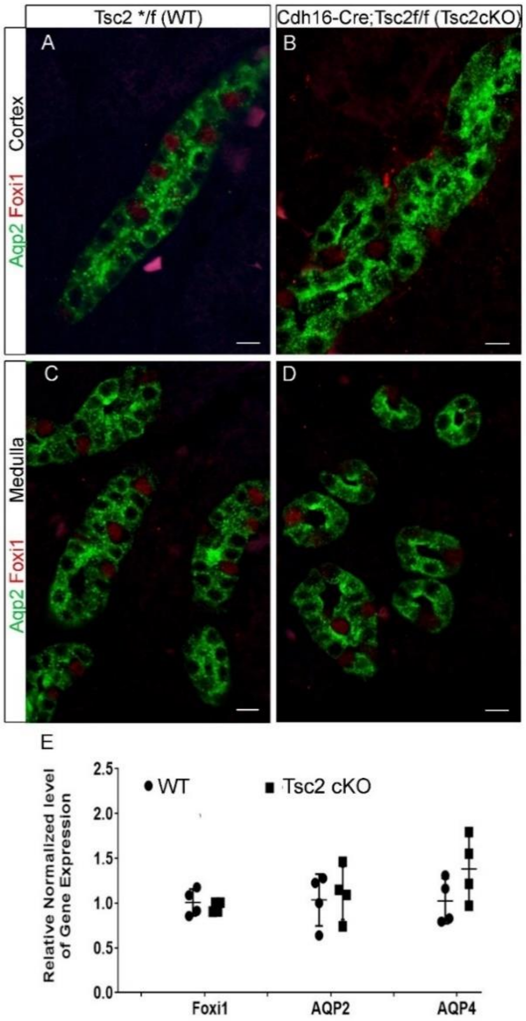


**Supplementary Figure S3. Inactivation of *Tsc2* in the developing kidney collecting ducts does not result in an increase in *Foxi1* expression in post-natal day 0 *Cdh16-Cre; Tsc2f/f* (Tsc2ckO) kidneys.** A-D. P0 kidney sections were stained for Foxi1 (red) and Aqp2 (green). Scale bars =10 $\mu$m. E. RT-qPCR using RNA extracted from P0 mouse kidneys. WT (n=4) kidneys were compared with Tsc2 cKO kidneys (n = 4). The expression levels of Foxi1, Aqp2 and Aqp4 were similar in WT and Tsc2cKO mouse kidneys.

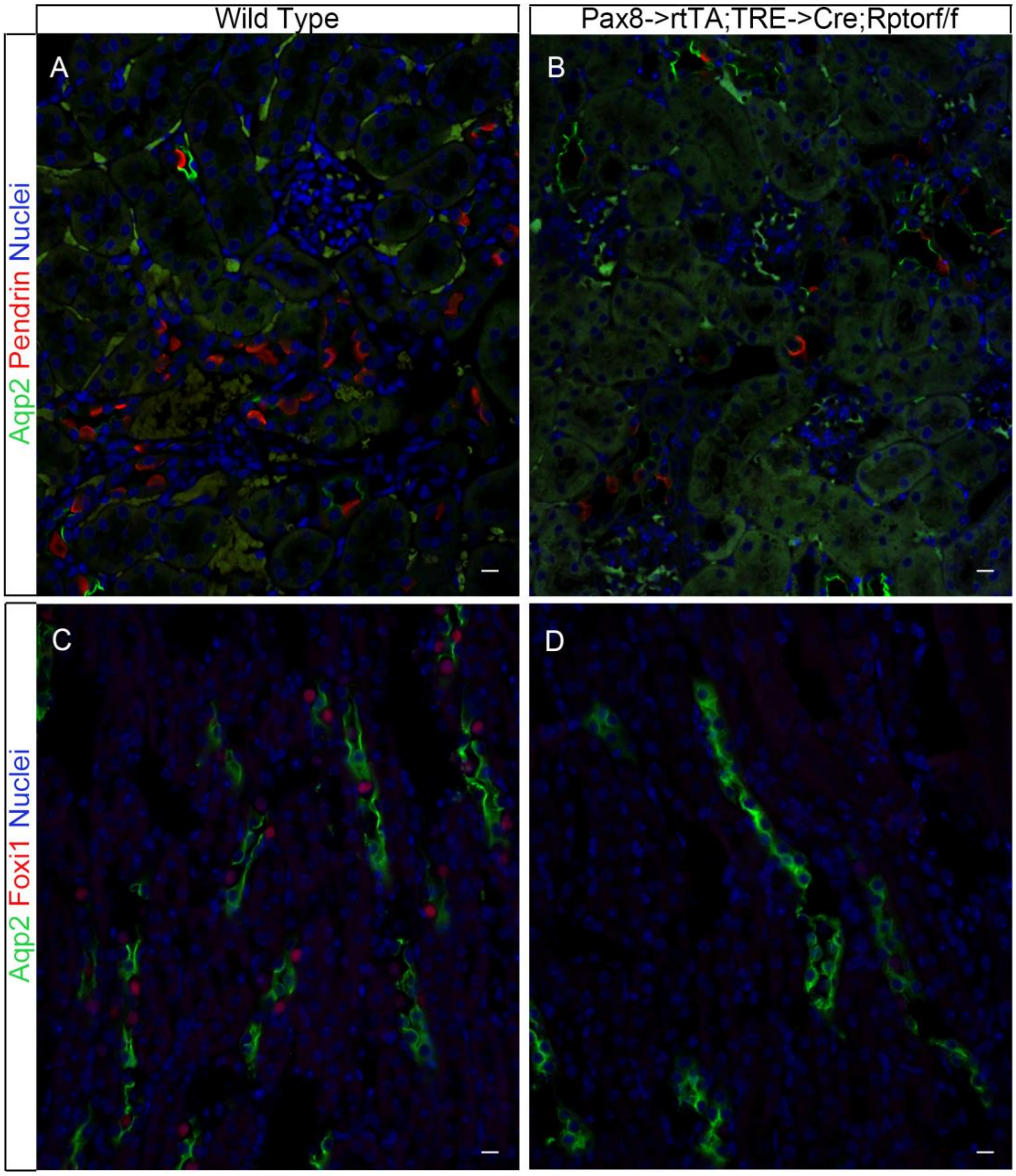


**Supplementary Figure S4. Raptor, a component of mTORC1, is required for maintenance of mature intercalated cell types**. Staining for IC and PC markers in adult mouse kidneys of wild type mice (A, C) and Raptor cKO mice (B, D) reveals a reduction in IC marker expression in Raptor cKO kidneys. **A & B**. Pendrin expression is reduced in ICs of Raptor cKO mouse kidneys. **C&D.** Foxi1 expression is reduced in Raptor cKO mouse kidneys. Scale bars-10µm.

**Supplementary Table 1:**

| Mouse referred to in manuscript | Official Name | Reference |
|---|---|---|
| Tet-O-Cre<br>Jax stock #006234 | B6.Cg-Tg(tetO-Cre)1Jaw/J | [1] |
| Elf5->rtTA-IRES-GFP | | [2] |
| Cdh16-> Cre<br>Jax stock #012237 | B6.Cg-Tg(Cdh16-cre)91Igr/J | [3] |
| Pax8->rtTA<br>Jax stock #007176 | B6.Cg-Tg(Pax8-rtTA2S*M2)1Koes/J | [4] |
| Hes1 $^{f/f}$ | | [5] |
| Tsc1 $^{f/f}$<br>Jax stock #005680 | Tsc1$^{tm1Djk}$/J | [6] |
| Tsc2 $^{f/f}$<br>Jax stock #027458 | Tsc2$^{tm1.1Mjg}$/J | [7] |
| Rosa $^{tdtomato}$<br>Jax stock #007909 | B6.Cg-Gt(ROSA)26Sor$^{tm9(CAG-tdTomato)Hze/J}$ | [8] |
| Raptor $^{f/f}$<br>Jax stock #013191 | B6.129S5-Rptor$^{tm1Lex}$/J | [9] |

**Supplementary Table 2:**

| Antibody Name | Company, Catalogue Number and/or Lot. Number | Dilution for IHC or amount used per ChIP | Reference for prior use or validation |
|---|---|---|---|
| Irs1 | Cell Signaling Technology, 2382 | 1:200 | |
| Phospho-S6 (Ser235/236) Ribosomal Protein | Cell Signaling Technology, 4858 | 1:200 | |
| Foxi1 | Abcam, Ab20454 | 1:100 | [10] |
| Aqp3 (C-18) | Santa Cruz, sc-9885 | 1:200 | |
| Pendrin (E-20) | Santa Cruz, sc-16894 | 1:100 | |
| Pendrin | BiCell, 20501 | 1:100 | |
| AE1 | Millipore, AB3500P | 1:200 | [11] |
| Aqp2 | Santa Cruz, sc-9882<br>Novus, NBP1-70378 | 1:200<br>1:200 | [12] |
| Aqp4 | Millipore, AB3594 | 1:200 | |
| c-Kit | Cell Signaling Technology, 3074 | 1:200 | [13] |
| α-Tubulin | Sigma-Aldrich, T9026 | 1:20,000 | |
| FLAG | Sigma-Aldrich, F1804 | 1:1000 | |
