## Supplementary Table 3 for "Kidney collecting duct cell type composition is regulated by Notch signaling via modulation of mTORC1"

| **Supplementary Table 3: Differentially expressed genes in whole adult mouse kidneys from** | | | | | |
| --- | --- | --- | --- | --- | --- |
| **9 wild type control littermates versus 7 Hes1 ckO (Pax8->rtTA;TRE->Cre; Hes1f/f).** | | | | | |
| All mice were given doxycycline in drinking water for 1 week prior to analysis. | | | | |  |
| **Ensemble_Id** | **Gene_name** | **DE_direction** | **Log Fold Change (LFC)** | **Adjusted p-value** | **Signaling Pathway** |
| ENSMUSG00000102004 | 4933402C06Rik | Down | -2.734590863 | 0.0058433 |  |
| ENSMUSG00000029608 | Rph3a | Down | -2.568324138 | 0.001237 |  |
| ENSMUSG00000105265 | Sox2ot | Down | -2.278958117 | 0.0407945 |  |
| ENSMUSG00000091813 | Ces2h | Down | -2.223865017 | 0.0056613 |  |
| ENSMUSG00000037418 | Best1 | Down | -1.920492358 | 0.036443 |  |
| ENSMUSG00000022002 | 4930564B18Rik | Down | -1.871557981 | 0.0442363 |  |
| ENSMUSG00000057074 | Ces1g | Down | -1.720005786 | 0.0203857 |  |
| ENSMUSG00000023328 | Ache | Down | -1.635747855 | 6.669E-09 |  |
| ENSMUSG00000046101 | Mcmdc2 | Down | -1.525356023 | 0.0486674 |  |
| ENSMUSG00000022508 | Bcl6 | Down | -1.46561501 | 0.035203 |  |
| ENSMUSG00000109532 | Gm4239 | Down | -1.428403358 | 0.0206313 |  |
| ENSMUSG00000071633 | Gm4952 | Down | -1.333751843 | 0.0090204 |  |
| ENSMUSG00000026725 | Tnn | Down | -1.209211417 | 0.0430761 |  |
| ENSMUSG00000078597 | Cyp4a12b | Down | -1.02756573 | 0.0381244 |  |
| **ENSMUSG00000022528** | **Hes1** | **Down** | **-0.984310426** | **5.638E-06** | **Notch** |
| ENSMUSG00000068303 | Spr-ps1 | Down | -0.978289266 | 0.0169728 |  |
| ENSMUSG00000015451 | C4a | Down | -0.952052651 | 0.0450022 |  |
| ENSMUSG00000023073 | Slc10a2 | Down | -0.893987948 | 0.0184725 |  |
| ENSMUSG00000051517 | Arhgef39 | Down | -0.857723039 | 0.0187788 |  |
| ENSMUSG00000039639 | Kcne1 | Down | -0.827639735 | 0.003874 |  |
| ENSMUSG00000020415 | Pttg1 | Down | -0.797191509 | 0.0061253 |  |
| ENSMUSG00000031891 | Hsd11b2 | Down | -0.766048036 | 6.576E-08 |  |
| ENSMUSG00000059602 | Syn3 | Down | -0.752031255 | 0.0032743 |  |
| ENSMUSG00000030498 | Gas2 | Down | -0.676186921 | 0.0056613 |  |
| ENSMUSG00000032815 | Fanca | Down | -0.65401088 | 0.0029965 |  |
| ENSMUSG00000025574 | Tk1 | Down | -0.645707397 | 0.0036763 |  |
| ENSMUSG00000047583 | Tyw3 | Down | -0.636082407 | 0.013448 |  |
| ENSMUSG00000004988 | Fxyd4 | Down | -0.61979937 | 0.0005059 |  |
| ENSMUSG00000038677 | Scube3 | Down | -0.562156859 | 0.0407945 |  |
| ENSMUSG00000063683 | Glyat | Down | -0.558190927 | 0.0026311 |  |
| ENSMUSG00000026348 | Acmsd | Down | -0.54455259 | 0.0058996 |  |
| ENSMUSG00000006585 | Cdt1 | Down | -0.509316319 | 0.0314129 |  |
| ENSMUSG00000030291 | Med21 | Down | -0.473908084 | 0.0098722 |  |
| ENSMUSG00000020774 | Aspa | Down | -0.470617551 | 2.241E-05 |  |
| ENSMUSG00000025395 | Prim1 | Down | -0.467700492 | 0.0161682 |  |
| ENSMUSG00000050730 | Arhgap42 | Down | -0.463285904 | 0.0486674 |  |
| ENSMUSG00000021306 | Gpr137b | Down | -0.414980511 | 0.0423792 |  |
| ENSMUSG00000047412 | Zbtb44 | Down | -0.400014561 | 0.0177079 |  |
| ENSMUSG00000079042 | Trim61 | Down | -0.395790498 | 0.0383677 |  |
| ENSMUSG00000031864 | Ints10 | Down | -0.387187674 | 0.0028855 |  |
| ENSMUSG00000027577 | Chrna4 | Down | -0.381770438 | 0.0246919 |  |
| ENSMUSG00000028030 | Tbck | Down | -0.377916321 | 0.001449 |  |
| ENSMUSG00000027605 | Acss2 | Down | -0.369508672 | 0.0246919 |  |
| ENSMUSG00000025194 | Abcc2 | Down | -0.365397419 | 0.0133186 |  |
| ENSMUSG00000074794 | Arrdc3 | Down | -0.36294066 | 0.0178193 |  |
| **ENSMUSG00000051177** | **Plcb1** | **Down** | **-0.346119794** | **0.0107159** | **Wnt** |
| ENSMUSG00000058690 | Ccser2 | Down | -0.343198606 | 0.0130132 |  |
| ENSMUSG00000022899 | Slc15a2 | Down | -0.34183185 | 0.0061253 |  |
| ENSMUSG00000038745 | Nlrp6 | Down | -0.340140974 | 0.0177599 |  |
| ENSMUSG00000029207 | Apbb2 | Down | -0.326539774 | 0.0251659 |  |
| ENSMUSG00000002032 | Tmem25 | Down | -0.305794049 | 0.0184725 |  |
| ENSMUSG00000058488 | Kl | Down | -0.300312702 | 0.0097682 |  |
| ENSMUSG00000024251 | Thada | Down | -0.294766748 | 0.0370578 |  |
| ENSMUSG00000075025 | Gm10804 | Down | -0.287598115 | 0.014809 |  |
| ENSMUSG00000062248 | Cks2 | Down | -0.265927806 | 0.045676 |  |
| ENSMUSG00000060961 | Slc4a4 | Down | -0.264837514 | 0.0496141 |  |
| ENSMUSG00000022132 | Cldn10 | Down | -0.25796008 | 0.0162469 |  |
| ENSMUSG00000029802 | Abcg2 | Down | -0.251879703 | 0.0308362 |  |
| ENSMUSG00000023150 | Ivns1abp | Down | -0.239547355 | 0.0489779 |  |
| ENSMUSG00000027679 | Dnajc19 | Down | -0.237308565 | 0.0182261 |  |
| ENSMUSG00000024924 | Vldlr | Down | -0.162900879 | 0.0247122 |  |
| ENSMUSG00000034108 | Ccs | UP | 0.188166784 | 0.0488935 |  |
| ENSMUSG00000007659 | Bcl2l1 | UP | 0.191730933 | 0.02987 |  |
| ENSMUSG00000038178 | Slc43a2 | UP | 0.19371735 | 0.0162469 |  |
| ENSMUSG00000019087 | Atp6ap1 | UP | 0.196863925 | 0.0085404 |  |
| ENSMUSG00000024942 | Capn1 | UP | 0.215199796 | 0.0130966 |  |
| ENSMUSG00000038712 | Fam63a | UP | 0.216155782 | 0.0075057 |  |
| ENSMUSG00000032220 | Myo1e | UP | 0.238636317 | 0.0107159 |  |
| ENSMUSG00000048878 | Hexim1 | UP | 0.244596285 | 0.001237 |  |
| ENSMUSG00000021665 | Hexb | UP | 0.245958086 | 0.0168585 |  |
| ENSMUSG00000028833 | Ncdn | UP | 0.251050173 | 0.0308362 |  |
| ENSMUSG00000001995 | Sipa1l2 | UP | 0.255918427 | 0.0418911 |  |
| ENSMUSG00000001089 | Luzp1 | UP | 0.264595544 | 0.0108635 |  |
| ENSMUSG00000021948 | Prkcd | UP | 0.265189884 | 0.0004249 |  |
| ENSMUSG00000017765 | Slc12a4 | UP | 0.273217818 | 0.0177079 |  |
| ENSMUSG00000073139 | BC023829 | UP | 0.275105304 | 0.0102169 |  |
| ENSMUSG00000004040 | Stat3 | UP | 0.280889531 | 0.023312 |  |
| ENSMUSG00000017132 | Cyth1 | UP | 0.296358062 | 0.0308922 |  |
| ENSMUSG00000022895 | Ets2 | UP | 0.297305768 | 0.0013196 |  |
| ENSMUSG00000027646 | Src | UP | 0.298438154 | 0.0069789 |  |
| ENSMUSG00000031007 | Atp6ap2 | UP | 0.298931466 | 0.0061099 |  |
| ENSMUSG00000032802 | Srxn1 | UP | 0.300882306 | 0.0188558 |  |
| ENSMUSG00000025314 | Ptprj | UP | 0.313726701 | 0.0203857 |  |
| ENSMUSG00000042524 | Sun2 | UP | 0.31430503 | 0.0058996 |  |
| ENSMUSG00000062300 | Pvrl2 | UP | 0.314668 | 0.0460454 |  |
| ENSMUSG00000050390 | C77080 | UP | 0.317335205 | 0.0012008 |  |
| ENSMUSG00000006463 | Zdhhc24 | UP | 0.317540752 | 0.0197379 |  |
| **ENSMUSG00000024913** | **Lrp5** | **UP** | **0.320003496** | **0.0134185** | **Wnt** |
| ENSMUSG00000018143 | Mafk | UP | 0.323344517 | 0.0437829 |  |
| ENSMUSG00000032261 | Sh3bgrl2 | UP | 0.323712892 | 0.0097477 |  |
| ENSMUSG00000063317 | Usp31 | UP | 0.341602785 | 0.0168585 |  |
| ENSMUSG00000038244 | Mical2 | UP | 0.343005087 | 0.0247122 |  |
| ENSMUSG00000074918 | Inafm2 | UP | 0.345222745 | 0.0075057 |  |
| ENSMUSG00000017707 | Serinc3 | UP | 0.347640965 | 0.0024549 |  |
| ENSMUSG00000024074 | Crim1 | UP | 0.355755942 | 0.036443 |  |
| ENSMUSG00000025509 | Pnpla2 | UP | 0.357424017 | 0.0488926 |  |
| ENSMUSG00000015839 | Nfe2l2 | UP | 0.361080739 | 0.0265442 |  |
| ENSMUSG00000026380 | Tfcp2l1 | UP | 0.363300525 | 3.793E-06 |  |
| ENSMUSG00000067825 | Pex26 | UP | 0.36348189 | 0.0001671 |  |
| ENSMUSG00000029622 | Arpc1b | UP | 0.364276431 | 0.0130132 |  |
| ENSMUSG00000036617 | Etl4 | UP | 0.36521396 | 0.0071537 |  |
| ENSMUSG00000036167 | Pphln1 | UP | 0.367170044 | 0.028281 |  |
| ENSMUSG00000027624 | Epb4.1l1 | UP | 0.367296671 | 0.0203857 |  |
| ENSMUSG00000005251 | Ripk4 | UP | 0.368107224 | 0.0038516 |  |
| ENSMUSG00000035547 | Capn5 | UP | 0.369624865 | 0.0193559 |  |
| ENSMUSG00000030340 | Scnn1a | UP | 0.375203401 | 0.0308362 |  |
| ENSMUSG00000026479 | Lamc2 | UP | 0.37582023 | 0.0128108 |  |
| ENSMUSG00000041143 | Tmco4 | UP | 0.379136874 | 0.0203857 |  |
| ENSMUSG00000054027 | Nt5dc3 | UP | 0.379818017 | 0.0250814 |  |
| ENSMUSG00000004044 | Ptrf | UP | 0.382205983 | 0.0266844 |  |
| ENSMUSG00000003309 | Ap1m2 | UP | 0.385967307 | 0.0181214 |  |
| ENSMUSG00000025225 | Nfkb2 | UP | 0.386813531 | 0.0095333 |  |
| ENSMUSG00000032194 | Kank2 | UP | 0.391765896 | 0.0160893 |  |
| ENSMUSG00000060098 | Prmt7 | UP | 0.392516593 | 0.0056613 |  |
| **ENSMUSG00000021466** | **Ptch1** | **UP** | **0.39825017** | **0.0003563** | **Hedgehog** |
| ENSMUSG00000020592 | Sdc1 | UP | 0.399579523 | 0.0130935 |  |
| ENSMUSG00000000308 | Ckmt1 | UP | 0.404502577 | 0.0181214 |  |
| ENSMUSG00000043144 | Aqp6 | UP | 0.404557092 | 0.0181214 |  |
| **ENSMUSG00000020176** | **Grb10** | **UP** | **0.408052741** | **0.012055** | **mTOR** |
| ENSMUSG00000050335 | Lgals3 | UP | 0.408614423 | 0.0061518 |  |
| ENSMUSG00000057969 | Sema3b | UP | 0.408840533 | 0.0004147 |  |
| ENSMUSG00000006373 | Pgrmc1 | UP | 0.408972273 | 0.0084563 |  |
| ENSMUSG00000025366 | Esyt1 | UP | 0.410044256 | 0.0381244 |  |
| ENSMUSG00000018217 | Pmp22 | UP | 0.412152694 | 0.0203857 |  |
| ENSMUSG00000067586 | S1pr3 | UP | 0.41313788 | 0.0370293 |  |
| ENSMUSG00000059248 | Sept9 | UP | 0.416050886 | 1.532E-05 |  |
| ENSMUSG00000001435 | Col18a1 | UP | 0.416354583 | 0.0023928 |  |
| ENSMUSG00000029994 | Anxa4 | UP | 0.418862038 | 0.0024549 |  |
| ENSMUSG00000027134 | Lpcat4 | UP | 0.422831955 | 0.0426574 |  |
| ENSMUSG00000021493 | Pdlim7 | UP | 0.424458304 | 0.0335411 |  |
| ENSMUSG00000036853 | Mcoln3 | UP | 0.425759609 | 0.0087545 |  |
| ENSMUSG00000037110 | Ralgapa2 | UP | 0.427335987 | 0.0002066 |  |
| ENSMUSG00000002265 | Peg3 | UP | 0.439082158 | 0.0005996 |  |
| ENSMUSG00000027188 | Pamr1 | UP | 0.440703059 | 0.0430761 |  |
| ENSMUSG00000074622 | Mafb | UP | 0.441376161 | 0.0229235 |  |
| ENSMUSG00000008305 | Tle1 | UP | 0.441566055 | 0.0435077 |  |
| ENSMUSG00000034382 | AI661453 | UP | 0.443714831 | 0.0020755 |  |
| ENSMUSG00000030727 | Rabep2 | UP | 0.448224609 | 2.241E-05 |  |
| ENSMUSG00000050052 | Tdrp | UP | 0.449106321 | 0.0005504 |  |
| ENSMUSG00000040703 | Cyp2s1 | UP | 0.455000447 | 0.0236709 |  |
| ENSMUSG00000005973 | Rcn1 | UP | 0.457796042 | 0.0047668 |  |
| **ENSMUSG00000036867** | **Smad6** | **UP** | **0.462782018** | **0.036443** | **TGF-beta** |
| ENSMUSG00000021061 | Sptb | UP | 0.473219562 | 0.0268423 |  |
| **ENSMUSG00000050965** | **Prkca** | **UP** | **0.47774662** | **0.0141656** | **Wnt** |
| **ENSMUSG00000032744** | **Heyl** | **UP** | **0.478122555** | **0.0189548** | **Notch** |
| ENSMUSG00000041598 | Cdc42ep4 | UP | 0.480367264 | 0.013448 |  |
| **ENSMUSG00000040010** | **Slc7a5** | **UP** | **0.482152353** | **0.0117863** | **mTOR** |
| ENSMUSG00000022178 | Ajuba | UP | 0.484793091 | 0.0038124 |  |
| ENSMUSG00000031995 | St14 | UP | 0.48956026 | 0.0001112 |  |
| ENSMUSG00000026303 | Mlph | UP | 0.490494916 | 0.0482496 |  |
| ENSMUSG00000024479 | Mal2 | UP | 0.491201019 | 0.0004746 |  |
| ENSMUSG00000026879 | Gsn | UP | 0.494724771 | 0.0236709 |  |
| ENSMUSG00000035413 | Tmem98 | UP | 0.498431348 | 0.0047146 |  |
| ENSMUSG00000037638 | Zbtb42 | UP | 0.49983493 | 0.0045915 |  |
| ENSMUSG00000036473 | Tbc1d24 | UP | 0.500008198 | 0.0006768 |  |
| ENSMUSG00000076431 | Sox4 | UP | 0.512172997 | 0.0045894 |  |
| ENSMUSG00000041313 | Slc7a1 | UP | 0.51303399 | 0.0001923 |  |
| ENSMUSG00000027669 | Gnb4 | UP | 0.514945316 | 0.0045915 |  |
| ENSMUSG00000022912 | Pros1 | UP | 0.515695329 | 0.0445749 |  |
| ENSMUSG00000023039 | Krt7 | UP | 0.518635368 | 0.0176824 |  |
| ENSMUSG00000037408 | Cnnm4 | UP | 0.519146576 | 0.0423792 |  |
| ENSMUSG00000023972 | Ptk7 | UP | 0.51941513 | 0.0112464 |  |
| ENSMUSG00000042115 | Klhdc8a | UP | 0.527600576 | 0.0001112 |  |
| ENSMUSG00000042523 | Dnal1 | UP | 0.528252342 | 0.0001646 |  |
| ENSMUSG00000020911 | Krt19 | UP | 0.535513484 | 0.034497 |  |
| ENSMUSG00000038578 | Susd1 | UP | 0.548606263 | 0.0019387 |  |
| ENSMUSG00000006930 | Hap1 | UP | 0.552457037 | 0.0036718 |  |
| ENSMUSG00000040093 | Bmf | UP | 0.554744512 | 0.0008085 |  |
| ENSMUSG00000031822 | Gse1 | UP | 0.561270181 | 1.054E-06 |  |
| **ENSMUSG00000027314** | **Dll4** | **UP** | **0.562392524** | **0.0058752** | **Notch** |
| ENSMUSG00000061451 | Tmem151a | UP | 0.564061618 | 0.0468359 |  |
| ENSMUSG00000036528 | Ppfibp2 | UP | 0.568128827 | 0.0002066 |  |
| ENSMUSG00000049866 | Arl4c | UP | 0.568595674 | 0.0130935 |  |
| ENSMUSG00000007097 | Atp1a2 | UP | 0.568962511 | 0.0243702 |  |
| ENSMUSG00000066438 | Plekhd1 | UP | 0.569700473 | 0.036443 |  |
| ENSMUSG00000039865 | Slc44a3 | UP | 0.578337825 | 0.0099422 |  |
| ENSMUSG00000026421 | Csrp1 | UP | 0.582779057 | 0.0001642 |  |
| ENSMUSG00000046318 | Ccbe1 | UP | 0.585361098 | 0.0035767 |  |
| ENSMUSG00000019817 | Plagl1 | UP | 0.589228299 | 2.315E-05 |  |
| ENSMUSG00000057604 | Lmcd1 | UP | 0.590031408 | 0.004569 |  |
| ENSMUSG00000020123 | Avpr1a | UP | 0.600224415 | 0.0242193 |  |
| ENSMUSG00000020814 | Mxra7 | UP | 0.608334936 | 0.0339685 |  |
| ENSMUSG00000033590 | Myo5c | UP | 0.613749903 | 0.0023117 |  |
| ENSMUSG00000027799 | Nbea | UP | 0.61783141 | 2.186E-10 |  |
| ENSMUSG00000024066 | Xdh | UP | 0.622870432 | 0.0211084 |  |
| ENSMUSG00000010154 | Spire2 | UP | 0.628977647 | 0.0007217 |  |
| ENSMUSG00000009545 | Kcnq1 | UP | 0.636473519 | 0.0004136 |  |
| ENSMUSG00000009394 | Syn2 | UP | 0.658149008 | 0.0108635 |  |
| ENSMUSG00000018569 | Cldn7 | UP | 0.659854138 | 0.0049517 |  |
| ENSMUSG00000050074 | Spink8 | UP | 0.676976969 | 0.0002014 |  |
| ENSMUSG00000021728 | Emb | UP | 0.681569827 | 0.0014548 |  |
| ENSMUSG00000032942 | Ucp3 | UP | 0.683712207 | 0.0223558 |  |
| **ENSMUSG00000036856** | **Wnt4** | **UP** | **0.700974885** | **0.0056613** | **Wnt** |
| ENSMUSG00000023827 | Agpat4 | UP | 0.703821708 | 0.0100933 |  |
| ENSMUSG00000045827 | Serpinb9 | UP | 0.705632946 | 1.733E-06 |  |
| ENSMUSG00000005672 | Kit | UP | 0.709090132 | 8.151E-06 |  |
| ENSMUSG00000029153 | Ociad2 | UP | 0.712399247 | 1.697E-05 |  |
| ENSMUSG00000031765 | Mt1 | UP | 0.723992372 | 0.0049517 |  |
| ENSMUSG00000008153 | Clstn3 | UP | 0.728642266 | 0.0104164 |  |
| ENSMUSG00000023043 | Krt18 | UP | 0.737632599 | 2.906E-07 |  |
| ENSMUSG00000004552 | Ctse | UP | 0.738982477 | 0.0203857 |  |
| ENSMUSG00000040711 | Sh3pxd2b | UP | 0.749654556 | 2.241E-05 |  |
| ENSMUSG00000024331 | Dsc2 | UP | 0.76481701 | 5.75E-05 |  |
| **ENSMUSG00000027276** | **Jag1** | **UP** | **0.765439995** | **9.102E-10** | **Notch** |
| ENSMUSG00000038732 | Mboat1 | UP | 0.793570776 | 0.0108748 |  |
| ENSMUSG00000024736 | Tmem132a | UP | 0.805446235 | 4.002E-08 |  |
| ENSMUSG00000020758 | Itgb4 | UP | 0.80567261 | 2.292E-10 |  |
| ENSMUSG00000054951 | 9130008F23Rik | UP | 0.814906707 | 0.036656 |  |
| ENSMUSG00000051596 | Otop1 | UP | 0.819265997 | 0.0293981 |  |
| ENSMUSG00000042622 | Maff | UP | 0.836710615 | 0.0058654 |  |
| ENSMUSG00000044313 | Mab21l3 | UP | 0.839324834 | 0.011715 |  |
| ENSMUSG00000039960 | Rhou | UP | 0.840530526 | 9.882E-07 |  |
| **ENSMUSG00000006269** | **Atp6v1b1** | **UP** | **0.844506537** | **4.525E-08** | **Notch** |
| ENSMUSG00000063430 | Wscd2 | UP | 0.84757105 | 2.507E-09 |  |
| ENSMUSG00000027513 | Pck1 | UP | 0.850830075 | 2.484E-05 |  |
| ENSMUSG00000025161 | Slc16a3 | UP | 0.852832765 | 0.0108635 |  |
| **ENSMUSG00000055980** | **Irs1** | **UP** | **0.855223203** | **1.687E-07** | **mTOR** |
| ENSMUSG00000032373 | Car12 | UP | 0.860352758 | 2.439E-28 |  |
| ENSMUSG00000037185 | Krt80 | UP | 0.861564028 | 5.523E-07 |  |
| ENSMUSG00000029765 | Plxna4 | UP | 0.862361648 | 0.0001593 |  |
| ENSMUSG00000047501 | Cldn4 | UP | 0.866789425 | 1.316E-05 |  |
| ENSMUSG00000015702 | Anxa9 | UP | 0.867178563 | 0.0473203 |  |
| ENSMUSG00000020798 | Spns3 | UP | 0.885569937 | 0.0436481 |  |
| ENSMUSG00000018727 | Cpsf4l | UP | 0.908351668 | 2.241E-05 |  |
| ENSMUSG00000060548 | Tnfrsf19 | UP | 0.924922043 | 0.0160893 |  |
| ENSMUSG00000028076 | Cd1d1 | UP | 0.927641079 | 0.0407945 |  |
| ENSMUSG00000022265 | Ank | UP | 0.942883371 | 2.078E-11 |  |
| ENSMUSG00000042842 | Serpinb6b | UP | 0.962757724 | 2.15E-05 |  |
| ENSMUSG00000044156 | Hepacam2 | UP | 0.966792541 | 2.334E-09 |  |
| ENSMUSG00000040564 | Apoc1 | UP | 0.969438823 | 0.0061518 |  |
| ENSMUSG00000055485 | Soga1 | UP | 0.985290868 | 1.191E-13 |  |
| ENSMUSG00000052760 | A630001G21Rik | UP | 0.997187462 | 0.0060229 |  |
| ENSMUSG00000041773 | Enc1 | UP | 1.009559314 | 1.815E-09 |  |
| ENSMUSG00000061527 | Krt5 | UP | 1.013656947 | 0.0014273 |  |
| ENSMUSG00000101304 | Plet1os | UP | 1.013988163 | 0.00872 |  |
| ENSMUSG00000039323 | Igfbp2 | UP | 1.034708474 | 0.0008148 |  |
| ENSMUSG00000024990 | Rbp4 | UP | 1.037396918 | 0.0236709 |  |
| ENSMUSG00000059430 | Actg2 | UP | 1.041099458 | 0.0129262 |  |
| ENSMUSG00000002699 | Lcp2 | UP | 1.063547582 | 0.0308362 |  |
| ENSMUSG00000021303 | Gng4 | UP | 1.064926016 | 0.0061518 |  |
| **ENSMUSG00000026394** | **Atp6v1g3** | **UP** | **1.07636153** | **1.27E-11** | **mTOR** |
| ENSMUSG00000036006 | Fam65b | UP | 1.103473199 | 0.0001275 |  |
| ENSMUSG00000032068 | Plet1 | UP | 1.106096858 | 6.591E-12 |  |
| ENSMUSG00000097184 | 4632428C04Rik | UP | 1.107119972 | 0.0171002 |  |
| ENSMUSG00000034330 | Plcg2 | UP | 1.109811535 | 1.553E-17 |  |
| ENSMUSG00000006457 | Actn3 | UP | 1.113395162 | 0.0005921 |  |
| ENSMUSG00000063296 | Tmem117 | UP | 1.136334534 | 3.456E-12 |  |
| ENSMUSG00000021280 | Exoc3l4 | UP | 1.147289373 | 5.924E-11 |  |
| ENSMUSG00000029254 | Stap1 | UP | 1.149655737 | 0.0036957 |  |
| ENSMUSG00000096573 | 1700009J07Rik | UP | 1.153580802 | 0.007559 |  |
| ENSMUSG00000051985 | Igfn1 | UP | 1.160467469 | 0.0014863 |  |
| **ENSMUSG00000028238** | **Atp6v0d2** | **UP** | **1.185468171** | **2.7E-15** | **mTOR** |
| ENSMUSG00000021070 | Bdkrb2 | UP | 1.192795096 | 0.0056823 |  |
| ENSMUSG00000039533 | Mmd2 | UP | 1.222389458 | 0.0001782 |  |
| ENSMUSG00000109764 | Klkb1 | UP | 1.247781642 | 3.191E-08 |  |
| ENSMUSG00000049999 | Ppp1r3d | UP | 1.294314006 | 0.0207635 |  |
| ENSMUSG00000002588 | Pon1 | UP | 1.298544412 | 0.0263924 |  |
| **ENSMUSG00000014773** | **Dll1** | **UP** | **1.305757657** | **5.606E-25** | **Notch** |
| ENSMUSG00000023247 | Guca2a | UP | 1.361666324 | 1.047E-06 |  |
| ENSMUSG00000022871 | Fetub | UP | 1.37791598 | 0.0206313 |  |
| **ENSMUSG00000020566** | **Atp6v1c2** | **UP** | **1.384801629** | **3.564E-17** | **mTOR** |
| **ENSMUSG00000032726** | **Bmp8a** | **UP** | **1.391943697** | **0.0039002** | **TGF-beta** |
| ENSMUSG00000051728 | 4930563D23Rik | UP | 1.402985948 | 0.001449 |  |
| ENSMUSG00000035735 | Dagla | UP | 1.424110229 | 7.624E-15 |  |
| ENSMUSG00000086243 | Gm12121 | UP | 1.428461442 | 2.039E-09 |  |
| ENSMUSG00000042306 | S100a14 | UP | 1.442052992 | 9.57E-05 |  |
| ENSMUSG00000020651 | Slc26a4 | UP | 1.470658449 | 7.261E-23 |  |
| ENSMUSG00000022229 | Atp12a | UP | 1.488648561 | 0.0011216 |  |
| ENSMUSG00000020080 | Hkdc1 | UP | 1.522628769 | 1.004E-12 |  |
| ENSMUSG00000024743 | Syt7 | UP | 1.533894963 | 3.208E-19 |  |
| ENSMUSG00000037833 | Sh2d4b | UP | 1.547056446 | 5.924E-11 |  |
| ENSMUSG00000026347 | Tmem163 | UP | 1.554183568 | 7.809E-13 |  |
| ENSMUSG00000115772 | Gm48935 | UP | 1.650520219 | 0.0175447 |  |
| ENSMUSG00000116967 | Smim34 | UP | 1.68694139 | 0.0058996 |  |
| ENSMUSG00000028072 | Ntrk1 | UP | 1.699660165 | 4.104E-11 |  |
| ENSMUSG00000024485 | Slc4a9 | UP | 1.705737849 | 1.168E-22 |  |
| ENSMUSG00000032315 | Cyp1a1 | UP | 1.751045654 | 2.147E-08 |  |
| ENSMUSG00000005640 | Insrr | UP | 1.782949183 | 2.4E-26 |  |
| ENSMUSG00000047861 | Foxi1 | UP | 1.829909861 | 1.466E-33 |  |
| ENSMUSG00000020010 | Vnn3 | UP | 2.010600944 | 0.0468399 |  |
| ENSMUSG00000026414 | Tnnt2 | UP | 2.067704823 | 0.0064472 |  |
| ENSMUSG00000027524 | Edn3 | UP | 2.094319428 | 0.0003868 |  |
| ENSMUSG00000111225 | Gm47790 | UP | 2.323340034 | 5.542E-07 |  |
| ENSMUSG00000074628 | Tldc2 | UP | 2.370054465 | 6.24E-37 |  |
| ENSMUSG00000111014 | Gm47795 | UP | 2.448303257 | 7.261E-21 |  |
| ENSMUSG00000020256 | Aldh1l2 | UP | 2.488540518 | 1.449E-21 |  |
| ENSMUSG00000085684 | 4930469K13Rik | UP | 2.497706564 | 1.262E-16 |  |
| ENSMUSG00000050100 | Hmx2 | UP | 2.52209301 | 1.469E-51 |  |
| **ENSMUSG00000033227** | **Wnt6** | **UP** | **2.628154259** | **2.435E-10** | **Wnt** |
| ENSMUSG00000040148 | Hmx3 | UP | 2.908269116 | 7.198E-14 |  |
