## Supplementary Table 4 for "Kidney collecting duct cell type composition is regulated by Notch signaling via modulation of mTORC1"

| **Supplementary Table 4: Hes1-FLAG ChIP-seq results from mpkCCDc14 cells expressing inducible Hes1-FLAG.** | | | | | |
| --- | --- | --- | --- | --- | --- |
| **Location of Peak** | **Start** | **End** | **ENSEMBL** | **SYMBOL** | **GENENAME** |
| chr1 | 4807570 | 4808237 | ENSMUSG00000025903 | Lypla1 | lysophospholipase 1 |
| chr1 | 4857438 | 4858361 | ENSMUSG00000033813 | Tcea1 | transcription elongation factor A (SII) 1 |
| chr1 | 7088349 | 7089444 | ENSMUSG00000051285 | Pcmtd1 | protein-L-isoaspartate (D-aspartate) O-methyltransferase domain containing 1 |
| chr1 | 9545391 | 9546309 | ENSMUSG00000061024 | Rrs1 | ribosome biogenesis regulator 1 |
| chr1 | 9699721 | 9700859 | ENSMUSG00000025912 | Mybl1 | myeloblastosis oncogene-like 1 |
| chr1 | 9797842 | 9798553 | ENSMUSG00000025915 | Sgk3 | serum/glucocorticoid regulated kinase 3 |
| chr1 | 10039595 | 10040269 | ENSMUSG00000056763 | Cspp1 | centrosome and spindle pole associated protein 1 |
| chr1 | 10232014 | 10233521 | ENSMUSG00000067851 | Arfgef1 | ADP-ribosylation factor guanine nucleotide-exchange factor 1(brefeldin A-inhibited) |
| chr1 | 13660111 | 13660795 | ENSMUSG00000025937 | Lactb2 | lactamase, beta 2 |
| chr1 | 16104868 | 16106501 | ENSMUSG00000025921 | Rdh10 | retinol dehydrogenase 10 (all-trans) |
| chr1 | 23255895 | 23257141 | ENSMUSG00000065405 | Mir30a | microRNA 30a |
| chr1 | 23921704 | 23922570 | ENSMUSG00000026155 | Smap1 | small ArfGAP 1 |
| chr1 | 30872890 | 30874065 | ENSMUSG00000048874 | Phf3 | PHD finger protein 3 |
| chr1 | 30949007 | 30950262 | ENSMUSG00000026064 | Ptp4a1 | protein tyrosine phosphatase 4a1 |
| chr1 | 33719768 | 33720540 | ENSMUSG00000004768 | Rab23 | RAB23, member RAS oncogene family |
| chr1 | 33813745 | 33814696 | ENSMUSG00000042197 | Zfp451 | zinc finger protein 451 |
| chr1 | 33907746 | 33908386 | ENSMUSG00000026131 | Dst | dystonin |
| chr1 | 34842517 | 34843300 | ENSMUSG00000037503 | Fam168b | family with sequence similarity 168, member B |
| chr1 | 36368654 | 36369208 | ENSMUSG00000010453 | Kansl3 | KAT8 regulatory NSL complex subunit 3 |
| chr1 | 36471244 | 36472067 | ENSMUSG00000037408 | Cnnm4 | cyclin M4 |
| chr1 | 36511497 | 36513076 | ENSMUSG00000001138 | Cnnm3 | cyclin M3 |
| chr1 | 37429817 | 37430543 | ENSMUSG00000026111 | Unc50 | unc-50 homolog |
| chr1 | 37540425 | 37541096 | ENSMUSG00000026110 | Mgat4a | mannoside acetylglucosaminyltransferase 4, isoenzyme A |
| chr1 | 37719373 | 37720140 | ENSMUSG00000026090 | Cracdl | capping protein inhibiting regulator of actin like |
| chr1 | 38128802 | 38130338 | ENSMUSG00000026082 | Rev1 | REV1, DNA directed polymerase |
| chr1 | 39535554 | 39536444 | ENSMUSG00000003135 | Cnot11 | CCR4-NOT transcription complex, subunit 11 |
| chr1 | 39576483 | 39577768 | ENSMUSG00000048234 | Rnf149 | ring finger protein 149 |
| chr1 | 39900564 | 39901491 | ENSMUSG00000026074 | Map4k4 | mitogen-activated protein kinase kinase kinase kinase 4 |
| chr1 | 43163735 | 43164184 | ENSMUSG00000008136 | Fhl2 | four and a half LIM domains 2 |
| chr1 | 43445497 | 43446570 | ENSMUSG00000066877 | Nck2 | non-catalytic region of tyrosine kinase adaptor protein 2 |
| chr1 | 43826899 | 43828090 | ENSMUSG00000057363 | Uxs1 | UDP-glucuronate decarboxylase 1 |
| chr1 | 43933548 | 43934306 | ENSMUSG00000041763 | Tpp2 | tripeptidyl peptidase II |
| chr1 | 44102015 | 44102629 | ENSMUSG00000026049 | Tex30 | testis expressed 30 |
| chr1 | 46852708 | 46853894 | ENSMUSG00000025986 | Slc39a10 | solute carrier family 39 (zinc transporter), member 10 |
| chr1 | 51915584 | 51916089 | ENSMUSG00000018417 | Myo1b | myosin IB |
| chr1 | 52232040 | 52233687 | ENSMUSG00000026103 | Gls | glutaminase |
| chr1 | 52499773 | 52501001 | ENSMUSG00000002881 | Nab1 | Ngfi-A binding protein 1 |
| chr1 | 52726472 | 52727494 | ENSMUSG00000041439 | Mfsd6 | major facilitator superfamily domain containing 6 |
| chr1 | 53296724 | 53297482 | ENSMUSG00000026098 | Pms1 | PMS1 homolog 1, mismatch repair system component |
| chr1 | 53784595 | 53785232 | ENSMUSG00000026094 | Stk17b | serine/threonine kinase 17b (apoptosis-inducing) |
| chr1 | 54925681 | 54926624 | ENSMUSG00000052331 | Ankrd44 | ankyrin repeat domain 44 |
| chr1 | 55087227 | 55088621 | ENSMUSG00000025980 | Hspd1 | heat shock protein 1 (chaperonin) |
| chr1 | 55130963 | 55131916 | ENSMUSG00000025979 | Mob4 | MOB family member 4, phocein |
| chr1 | 56972014 | 56973511 | ENSMUSG00000038331 | Satb2 | special AT-rich sequence binding protein 2 |
| chr1 | 58393050 | 58393914 | ENSMUSG00000051223 | Bzw1 | basic leucine zipper and W2 domains 1 |
| chr1 | 58973203 | 58974231 | ENSMUSG00000026027 | Stradb | STE20-related kinase adaptor beta |
| chr1 | 59481990 | 59483265 | ENSMUSG00000041075 | Fzd7 | frizzled class receptor 7 |
| chr1 | 59912525 | 59913973 | ENSMUSG00000041040 | Fam117b | family with sequence similarity 117, member B |
| chr1 | 60098197 | 60098809 | ENSMUSG00000026017 | Carf | calcium response factor |
| chr1 | 60180151 | 60181169 | ENSMUSG00000073664 | Nbeal1 | neurobeachin like 1 |
| chr1 | 60343086 | 60343687 | ENSMUSG00000049439 | Cyp20a1 | cytochrome P450, family 20, subfamily a, polypeptide 1 |
| chr1 | 60566143 | 60567270 | ENSMUSG00000026014 | Raph1 | Ras association (RalGDS/AF-6) and pleckstrin homology domains 1 |
| chr1 | 62702887 | 62703862 | ENSMUSG00000025969 | Nrp2 | neuropilin 2 |
| chr1 | 62706286 | 62707207 | ENSMUSG00000025969 | Nrp2 | neuropilin 2 |
| chr1 | 63113423 | 63114723 | ENSMUSG00000084799 | Ino80dos | INO80 complex subunit D, opposite strand |
| chr1 | 64690512 | 64691568 | ENSMUSG00000070871 | Ccnyl1 | cyclin Y-like 1 |
| chr1 | 64736876 | 64737980 | ENSMUSG00000045005 | Fzd5 | frizzled class receptor 5 |
| chr1 | 64737980 | 64738477 | ENSMUSG00000045005 | Fzd5 | frizzled class receptor 5 |
| chr1 | 66817108 | 66817989 | ENSMUSG00000026004 | Kansl1l | KAT8 regulatory NSL complex subunit 1-like |
| chr1 | 66862773 | 66863539 | ENSMUSG00000026003 | Acadl | acyl-Coenzyme A dehydrogenase, long-chain |
| chr1 | 69687092 | 69687951 | ENSMUSG00000025997 | Ikzf2 | IKAROS family zinc finger 2 |
| chr1 | 72211879 | 72212516 | ENSMUSG00000039395 | Mreg | melanoregulin |
| chr1 | 74236245 | 74237222 | ENSMUSG00000006304 | Arpc2 | actin related protein 2/3 complex, subunit 2 |
| chr1 | 74391241 | 74392288 | ENSMUSG00000026176 | Ctdsp1 | CTD (carboxy-terminal domain, RNA polymerase II, polypeptide A) small phosphatase 1 |
| chr1 | 74661655 | 74662268 | ENSMUSG00000033257 | Ttll4 | tubulin tyrosine ligase-like family, member 4 |
| chr1 | 75141412 | 75142464 | ENSMUSG00000033159 | Cnppd1 | cyclin Pas1/PHO80 domain containing 1 |
| chr1 | 75478689 | 75479835 | ENSMUSG00000051703 | Tmem198 | transmembrane protein 198 |
| chr1 | 75505318 | 75506443 | ENSMUSG00000026211 | Obsl1 | obscurin-like 1 |
| chr1 | 75506443 | 75506677 | ENSMUSG00000026211 | Obsl1 | obscurin-like 1 |
| chr1 | 78657652 | 78658526 | ENSMUSG00000079470 | Utp14b | UTP14B small subunit processome component |
| chr1 | 80339730 | 80340989 | ENSMUSG00000004364 | Cul3 | cullin 3 |
| chr1 | 82290702 | 82291891 | ENSMUSG00000055980 | Irs1 | insulin receptor substrate 1 |
| chr1 | 86358536 | 86359996 | ENSMUSG00000026234 | Ncl | nucleolin |
| chr1 | 86442140 | 86443020 | ENSMUSG00000036574 | Tex44 | testis expressed 44 |
| chr1 | 86503926 | 86504536 | ENSMUSG00000026238 | Ptma | prothymosin alpha |
| chr1 | 87213751 | 87214527 | ENSMUSG00000026254 | Eif4e2 | eukaryotic translation initiation factor 4E member 2 |
| chr1 | 87755845 | 87756520 | ENSMUSG00000026289 | Atg16l1 | autophagy related 16-like 1 (S. cerevisiae) |
| chr1 | 87853116 | 87853922 | ENSMUSG00000070738 | Dgkd | diacylglycerol kinase, delta |
| chr1 | 88277154 | 88277852 | ENSMUSG00000036251 | Trpm8 | transient receptor potential cation channel, subfamily M, member 8 |
| chr1 | 88700966 | 88702544 | ENSMUSG00000049866 | Arl4c | ADP-ribosylation factor-like 4C |
| chr1 | 89070764 | 89071366 | ENSMUSG00000036206 | Sh3bp4 | SH3-domain binding protein 4 |
| chr1 | 89454380 | 89456081 | ENSMUSG00000055013 | Agap1 | ArfGAP with GTPase domain, ankyrin repeat and PH domain 1 |
| chr1 | 89579931 | 89581631 | ENSMUSG00000055013 | Agap1 | ArfGAP with GTPase domain, ankyrin repeat and PH domain 1 |
| chr1 | 91366261 | 91366861 | ENSMUSG00000047443 | Erfe | erythroferrone |
| chr1 | 91494387 | 91494986 | ENSMUSG00000034292 | Traf3ip1 | TRAF3 interacting protein 1 |
| chr1 | 92831386 | 92832868 | ENSMUSG00000034220 | Gpc1 | glypican 1 |
| chr1 | 92906897 | 92907493 | ENSMUSG00000047067 | Dusp28 | dual specificity phosphatase 28 |
| chr1 | 92910485 | 92911535 | ENSMUSG00000026269 | Rnpepl1 | arginyl aminopeptidase (aminopeptidase B)-like 1 |
| chr1 | 93235482 | 93236550 | ENSMUSG00000047793 | Sned1 | sushi, nidogen and EGF-like domains 1 |
| chr1 | 93511750 | 93512484 | ENSMUSG00000034066 | Farp2 | FERM, RhoGEF and pleckstrin domain protein 2 |
| chr1 | 93634636 | 93635745 | ENSMUSG00000026277 | Stk25 | serine/threonine kinase 25 (yeast) |
| chr1 | 93685419 | 93686962 | ENSMUSG00000026278 | Bok | BCL2-related ovarian killer |
| chr1 | 93753950 | 93755450 | ENSMUSG00000026280 | Atg4b | autophagy related 4B, cysteine peptidase |
| chr1 | 97660595 | 97662376 | ENSMUSG00000044768 | Macir | macrophage immunometabolism regulator |
| chr1 | 97976807 | 97977334 | ENSMUSG00000026335 | Pam | peptidylglycine alpha-amidating monooxygenase |
| chr1 | 98094953 | 98095843 | ENSMUSG00000026335 | Pam | peptidylglycine alpha-amidating monooxygenase |
| chr1 | 105990321 | 105991736 | ENSMUSG00000038866 | Zcchc2 | zinc finger, CCHC domain containing 2 |
| chr1 | 106171616 | 106173216 | ENSMUSG00000044340 | Phlpp1 | PH domain and leucine rich repeat protein phosphatase 1 |
| chr1 | 119504209 | 119505140 | ENSMUSG00000004451 | Ralb | v-ral simian leukemia viral oncogene B |
| chr1 | 119525908 | 119526458 | ENSMUSG00000098923 | Tmem185b | transmembrane protein 185B |
| chr1 | 119836874 | 119837727 | ENSMUSG00000026384 | Ptpn4 | protein tyrosine phosphatase, non-receptor type 4 |
| chr1 | 125435159 | 125435891 | ENSMUSG00000026341 | Actr3 | ARP3 actin-related protein 3 |
| chr1 | 127773907 | 127774725 | ENSMUSG00000102014 | 2900009J06Rik | RIKEN cDNA 2900009J06 gene |
| chr1 | 128244057 | 128244938 | ENSMUSG00000026353 | Ubxn4 | UBX domain protein 4 |
| chr1 | 131096745 | 131098036 | ENSMUSG00000016528 | Mapkapk2 | MAP kinase-activated protein kinase 2 |
| chr1 | 131137771 | 131138501 | ENSMUSG00000016526 | Dyrk3 | dual-specificity tyrosine-(Y)-phosphorylation regulated kinase 3 |
| chr1 | 131962635 | 131963615 | ENSMUSG00000026435 | Slc45a3 | solute carrier family 45, member 3 |
| chr1 | 132007319 | 132008512 | ENSMUSG00000026436 | Elk4 | ELK4, member of ETS oncogene family |
| chr1 | 132190303 | 132191321 | ENSMUSG00000079330 | Lemd1 | LEM domain containing 1 |
| chr1 | 132298525 | 132299462 | ENSMUSG00000042115 | Klhdc8a | kelch domain containing 8A |
| chr1 | 133045931 | 133046759 | ENSMUSG00000026447 | Pik3c2b | phosphatidylinositol-4-phosphate 3-kinase catalytic subunit type 2 beta |
| chr1 | 133131017 | 133132308 | ENSMUSG00000046062 | Ppp1r15b | protein phosphatase 1, regulatory subunit 15B |
| chr1 | 133423401 | 133424374 | ENSMUSG00000070643 | Sox13 | SRY (sex determining region Y)-box 13 |
| chr1 | 134415169 | 134415989 | ENSMUSG00000026457 | Adipor1 | adiponectin receptor 1 |
| chr1 | 135146537 | 135147638 | ENSMUSG00000026426 | Arl8a | ADP-ribosylation factor-like 8A |
| chr1 | 135687863 | 135688530 | ENSMUSG00000009418 | Nav1 | neuron navigator 1 |
| chr1 | 135688530 | 135689096 | ENSMUSG00000009418 | Nav1 | neuron navigator 1 |
| chr1 | 135765959 | 135766853 | ENSMUSG00000041801 | Phlda3 | pleckstrin homology like domain, family A, member 3 |
| chr1 | 135820721 | 135821296 | ENSMUSG00000041782 | Lad1 | ladinin |
| chr1 | 136345340 | 136346382 | ENSMUSG00000041570 | Camsap2 | calmodulin regulated spectrin-associated protein family, member 2 |
| chr1 | 136415168 | 136415795 | ENSMUSG00000097403 | 9230116N13Rik | RIKEN cDNA 9230116N13 gene |
| chr1 | 138619141 | 138619922 | ENSMUSG00000026393 | Nek7 | NIMA (never in mitosis gene a)-related expressed kinase 7 |
| chr1 | 138962813 | 138964180 | ENSMUSG00000056268 | Dennd1b | DENN/MADD domain containing 1B |
| chr1 | 139422044 | 139422609 | ENSMUSG00000033964 | Zbtb41 | zinc finger and BTB domain containing 41 |
| chr1 | 143702009 | 143703218 | ENSMUSG00000026361 | Cdc73 | cell division cycle 73, Paf1/RNA polymerase II complex component |
| chr1 | 143739334 | 143739926 | ENSMUSG00000018196 | Glrx2 | glutaredoxin 2 (thioltransferase) |
| chr1 | 143776707 | 143777973 | ENSMUSG00000018199 | Ro60 | Ro60, Y RNA binding protein |
| chr1 | 150392828 | 150393836 | ENSMUSG00000006005 | Tpr | translocated promoter region, nuclear basket protein |
| chr1 | 151344099 | 151345573 | ENSMUSG00000023150 | Ivns1abp | influenza virus NS1A binding protein |
| chr1 | 151500127 | 151501091 | ENSMUSG00000026484 | Rnf2 | ring finger protein 2 |
| chr1 | 152089789 | 152090625 | ENSMUSG00000032666 | 1700025G04Rik | RIKEN cDNA 1700025G04 gene |
| chr1 | 152090625 | 152090964 | ENSMUSG00000032666 | 1700025G04Rik | RIKEN cDNA 1700025G04 gene |
| chr1 | 152765987 | 152767283 | ENSMUSG00000026482 | Rgl1 | ral guanine nucleotide dissociation stimulator,-like 1 |
| chr1 | 155098241 | 155099493 | ENSMUSG00000056708 | Ier5 | immediate early response 5 |
| chr1 | 155558098 | 155558775 | ENSMUSG00000033701 | Acbd6 | acyl-Coenzyme A binding domain containing 6 |
| chr1 | 156718397 | 156719222 | ENSMUSG00000033557 | Fam20b | family with sequence similarity 20, member B |
| chr1 | 156938895 | 156940024 | ENSMUSG00000026594 | Ralgps2 | Ral GEF with PH domain and SH3 binding motif 2 |
| chr1 | 157411684 | 157412717 | ENSMUSG00000070565 | Rasal2 | RAS protein activator like 2 |
| chr1 | 160212273 | 160212919 | ENSMUSG00000014226 | Cacybp | calcyclin binding protein |
| chr1 | 160906050 | 160907273 | ENSMUSG00000040423 | Rc3h1 | RING CCCH (C3H) domains 1 |
| chr1 | 161876138 | 161877050 | ENSMUSG00000040297 | Suco | SUN domain containing ossification factor |
| chr1 | 162739855 | 162740418 | ENSMUSG00000040225 | Prrc2c | proline-rich coiled-coil 2C |
| chr1 | 164248624 | 164249736 | ENSMUSG00000040918 | Slc19a2 | solute carrier family 19 (thiamine transporter), member 2 |
| chr1 | 164457347 | 164458553 | ENSMUSG00000026576 | Atp1b1 | ATPase, Na+/K+ transporting, beta 1 polypeptide |
| chr1 | 164458553 | 164459076 | ENSMUSG00000026576 | Atp1b1 | ATPase, Na+/K+ transporting, beta 1 polypeptide |
| chr1 | 165193877 | 165194522 | ENSMUSG00000040848 | Sft2d2 | SFT2 domain containing 2 |
| chr1 | 165281709 | 165282479 | ENSMUSG00000040836 | Gpr161 | G protein-coupled receptor 161 |
| chr1 | 165633964 | 165634853 | ENSMUSG00000026566 | Mpzl1 | myelin protein zero-like 1 |
| chr1 | 167283525 | 167285338 | ENSMUSG00000026558 | Uck2 | uridine-cytidine kinase 2 |
| chr1 | 170174364 | 170175364 | ENSMUSG00000026670 | Uap1 | UDP-N-acetylglucosamine pyrophosphorylase 1 |
| chr1 | 177257521 | 177259013 | ENSMUSG00000019699 | Akt3 | thymoma viral proto-oncogene 3 |
| chr1 | 177441449 | 177443243 | ENSMUSG00000063659 | Zbtb18 | zinc finger and BTB domain containing 18 |
| chr1 | 177795870 | 177796805 | ENSMUSG00000015961 | Adss | adenylosuccinate synthetase, non muscle |
| chr1 | 178318188 | 178319012 | ENSMUSG00000026500 | Cox20 | cytochrome c oxidase assembly protein 20 |
| chr1 | 178336395 | 178338128 | ENSMUSG00000039630 | Hnrnpu | heterogeneous nuclear ribonucleoprotein U |
| chr1 | 179546011 | 179546938 | ENSMUSG00000026492 | Tfb2m | transcription factor B2, mitochondrial |
| chr1 | 179667966 | 179668734 | ENSMUSG00000038936 | Sccpdh | saccharopine dehydrogenase (putative) |
| chr1 | 179802821 | 179803872 | ENSMUSG00000026491 | Ahctf1 | AT hook containing transcription factor 1 |
| chr1 | 179959609 | 179960644 | ENSMUSG00000026490 | Cdc42bpa | CDC42 binding protein kinase alpha |
| chr1 | 180255961 | 180256429 | ENSMUSG00000010609 | Psen2 | presenilin 2 |
| chr1 | 180813098 | 180814411 | ENSMUSG00000060743 | H3f3a | H3.3 histone A |
| chr1 | 181210889 | 181212476 | ENSMUSG00000038733 | Wdr26 | WD repeat domain 26 |
| chr1 | 181841483 | 181842887 | ENSMUSG00000004880 | Lbr | lamin B receptor |
| chr1 | 182019300 | 182020331 | ENSMUSG00000022995 | Enah | ENAH actin regulator |
| chr1 | 182282058 | 182282935 | ENSMUSG00000038633 | Degs1 | delta(4)-desaturase, sphingolipid 1 |
| chr1 | 182516908 | 182517825 | ENSMUSG00000026509 | Capn2 | calpain 2 |
| chr1 | 184033382 | 184034622 | ENSMUSG00000039384 | Dusp10 | dual specificity phosphatase 10 |
| chr1 | 184845372 | 184846088 | ENSMUSG00000073481 | Mtarc2 | mitochondrial amidoxime reducing component 2 |
| chr1 | 185203930 | 185204640 | ENSMUSG00000039318 | Rab3gap2 | RAB3 GTPase activating protein subunit 2 |
| chr1 | 185362788 | 185363498 | ENSMUSG00000026615 | Eprs | glutamyl-prolyl-tRNA synthetase |
| chr1 | 186705097 | 186705708 | ENSMUSG00000039239 | Tgfb2 | transforming growth factor, beta 2 |
| chr1 | 189727541 | 189728975 | ENSMUSG00000026604 | Ptpn14 | protein tyrosine phosphatase, non-receptor type 14 |
| chr1 | 189921884 | 189922657 | ENSMUSG00000026603 | Smyd2 | SET and MYND domain containing 2 |
| chr1 | 190928391 | 190929236 | ENSMUSG00000026634 | Angel2 | angel homolog 2 |
| chr1 | 191025111 | 191026417 | ENSMUSG00000097845 | A230020J21Rik | RIKEN cDNA A230020J21 gene |
| chr1 | 191182384 | 191183776 | ENSMUSG00000026628 | Atf3 | activating transcription factor 3 |
| chr1 | 191396123 | 191397295 | ENSMUSG00000026626 | Ppp2r5a | protein phosphatase 2, regulatory subunit B', alpha |
| chr1 | 191717773 | 191718813 | ENSMUSG00000026623 | Lpgat1 | lysophosphatidylglycerol acyltransferase 1 |
| chr1 | 191718813 | 191719248 | ENSMUSG00000026623 | Lpgat1 | lysophosphatidylglycerol acyltransferase 1 |
| chr1 | 191906193 | 191907818 | ENSMUSG00000037434 | Slc30a1 | solute carrier family 30 (zinc transporter), member 1 |
| chr1 | 192854796 | 192855978 |  | Gm15867 | predicted gene 15867 |
| chr1 | 194976223 | 194976695 | ENSMUSG00000096929 | A330023F24Rik | RIKEN cDNA A330023F24 gene |
| chr1 | 194976695 | 194976943 | ENSMUSG00000097325 | Gm16897 | predicted gene, 16897 |
| chr10 | 3973115 | 3973981 | ENSMUSG00000040675 | Mthfd1l | methylenetetrahydrofolate dehydrogenase (NADP+ dependent) 1-like |
| chr10 | 7211032 | 7212613 | ENSMUSG00000015202 | Cnksr3 | Cnksr family member 3 |
| chr10 | 7589613 | 7590706 | ENSMUSG00000019796 | Lrp11 | low density lipoprotein receptor-related protein 11 |
| chr10 | 7662589 | 7663619 | ENSMUSG00000019795 | Pcmt1 | protein-L-isoaspartate (D-aspartate) O-methyltransferase 1 |
| chr10 | 7680824 | 7681971 | ENSMUSG00000040021 | Lats1 | large tumor suppressor |
| chr10 | 7955091 | 7956966 | ENSMUSG00000015755 | Tab2 | TGF-beta activated kinase 1/MAP3K7 binding protein 2 |
| chr10 | 8885417 | 8886763 | ENSMUSG00000015305 | Sash1 | SAM and SH3 domain containing 1 |
| chr10 | 9674376 | 9675580 | ENSMUSG00000060487 | Samd5 | sterile alpha motif domain containing 5 |
| chr10 | 9900417 | 9901548 | ENSMUSG00000019790 | Stxbp5 | syntaxin binding protein 5 (tomosyn) |
| chr10 | 11280618 | 11282161 | ENSMUSG00000047648 | Fbxo30 | F-box protein 30 |
| chr10 | 12868021 | 12868932 | ENSMUSG00000019820 | Utrn | utrophin |
| chr10 | 13323904 | 13324355 | ENSMUSG00000062866 | Phactr2 | phosphatase and actin regulator 2 |
| chr10 | 14544739 | 14545674 | ENSMUSG00000039116 | Adgrg6 | adhesion G protein-coupled receptor G6 |
| chr10 | 17947080 | 17948403 | ENSMUSG00000039879 | Heca | hdc homolog, cell cycle regulator |
| chr10 | 18055397 | 18056390 | ENSMUSG00000019854 | Reps1 | RalBP1 associated Eps domain containing protein |
| chr10 | 18318358 | 18319595 | ENSMUSG00000039835 | Nhsl1 | NHS-like 1 |
| chr10 | 18844684 | 18845486 | ENSMUSG00000019851 | Perp | PERP, TP53 apoptosis effector |
| chr10 | 19933460 | 19935092 | ENSMUSG00000071369 | Map3k5 | mitogen-activated protein kinase kinase kinase 5 |
| chr10 | 20148400 | 20149719 | ENSMUSG00000019996 | Map7 | microtubule-associated protein 7 |
| chr10 | 20312010 | 20313309 | ENSMUSG00000037608 | Bclaf1 | BCL2-associated transcription factor 1 |
| chr10 | 20952385 | 20953000 | ENSMUSG00000019986 | Ahi1 | Abelson helper integration site 1 |
| chr10 | 21993890 | 21994810 | ENSMUSG00000019970 | Sgk1 | serum/glucocorticoid regulated kinase 1 |
| chr10 | 22148265 | 22149585 | ENSMUSG00000097327 | E030030I06Rik | RIKEN cDNA E030030I06 gene |
| chr10 | 24595316 | 24596622 |  | Gm15270 | predicted gene 15270 |
| chr10 | 25296696 | 25297299 | ENSMUSG00000039166 | Akap7 | A kinase (PRKA) anchor protein 7 |
| chr10 | 25359600 | 25360572 | ENSMUSG00000019978 | Epb41l2 | erythrocyte membrane protein band 4.1 like 2 |
| chr10 | 29313018 | 29313702 | ENSMUSG00000019883 | Echdc1 | enoyl Coenzyme A hydratase domain containing 1 |
| chr10 | 29361372 | 29362373 | ENSMUSG00000038876 | Rnf146 | ring finger protein 146 |
| chr10 | 30200220 | 30200606 | ENSMUSG00000075266 | Cenpw | centromere protein W |
| chr10 | 30600313 | 30600914 | ENSMUSG00000019792 | Trmt11 | tRNA methyltransferase 11 |
| chr10 | 30618099 | 30618603 | ENSMUSG00000019791 | Hint3 | histidine triad nucleotide binding protein 3 |
| chr10 | 30802489 | 30803560 | ENSMUSG00000039697 | Ncoa7 | nuclear receptor coactivator 7 |
| chr10 | 30842311 | 30843251 | ENSMUSG00000019789 | Hey2 | hairy/enhancer-of-split related with YRPW motif 2 |
| chr10 | 31608424 | 31609936 | ENSMUSG00000063760 | Rnf217 | ring finger protein 217 |
| chr10 | 34019173 | 34020150 | ENSMUSG00000019782 | Rwdd1 | RWD domain containing 1 |
| chr10 | 34418187 | 34418665 | ENSMUSG00000039480 | Nt5dc1 | 5'-nucleotidase domain containing 1 |
| chr10 | 36973975 | 36975164 | ENSMUSG00000019777 | Hdac2 | histone deacetylase 2 |
| chr10 | 37136007 | 37137328 | ENSMUSG00000069662 | Marcks | myristoylated alanine rich protein kinase C substrate |
| chr10 | 39369385 | 39370554 | ENSMUSG00000019843 | Fyn | Fyn proto-oncogene |
| chr10 | 39898783 | 39899348 |  | Mfsd4b4 | major facilitator superfamily domain containing 4B4 |
| chr10 | 40141759 | 40142498 | ENSMUSG00000019838 | Slc16a10 | solute carrier family 16 (monocarboxylic acid transporters), member 10 |
| chr10 | 40300726 | 40302648 | ENSMUSG00000063953 | Amd2 | S-adenosylmethionine decarboxylase 2 |
| chr10 | 40348966 | 40350051 | ENSMUSG00000038481 | Cdk19 | cyclin-dependent kinase 19 |
| chr10 | 40883436 | 40884611 | ENSMUSG00000019831 | Wasf1 | WASP family, member 1 |
| chr10 | 41519044 | 41520175 | ENSMUSG00000019818 | Cd164 | CD164 antigen |
| chr10 | 41809871 | 41810790 | ENSMUSG00000019813 | Cep57l1 | centrosomal protein 57-like 1 |
| chr10 | 41887148 | 41887861 | ENSMUSG00000038332 | Sesn1 | sestrin 1 |
| chr10 | 42501900 | 42503108 | ENSMUSG00000019804 | Snx3 | sorting nexin 3 |
| chr10 | 42761390 | 42762507 | ENSMUSG00000019802 | Sec63 | SEC63-like (S. cerevisiae) |
| chr10 | 43478566 | 43480214 | ENSMUSG00000038214 | Bend3 | BEN domain containing 3 |
| chr10 | 43540569 | 43541051 | ENSMUSG00000019797 | Mtres1 | mitochondrial transcription rescue factor 1 |
| chr10 | 43578270 | 43580149 | ENSMUSG00000047139 | Cd24a | CD24a antigen |
| chr10 | 44268158 | 44268808 | ENSMUSG00000038160 | Atg5 | autophagy related 5 |
| chr10 | 45066657 | 45067705 | ENSMUSG00000019849 | Prep | prolyl endopeptidase |
| chr10 | 45577716 | 45578551 | ENSMUSG00000038822 | Hace1 | HECT domain and ankyrin repeat containing, E3 ubiquitin protein ligase 1 |
| chr10 | 50592422 | 50593286 | ENSMUSG00000038774 | Ascc3 | activating signal cointegrator 1 complex subunit 3 |
| chr10 | 50894159 | 50895633 | ENSMUSG00000019913 | Sim1 | single-minded family bHLH transcription factor 1 |
| chr10 | 52233500 | 52234309 | ENSMUSG00000019891 | Dcbld1 | discoidin, CUB and LCCL domain containing 1 |
| chr10 | 52381454 | 52382296 | ENSMUSG00000019861 | Gopc | golgi associated PDZ and coiled-coil motif containing |
| chr10 | 52417273 | 52418290 | ENSMUSG00000023068 | Nus1 | NUS1 dehydrodolichyl diphosphate synthase subunit |
| chr10 | 53596472 | 53598010 | ENSMUSG00000019857 | Asf1a | anti-silencing function 1A histone chaperone |
| chr10 | 53629337 | 53630199 | ENSMUSG00000058298 | Mcm9 | minichromosome maintenance 9 homologous recombination repair factor |
| chr10 | 54074557 | 54076068 | ENSMUSG00000097069 | Gm16998 | predicted gene, 16998 |
| chr10 | 58323076 | 58323935 | ENSMUSG00000019920 | Lims1 | LIM and senescent cell antigen-like domains 1 |
| chr10 | 58446644 | 58447549 | ENSMUSG00000003226 | Ranbp2 | RAN binding protein 2 |
| chr10 | 59323067 | 59324007 | ENSMUSG00000019916 | P4ha1 | procollagen-proline, 2-oxoglutarate 4-dioxygenase (proline 4-hydroxylase), alpha 1 polypeptide |
| chr10 | 59615910 | 59616869 | ENSMUSG00000009647 | Mcu | mitochondrial calcium uniporter |
| chr10 | 61146265 | 61148050 | ENSMUSG00000020097 | Sgpl1 | sphingosine phosphate lyase 1 |
| chr10 | 61452043 | 61452905 | ENSMUSG00000020091 | Eif4ebp2 | eukaryotic translation initiation factor 4E binding protein 2 |
| chr10 | 61475625 | 61476374 | ENSMUSG00000037151 | Lrrc20 | leucine rich repeat containing 20 |
| chr10 | 61648330 | 61648992 | ENSMUSG00000020089 | Ppa1 | pyrophosphatase (inorganic) 1 |
| chr10 | 61679924 | 61680779 | ENSMUSG00000020088 | Sar1a | secretion associated Ras related GTPase 1A |
| chr10 | 61695426 | 61696643 | ENSMUSG00000020087 | Tysnd1 | trypsin domain containing 1 |
| chr10 | 62486234 | 62486890 | ENSMUSG00000020078 | Vps26a | VPS26 retromer complex component A |
| chr10 | 62650495 | 62651742 | ENSMUSG00000020076 | Ddx50 | DExD box helicase 50 |
| chr10 | 62725607 | 62726301 | ENSMUSG00000036923 | Stox1 | storkhead box 1 |
| chr10 | 62791696 | 62792582 | ENSMUSG00000020074 | Ccar1 | cell division cycle and apoptosis regulator 1 |
| chr10 | 62946672 | 62947479 | ENSMUSG00000036875 | Dna2 | DNA replication helicase/nuclease 2 |
| chr10 | 63023161 | 63024329 | ENSMUSG00000020069 | Hnrnph3 | heterogeneous nuclear ribonucleoprotein H3 |
| chr10 | 63338234 | 63339341 | ENSMUSG00000020063 | Sirt1 | sirtuin 1 |
| chr10 | 67096148 | 67097384 | ENSMUSG00000037876 | Jmjd1c | jumonji domain containing 1C |
| chr10 | 67126522 | 67127646 | ENSMUSG00000037876 | Jmjd1c | jumonji domain containing 1C |
| chr10 | 67284508 | 67285543 | ENSMUSG00000075000 | Nrbf2 | nuclear receptor binding factor 2 |
| chr10 | 67547833 | 67549359 | ENSMUSG00000057134 | Ado | 2-aminoethanethiol (cysteamine) dioxygenase |
| chr10 | 69212525 | 69213768 | ENSMUSG00000090622 | A930033H14Rik | RIKEN cDNA A930033H14 gene |
| chr10 | 69398295 | 69399658 | ENSMUSG00000069601 | Ank3 | ankyrin 3, epithelial |
| chr10 | 70096364 | 70098208 | ENSMUSG00000048701 | Ccdc6 | coiled-coil domain containing 6 |
| chr10 | 70598857 | 70600010 | ENSMUSG00000037747 | Phyhipl | phytanoyl-CoA hydroxylase interacting protein-like |
| chr10 | 71158927 | 71160143 | ENSMUSG00000014329 | Bicc1 | BicC family RNA binding protein 1 |
| chr10 | 71284498 | 71285576 | ENSMUSG00000019927 | Ube2d1 | ubiquitin-conjugating enzyme E2D 1 |
| chr10 | 71344382 | 71345338 | ENSMUSG00000037710 | Cisd1 | CDGSH iron sulfur domain 1 |
| chr10 | 71347314 | 71348599 | ENSMUSG00000060733 | Ipmk | inositol polyphosphate multikinase |
| chr10 | 75060352 | 75061628 | ENSMUSG00000009681 | Bcr | BCR activator of RhoGEF and GTPase |
| chr10 | 75211803 | 75212469 | ENSMUSG00000033444 | Specc1l | sperm antigen with calponin homology and coiled-coil domains 1-like |
| chr10 | 75932127 | 75932609 | ENSMUSG00000000901 | Mmp11 | matrix metallopeptidase 11 |
| chr10 | 76032510 | 76032970 | ENSMUSG00000049764 | Zfp280b | zinc finger protein 280B |
| chr10 | 76344615 | 76345810 | ENSMUSG00000020231 | Dip2a | disco interacting protein 2 homolog A |
| chr10 | 76961641 | 76962130 | ENSMUSG00000001120 | Pcbp3 | poly(rC) binding protein 3 |
| chr10 | 77258799 | 77259772 | ENSMUSG00000078444 | Gm10941 | predicted gene 10941 |
| chr10 | 77417767 | 77418535 | ENSMUSG00000020262 | Adarb1 | adenosine deaminase, RNA-specific, B1 |
| chr10 | 77515509 | 77516138 | ENSMUSG00000032977 | Fam207a | family with sequence similarity 207, member A |
| chr10 | 77606016 | 77606841 | ENSMUSG00000020265 | Sumo3 | small ubiquitin-like modifier 3 |
| chr10 | 77622129 | 77622678 | ENSMUSG00000009293 | Ube2g2 | ubiquitin-conjugating enzyme E2G 2 |
| chr10 | 77978338 | 77978898 | ENSMUSG00000020284 | Cfap410 | cilia and flagella associated protein 410 |
| chr10 | 78244032 | 78245107 | ENSMUSG00000000374 | Trappc10 | trafficking protein particle complex 10 |
| chr10 | 78351521 | 78352672 | ENSMUSG00000001211 | Agpat3 | 1-acylglycerol-3-phosphate O-acyltransferase 3 |
| chr10 | 78464382 | 78465256 | ENSMUSG00000032788 | Pdxk | pyridoxal (pyridoxine, vitamin B6) kinase |
| chr10 | 79554387 | 79555248 | ENSMUSG00000042570 | Mier2 | MIER family member 2 |
| chr10 | 79681685 | 79683075 | ENSMUSG00000020307 | Cdc34 | cell division cycle 34 |
| chr10 | 79704170 | 79705117 | ENSMUSG00000023175 | Bsg | basigin |
| chr10 | 79716666 | 79717390 | ENSMUSG00000020331 | Hcn2 | hyperpolarization-activated, cyclic nucleotide-gated K+ 2 |
| chr10 | 79766188 | 79767440 | ENSMUSG00000035890 | Rnf126 | ring finger protein 126 |
| chr10 | 79854006 | 79855795 | ENSMUSG00000006498 | Ptbp1 | polypyrimidine tract binding protein 1 |
| chr10 | 79936972 | 79937507 | ENSMUSG00000019564 | Arid3a | AT rich interactive domain 3A (BRIGHT-like) |
| chr10 | 80039410 | 80039901 | ENSMUSG00000004667 | Polr2e | polymerase (RNA) II (DNA directed) polypeptide E |
| chr10 | 80053298 | 80054646 | ENSMUSG00000075706 | Gpx4 | glutathione peroxidase 4 |
| chr10 | 80074864 | 80075576 | ENSMUSG00000035673 | Sbno2 | strawberry notch 2 |
| chr10 | 80138808 | 80139505 | ENSMUSG00000035640 | Cbarp | calcium channel, voltage-dependent, beta subunit associated regulatory protein |
| chr10 | 80167850 | 80168412 | ENSMUSG00000045193 | Cirbp | cold inducible RNA binding protein |
| chr10 | 80226299 | 80227096 | ENSMUSG00000020156 | Pwwp3a | PWWP domain containing 3A, DNA repair factor |
| chr10 | 80386265 | 80387797 | ENSMUSG00000048696 | Mex3d | mex3 RNA binding family member D |
| chr10 | 80398575 | 80400206 | ENSMUSG00000035478 | Mbd3 | methyl-CpG binding domain protein 3 |
| chr10 | 80602769 | 80603310 | ENSMUSG00000113640 | Adat3 | adenosine deaminase, tRNA-specific 3 |
| chr10 | 80754820 | 80755809 | ENSMUSG00000061589 | Dot1l | DOT1-like, histone H3 methyltransferase (S. cerevisiae) |
| chr10 | 81137725 | 81138692 | ENSMUSG00000035011 | Zbtb7a | zinc finger and BTB domain containing 7a |
| chr10 | 81192274 | 81193041 | ENSMUSG00000034974 | Dapk3 | death-associated protein kinase 3 |
| chr10 | 81357335 | 81358196 | ENSMUSG00000034854 | Mfsd12 | major facilitator superfamily domain containing 12 |
| chr10 | 81429773 | 81431331 | ENSMUSG00000055053 | Nfic | nuclear factor I/C |
| chr10 | 81495616 | 81496517 | ENSMUSG00000020238 | Ncln | nicalin |
| chr10 | 81544460 | 81545540 | ENSMUSG00000034781 | Gna11 | guanine nucleotide binding protein, alpha 11 |
| chr10 | 81643301 | 81644063 | ENSMUSG00000054708 | Ankrd24 | ankyrin repeat domain 24 |
| chr10 | 82629667 | 82630304 | ENSMUSG00000034674 | Tdg | thymine DNA glycosylase |
| chr10 | 82763279 | 82764312 | ENSMUSG00000020248 | Nfyb | nuclear transcription factor-Y beta |
| chr10 | 83337127 | 83338102 | ENSMUSG00000034591 | Slc41a2 | solute carrier family 41, member 2 |
| chr10 | 83543713 | 83544442 | ENSMUSG00000034560 | Washc4 | WASH complex subunit 4 |
| chr10 | 84440179 | 84441484 | ENSMUSG00000020032 | Nuak1 | NUAK family, SNF1-like kinase, 1 |
| chr10 | 85102350 | 85103621 | ENSMUSG00000060935 | Tmem263 | transmembrane protein 263 |
| chr10 | 85184218 | 85185653 | ENSMUSG00000020038 | Cry1 | cryptochrome 1 (photolyase-like) |
| chr10 | 85916318 | 85917322 | ENSMUSG00000035529 | Prdm4 | PR domain containing 4 |
| chr10 | 85928416 | 85928799 | ENSMUSG00000085111 | Ascl4 | achaete-scute family bHLH transcription factor 4 |
| chr10 | 86021661 | 86022555 | ENSMUSG00000001786 | Fbxo7 | F-box protein 7 |
| chr10 | 86704778 | 86705848 | ENSMUSG00000044937 | Ttc41 | tetratricopeptide repeat domain 41 |
| chr10 | 86778429 | 86779700 | ENSMUSG00000054027 | Nt5dc3 | 5'-nucleotidase domain containing 3 |
| chr10 | 88378938 | 88379972 | ENSMUSG00000035311 | Gnptab | N-acetylglucosamine-1-phosphate transferase, alpha and beta subunits |
| chr10 | 91082116 | 91083202 | ENSMUSG00000019979 | Apaf1 | apoptotic peptidase activating factor 1 |
| chr10 | 91170601 | 91172118 | ENSMUSG00000019961 | Tmpo | thymopoietin |
| chr10 | 92721974 | 92722597 | ENSMUSG00000019988 | Nedd1 | neural precursor cell expressed, developmentally down-regulated gene 1 |
| chr10 | 93160319 | 93162052 | ENSMUSG00000020015 | Cdk17 | cyclin-dependent kinase 17 |
| chr10 | 94035726 | 94036581 | ENSMUSG00000020021 | Fgd6 | FYVE, RhoGEF and PH domain containing 6 |
| chr10 | 94514595 | 94515695 | ENSMUSG00000020023 | Tmcc3 | transmembrane and coiled coil domains 3 |
| chr10 | 94688248 | 94689267 |  | Cep83os | centrosomal protein 83, opposite strand |
| chr10 | 95415635 | 95417020 | ENSMUSG00000020027 | Socs2 | suppressor of cytokine signaling 2 |
| chr10 | 95417020 | 95417621 | ENSMUSG00000020027 | Socs2 | suppressor of cytokine signaling 2 |
| chr10 | 95514549 | 95515977 | ENSMUSG00000074781 | Ube2n | ubiquitin-conjugating enzyme E2N |
| chr10 | 95563215 | 95564343 | ENSMUSG00000020029 | Nudt4 | nudix (nucleoside diphosphate linked moiety X)-type motif 4 |
| chr10 | 95940328 | 95941519 | ENSMUSG00000036499 | Eea1 | early endosome antigen 1 |
| chr10 | 99106849 | 99108449 | ENSMUSG00000090035 | Galnt4 | polypeptide N-acetylgalactosaminyltransferase 4 |
| chr10 | 105573442 | 105575007 | ENSMUSG00000036019 | Tmtc2 | transmembrane and tetratricopeptide repeat containing 2 |
| chr10 | 108161493 | 108163054 | ENSMUSG00000019907 | Ppp1r12a | protein phosphatase 1, regulatory subunit 12A |
| chr10 | 108331691 | 108333522 | ENSMUSG00000035873 | Pawr | PRKC, apoptosis, WT1, regulator |
| chr10 | 110744849 | 110745899 | ENSMUSG00000020185 | E2f7 | E2F transcription factor 7 |
| chr10 | 111009520 | 111010522 | ENSMUSG00000035798 | Zdhhc17 | zinc finger, DHHC domain containing 17 |
| chr10 | 111473018 | 111473896 | ENSMUSG00000058799 | Nap1l1 | nucleosome assembly protein 1-like 1 |
| chr10 | 111506082 | 111507886 | ENSMUSG00000020205 | Phlda1 | pleckstrin homology like domain, family A, member 1 |
| chr10 | 115250975 | 115251683 | ENSMUSG00000020130 | Tbc1d15 | TBC1 domain family, member 15 |
| chr10 | 115384015 | 115385900 | ENSMUSG00000034163 | Zfc3h1 | zinc finger, C3H1-type containing |
| chr10 | 116580543 | 116582276 | ENSMUSG00000020166 | Cnot2 | CCR4-NOT transcription complex, subunit 2 |
| chr10 | 116949586 | 116950643 | ENSMUSG00000064181 | Rab3ip | RAB3A interacting protein |
| chr10 | 117223995 | 117224608 | ENSMUSG00000020171 | Yeats4 | YEATS domain containing 4 |
| chr10 | 117375932 | 117377100 | ENSMUSG00000055531 | Cpsf6 | cleavage and polyadenylation specific factor 6 |
| chr10 | 117629052 | 117630065 | ENSMUSG00000020183 | Cpm | carboxypeptidase M |
| chr10 | 117709893 | 117710918 | ENSMUSG00000020184 | Mdm2 | transformed mouse 3T3 cell double minute 2 |
| chr10 | 117745632 | 117746470 | ENSMUSG00000060181 | Slc35e3 | solute carrier family 35, member E3 |
| chr10 | 117845002 | 117846455 | ENSMUSG00000052681 | Rap1b | RAS related protein 1b |
| chr10 | 118867973 | 118869628 | ENSMUSG00000028630 | Dyrk2 | dual-specificity tyrosine-(Y)-phosphorylation regulated kinase 2 |
| chr10 | 119239134 | 119240444 | ENSMUSG00000020114 | Cand1 | cullin associated and neddylation disassociated 1 |
| chr10 | 120898364 | 120899338 | ENSMUSG00000051236 | Msrb3 | methionine sulfoxide reductase B3 |
| chr10 | 121351203 | 121351630 | ENSMUSG00000052302 | Tbc1d30 | TBC1 domain family, member 30 |
| chr10 | 121475930 | 121476568 | ENSMUSG00000025795 | Rassf3 | Ras association (RalGDS/AF-6) domain family member 3 |
| chr10 | 121586272 | 121587080 | ENSMUSG00000020115 | Tbk1 | TANK-binding kinase 1 |
| chr10 | 121625827 | 121626782 | ENSMUSG00000034667 | Xpot | exportin, tRNA (nuclear export receptor for tRNAs) |
| chr10 | 121739633 | 121740371 | ENSMUSG00000053684 | BC048403 | cDNA sequence BC048403 |
| chr10 | 122096637 | 122097200 | ENSMUSG00000034620 | Rxylt1 | ribitol xylosyltransferase 1 |
| chr10 | 122985190 | 122986757 | ENSMUSG00000065406 | Mirlet7i | microRNA let7i |
| chr10 | 123196236 | 123197434 | ENSMUSG00000020124 | Usp15 | ubiquitin specific peptidase 15 |
| chr10 | 126900879 | 126901484 | ENSMUSG00000025436 | Atp23 | ATP23 metallopeptidase and ATP synthase assembly factor homolog |
| chr10 | 127063284 | 127064211 | ENSMUSG00000006728 | Cdk4 | cyclin-dependent kinase 4 |
| chr10 | 127164808 | 127166495 | ENSMUSG00000006731 | B4galnt1 | beta-1,4-N-acetyl-galactosaminyl transferase 1 |
| chr10 | 127380279 | 127381315 | ENSMUSG00000025404 | R3hdm2 | R3H domain containing 2 |
| chr10 | 127641963 | 127642757 | ENSMUSG00000002147 | Stat6 | signal transducer and activator of transcription 6 |
| chr10 | 127667710 | 127668497 | ENSMUSG00000025402 | Nab2 | Ngfi-A binding protein 2 |
| chr10 | 127739166 | 127740028 | ENSMUSG00000044617 | Zbtb39 | zinc finger and BTB domain containing 39 |
| chr10 | 128058537 | 128059933 | ENSMUSG00000071072 | Ptges3 | prostaglandin E synthase 3 |
| chr10 | 128092518 | 128093454 | ENSMUSG00000040054 | Baz2a | bromodomain adjacent to zinc finger domain, 2A |
| chr10 | 128193992 | 128195031 | ENSMUSG00000044005 | Gls2 | glutaminase 2 (liver, mitochondrial) |
| chr10 | 128368268 | 128369287 | ENSMUSG00000039914 | Coq10a | coenzyme Q10A |
| chr10 | 128376735 | 128377434 | ENSMUSG00000014498 | Ankrd52 | ankyrin repeat domain 52 |
| chr10 | 128458584 | 128459977 | ENSMUSG00000098957 | Mir8105 | microRNA 8105 |
| chr10 | 128565237 | 128566581 | ENSMUSG00000025364 | Pa2g4 | proliferation-associated 2G4 |
| chr10 | 128743635 | 128744241 | ENSMUSG00000025357 | Dgka | diacylglycerol kinase, alpha |
| chr10 | 128909566 | 128910707 | ENSMUSG00000025351 | Cd63 | CD63 antigen |
| chr11 | 3266135 | 3266705 | ENSMUSG00000020457 | Drg1 | developmentally regulated GTP binding protein 1 |
| chr11 | 3289857 | 3291642 | ENSMUSG00000020453 | Patz1 | POZ (BTB) and AT hook containing zinc finger 1 |
| chr11 | 3648592 | 3649767 | ENSMUSG00000056579 | Tug1 | taurine upregulated gene 1 |
| chr11 | 4186526 | 4187210 | ENSMUSG00000034412 | Tbc1d10a | TBC1 domain family, member 10a |
| chr11 | 4266501 | 4267200 | ENSMUSG00000034394 | Lif | leukemia inhibitory factor |
| chr11 | 5443773 | 5444985 | ENSMUSG00000041961 | Znrf3 | zinc and ring finger 3 |
| chr11 | 5520634 | 5521494 | ENSMUSG00000020484 | Xbp1 | X-box binding protein 1 |
| chr11 | 5707109 | 5707859 | ENSMUSG00000020477 | Mrps24 | mitochondrial ribosomal protein S24 |
| chr11 | 5740730 | 5741523 | ENSMUSG00000049680 | Urgcp | upregulator of cell proliferation |
| chr11 | 5761839 | 5762542 | ENSMUSG00000049680 | Urgcp | upregulator of cell proliferation |
| chr11 | 6415496 | 6416653 | ENSMUSG00000071866 | Ppia | peptidylprolyl isomerase A |
| chr11 | 6444022 | 6444708 | ENSMUSG00000041126 | H2az2 | H2A.Z histone variant 2 |
| chr11 | 6474792 | 6476360 | ENSMUSG00000094483 | Purb | purine rich element binding protein B |
| chr11 | 17159095 | 17160012 | ENSMUSG00000033953 | Ppp3r1 | protein phosphatase 3, regulatory subunit B, alpha isoform (calcineurin B, type I) |
| chr11 | 19919867 | 19920696 | ENSMUSG00000045671 | Spred2 | sprouty-related EVH1 domain containing 2 |
| chr11 | 19924131 | 19925901 | ENSMUSG00000045671 | Spred2 | sprouty-related EVH1 domain containing 2 |
| chr11 | 20112220 | 20113236 | ENSMUSG00000020152 | Actr2 | ARP2 actin-related protein 2 |
| chr11 | 20248841 | 20249731 | ENSMUSG00000044066 | Cep68 | centrosomal protein 68 |
| chr11 | 20542449 | 20543971 | ENSMUSG00000049800 | Sertad2 | SERTA domain containing 2 |
| chr11 | 20631693 | 20632804 | ENSMUSG00000049800 | Sertad2 | SERTA domain containing 2 |
| chr11 | 20740847 | 20741824 | ENSMUSG00000049659 | Aftph | aftiphilin |
| chr11 | 21091230 | 21091969 | ENSMUSG00000020134 | Peli1 | pellino 1 |
| chr11 | 21238668 | 21239665 | ENSMUSG00000020128 | Vps54 | VPS54 GARP complex subunit |
| chr11 | 22858900 | 22860177 | ENSMUSG00000051650 | B3gnt2 | UDP-GlcNAc:betaGal beta-1,3-N-acetylglucosaminyltransferase 2 |
| chr11 | 23067645 | 23068079 | ENSMUSG00000049811 | Fam161a | family with sequence similarity 161, member A |
| chr11 | 23305696 | 23307385 | ENSMUSG00000056342 | Usp34 | ubiquitin specific peptidase 34 |
| chr11 | 23633095 | 23633931 | ENSMUSG00000042208 | 0610010F05Rik | RIKEN cDNA 0610010F05 gene |
| chr11 | 23770149 | 23771443 | ENSMUSG00000020275 | Rel | reticuloendotheliosis oncogene |
| chr11 | 26593200 | 26593833 | ENSMUSG00000064090 | Vrk2 | vaccinia related kinase 2 |
| chr11 | 29373352 | 29374217 | ENSMUSG00000032740 | Ccdc88a | coiled coil domain containing 88A |
| chr11 | 32221122 | 32222528 | ENSMUSG00000020282 | Rhbdf1 | rhomboid 5 homolog 1 |
| chr11 | 32347590 | 32348158 | ENSMUSG00000040711 | Sh3pxd2b | SH3 and PX domains 2B |
| chr11 | 32454598 | 32455839 | ENSMUSG00000044949 | Ubtd2 | ubiquitin domain containing 2 |
| chr11 | 32532769 | 32533842 | ENSMUSG00000020272 | Stk10 | serine/threonine kinase 10 |
| chr11 | 32587872 | 32588330 | ENSMUSG00000020272 | Stk10 | serine/threonine kinase 10 |
| chr11 | 33146892 | 33148396 | ENSMUSG00000057967 | Fgf18 | fibroblast growth factor 18 |
| chr11 | 35979860 | 35980824 | ENSMUSG00000018849 | Wwc1 | WW, C2 and coiled-coil domain containing 1 |
| chr11 | 43681730 | 43682875 | ENSMUSG00000044950 | Pwwp2a | PWWP domain containing 2A |
| chr11 | 45851701 | 45852625 | ENSMUSG00000006169 | Clint1 | clathrin interactor 1 |
| chr11 | 45944394 | 45944976 | ENSMUSG00000044847 | Lsm11 | U7 snRNP-specific Sm-like protein LSM11 |
| chr11 | 48837291 | 48838390 | ENSMUSG00000040350 | Trim7 | tripartite motif-containing 7 |
| chr11 | 49243770 | 49244397 | ENSMUSG00000020346 | Mgat1 | mannoside acetylglucosaminyltransferase 1 |
| chr11 | 50024891 | 50025737 | ENSMUSG00000020376 | Rnf130 | ring finger protein 130 |
| chr11 | 50131068 | 50131652 | ENSMUSG00000036644 | Tbc1d9b | TBC1 domain family, member 9B |
| chr11 | 50224535 | 50226183 | ENSMUSG00000036620 | Mgat4b | mannoside acetylglucosaminyltransferase 4, isoenzyme B |
| chr11 | 50325061 | 50326228 | ENSMUSG00000020368 | Canx | calnexin |
| chr11 | 50430549 | 50431027 | ENSMUSG00000020375 | Rufy1 | RUN and FYVE domain containing 1 |
| chr11 | 51605882 | 51607017 | ENSMUSG00000020358 | Hnrnpab | heterogeneous nuclear ribonucleoprotein A/B |
| chr11 | 51650252 | 51651109 | ENSMUSG00000001053 | N4bp3 | NEDD4 binding protein 3 |
| chr11 | 52004126 | 52004558 | ENSMUSG00000020389 | Cdkl3 | cyclin-dependent kinase-like 3 |
| chr11 | 52098504 | 52099699 | ENSMUSG00000020349 | Ppp2ca | protein phosphatase 2 (formerly 2A), catalytic subunit, alpha isoform |
| chr11 | 52231871 | 52232461 | ENSMUSG00000036309 | Skp1 | S-phase kinase-associated protein 1 |
| chr11 | 52283089 | 52283548 | ENSMUSG00000000782 | Tcf7 | transcription factor 7, T cell specific |
| chr11 | 52360455 | 52361890 | ENSMUSG00000020402 | Vdac1 | voltage-dependent anion channel 1 |
| chr11 | 53350238 | 53351315 | ENSMUSG00000049470 | Aff4 | AF4/FMR2 family, member 4 |
| chr11 | 53769791 | 53770779 | ENSMUSG00000018899 | Irf1 | interferon regulatory factor 1 |
| chr11 | 53891131 | 53891925 | ENSMUSG00000018900 | Slc22a5 | solute carrier family 22 (organic cation transporter), member 5 |
| chr11 | 54437883 | 54438777 | ENSMUSG00000035992 | Fnip1 | folliculin interacting protein 1 |
| chr11 | 54866177 | 54867148 | ENSMUSG00000020267 | Hint1 | histidine triad nucleotide binding protein 1 |
| chr11 | 57801283 | 57802490 | ENSMUSG00000020519 | Sap30l | SAP30-like |
| chr11 | 58307021 | 58307875 | ENSMUSG00000037243 | Zfp692 | zinc finger protein 692 |
| chr11 | 58322823 | 58323512 | ENSMUSG00000049755 | Zfp672 | zinc finger protein 672 |
| chr11 | 58938264 | 58939112 | ENSMUSG00000020496 | Rnf187 | ring finger protein 187 |
| chr11 | 58978015 | 58978798 | ENSMUSG00000020455 | Trim11 | tripartite motif-containing 11 |
| chr11 | 59163284 | 59163860 | ENSMUSG00000049287 | Iba57 | IBA57 homolog, iron-sulfur cluster assembly |
| chr11 | 59227661 | 59228534 | ENSMUSG00000048076 | Arf1 | ADP-ribosylation factor 1 |
| chr11 | 59306616 | 59308337 | ENSMUSG00000000126 | Wnt9a | wingless-type MMTV integration site family, member 9A |
| chr11 | 59662129 | 59663065 | ENSMUSG00000005417 | Mprip | myosin phosphatase Rho interacting protein |
| chr11 | 59847832 | 59848461 | ENSMUSG00000032615 | Nt5m | 5',3'-nucleotidase, mitochondrial |
| chr11 | 60104767 | 60106677 | ENSMUSG00000062115 | Rai1 | retinoic acid induced 1 |
| chr11 | 60352553 | 60353064 | ENSMUSG00000000538 | Tom1l2 | target of myb1-like 2 (chicken) |
| chr11 | 60416875 | 60418022 | ENSMUSG00000042709 | Atpaf2 | ATP synthase mitochondrial F1 complex assembly factor 2 |
| chr11 | 60699561 | 60700169 | ENSMUSG00000020536 | Llgl1 | LLGL1 scribble cell polarity complex component |
| chr11 | 60878192 | 60879284 | ENSMUSG00000043284 | Tmem11 | transmembrane protein 11 |
| chr11 | 60913294 | 60913912 | ENSMUSG00000018931 | Natd1 | N-acetyltransferase domain containing 1 |
| chr11 | 60931436 | 60932530 | ENSMUSG00000042549 | Map2k3os | mitogen-activated protein kinase kinase 3, opposite strand |
| chr11 | 61174496 | 61175522 | ENSMUSG00000042506 | Usp22 | ubiquitin specific peptidase 22 |
| chr11 | 61493918 | 61494590 | ENSMUSG00000001034 | Mapk7 | mitogen-activated protein kinase 7 |
| chr11 | 61578870 | 61579791 | ENSMUSG00000001036 | Epn2 | epsin 2 |
| chr11 | 61854096 | 61855427 | ENSMUSG00000004798 | Ulk2 | unc-51 like kinase 2 |
| chr11 | 61929980 | 61930480 | ENSMUSG00000047804 | Akap10 | A kinase (PRKA) anchor protein 10 |
| chr11 | 62076998 | 62077698 | ENSMUSG00000042331 | Specc1 | sperm antigen with calponin homology and coiled-coil domains 1 |
| chr11 | 62281200 | 62281861 | ENSMUSG00000014243 | Zswim7 | zinc finger SWIM-type containing 7 |
| chr11 | 62456688 | 62458226 | ENSMUSG00000018501 | Ncor1 | nuclear receptor co-repressor 1 |
| chr11 | 62538612 | 62539329 | ENSMUSG00000018509 | Cenpv | centromere protein V |
| chr11 | 62647999 | 62648985 | ENSMUSG00000046417 | Lrrc75a | leucine rich repeat containing 75A |
| chr11 | 65787589 | 65788507 | ENSMUSG00000033352 | Map2k4 | mitogen-activated protein kinase kinase 4 |
| chr11 | 65807059 | 65807595 | ENSMUSG00000018347 | Zkscan6 | zinc finger with KRAB and SCAN domains 6 |
| chr11 | 68852555 | 68853324 | ENSMUSG00000018736 | Ndel1 | nudE neurodevelopment protein 1 like 1 |
| chr11 | 69088432 | 69089257 | ENSMUSG00000020894 | Vamp2 | vesicle-associated membrane protein 2 |
| chr11 | 69117913 | 69118343 | ENSMUSG00000023781 | Hes7 | hes family bHLH transcription factor 7 |
| chr11 | 69122299 | 69123252 | ENSMUSG00000023781 | Hes7 | hes family bHLH transcription factor 7 |
| chr11 | 69632716 | 69633463 | ENSMUSG00000018765 | Fxr2 | fragile X mental retardation, autosomal homolog 2 |
| chr11 | 69681689 | 69682198 | ENSMUSG00000005204 | Senp3 | SUMO/sentrin specific peptidase 3 |
| chr11 | 69804172 | 69804575 | ENSMUSG00000089876 | Tmem102 | transmembrane protein 102 |
| chr11 | 70237467 | 70238333 | ENSMUSG00000020831 | 0610010K14Rik | RIKEN cDNA 0610010K14 gene |
| chr11 | 70432170 | 70432963 | ENSMUSG00000060216 | Arrb2 | arrestin, beta 2 |
| chr11 | 70562633 | 70563305 | ENSMUSG00000020827 | Mink1 | misshapen-like kinase 1 (zebrafish) |
| chr11 | 70563305 | 70563795 | ENSMUSG00000020827 | Mink1 | misshapen-like kinase 1 (zebrafish) |
| chr11 | 70653691 | 70654901 | ENSMUSG00000018293 | Pfn1 | profilin 1 |
| chr11 | 70654901 | 70656219 | ENSMUSG00000018293 | Pfn1 | profilin 1 |
| chr11 | 70764095 | 70764760 | ENSMUSG00000043602 | Zfp3 | zinc finger protein 3 |
| chr11 | 70844688 | 70845488 | ENSMUSG00000020817 | Rabep1 | rabaptin, RAB GTPase binding effector protein 1 |
| chr11 | 70982621 | 70983170 | ENSMUSG00000018446 | C1qbp | complement component 1, q subcomponent binding protein |
| chr11 | 71003757 | 71004567 | ENSMUSG00000040620 | Dhx33 | DEAH (Asp-Glu-Ala-His) box polypeptide 33 |
| chr11 | 72410716 | 72411796 | ENSMUSG00000045667 | Smtnl2 | smoothelin-like 2 |
| chr11 | 72488635 | 72490297 | ENSMUSG00000040447 | Spns2 | spinster homolog 2 |
| chr11 | 72606991 | 72608708 | ENSMUSG00000020794 | Ube2g1 | ubiquitin-conjugating enzyme E2G 1 |
| chr11 | 74637888 | 74638702 | ENSMUSG00000020741 | Cluh | clustered mitochondria (cluA/CLU1) homolog |
| chr11 | 74649286 | 74650019 | ENSMUSG00000020741 | Cluh | clustered mitochondria (cluA/CLU1) homolog |
| chr11 | 74724045 | 74724751 | ENSMUSG00000020745 | Pafah1b1 | platelet-activating factor acetylhydrolase, isoform 1b, subunit 1 |
| chr11 | 74896336 | 74897858 | ENSMUSG00000038351 | Sgsm2 | small G protein signaling modulator 2 |
| chr11 | 74925448 | 74926338 | ENSMUSG00000038290 | Smg6 | Smg-6 homolog, nonsense mediated mRNA decay factor (C. elegans) |
| chr11 | 75679151 | 75680137 | ENSMUSG00000017776 | Crk | v-crk avian sarcoma virus CT10 oncogene homolog |
| chr11 | 75966468 | 75967264 | ENSMUSG00000020847 | Rph3al | rabphilin 3A-like (without C2 domains) |
| chr11 | 76027058 | 76028033 | ENSMUSG00000020846 | Rflnb | refilin B |
| chr11 | 76201790 | 76202471 | ENSMUSG00000069808 | Tlcd3a | TLC domain containing 3A |
| chr11 | 76398440 | 76399475 | ENSMUSG00000020844 | Nxn | nucleoredoxin |
| chr11 | 76494371 | 76494983 | ENSMUSG00000017631 | Abr | active BCR-related gene |
| chr11 | 76846151 | 76847171 | ENSMUSG00000020841 | Cpd | carboxypeptidase D |
| chr11 | 77215830 | 77216720 | ENSMUSG00000037926 | Ssh2 | slingshot protein phosphatase 2 |
| chr11 | 77513073 | 77513599 | ENSMUSG00000044328 | Trp53i13 | transformation related protein 53 inducible protein 13 |
| chr11 | 77606969 | 77608041 | ENSMUSG00000017291 | Taok1 | TAO kinase 1 |
| chr11 | 77763038 | 77764359 | ENSMUSG00000000631 | Myo18a | myosin XVIIIA |
| chr11 | 78031847 | 78032853 | ENSMUSG00000087050 | Dhrs13os | dehydrogenase/reductase (SDR family) member 13, opposite strand |
| chr11 | 78164840 | 78166013 | ENSMUSG00000017386 | Traf4 | TNF receptor associated factor 4 |
| chr11 | 78177909 | 78179098 | ENSMUSG00000099101 | Mir7653 | microRNA 7653 |
| chr11 | 78261324 | 78262274 | ENSMUSG00000010277 | 2610507B11Rik | RIKEN cDNA 2610507B11 gene |
| chr11 | 80080693 | 80081137 | ENSMUSG00000017561 | Crlf3 | cytokine receptor-like factor 3 |
| chr11 | 80208960 | 80209552 | ENSMUSG00000017686 | Rhot1 | ras homolog family member T1 |
| chr11 | 80476532 | 80477293 | ENSMUSG00000048895 | Cdk5r1 | cyclin-dependent kinase 5, regulatory subunit 1 (p35) |
| chr11 | 82780998 | 82781845 | ENSMUSG00000085175 | Gm11423 | predicted gene 11423 |
| chr11 | 82870385 | 82871306 | ENSMUSG00000020696 | Rffl | ring finger and FYVE like domain containing protein |
| chr11 | 83964245 | 83964843 | ENSMUSG00000034940 | Synrg | synergin, gamma |
| chr11 | 84067872 | 84068804 | ENSMUSG00000018648 | Dusp14 | dual specificity phosphatase 14 |
| chr11 | 84129191 | 84129884 | ENSMUSG00000020532 | Acaca | acetyl-Coenzyme A carboxylase alpha |
| chr11 | 84179287 | 84180589 | ENSMUSG00000020532 | Acaca | acetyl-Coenzyme A carboxylase alpha |
| chr11 | 84513124 | 84513679 | ENSMUSG00000018697 | Aatf | apoptosis antagonizing transcription factor |
| chr11 | 84818725 | 84819604 | ENSMUSG00000018405 | Mrm1 | mitochondrial rRNA methyltransferase 1 |
| chr11 | 84869374 | 84871087 | ENSMUSG00000020530 | Ggnbp2 | gametogenetin binding protein 2 |
| chr11 | 84879764 | 84880836 | ENSMUSG00000020527 | Myo19 | myosin XIX |
| chr11 | 85234378 | 85235219 | ENSMUSG00000085628 | Appbp2os | amyloid beta precursor protein (cytoplasmic tail) binding protein 2, opposite strand |
| chr11 | 86757262 | 86758344 | ENSMUSG00000047126 | Cltc | clathrin, heavy polypeptide (Hc) |
| chr11 | 86992967 | 86994023 | ENSMUSG00000018427 | Ypel2 | yippee like 2 |
| chr11 | 87126815 | 87127692 | ENSMUSG00000018548 | Trim37 | tripartite motif-containing 37 |
| chr11 | 87986552 | 87987833 | ENSMUSG00000020483 | Dynll2 | dynein light chain LC8-type 2 |
| chr11 | 88098994 | 88099919 | ENSMUSG00000018378 | Cuedc1 | CUE domain containing 1 |
| chr11 | 88717739 | 88718921 |  | C030037D09Rik | RIKEN cDNA C030037D09 gene |
| chr11 | 88863965 | 88864932 | ENSMUSG00000018428 | Akap1 | A kinase (PRKA) anchor protein 1 |
| chr11 | 93884958 | 93886567 | ENSMUSG00000059474 | Mbtd1 | mbt domain containing 1 |
| chr11 | 93995813 | 93996799 | ENSMUSG00000020859 | Spag9 | sperm associated antigen 9 |
| chr11 | 94210176 | 94212175 | ENSMUSG00000037573 | Tob1 | transducer of ErbB-2.1 |
| chr11 | 94327895 | 94328613 | ENSMUSG00000020864 | Ankrd40 | ankyrin repeat domain 40 |
| chr11 | 94548581 | 94549417 | ENSMUSG00000039096 | Rsad1 | radical S-adenosyl methionine domain containing 1 |
| chr11 | 94676978 | 94677602 | ENSMUSG00000020868 | Xylt2 | xylosyltransferase II |
| chr11 | 95145886 | 95146380 | ENSMUSG00000020871 | Dlx4 | distal-less homeobox 4 |
| chr11 | 95712595 | 95713602 | ENSMUSG00000050860 | Phospho1 | phosphatase, orphan 1 |
| chr11 | 95823966 | 95824704 | ENSMUSG00000050860 | Phospho1 | phosphatase, orphan 1 |
| chr11 | 95859252 | 95859856 | ENSMUSG00000086015 | 4833417C18Rik | RIKEN cDNA 4833417C18 gene |
| chr11 | 96271056 | 96272862 | ENSMUSG00000020875 | Hoxb9 | homeobox B9 |
| chr11 | 96298927 | 96299671 | ENSMUSG00000000690 | Hoxb6 | homeobox B6 |
| chr11 | 96352670 | 96353641 | ENSMUSG00000075588 | Hoxb2 | homeobox B2 |
| chr11 | 96789010 | 96789758 | ENSMUSG00000018666 | Cbx1 | chromobox 1 |
| chr11 | 97279718 | 97280954 | ENSMUSG00000001441 | Npepps | aminopeptidase puromycin sensitive |
| chr11 | 97362150 | 97363295 | ENSMUSG00000038485 | Socs7 | suppressor of cytokine signaling 7 |
| chr11 | 97659973 | 97660719 | ENSMUSG00000038437 | Mllt6 | myeloid/lymphoid or mixed-lineage leukemia; translocated to, 6 |
| chr11 | 97699887 | 97700676 | ENSMUSG00000018537 | Pcgf2 | polycomb group ring finger 2 |
| chr11 | 98338446 | 98339191 | ENSMUSG00000038255 | Neurod2 | neurogenic differentiation 2 |
| chr11 | 98412210 | 98413107 | ENSMUSG00000062312 | Erbb2 | erb-b2 receptor tyrosine kinase 2 |
| chr11 | 98809599 | 98810257 | ENSMUSG00000075576 | Gm12359 | predicted gene 12359 |
| chr11 | 100850126 | 100851152 | ENSMUSG00000020919 | Stat5b | signal transducer and activator of transcription 5B |
| chr11 | 100969104 | 100970752 | ENSMUSG00000004044 | Cavin1 | caveolae associated 1 |
| chr11 | 101119881 | 101120491 | ENSMUSG00000035198 | Tubg1 | tubulin, gamma 1 |
| chr11 | 101464979 | 101466448 | ENSMUSG00000001313 | Rnd2 | Rho family GTPase 2 |
| chr11 | 101784351 | 101785337 | ENSMUSG00000017724 | Etv4 | ets variant 4 |
| chr11 | 102276381 | 102276830 | ENSMUSG00000034768 | Asb16 | ankyrin repeat and SOCS box-containing 16 |
| chr11 | 102604398 | 102605482 | ENSMUSG00000050288 | Fzd2 | frizzled class receptor 2 |
| chr11 | 102818488 | 102819615 | ENSMUSG00000034520 | Gjc1 | gap junction protein, gamma 1 |
| chr11 | 103266775 | 103267373 | ENSMUSG00000020941 | Map3k14 | mitogen-activated protein kinase kinase kinase 14 |
| chr11 | 103311933 | 103312453 | ENSMUSG00000034255 | Arhgap27 | Rho GTPase activating protein 27 |
| chr11 | 104441472 | 104442997 | ENSMUSG00000018412 | Kansl1 | KAT8 regulatory NSL complex subunit 1 |
| chr11 | 104468717 | 104469366 | ENSMUSG00000018412 | Kansl1 | KAT8 regulatory NSL complex subunit 1 |
| chr11 | 105943699 | 105944609 | ENSMUSG00000019590 | Cyb561 | cytochrome b-561 |
| chr11 | 106083854 | 106085350 | ENSMUSG00000020700 | Map3k3 | mitogen-activated protein kinase kinase kinase 3 |
| chr11 | 106487129 | 106488177 | ENSMUSG00000020715 | Ern1 | endoplasmic reticulum (ER) to nucleus signalling 1 |
| chr11 | 107027689 | 107028351 | ENSMUSG00000078607 | 1810010H24Rik | RIKEN cDNA 1810010H24 gene |
| chr11 | 107469163 | 107470970 | ENSMUSG00000040430 | Pitpnc1 | phosphatidylinositol transfer protein, cytoplasmic 1 |
| chr11 | 108342826 | 108344302 | ENSMUSG00000000049 | Apoh | apolipoprotein H |
| chr11 | 109362296 | 109363629 | ENSMUSG00000056687 | Gm11696 | predicted gene 11696 |
| chr11 | 109473003 | 109474107 | ENSMUSG00000041920 | Slc16a6 | solute carrier family 16 (monocarboxylic acid transporters), member 6 |
| chr11 | 109650803 | 109651867 | ENSMUSG00000020612 | Prkar1a | protein kinase, cAMP dependent regulatory, type I, alpha |
| chr11 | 112782438 | 112783141 | ENSMUSG00000000567 | Sox9 | SRY (sex determining region Y)-box 9 |
| chr11 | 113750830 | 113752044 | ENSMUSG00000041598 | Cdc42ep4 | CDC42 effector protein (Rho GTPase binding) 4 |
| chr11 | 115187239 | 115187680 | ENSMUSG00000045980 | Tmem104 | transmembrane protein 104 |
| chr11 | 115381717 | 115382429 | ENSMUSG00000050910 | Cdr2l | cerebellar degeneration-related protein 2-like |
| chr11 | 115535266 | 115536648 | ENSMUSG00000020738 | Sumo2 | small ubiquitin-like modifier 2 |
| chr11 | 115603414 | 115604668 | ENSMUSG00000020740 | Gga3 | golgi associated, gamma adaptin ear containing, ARF binding protein 3 |
| chr11 | 115814606 | 115815184 | ENSMUSG00000020781 | Tsen54 | tRNA splicing endonuclease subunit 54 |
| chr11 | 115974462 | 115975643 | ENSMUSG00000020758 | Itgb4 | integrin beta 4 |
| chr11 | 116108736 | 116110437 | ENSMUSG00000020773 | Trim47 | tripartite motif-containing 47 |
| chr11 | 116130418 | 116131098 | ENSMUSG00000054517 | Trim65 | tripartite motif-containing 65 |
| chr11 | 116198610 | 116199206 | ENSMUSG00000020777 | Acox1 | acyl-Coenzyme A oxidase 1, palmitoyl |
| chr11 | 116412239 | 116413386 | ENSMUSG00000052949 | Rnf157 | ring finger protein 157 |
| chr11 | 116433946 | 116434901 | ENSMUSG00000050628 | Ubald2 | UBA-like domain containing 2 |
| chr11 | 116489719 | 116490511 | ENSMUSG00000015869 | Prpsap1 | phosphoribosyl pyrophosphate synthetase-associated protein 1 |
| chr11 | 116842653 | 116843945 | ENSMUSG00000056962 | Jmjd6 | jumonji domain containing 6 |
| chr11 | 116852047 | 116853824 | ENSMUSG00000034120 | Srsf2 | serine and arginine-rich splicing factor 2 |
| chr11 | 117114793 | 117115708 | ENSMUSG00000020823 | Sec14l1 | SEC14-like lipid binding 1 |
| chr11 | 117809661 | 117810794 | ENSMUSG00000048277 | Syngr2 | synaptogyrin 2 |
| chr11 | 117968858 | 117969727 | ENSMUSG00000053113 | Socs3 | suppressor of cytokine signaling 3 |
| chr11 | 117986649 | 117987435 | ENSMUSG00000017715 | Pgs1 | phosphatidylglycerophosphate synthase 1 |
| chr11 | 118247764 | 118249124 | ENSMUSG00000017132 | Cyth1 | cytohesin 1 |
| chr11 | 118289809 | 118290490 | ENSMUSG00000033909 | Usp36 | ubiquitin specific peptidase 36 |
| chr11 | 118354803 | 118355653 | ENSMUSG00000017466 | Timp2 | tissue inhibitor of metalloproteinase 2 |
| chr11 | 118418546 | 118419257 | ENSMUSG00000025575 | Cant1 | calcium activated nucleotidase 1 |
| chr11 | 119022750 | 119023665 | ENSMUSG00000025577 | Cbx2 | chromobox 2 |
| chr11 | 119085466 | 119086631 | ENSMUSG00000039989 | Cbx4 | chromobox 4 |
| chr11 | 119942325 | 119943831 | ENSMUSG00000025372 | Baiap2 | brain-specific angiogenesis inhibitor 1-associated protein 2 |
| chr11 | 120189266 | 120190216 |  | 2810410L24Rik | RIKEN cDNA 2810410L24 gene |
| chr11 | 120230818 | 120231429 |  | 2900052L18Rik | RIKEN cDNA 2900052L18 gene |
| chr11 | 120236309 | 120237945 | ENSMUSG00000039741 | Bahcc1 | BAH domain and coiled-coil containing 1 |
| chr11 | 120261429 | 120262190 | ENSMUSG00000039741 | Bahcc1 | BAH domain and coiled-coil containing 1 |
| chr11 | 120572553 | 120573658 | ENSMUSG00000025130 | P4hb | prolyl 4-hydroxylase, beta polypeptide |
| chr11 | 120581067 | 120582068 | ENSMUSG00000025132 | Arhgdia | Rho GDP dissociation inhibitor (GDI) alpha |
| chr11 | 120597665 | 120598887 | ENSMUSG00000025134 | Alyref | Aly/REF export factor |
| chr11 | 120632760 | 120633833 | ENSMUSG00000051510 | Mafg | v-maf musculoaponeurotic fibrosarcoma oncogene family, protein G (avian) |
| chr11 | 120672873 | 120673490 | ENSMUSG00000025142 | Aspscr1 | alveolar soft part sarcoma chromosome region, candidate 1 (human) |
| chr11 | 120783786 | 120784923 | ENSMUSG00000025158 | Rfng | RFNG O-fucosylpeptide 3-beta-N-acetylglucosaminyltransferase |
| chr11 | 120822944 | 120824931 | ENSMUSG00000025153 | Fasn | fatty acid synthase |
| chr11 | 121259602 | 121260769 | ENSMUSG00000039275 | Foxk2 | forkhead box K2 |
| chr11 | 121353748 | 121354674 | ENSMUSG00000025173 | Wdr45b | WD repeat domain 45B |
| chr11 | 121672764 | 121673500 | ENSMUSG00000046605 | B3gntl1 | UDP-GlcNAc:betaGal beta-1,3-N-acetylglucosaminyltransferase-like 1 |
| chr11 | 121702160 | 121703218 | ENSMUSG00000039208 | Metrnl | meteorin, glial cell differentiation regulator-like |
| chr12 | 3309061 | 3310465 | ENSMUSG00000020671 | Rab10 | RAB10, member RAS oncogene family |
| chr12 | 3426539 | 3427741 | ENSMUSG00000071456 | 1110002L01Rik | RIKEN cDNA 1110002L01 gene |
| chr12 | 3571830 | 3573449 | ENSMUSG00000071454 | Dtnb | dystrobrevin, beta |
| chr12 | 3806140 | 3808045 | ENSMUSG00000020661 | Dnmt3a | DNA methyltransferase 3A |
| chr12 | 4476181 | 4477482 | ENSMUSG00000020647 | Ncoa1 | nuclear receptor coactivator 1 |
| chr12 | 4592594 | 4593810 | ENSMUSG00000020640 | Itsn2 | intersectin 2 |
| chr12 | 4906966 | 4907865 | ENSMUSG00000020634 | Ubxn2a | UBX domain protein 2A |
| chr12 | 4917260 | 4918326 | ENSMUSG00000052812 | Atad2b | ATPase family, AAA domain containing 2B |
| chr12 | 8498774 | 8500442 | ENSMUSG00000054364 | Rhob | ras homolog family member B |
| chr12 | 8673836 | 8675152 | ENSMUSG00000020594 | Pum2 | pumilio RNA-binding family member 2 |
| chr12 | 8771350 | 8773030 | ENSMUSG00000020592 | Sdc1 | syndecan 1 |
| chr12 | 8773030 | 8773360 | ENSMUSG00000020592 | Sdc1 | syndecan 1 |
| chr12 | 9029895 | 9030369 | ENSMUSG00000066637 | Ttc32 | tetratricopeptide repeat domain 32 |
| chr12 | 11265410 | 11266785 | ENSMUSG00000051235 | Gen1 | GEN1, Holliday junction 5' flap endonuclease |
| chr12 | 16894109 | 16895807 | ENSMUSG00000020580 | Rock2 | Rho-associated coiled-coil containing protein kinase 2 |
| chr12 | 16999747 | 17000473 | ENSMUSG00000045679 | Pqlc3 | PQ loop repeat containing |
| chr12 | 17266454 | 17267072 | ENSMUSG00000020571 | Pdia6 | protein disulfide isomerase associated 6 |
| chr12 | 17544513 | 17545491 | ENSMUSG00000011179 | Odc1 | ornithine decarboxylase, structural 1 |
| chr12 | 17881593 | 17882977 | ENSMUSG00000071379 | Hpcal1 | hippocalcin-like 1 |
| chr12 | 20920348 | 20921436 | ENSMUSG00000052632 | Asap2 | ArfGAP with SH3 domain, ankyrin repeat and PH domain 2 |
| chr12 | 21111309 | 21112828 | ENSMUSG00000052632 | Asap2 | ArfGAP with SH3 domain, ankyrin repeat and PH domain 2 |
| chr12 | 21373121 | 21374114 | ENSMUSG00000052593 | Adam17 | a disintegrin and metallopeptidase domain 17 |
| chr12 | 21416941 | 21418051 | ENSMUSG00000076432 | Ywhaq | tyrosine 3-monooxygenase/tryptophan 5-monooxygenase activation protein theta |
| chr12 | 24571816 | 24572768 | ENSMUSG00000020656 | Grhl1 | grainyhead like transcription factor 1 |
| chr12 | 24650880 | 24652157 | ENSMUSG00000020653 | Klf11 | Kruppel-like factor 11 |
| chr12 | 24707958 | 24708732 | ENSMUSG00000020649 | Rrm2 | ribonucleotide reductase M2 |
| chr12 | 24831286 | 24832277 | ENSMUSG00000020646 | Mboat2 | membrane bound O-acyltransferase domain containing 2 |
| chr12 | 25098261 | 25099913 | ENSMUSG00000020644 | Id2 | inhibitor of DNA binding 2 |
| chr12 | 27341224 | 27342497 | ENSMUSG00000063632 | Sox11 | SRY (sex determining region Y)-box 11 |
| chr12 | 28649489 | 28649982 | ENSMUSG00000020630 | Rnaseh1 | ribonuclease H1 |
| chr12 | 29937836 | 29939138 | ENSMUSG00000020674 | Pxdn | peroxidasin |
| chr12 | 31634166 | 31635067 | ENSMUSG00000020650 | Bcap29 | B cell receptor associated protein 29 |
| chr12 | 32060536 | 32061691 | ENSMUSG00000002997 | Prkar2b | protein kinase, cAMP dependent regulatory, type II beta |
| chr12 | 32378652 | 32380012 | ENSMUSG00000090946 | Ccdc71l | coiled-coil domain containing 71 like |
| chr12 | 32819863 | 32821188 | ENSMUSG00000020572 | Nampt | nicotinamide phosphoribosyltransferase |
| chr12 | 32953704 | 32954745 | ENSMUSG00000020570 | Sypl | synaptophysin-like protein |
| chr12 | 33957209 | 33958876 | ENSMUSG00000035799 | Twist1 | twist basic helix-loop-helix transcription factor 1 |
| chr12 | 35046668 | 35048273 | ENSMUSG00000020590 | Snx13 | sorting nexin 13 |
| chr12 | 35533996 | 35535248 | ENSMUSG00000019256 | Ahr | aryl-hydrocarbon receptor |
| chr12 | 40037119 | 40038439 | ENSMUSG00000047446 | Arl4a | ADP-ribosylation factor-like 4A |
| chr12 | 40133503 | 40134350 | ENSMUSG00000002565 | Scin | scinderin |
| chr12 | 40222227 | 40223474 | ENSMUSG00000089371 | Mir1938 | microRNA 1938 |
| chr12 | 40445343 | 40446669 | ENSMUSG00000035954 | Dock4 | dedicator of cytokinesis 4 |
| chr12 | 44269026 | 44269544 | ENSMUSG00000036257 | Pnpla8 | patatin-like phospholipase domain containing 8 |
| chr12 | 45073722 | 45074922 | ENSMUSG00000046314 | Stxbp6 | syntaxin binding protein 6 (amisyn) |
| chr12 | 46818534 | 46819848 | ENSMUSG00000021047 | Nova1 | NOVA alternative splicing regulator 1 |
| chr12 | 51347920 | 51348625 | ENSMUSG00000035293 | G2e3 | G2/M-phase specific E3 ubiquitin ligase |
| chr12 | 51690796 | 51692521 | ENSMUSG00000020955 | Ap4s1 | adaptor-related protein complex AP-4, sigma 1 |
| chr12 | 51828488 | 51830199 | ENSMUSG00000035247 | Hectd1 | HECT domain E3 ubiquitin protein ligase 1 |
| chr12 | 51970834 | 51971528 | ENSMUSG00000035181 | Heatr5a | HEAT repeat containing 5A |
| chr12 | 52006163 | 52006662 | ENSMUSG00000020956 | Dtd2 | D-tyrosyl-tRNA deacylase 2 |
| chr12 | 54655994 | 54656950 | ENSMUSG00000044408 | Sptssa | serine palmitoyltransferase, small subunit A |
| chr12 | 54862138 | 54863149 | ENSMUSG00000062929 | Cfl2 | cofilin 2, muscle |
| chr12 | 54984968 | 54986398 | ENSMUSG00000035021 | Baz1a | bromodomain adjacent to zinc finger domain 1A |
| chr12 | 55491775 | 55492837 | ENSMUSG00000021025 | Nfkbia | nuclear factor of kappa light polypeptide gene enhancer in B cells inhibitor, alpha |
| chr12 | 55820428 | 55821198 | ENSMUSG00000021027 | Ralgapa1 | Ral GTPase activating protein, alpha subunit 1 |
| chr12 | 55836251 | 55837028 | ENSMUSG00000012076 | Brms1l | breast cancer metastasis-suppressor 1-like |
| chr12 | 57230141 | 57231079 | ENSMUSG00000079104 | Prps1l3 | phosphoribosyl pyrophosphate synthetase 1-like 3 |
| chr12 | 57545391 | 57546683 | ENSMUSG00000035451 | Foxa1 | forkhead box A1 |
| chr12 | 57546683 | 57547250 | ENSMUSG00000035451 | Foxa1 | forkhead box A1 |
| chr12 | 59130761 | 59131921 | ENSMUSG00000021000 | Mia2 | MIA SH3 domain ER export factor 2 |
| chr12 | 59218684 | 59220215 | ENSMUSG00000035329 | Fbxo33 | F-box protein 33 |
| chr12 | 64917660 | 64918313 | ENSMUSG00000047227 | Gm527 | predicted gene 527 |
| chr12 | 65036139 | 65036821 | ENSMUSG00000035597 | Prpf39 | pre-mRNA processing factor 39 |
| chr12 | 65073363 | 65074331 | ENSMUSG00000020949 | Fkbp3 | FK506 binding protein 3 |
| chr12 | 65075445 | 65076081 | ENSMUSG00000055884 | Fancm | Fanconi anemia, complementation group M |
| chr12 | 69158637 | 69159317 | ENSMUSG00000034892 | Rps29 | ribosomal protein S29 |
| chr12 | 69183658 | 69185042 | ENSMUSG00000043998 | Mgat2 | mannoside acetylglucosaminyltransferase 2 |
| chr12 | 69196500 | 69198392 | ENSMUSG00000020973 | Dnaaf2 | dynein, axonemal assembly factor 2 |
| chr12 | 69296520 | 69297427 | ENSMUSG00000020978 | Klhdc2 | kelch domain containing 2 |
| chr12 | 69371428 | 69373369 | ENSMUSG00000044147 | Arf6 | ADP-ribosylation factor 6 |
| chr12 | 69582675 | 69583099 | ENSMUSG00000049882 | Vcpkmt | valosin containing protein lysine (K) methyltransferase |
| chr12 | 69681077 | 69682423 | ENSMUSG00000034801 | Sos2 | SOS Ras/Rho guanine nucleotide exchange factor 2 |
| chr12 | 69724536 | 69725386 | ENSMUSG00000020988 | L2hgdh | L-2-hydroxyglutarate dehydrogenase |
| chr12 | 69892373 | 69893478 | ENSMUSG00000034761 | Map4k5 | mitogen-activated protein kinase kinase kinase kinase 5 |
| chr12 | 69986303 | 69987599 | ENSMUSG00000021067 | Sav1 | salvador family WW domain containing 1 |
| chr12 | 70111115 | 70112183 | ENSMUSG00000021068 | Nin | ninein |
| chr12 | 70825367 | 70826372 | ENSMUSG00000048285 | Frmd6 | FERM domain containing 6 |
| chr12 | 70974583 | 70975308 | ENSMUSG00000060073 | Psma3 | proteasome subunit alpha 3 |
| chr12 | 71016002 | 71017129 | ENSMUSG00000048118 | Arid4a | AT rich interactive domain 4A (RBP1-like) |
| chr12 | 72760351 | 72762153 | ENSMUSG00000021096 | Ppm1a | protein phosphatase 1A, magnesium dependent, alpha isoform |
| chr12 | 73111850 | 73113123 | ENSMUSG00000034460 | Six4 | sine oculis-related homeobox 4 |
| chr12 | 73907384 | 73908834 | ENSMUSG00000021109 | Hif1a | hypoxia inducible factor 1, alpha subunit |
| chr12 | 75595252 | 75596808 | ENSMUSG00000021051 | Ppp2r5e | protein phosphatase 2, regulatory subunit B', epsilon |
| chr12 | 75668890 | 75670097 | ENSMUSG00000045690 | Wdr89 | WD repeat domain 89 |
| chr12 | 75734819 | 75736043 | ENSMUSG00000021054 | Sgpp1 | sphingosine-1-phosphate phosphatase 1 |
| chr12 | 75817895 | 75819880 | ENSMUSG00000063450 | Syne2 | spectrin repeat containing, nuclear envelope 2 |
| chr12 | 76533010 | 76534292 | ENSMUSG00000092738 | Mir5135 | microRNA 5135 |
| chr12 | 76927084 | 76927865 | ENSMUSG00000059436 | Max | Max protein |
| chr12 | 76961517 | 76962804 | ENSMUSG00000059436 | Max | Max protein |
| chr12 | 77237899 | 77239811 | ENSMUSG00000021065 | Fut8 | fucosyltransferase 8 |
| chr12 | 78748474 | 78749646 | ENSMUSG00000021112 | Mpp5 | membrane protein, palmitoylated 5 (MAGUK p55 subfamily member 5) |
| chr12 | 79028983 | 79029715 | ENSMUSG00000060716 | Plekhh1 | pleckstrin homology domain containing, family H (with MyTH4 domain) member 1 |
| chr12 | 80132336 | 80132957 |  | 2310015A10Rik | RIKEN cDNA 2310015A10 gene |
| chr12 | 80436043 | 80437047 | ENSMUSG00000049106 | Dcaf5 | DDB1 and CUL4 associated factor 5 |
| chr12 | 80790284 | 80791443 | ENSMUSG00000021133 | Susd6 | sushi domain containing 6 |
| chr12 | 80945522 | 80946480 | ENSMUSG00000021134 | Srsf5 | serine and arginine-rich splicing factor 5 |
| chr12 | 81484689 | 81485279 | ENSMUSG00000091803 | Cox16 | cytochrome c oxidase assembly protein 16 |
| chr12 | 81630480 | 81632303 | ENSMUSG00000042734 | Ttc9 | tetratricopeptide repeat domain 9 |
| chr12 | 81859762 | 81861233 | ENSMUSG00000021140 | Pcnx | pecanex homolog |
| chr12 | 82169201 | 82170432 | ENSMUSG00000042700 | Sipa1l1 | signal-induced proliferation-associated 1 like 1 |
| chr12 | 83987762 | 83988620 | ENSMUSG00000021226 | Acot2 | acyl-CoA thioesterase 2 |
| chr12 | 84192424 | 84194394 | ENSMUSG00000042507 | Mideas | mitotic deacetylase associated SANT domain protein |
| chr12 | 84218417 | 84219158 | ENSMUSG00000042507 | Mideas | mitotic deacetylase associated SANT domain protein |
| chr12 | 84408509 | 84409191 | ENSMUSG00000021236 | Entpd5 | ectonucleoside triphosphate diphosphohydrolase 5 |
| chr12 | 85110670 | 85111444 | ENSMUSG00000004789 | Dlst | dihydrolipoamide S-succinyltransferase (E2 component of 2-oxo-glutarate complex) |
| chr12 | 85338800 | 85339518 | ENSMUSG00000034290 | Nek9 | NIMA (never in mitosis gene a)-related expressed kinase 9 |
| chr12 | 85374055 | 85374807 | ENSMUSG00000021248 | Tmed10 | transmembrane p24 trafficking protein 10 |
| chr12 | 85472009 | 85473047 | ENSMUSG00000021250 | Fos | FBJ osteosarcoma oncogene |
| chr12 | 85473047 | 85474771 | ENSMUSG00000021250 | Fos | FBJ osteosarcoma oncogene |
| chr12 | 85485900 | 85486450 | ENSMUSG00000021250 | Fos | FBJ osteosarcoma oncogene |
| chr12 | 85598638 | 85600085 | ENSMUSG00000034271 | Jdp2 | Jun dimerization protein 2 |
| chr12 | 85824412 | 85825411 | ENSMUSG00000012609 | Ttll5 | tubulin tyrosine ligase-like family, member 5 |
| chr12 | 86678623 | 86679209 | ENSMUSG00000021256 | Vash1 | vasohibin 1 |
| chr12 | 86725756 | 86726614 | ENSMUSG00000021257 | Angel1 | angel homolog 1 |
| chr12 | 86887996 | 86889277 | ENSMUSG00000034168 | Irf2bpl | interferon regulatory factor 2 binding protein-like |
| chr12 | 86946747 | 86948292 | ENSMUSG00000034157 | Cipc | CLOCK interacting protein, circadian |
| chr12 | 87199758 | 87200343 | ENSMUSG00000090812 | Samd15 | sterile alpha motif domain containing 15 |
| chr12 | 91588902 | 91590709 | ENSMUSG00000020962 | Gtf2a1 | general transcription factor II A, 1 |
| chr12 | 98258848 | 98259394 | ENSMUSG00000021003 | Galc | galactosylceramidase |
| chr12 | 98574478 | 98575185 | ENSMUSG00000033854 | Kcnk10 | potassium channel, subfamily K, member 10 |
| chr12 | 98737043 | 98737829 | ENSMUSG00000021009 | Ptpn21 | protein tyrosine phosphatase, non-receptor type 21 |
| chr12 | 98746773 | 98747446 | ENSMUSG00000021012 | Zc3h14 | zinc finger CCCH type containing 14 |
| chr12 | 98900891 | 98901512 | ENSMUSG00000051166 | Eml5 | echinoderm microtubule associated protein like 5 |
| chr12 | 99391997 | 99393584 | ENSMUSG00000033713 | Foxn3 | forkhead box N3 |
| chr12 | 100199205 | 100201060 | ENSMUSG00000001175 | Calm1 | calmodulin 1 |
| chr12 | 100520245 | 100521085 | ENSMUSG00000033530 | Ttc7b | tetratricopeptide repeat domain 7B |
| chr12 | 101028440 | 101029559 | ENSMUSG00000021182 | Ccdc88c | coiled-coil domain containing 88C |
| chr12 | 101082479 | 101084188 | ENSMUSG00000097121 | D130020L05Rik | RIKEN cDNA D130020L05 gene |
| chr12 | 101717981 | 101718896 | ENSMUSG00000021187 | Tc2n | tandem C2 domains, nuclear |
| chr12 | 102282821 | 102284063 | ENSMUSG00000044456 | Rin3 | Ras and Rab interactor 3 |
| chr12 | 102468928 | 102469619 | ENSMUSG00000021192 | Golga5 | golgi autoantigen, golgin subfamily a, 5 |
| chr12 | 102704450 | 102705512 | ENSMUSG00000057963 | Itpk1 | inositol 1,3,4-triphosphate 5/6 kinase |
| chr12 | 102743437 | 102744202 | ENSMUSG00000046675 | Tmem251 | transmembrane protein 251 |
| chr12 | 102877361 | 102878699 | ENSMUSG00000041702 | Btbd7 | BTB (POZ) domain containing 7 |
| chr12 | 104751272 | 104752458 | ENSMUSG00000041415 | Dicer1 | dicer 1, ribonuclease type III |
| chr12 | 105032027 | 105033424 | ENSMUSG00000113722 | Snhg10 | small nucleolar RNA host gene 10 |
| chr12 | 105684564 | 105685891 | ENSMUSG00000044715 | Gskip | GSK3B interacting protein |
| chr12 | 105784222 | 105785410 | ENSMUSG00000021111 | Papola | poly (A) polymerase alpha |
| chr12 | 108178359 | 108179513 | ENSMUSG00000056770 | Setd3 | SET domain containing 3 |
| chr12 | 108274610 | 108275803 | ENSMUSG00000084883 | Ccdc85c | coiled-coil domain containing 85C |
| chr12 | 108422928 | 108423574 | ENSMUSG00000058070 | Eml1 | echinoderm microtubule associated protein like 1 |
| chr12 | 108554604 | 108555670 |  | Gm16596 | predicted gene, 16596 |
| chr12 | 108835453 | 108836585 | ENSMUSG00000048856 | Slc25a47 | solute carrier family 25, member 47 |
| chr12 | 110601130 | 110602416 | ENSMUSG00000018707 | Dync1h1 | dynein cytoplasmic 1 heavy chain 1 |
| chr12 | 110695714 | 110696778 | ENSMUSG00000021270 | Hsp90aa1 | heat shock protein 90, alpha (cytosolic), class A member 1 |
| chr12 | 110850093 | 110850834 | ENSMUSG00000021271 | Zfp839 | zinc finger protein 839 |
| chr12 | 110889171 | 110889745 | ENSMUSG00000021275 | Tecpr2 | tectonin beta-propeller repeat containing 2 |
| chr12 | 110978483 | 110979427 | ENSMUSG00000037904 | Ankrd9 | ankyrin repeat domain 9 |
| chr12 | 111038686 | 111040401 | ENSMUSG00000037896 | Rcor1 | REST corepressor 1 |
| chr12 | 111165990 | 111167049 | ENSMUSG00000021277 | Traf3 | TNF receptor-associated factor 3 |
| chr12 | 111376684 | 111378403 | ENSMUSG00000021279 | Cdc42bpb | CDC42 binding protein kinase beta |
| chr12 | 111537398 | 111539273 | ENSMUSG00000021282 | Eif5 | eukaryotic translation initiation factor 5 |
| chr12 | 111574378 | 111575471 | ENSMUSG00000096995 | 2810029C07Rik | RIKEN cDNA 2810029C07 gene |
| chr12 | 111712545 | 111713650 | ENSMUSG00000037787 | Coa8 | cytochrome c oxidase assembly factor 8 |
| chr12 | 111758400 | 111760041 | ENSMUSG00000021288 | Klc1 | kinesin light chain 1 |
| chr12 | 111813579 | 111814706 | ENSMUSG00000021286 | Zfyve21 | zinc finger, FYVE domain containing 21 |
| chr12 | 111907470 | 111908417 | ENSMUSG00000021285 | Ppp1r13b | protein phosphatase 1, regulatory subunit 13B |
| chr12 | 112130143 | 112130971 | ENSMUSG00000065574 | Mir203 | microRNA 203 |
| chr12 | 112588417 | 112589437 | ENSMUSG00000037679 | Inf2 | inverted formin, FH2 and WH2 domain containing |
| chr12 | 112644341 | 112645398 | ENSMUSG00000064326 | Siva1 | SIVA1, apoptosis-inducing factor |
| chr12 | 112673599 | 112674622 | ENSMUSG00000001729 | Akt1 | thymoma viral proto-oncogene 1 |
| chr12 | 112678647 | 112679829 | ENSMUSG00000037638 | Zbtb42 | zinc finger and BTB domain containing 42 |
| chr12 | 112693311 | 112693884 | ENSMUSG00000037638 | Zbtb42 | zinc finger and BTB domain containing 42 |
| chr12 | 112828794 | 112829762 | ENSMUSG00000047832 | Cdca4 | cell division cycle associated 4 |
| chr12 | 112976439 | 112977228 | ENSMUSG00000002803 | Btbd6 | BTB (POZ) domain containing 6 |
| chr12 | 112999914 | 113000985 | ENSMUSG00000011158 | Brf1 | BRF1, RNA polymerase III transcription initiation factor 90 kDa subunit |
| chr12 | 113013950 | 113015243 | ENSMUSG00000021143 | Pacs2 | phosphofurin acidic cluster sorting protein 2 |
| chr12 | 113139979 | 113140982 | ENSMUSG00000006356 | Crip2 | cysteine rich protein 2 |
| chr12 | 116280549 | 116282298 | ENSMUSG00000021171 | Esyt2 | extended synaptotagmin-like protein 2 |
| chr12 | 117843327 | 117844299 | ENSMUSG00000021175 | Cdca7l | cell division cycle associated 7 like |
| chr12 | 118301084 | 118302595 | ENSMUSG00000025323 | Sp4 | trans-acting transcription factor 4 |
| chr12 | 118836553 | 118837206 | ENSMUSG00000048562 | Sp8 | trans-acting transcription factor 8 |
| chr13 | 3537553 | 3538669 | ENSMUSG00000021218 | Gdi2 | guanosine diphosphate (GDP) dissociation inhibitor 2 |
| chr13 | 9093448 | 9094886 | ENSMUSG00000033499 | Larp4b | La ribonucleoprotein domain family, member 4B |
| chr13 | 9276390 | 9277443 | ENSMUSG00000048264 | Dip2c | disco interacting protein 2 homolog C |
| chr13 | 9763786 | 9765220 | ENSMUSG00000021156 | Zmynd11 | zinc finger, MYND domain containing 11 |
| chr13 | 13954113 | 13955232 | ENSMUSG00000039242 | B3galnt2 | UDP-GalNAc:betaGlcNAc beta 1,3-galactosaminyltransferase, polypeptide 2 |
| chr13 | 15462998 | 15464547 | ENSMUSG00000021318 | Gli3 | GLI-Kruppel family member GLI3 |
| chr13 | 17943655 | 17944456 | ENSMUSG00000008859 | Rala | v-ral simian leukemia viral oncogene A (ras related) |
| chr13 | 19395423 | 19395958 | ENSMUSG00000003062 | Stard3nl | STARD3 N-terminal like |
| chr13 | 21179985 | 21181184 | ENSMUSG00000021326 | Trim27 | tripartite motif-containing 27 |
| chr13 | 21735016 | 21735491 | ENSMUSG00000067455 | H4c11 | H4 clustered histone 11 |
| chr13 | 22040978 | 22041345 | ENSMUSG00000060639 | H4c9 | H4 clustered histone 9 |
| chr13 | 23531076 | 23531811 | ENSMUSG00000060981 | H4c8 | H4 clustered histone 8 |
| chr13 | 23533866 | 23534371 | ENSMUSG00000061991 | H2ac10 | H2A clustered histone 10 |
| chr13 | 23535269 | 23535613 | ENSMUSG00000099517 | H3c8 | H3 clustered histone 8 |
| chr13 | 23622212 | 23622658 | ENSMUSG00000051627 | H1f4 | H1.4 linker histone, cluster member |
| chr13 | 23751077 | 23751635 | ENSMUSG00000061615 | H2ac4 | H2A clustered histone 4 |
| chr13 | 23756948 | 23757443 | ENSMUSG00000069266 | H4c2 | H4 clustered histone 2 |
| chr13 | 24280709 | 24281494 | ENSMUSG00000021338 | Carmil1 | capping protein regulator and myosin 1 linker 1 |
| chr13 | 24831414 | 24832295 | ENSMUSG00000035958 | Tdp2 | tyrosyl-DNA phosphodiesterase 2 |
| chr13 | 25019856 | 25020476 | ENSMUSG00000021339 | Mrs2 | MRS2 magnesium transporter |
| chr13 | 30136218 | 30136810 | ENSMUSG00000038732 | Mboat1 | membrane bound O-acyltransferase domain containing 1 |
| chr13 | 32338105 | 32338932 | ENSMUSG00000038372 | Gmds | GDP-mannose 4, 6-dehydratase |
| chr13 | 32801814 | 32803072 | ENSMUSG00000021400 | Wrnip1 | Werner helicase interacting protein 1 |
| chr13 | 34344094 | 34345539 | ENSMUSG00000038267 | Slc22a23 | solute carrier family 22, member 23 |
| chr13 | 34651731 | 34653214 | ENSMUSG00000086786 | Gm15908 | predicted gene 15908 |
| chr13 | 35740634 | 35742349 | ENSMUSG00000059288 | Cdyl | chromodomain protein, Y chromosome-like |
| chr13 | 36116971 | 36118118 | ENSMUSG00000046573 | Lyrm4 | LYR motif containing 4 |
| chr13 | 37993540 | 37994325 | ENSMUSG00000021427 | Ssr1 | signal sequence receptor, alpha |
| chr13 | 38151175 | 38152734 | ENSMUSG00000054889 | Dsp | desmoplakin |
| chr13 | 40730476 | 40731890 | ENSMUSG00000021359 | Tfap2a | transcription factor AP-2, alpha |
| chr13 | 41605422 | 41606457 | ENSMUSG00000087370 | Tmem170b | transmembrane protein 170B |
| chr13 | 42051683 | 42052803 | ENSMUSG00000021366 | Hivep1 | human immunodeficiency virus type I enhancer binding protein 1 |
| chr13 | 43303231 | 43304379 | ENSMUSG00000051335 | Gfod1 | glucose-fructose oxidoreductase domain containing 1 |
| chr13 | 43304379 | 43305086 | ENSMUSG00000051335 | Gfod1 | glucose-fructose oxidoreductase domain containing 1 |
| chr13 | 43480359 | 43481680 | ENSMUSG00000038546 | Ranbp9 | RAN binding protein 9 |
| chr13 | 43559568 | 43560507 | ENSMUSG00000021371 | Mcur1 | mitochondrial calcium uniporter regulator 1 |
| chr13 | 44733368 | 44734190 | ENSMUSG00000038518 | Jarid2 | jumonji, AT rich interactive domain 2 |
| chr13 | 45001538 | 45002285 | ENSMUSG00000057531 | Dtnbp1 | dystrobrevin binding protein 1 |
| chr13 | 46669454 | 46670271 | ENSMUSG00000069237 | Fam8a1 | family with sequence similarity 8, member A1 |
| chr13 | 46726901 | 46728044 | ENSMUSG00000021374 | Nup153 | nucleoporin 153 |
| chr13 | 46929220 | 46930496 | ENSMUSG00000021375 | Kif13a | kinesin family member 13A |
| chr13 | 47105345 | 47106668 | ENSMUSG00000021377 | Dek | DEK proto-oncogene (DNA binding) |
| chr13 | 48625182 | 48625885 | ENSMUSG00000038042 | Ptpdc1 | protein tyrosine phosphatase domain containing 1 |
| chr13 | 48966848 | 48968716 | ENSMUSG00000038014 | Fam120a | family with sequence similarity 120, member A |
| chr13 | 49187219 | 49188064 | ENSMUSG00000037966 | Ninj1 | ninjurin 1 |
| chr13 | 51100243 | 51101552 | ENSMUSG00000021395 | Spin1 | spindlin 1 |
| chr13 | 51792676 | 51793734 | ENSMUSG00000021451 | Sema4d | sema domain, immunoglobulin domain (Ig), transmembrane domain (TM) and short cytoplasmic domain, (semaphorin) 4D |
| chr13 | 52929240 | 52929834 | ENSMUSG00000021460 | Auh | AU RNA binding protein/enoyl-coenzyme A hydratase |
| chr13 | 54503344 | 54504455 | ENSMUSG00000043183 | Simc1 | SUMO-interacting motifs containing 1 |
| chr13 | 55099718 | 55100389 | ENSMUSG00000025878 | Uimc1 | ubiquitin interaction motif containing 1 |
| chr13 | 55471579 | 55472881 | ENSMUSG00000098341 | Mir6944 | microRNA 6944 |
| chr13 | 55692819 | 55693693 | ENSMUSG00000045767 | B230219D22Rik | RIKEN cDNA B230219D22 gene |
| chr13 | 55727063 | 55728186 | ENSMUSG00000021496 | Pcbd2 | pterin 4 alpha carbinolamine dehydratase/dimerization cofactor of hepatocyte nuclear factor 1 alpha (TCF1) 2 |
| chr13 | 56134695 | 56135750 | ENSMUSG00000015937 | Macroh2a1 | macroH2A.1 histone |
| chr13 | 56438252 | 56438835 | ENSMUSG00000021509 | Slc25a48 | solute carrier family 25, member 48 |
| chr13 | 56438835 | 56439082 | ENSMUSG00000021509 | Slc25a48 | solute carrier family 25, member 48 |
| chr13 | 56702852 | 56703847 | ENSMUSG00000021540 | Smad5 | SMAD family member 5 |
| chr13 | 58214966 | 58215882 | ENSMUSG00000005312 | Ubqln1 | ubiquilin 1 |
| chr13 | 58273578 | 58274671 | ENSMUSG00000021552 | Gkap1 | G kinase anchoring protein 1 |
| chr13 | 58401253 | 58403090 | ENSMUSG00000021546 | Hnrnpk | heterogeneous nuclear ribonucleoprotein K |
| chr13 | 59556638 | 59557713 | ENSMUSG00000021557 | Agtpbp1 | ATP/GTP binding protein 1 |
| chr13 | 60175741 | 60177295 | ENSMUSG00000052957 | Gas1 | growth arrest specific 1 |
| chr13 | 60177295 | 60177837 | ENSMUSG00000052957 | Gas1 | growth arrest specific 1 |
| chr13 | 63239792 | 63240207 | ENSMUSG00000021458 | Aopep | aminopeptidase O |
| chr13 | 63567788 | 63568290 | ENSMUSG00000021466 | Ptch1 | patched 1 |
| chr13 | 63815098 | 63815700 | ENSMUSG00000021470 | Ercc6l2 | excision repair cross-complementing rodent repair deficiency, complementation group 6 like 2 |
| chr13 | 64152250 | 64153570 | ENSMUSG00000044934 | Zfp367 | zinc finger protein 367 |
| chr13 | 64161588 | 64162495 | ENSMUSG00000021476 | Habp4 | hyaluronic acid binding protein 4 |
| chr13 | 69533314 | 69534321 | ENSMUSG00000034575 | Tent4a | terminal nucleotidyltransferase 4A |
| chr13 | 69534321 | 69535426 | ENSMUSG00000034575 | Tent4a | terminal nucleotidyltransferase 4A |
| chr13 | 69610682 | 69611870 | ENSMUSG00000021594 | Srd5a1 | steroid 5 alpha-reductase 1 |
| chr13 | 70637396 | 70637907 | ENSMUSG00000034525 | Ice1 | interactor of little elongation complex ELL subunit 1 |
| chr13 | 73603726 | 73604522 | ENSMUSG00000021610 | Clptm1l | CLPTM1-like |
| chr13 | 75943020 | 75944092 | ENSMUSG00000021589 | Rhobtb3 | Rho-related BTB domain containing 3 |
| chr13 | 78198181 | 78199726 | ENSMUSG00000069171 | Nr2f1 | nuclear receptor subfamily 2, group F, member 1 |
| chr13 | 80883123 | 80883468 | ENSMUSG00000074794 | Arrdc3 | arrestin domain containing 3 |
| chr13 | 80885544 | 80886275 | ENSMUSG00000074794 | Arrdc3 | arrestin domain containing 3 |
| chr13 | 81710870 | 81711763 | ENSMUSG00000035834 | Polr3g | polymerase (RNA) III (DNA directed) polypeptide G |
| chr13 | 83730503 | 83731393 | ENSMUSG00000050334 | C130071C03Rik | RIKEN cDNA C130071C03 gene |
| chr13 | 85288504 | 85289693 | ENSMUSG00000021549 | Rasa1 | RAS p21 protein activator 1 |
| chr13 | 91223748 | 91224264 | ENSMUSG00000021619 | Atg10 | autophagy related 10 |
| chr13 | 91460458 | 91462268 | ENSMUSG00000003992 | Ssbp2 | single-stranded DNA binding protein 2 |
| chr13 | 93498376 | 93499948 | ENSMUSG00000021690 | Jmy | junction-mediating and regulatory protein |
| chr13 | 94284840 | 94285706 | ENSMUSG00000021687 | Scamp1 | secretory carrier membrane protein 1 |
| chr13 | 97240984 | 97241919 | ENSMUSG00000041773 | Enc1 | ectodermal-neural cortex 1 |
| chr13 | 97241919 | 97242266 | ENSMUSG00000041773 | Enc1 | ectodermal-neural cortex 1 |
| chr13 | 98814480 | 98815708 | ENSMUSG00000041685 | Fcho2 | FCH domain only 2 |
| chr13 | 100107850 | 100108518 | ENSMUSG00000021643 | Serf1 | small EDRK-rich factor 1 |
| chr13 | 103920203 | 103921751 | ENSMUSG00000021709 | Erbin | Erbb2 interacting protein |
| chr13 | 107021597 | 107022730 | ENSMUSG00000071181 | 3830408C21Rik | RIKEN cDNA 3830408C21 gene |
| chr13 | 107889904 | 107891519 | ENSMUSG00000032846 | Zswim6 | zinc finger SWIM-type containing 6 |
| chr13 | 111807939 | 111809831 | ENSMUSG00000021754 | Map3k1 | mitogen-activated protein kinase kinase kinase 1 |
| chr13 | 114458423 | 114459440 | ENSMUSG00000021765 | Fst | follistatin |
| chr13 | 117025494 | 117025992 | ENSMUSG00000021725 | Parp8 | poly (ADP-ribose) polymerase family, member 8 |
| chr14 | 8097983 | 8098740 | ENSMUSG00000033885 | Pxk | PX domain containing serine/threonine kinase |
| chr14 | 8213902 | 8214757 | ENSMUSG00000021752 | Kctd6 | potassium channel tetramerisation domain containing 6 |
| chr14 | 14012230 | 14013391 | ENSMUSG00000021738 | Atxn7 | ataxin 7 |
| chr14 | 14703008 | 14703745 | ENSMUSG00000021733 | Slc4a7 | solute carrier family 4, sodium bicarbonate cotransporter, member 7 |
| chr14 | 16364423 | 16365287 | ENSMUSG00000017485 | Top2b | topoisomerase (DNA) II beta |
| chr14 | 18238194 | 18239464 | ENSMUSG00000021775 | Nr1d2 | nuclear receptor subfamily 1, group D, member 2 |
| chr14 | 20545623 | 20546661 | ENSMUSG00000021816 | Ppp3cb | protein phosphatase 3, catalytic subunit, beta isoform |
| chr14 | 20694881 | 20695900 | ENSMUSG00000039357 | Fut11 | fucosyltransferase 11 |
| chr14 | 20702621 | 20703371 | ENSMUSG00000063787 | Chchd1 | coiled-coil-helix-coiled-coil-helix domain containing 1 |
| chr14 | 20733921 | 20734589 | ENSMUSG00000039308 | Ndst2 | N-deacetylase/N-sulfotransferase (heparan glucosaminyl) 2 |
| chr14 | 21075882 | 21076414 | ENSMUSG00000039197 | Adk | adenosine kinase |
| chr14 | 21831018 | 21831841 | ENSMUSG00000021771 | Vdac2 | voltage-dependent anion channel 2 |
| chr14 | 24244674 | 24245925 | ENSMUSG00000021782 | Dlg5 | discs large MAGUK scaffold protein 5 |
| chr14 | 26533960 | 26534970 | ENSMUSG00000021870 | Slmap | sarcolemma associated protein |
| chr14 | 26637801 | 26639087 | ENSMUSG00000021877 | Arf4 | ADP-ribosylation factor 4 |
| chr14 | 26969752 | 26970996 | ENSMUSG00000040760 | Appl1 | adaptor protein, phosphotyrosine interaction, PH domain and leucine zipper containing 1 |
| chr14 | 27428772 | 27429697 | ENSMUSG00000040651 | Tasor | transcription activation suppressor |
| chr14 | 30008840 | 30009376 | ENSMUSG00000015970 | Chdh | choline dehydrogenase |
| chr14 | 30549040 | 30549709 | ENSMUSG00000021957 | Tkt | transketolase |
| chr14 | 30715377 | 30716747 | ENSMUSG00000006527 | Sfmbt1 | Scm-like with four mbt domains 1 |
| chr14 | 31018653 | 31019656 | ENSMUSG00000042354 | Gnl3 | guanine nucleotide binding protein-like 3 (nucleolar) |
| chr14 | 31250897 | 31251729 | ENSMUSG00000021901 | Bap1 | Brca1 associated protein 1 |
| chr14 | 31829847 | 31831223 | ENSMUSG00000014496 | Ankrd28 | ankyrin repeat domain 28 |
| chr14 | 33320160 | 33320809 | ENSMUSG00000063506 | Arhgap22 | Rho GTPase activating protein 22 |
| chr14 | 33351032 | 33351565 | ENSMUSG00000063506 | Arhgap22 | Rho GTPase activating protein 22 |
| chr14 | 34310151 | 34311632 | ENSMUSG00000021794 | Glud1 | glutamate dehydrogenase 1 |
| chr14 | 34501878 | 34503397 | ENSMUSG00000021796 | Bmpr1a | bone morphogenetic protein receptor, type 1A |
| chr14 | 34548753 | 34549717 | ENSMUSG00000021797 | 9230112D13Rik | RIKEN cDNA 9230112D13 gene |
| chr14 | 34673254 | 34674839 | ENSMUSG00000041408 | Wapl | WAPL cohesin release factor |
| chr14 | 36968128 | 36968925 | ENSMUSG00000058690 | Ccser2 | coiled-coil serine rich 2 |
| chr14 | 45219231 | 45220780 | ENSMUSG00000021830 | Txndc16 | thioredoxin domain containing 16 |
| chr14 | 45317985 | 45319007 | ENSMUSG00000021831 | Ero1a | endoplasmic reticulum oxidoreductase 1 alpha |
| chr14 | 45351011 | 45351709 | ENSMUSG00000053205 | Styx | serine/threonine/tyrosine interaction protein |
| chr14 | 45388561 | 45389174 | ENSMUSG00000037722 | Gnpnat1 | glucosamine-phosphate N-acetyltransferase 1 |
| chr14 | 45529578 | 45530346 | ENSMUSG00000037712 | Fermt2 | fermitin family member 2 |
| chr14 | 45656935 | 45658575 | ENSMUSG00000093167 | Mir5131 | microRNA 5131 |
| chr14 | 46787787 | 46788449 | ENSMUSG00000015759 | Cnih1 | cornichon family AMPA receptor auxiliary protein 1 |
| chr14 | 46821750 | 46822421 | ENSMUSG00000062014 | Gmfb | glia maturation factor, beta |
| chr14 | 47276331 | 47277454 | ENSMUSG00000037572 | Wdhd1 | WD repeat and HMG-box DNA binding protein 1 |
| chr14 | 47472368 | 47473277 | ENSMUSG00000037536 | Fbxo34 | F-box protein 34 |
| chr14 | 47649164 | 47650198 | ENSMUSG00000021843 | Ktn1 | kinectin 1 |
| chr14 | 48446128 | 48446686 | ENSMUSG00000036339 | Tmem260 | transmembrane protein 260 |
| chr14 | 49065810 | 49066762 | ENSMUSG00000061244 | Exoc5 | exocyst complex component 5 |
| chr14 | 50930481 | 50931550 | ENSMUSG00000115338 | Pnp | purine-nucleoside phosphorylase |
| chr14 | 52019552 | 52020319 | ENSMUSG00000049295 | Zfp219 | zinc finger protein 219 |
| chr14 | 52196650 | 52197645 | ENSMUSG00000035726 | Supt16 | SPT16, facilitates chromatin remodeling subunit |
| chr14 | 52257300 | 52258052 | ENSMUSG00000053754 | Chd8 | chromodomain helicase DNA binding protein 8 |
| chr14 | 54876246 | 54877736 | ENSMUSG00000072494 | Ppp1r3e | protein phosphatase 1, regulatory subunit 3E |
| chr14 | 55642993 | 55643902 | ENSMUSG00000002320 | Tm9sf1 | transmembrane 9 superfamily member 1 |
| chr14 | 56811948 | 56812942 | ENSMUSG00000100017 | 2410022M11Rik | RIKEN cDNA 2410022M11 gene |
| chr14 | 57423949 | 57424423 | ENSMUSG00000040040 | Ift88 | intraflagellar transport 88 |
| chr14 | 57524856 | 57525704 | ENSMUSG00000050222 | Il17d | interleukin 17D |
| chr14 | 57664281 | 57665406 | ENSMUSG00000021952 | Xpo4 | exportin 4 |
| chr14 | 57745210 | 57746588 | ENSMUSG00000021959 | Lats2 | large tumor suppressor 2 |
| chr14 | 57797958 | 57799067 | ENSMUSG00000061104 | Sap18b | Sin3-associated polypeptide 18B |
| chr14 | 57825913 | 57826771 | ENSMUSG00000021965 | Ska3 | spindle and kinetochore associated complex subunit 3 |
| chr14 | 58071829 | 58072596 | ENSMUSG00000021974 | Fgf9 | fibroblast growth factor 9 |
| chr14 | 59200917 | 59201726 | ENSMUSG00000035469 | Rcbtb1 | regulator of chromosome condensation (RCC1) and BTB (POZ) domain containing protein 1 |
| chr14 | 60784457 | 60784874 | ENSMUSG00000021993 | Mipep | mitochondrial intermediate peptidase |
| chr14 | 61439003 | 61440368 | ENSMUSG00000021929 | Kpna3 | karyopherin (importin) alpha 3 |
| chr14 | 61680886 | 61682225 | ENSMUSG00000097589 | Dleu2 | deleted in lymphocytic leukemia, 2 |
| chr14 | 62760008 | 62761777 | ENSMUSG00000035161 | Ints6 | integrator complex subunit 6 |
| chr14 | 65097546 | 65098362 | ENSMUSG00000021978 | Extl3 | exostosin-like glycosyltransferase 3 |
| chr14 | 65358036 | 65359257 | ENSMUSG00000034522 | Zfp395 | zinc finger protein 395 |
| chr14 | 66868645 | 66869323 | ENSMUSG00000022048 | Dpysl2 | dihydropyrimidinase-like 2 |
| chr14 | 67008129 | 67009189 | ENSMUSG00000022051 | Bnip3l | BCL2/adenovirus E1B interacting protein 3-like |
| chr14 | 67715176 | 67716027 | ENSMUSG00000034327 | Kctd9 | potassium channel tetramerisation domain containing 9 |
| chr14 | 67932644 | 67933947 | ENSMUSG00000044447 | Dock5 | dedicator of cytokinesis 5 |
| chr14 | 70288845 | 70289690 | ENSMUSG00000022092 | Ppp3cc | protein phosphatase 3, catalytic subunit, gamma isoform |
| chr14 | 70350805 | 70351643 | ENSMUSG00000022094 | Slc39a14 | solute carrier family 39 (zinc transporter), member 14 |
| chr14 | 70519791 | 70520563 | ENSMUSG00000022098 | Bmp1 | bone morphogenetic protein 1 |
| chr14 | 70577552 | 70578308 | ENSMUSG00000045211 | Nudt18 | nudix (nucleoside diphosphate linked moiety X)-type motif 18 |
| chr14 | 70617966 | 70618598 | ENSMUSG00000022099 | Dmtn | dematin actin binding protein |
| chr14 | 70765824 | 70767075 | ENSMUSG00000022100 | Xpo7 | exportin 7 |
| chr14 | 72709147 | 72710464 | ENSMUSG00000033487 | Fndc3a | fibronectin type III domain containing 3A |
| chr14 | 73142952 | 73143739 | ENSMUSG00000022106 | Rcbtb2 | regulator of chromosome condensation (RCC1) and BTB (POZ) domain containing protein 2 |
| chr14 | 73325101 | 73326316 | ENSMUSG00000022105 | Rb1 | RB transcriptional corepressor 1 |
| chr14 | 75844707 | 75846029 |  | Gm4285 | predicted gene 4285 |
| chr14 | 76010516 | 76010949 | ENSMUSG00000067995 | Gtf2f2 | general transcription factor IIF, polypeptide 2 |
| chr14 | 76414116 | 76415706 | ENSMUSG00000022010 | Tsc22d1 | TSC22 domain family, member 1 |
| chr14 | 76415706 | 76416028 | ENSMUSG00000022010 | Tsc22d1 | TSC22 domain family, member 1 |
| chr14 | 76504245 | 76504624 | ENSMUSG00000022010 | Tsc22d1 | TSC22 domain family, member 1 |
| chr14 | 78536398 | 78537224 | ENSMUSG00000022016 | Akap11 | A kinase (PRKA) anchor protein 11 |
| chr14 | 78848866 | 78849491 | ENSMUSG00000058997 | Vwa8 | von Willebrand factor A domain containing 8 |
| chr14 | 79390282 | 79390858 | ENSMUSG00000022020 | Naa16 | N(alpha)-acetyltransferase 16, NatA auxiliary subunit |
| chr14 | 79426428 | 79426827 | ENSMUSG00000043881 | Kbtbd7 | kelch repeat and BTB (POZ) domain containing 7 |
| chr14 | 87415778 | 87417512 | ENSMUSG00000022019 | Tdrd3 | tudor domain containing 3 |
| chr14 | 99099254 | 99099875 | ENSMUSG00000022064 | Pibf1 | progesterone immunomodulatory binding factor 1 |
| chr14 | 99298321 | 99299841 | ENSMUSG00000005148 | Klf5 | Kruppel-like factor 5 |
| chr14 | 101607701 | 101609316 | ENSMUSG00000033083 | Tbc1d4 | TBC1 domain family, member 4 |
| chr14 | 103070078 | 103070568 | ENSMUSG00000022125 | Cln5 | ceroid-lipofuscinosis, neuronal 5 |
| chr14 | 103098698 | 103099860 | ENSMUSG00000022124 | Fbxl3 | F-box and leucine-rich repeat protein 3 |
| chr14 | 103345938 | 103347183 | ENSMUSG00000033004 | Mycbp2 | MYC binding protein 2, E3 ubiquitin protein ligase |
| chr14 | 103650054 | 103651135 | ENSMUSG00000055717 | Slain1 | SLAIN motif family, member 1 |
| chr14 | 104467353 | 104468369 | ENSMUSG00000048349 | Pou4f1 | POU domain, class 4, transcription factor 1 |
| chr14 | 104522087 | 104522847 | ENSMUSG00000022120 | Obi1 | ORC ubiquitin ligase 1 |
| chr14 | 105896847 | 105897721 | ENSMUSG00000022114 | Spry2 | sprouty RTK signaling antagonist 2 |
| chr14 | 118136830 | 118137669 | ENSMUSG00000022131 | Gpr180 | G protein-coupled receptor 180 |
| chr14 | 118705596 | 118706756 | ENSMUSG00000032849 | Abcc4 | ATP-binding cassette, sub-family C (CFTR/MRP), member 4 |
| chr14 | 118937482 | 118938609 | ENSMUSG00000022136 | Dnajc3 | DnaJ heat shock protein family (Hsp40) member C3 |
| chr14 | 120478038 | 120479188 | ENSMUSG00000051615 | Rap2a | RAS related protein 2a |
| chr14 | 120910533 | 120911834 | ENSMUSG00000030662 | Ipo5 | importin 5 |
| chr14 | 121035092 | 121036364 | ENSMUSG00000025555 | Farp1 | FERM, RhoGEF (Arhgef) and pleckstrin domain protein 1 (chondrocyte-derived) |
| chr14 | 121378216 | 121379890 | ENSMUSG00000063410 | Stk24 | serine/threonine kinase 24 |
| chr14 | 121797248 | 121798575 | ENSMUSG00000025558 | Dock9 | dedicator of cytokinesis 9 |
| chr14 | 122106838 | 122107708 |  | A330035P11Rik | RIKEN cDNA A330035P11 gene |
| chr14 | 122982595 | 122983643 | ENSMUSG00000041594 | Tmtc4 | transmembrane and tetratricopeptide repeat containing 4 |
| chr15 | 3979027 | 3979611 | ENSMUSG00000022184 | Fbxo4 | F-box protein 4 |
| chr15 | 4026380 | 4027518 |  | BC037032 | cDNA Sequence BC037032 |
| chr15 | 6707485 | 6708936 | ENSMUSG00000050310 | Rictor | RPTOR independent companion of MTOR, complex 2 |
| chr15 | 6708936 | 6709461 | ENSMUSG00000050310 | Rictor | RPTOR independent companion of MTOR, complex 2 |
| chr15 | 9070877 | 9072299 | ENSMUSG00000022253 | Nadk2 | NAD kinase 2, mitochondrial |
| chr15 | 10469757 | 10470657 | ENSMUSG00000044224 | Dnajc21 | DnaJ heat shock protein family (Hsp40) member C21 |
| chr15 | 12204659 | 12205797 | ENSMUSG00000039458 | Mtmr12 | myotubularin related protein 12 |
| chr15 | 12824263 | 12825254 | ENSMUSG00000022191 | Drosha | drosha, ribonuclease type III |
| chr15 | 25622534 | 25623453 | ENSMUSG00000022272 | Myo10 | myosin X |
| chr15 | 25842995 | 25843839 | ENSMUSG00000022270 | Retreg1 | reticulophagy regulator 1 |
| chr15 | 25984035 | 25985189 | ENSMUSG00000052253 | Zfp622 | zinc finger protein 622 |
| chr15 | 27466202 | 27467132 | ENSMUSG00000022265 | Ank | progressive ankylosis |
| chr15 | 31530401 | 31531428 | ENSMUSG00000039100 | Marchf6 | membrane associated ring-CH-type finger 6 |
| chr15 | 34082239 | 34083339 | ENSMUSG00000022255 | Mtdh | metadherin |
| chr15 | 35371088 | 35371907 | ENSMUSG00000037646 | Vps13b | vacuolar protein sorting 13B |
| chr15 | 36282401 | 36283519 | ENSMUSG00000022280 | Rnf19a | ring finger protein 19A |
| chr15 | 36608087 | 36609789 | ENSMUSG00000022283 | Pabpc1 | poly(A) binding protein, cytoplasmic 1 |
| chr15 | 36938533 | 36939068 | ENSMUSG00000062397 | Zfp706 | zinc finger protein 706 |
| chr15 | 37006768 | 37007938 | ENSMUSG00000062397 | Zfp706 | zinc finger protein 706 |
| chr15 | 38078047 | 38079196 | ENSMUSG00000037487 | Ubr5 | ubiquitin protein ligase E3 component n-recognin 5 |
| chr15 | 38470493 | 38471350 | ENSMUSG00000099110 | Mir6951 | microRNA 6951 |
| chr15 | 38473489 | 38473980 | ENSMUSG00000099110 | Mir6951 | microRNA 6951 |
| chr15 | 38518131 | 38519857 | ENSMUSG00000037458 | Azin1 | antizyme inhibitor 1 |
| chr15 | 38661759 | 38662262 | ENSMUSG00000022295 | Atp6v1c1 | ATPase, H+ transporting, lysosomal V1 subunit C1 |
| chr15 | 39006011 | 39006717 | ENSMUSG00000022297 | Fzd6 | frizzled class receptor 6 |
| chr15 | 39111971 | 39113264 | ENSMUSG00000022300 | Dcaf13 | DDB1 and CUL4 associated factor 13 |
| chr15 | 39943117 | 39944434 | ENSMUSG00000022305 | Lrp12 | low density lipoprotein-related protein 12 |
| chr15 | 41788720 | 41789670 | ENSMUSG00000022307 | Oxr1 | oxidation resistance 1 |
| chr15 | 44619028 | 44619936 | ENSMUSG00000022339 | Ebag9 | estrogen receptor-binding fragment-associated gene 9 |
| chr15 | 53345611 | 53346657 | ENSMUSG00000061731 | Ext1 | exostosin glycosyltransferase 1 |
| chr15 | 55112280 | 55112934 | ENSMUSG00000022419 | Deptor | DEP domain containing MTOR-interacting protein |
| chr15 | 57891985 | 57892648 | ENSMUSG00000022365 | Derl1 | Der1-like domain family, member 1 |
| chr15 | 58415330 | 58416108 | ENSMUSG00000037119 | Fam91a1 | family with sequence similarity 91, member A1 |
| chr15 | 58822682 | 58823943 | ENSMUSG00000062373 | Tmem65 | transmembrane protein 65 |
| chr15 | 58888655 | 58889723 | ENSMUSG00000022354 | Ndufb9 | NADH:ubiquinone oxidoreductase subunit B9 |
| chr15 | 59081058 | 59082545 | ENSMUSG00000022353 | Mtss1 | MTSS I-BAR domain containing 1 |
| chr15 | 59314671 | 59315672 | ENSMUSG00000022351 | Sqle | squalene epoxidase |
| chr15 | 63997478 | 63998641 | ENSMUSG00000022378 | Cyrib | CYFIP related Rac1 interactor B |
| chr15 | 64060020 | 64060717 | ENSMUSG00000022378 | Cyrib | CYFIP related Rac1 interactor B |
| chr15 | 64312249 | 64313128 | ENSMUSG00000022377 | Asap1 | ArfGAP with SH3 domain, ankyrin repeat and PH domain1 |
| chr15 | 64382140 | 64383165 | ENSMUSG00000022377 | Asap1 | ArfGAP with SH3 domain, ankyrin repeat and PH domain1 |
| chr15 | 66577394 | 66578148 | ENSMUSG00000072501 | Phf20l1 | PHD finger protein 20-like 1 |
| chr15 | 66968509 | 66969527 | ENSMUSG00000005125 | Ndrg1 | N-myc downstream regulated gene 1 |
| chr15 | 68362142 | 68363465 | ENSMUSG00000065437 | Mir30d | microRNA 30d |
| chr15 | 73090210 | 73090886 | ENSMUSG00000068391 | Chrac1 | chromatin accessibility complex 1 |
| chr15 | 73183889 | 73185471 | ENSMUSG00000036698 | Ago2 | argonaute RISC catalytic subunit 2 |
| chr15 | 73422514 | 73423833 | ENSMUSG00000022607 | Ptk2 | PTK2 protein tyrosine kinase 2 |
| chr15 | 74672352 | 74672871 | ENSMUSG00000022602 | Arc | activity regulated cytoskeletal-associated protein |
| chr15 | 74720942 | 74721847 | ENSMUSG00000056665 | Them6 | thioesterase superfamily member 6 |
| chr15 | 75940853 | 75941733 | ENSMUSG00000050846 | Zfp623 | zinc finger protein 623 |
| chr15 | 76069098 | 76069999 | ENSMUSG00000022568 | Scrib | scribbled planar cell polarity |
| chr15 | 76080260 | 76080926 | ENSMUSG00000002524 | Puf60 | poly-U binding splicing factor 60 |
| chr15 | 76091275 | 76091744 | ENSMUSG00000075590 | Nrbp2 | nuclear receptor binding protein 2 |
| chr15 | 76228604 | 76229839 | ENSMUSG00000022565 | Plec | plectin |
| chr15 | 76330947 | 76332156 | ENSMUSG00000022561 | Gpaa1 | GPI anchor attachment protein 1 |
| chr15 | 76368893 | 76369752 | ENSMUSG00000022554 | Hgh1 | HGH1 homolog |
| chr15 | 76457348 | 76458346 | ENSMUSG00000034161 | Scx | scleraxis |
| chr15 | 76476940 | 76478245 | ENSMUSG00000022557 | Bop1 | block of proliferation 1 |
| chr15 | 76537807 | 76539120 | ENSMUSG00000022559 | Fbxl6 | F-box and leucine-rich repeat protein 6 |
| chr15 | 76817080 | 76818399 | ENSMUSG00000033697 | Arhgap39 | Rho GTPase activating protein 39 |
| chr15 | 77305850 | 77307275 | ENSMUSG00000033565 | Rbfox2 | RNA binding protein, fox-1 homolog (C. elegans) 2 |
| chr15 | 78802229 | 78803842 | ENSMUSG00000033170 | Card10 | caspase recruitment domain family, member 10 |
| chr15 | 79028059 | 79029102 | ENSMUSG00000096210 | H1f0 | H1.0 linker histone |
| chr15 | 79108525 | 79109649 | ENSMUSG00000033039 | Micall1 | microtubule associated monooxygenase, calponin and LIM domain containing -like 1 |
| chr15 | 79686581 | 79687363 | ENSMUSG00000022426 | Josd1 | Josephin domain containing 1 |
| chr15 | 79687363 | 79688349 | ENSMUSG00000022426 | Josd1 | Josephin domain containing 1 |
| chr15 | 79741918 | 79742888 | ENSMUSG00000087543 | Gm16576 | predicted gene 16576 |
| chr15 | 79803962 | 79804866 | ENSMUSG00000022421 | Nptxr | neuronal pentraxin receptor |
| chr15 | 81522946 | 81523576 | ENSMUSG00000022400 | Rbx1 | ring-box 1 |
| chr15 | 81585291 | 81586818 | ENSMUSG00000055024 | Ep300 | E1A binding protein p300 |
| chr15 | 81693883 | 81694647 | ENSMUSG00000063765 | Chadl | chondroadherin-like |
| chr15 | 81810273 | 81811842 | ENSMUSG00000022389 | Tef | thyrotroph embryonic factor |
| chr15 | 82015944 | 82016755 | ENSMUSG00000022471 | Xrcc6 | X-ray repair complementing defective repair in Chinese hamster cells 6 |
| chr15 | 82147004 | 82148147 | ENSMUSG00000022463 | Srebf2 | sterol regulatory element binding factor 2 |
| chr15 | 82898681 | 82900088 | ENSMUSG00000100199 | Gm20324 | predicted gene, 20324 |
| chr15 | 83464091 | 83464768 | ENSMUSG00000016664 | Pacsin2 | protein kinase C and casein kinase substrate in neurons 2 |
| chr15 | 84854949 | 84855728 | ENSMUSG00000016624 | Phf21b | PHD finger protein 21B |
| chr15 | 84922905 | 84924020 | ENSMUSG00000016619 | Nup50 | nucleoporin 50 |
| chr15 | 85653361 | 85653985 |  | Lncppara | long noncoding RNA near Ppara |
| chr15 | 86033043 | 86034427 | ENSMUSG00000016028 | Celsr1 | cadherin, EGF LAG seven-pass G-type receptor 1 |
| chr15 | 86058438 | 86059176 | ENSMUSG00000035900 | Gramd4 | GRAM domain containing 4 |
| chr15 | 86185241 | 86186225 | ENSMUSG00000086424 | Gm15569 | predicted gene 15569 |
| chr15 | 88733026 | 88734453 | ENSMUSG00000022387 | Brd1 | bromodomain containing 1 |
| chr15 | 88751186 | 88752446 | ENSMUSG00000034333 | Zbed4 | zinc finger, BED type containing 4 |
| chr15 | 89075737 | 89076776 | ENSMUSG00000015363 | Trabd | TraB domain containing |
| chr15 | 89088889 | 89089810 | ENSMUSG00000035757 | Selenoo | selenoprotein O |
| chr15 | 89373547 | 89374117 | ENSMUSG00000091780 | Sco2 | SCO2 cytochrome c oxidase assembly protein |
| chr15 | 93274719 | 93275437 | ENSMUSG00000036197 | Gxylt1 | glucoside xylosyltransferase 1 |
| chr15 | 93336060 | 93337397 | ENSMUSG00000022634 | Yaf2 | YY1 associated factor 2 |
| chr15 | 93398100 | 93398723 | ENSMUSG00000036167 | Pphln1 | periphilin 1 |
| chr15 | 96459924 | 96461752 | ENSMUSG00000033228 | Scaf11 | SR-related CTD-associated factor 11 |
| chr15 | 96698472 | 96700325 | ENSMUSG00000022462 | Slc38a2 | solute carrier family 38, member 2 |
| chr15 | 96709366 | 96710375 | ENSMUSG00000022462 | Slc38a2 | solute carrier family 38, member 2 |
| chr15 | 98533468 | 98534402 | ENSMUSG00000022992 | Kansl2 | KAT8 regulatory NSL complex subunit 2 |
| chr15 | 98870982 | 98872175 | ENSMUSG00000048154 | Kmt2d | lysine (K)-specific methyltransferase 2D |
| chr15 | 98933825 | 98934947 | ENSMUSG00000023004 | Tuba1b | tubulin, alpha 1B |
| chr15 | 99126080 | 99126984 | ENSMUSG00000051934 | Spats2 | spermatogenesis associated, serine-rich 2 |
| chr15 | 99725586 | 99726136 | ENSMUSG00000023020 | Cox14 | cytochrome c oxidase assembly protein 14 |
| chr15 | 99771773 | 99772745 | ENSMUSG00000023021 | Cers5 | ceramide synthase 5 |
| chr15 | 99972538 | 99973867 | ENSMUSG00000023025 | Larp4 | La ribonucleoprotein domain family, member 4 |
| chr15 | 100038500 | 100039299 | ENSMUSG00000023026 | Dip2b | disco interacting protein 2 homolog B |
| chr15 | 100227539 | 100228604 | ENSMUSG00000023027 | Atf1 | activating transcription factor 1 |
| chr15 | 100422467 | 100423273 | ENSMUSG00000023030 | Slc11a2 | solute carrier family 11 (proton-coupled divalent metal ion transporters), member 2 |
| chr15 | 100636611 | 100637192 | ENSMUSG00000053559 | Smagp | small cell adhesion glycoprotein |
| chr15 | 101246237 | 101246744 | ENSMUSG00000023034 | Nr4a1 | nuclear receptor subfamily 4, group A, member 1 |
| chr15 | 101253900 | 101254509 | ENSMUSG00000023034 | Nr4a1 | nuclear receptor subfamily 4, group A, member 1 |
| chr15 | 101266574 | 101267160 | ENSMUSG00000023034 | Nr4a1 | nuclear receptor subfamily 4, group A, member 1 |
| chr15 | 102203160 | 102204273 | ENSMUSG00000023044 | Csad | cysteine sulfinic acid decarboxylase |
| chr15 | 102246871 | 102248090 | ENSMUSG00000001288 | Rarg | retinoic acid receptor, gamma |
| chr15 | 102279218 | 102279911 | ENSMUSG00000045665 | Mfsd5 | major facilitator superfamily domain containing 5 |
| chr15 | 102624953 | 102626106 | ENSMUSG00000099083 | Atf7 | activating transcription factor 7 |
| chr15 | 102670632 | 102671158 | ENSMUSG00000062683 | Atp5g2 | ATP synthase, H+ transporting, mitochondrial F0 complex, subunit C2 (subunit 9) |
| chr16 | 3884175 | 3885012 | ENSMUSG00000039789 | Zfp597 | zinc finger protein 597 |
| chr16 | 4077351 | 4078002 | ENSMUSG00000005981 | Trap1 | TNF receptor-associated protein 1 |
| chr16 | 4419857 | 4420829 | ENSMUSG00000005580 | Adcy9 | adenylate cyclase 9 |
| chr16 | 4878897 | 4880471 | ENSMUSG00000039568 | Ubald1 | UBA-like domain containing 1 |
| chr16 | 4963862 | 4964634 | ENSMUSG00000022515 | Anks3 | ankyrin repeat and sterile alpha motif domain containing 3 |
| chr16 | 5013211 | 5013697 | ENSMUSG00000022540 | Rogdi | rogdi homolog |
| chr16 | 5049450 | 5050914 | ENSMUSG00000039473 | Ubn1 | ubinuclein 1 |
| chr16 | 5131565 | 5132758 | ENSMUSG00000039457 | Ppl | periplakin |
| chr16 | 5233496 | 5233943 | ENSMUSG00000039427 | Alg1 | asparagine-linked glycosylation 1 (beta-1,4-mannosyltransferase) |
| chr16 | 5255359 | 5256176 | ENSMUSG00000022544 | Eef2kmt | eukaryotic elongation factor 2 lysine methyltransferase |
| chr16 | 8671460 | 8672575 | ENSMUSG00000008393 | Carhsp1 | calcium regulated heat stable protein 1 |
| chr16 | 8737722 | 8738386 | ENSMUSG00000022710 | Usp7 | ubiquitin specific peptidase 7 |
| chr16 | 8738700 | 8739264 | ENSMUSG00000022710 | Usp7 | ubiquitin specific peptidase 7 |
| chr16 | 8829261 | 8831158 | ENSMUSG00000022507 | 1810013L24Rik | RIKEN cDNA 1810013L24 gene |
| chr16 | 10542182 | 10543191 | ENSMUSG00000038055 | Dexi | dexamethasone-induced transcript |
| chr16 | 10834944 | 10835560 | ENSMUSG00000037991 | Rmi2 | RecQ mediated genome instability 2 |
| chr16 | 11066159 | 11066723 | ENSMUSG00000037972 | Snn | stannin |
| chr16 | 11134079 | 11135004 | ENSMUSG00000022498 | Txndc11 | thioredoxin domain containing 11 |
| chr16 | 11175831 | 11176689 | ENSMUSG00000037965 | Zc3h7a | zinc finger CCCH type containing 7 A |
| chr16 | 11253075 | 11254656 | ENSMUSG00000088544 | Mir1945 | microRNA 1945 |
| chr16 | 11322724 | 11323209 | ENSMUSG00000071669 | Snx29 | sorting nexin 29 |
| chr16 | 13109560 | 13110186 | ENSMUSG00000022545 | Ercc4 | excision repair cross-complementing rodent repair deficiency, complementation group 4 |
| chr16 | 13255963 | 13257204 | ENSMUSG00000009569 | Mrtfb | myocardin related transcription factor B |
| chr16 | 13780291 | 13781131 | ENSMUSG00000022682 | Rrn3 | RRN3 RNA polymerase I transcription factor homolog (yeast) |
| chr16 | 13902520 | 13903239 | ENSMUSG00000022679 | Mpv17l | Mpv17 transgene, kidney disease mutant-like |
| chr16 | 14162937 | 14163717 | ENSMUSG00000060657 | Marf1 | meiosis regulator and mRNA stability 1 |
| chr16 | 14361328 | 14361988 | ENSMUSG00000023088 | Abcc1 | ATP-binding cassette, sub-family C (CFTR/MRP), member 1 |
| chr16 | 15593977 | 15594623 | ENSMUSG00000022674 | Ube2v2 | ubiquitin-conjugating enzyme E2 variant 2 |
| chr16 | 15886893 | 15888824 | ENSMUSG00000071637 | Cebpd | CCAAT/enhancer binding protein (C/EBP), delta |
| chr16 | 16213099 | 16213746 | ENSMUSG00000041957 | Pkp2 | plakophilin 2 |
| chr16 | 16302782 | 16303755 | ENSMUSG00000022792 | Yars2 | tyrosyl-tRNA synthetase 2 (mitochondrial) |
| chr16 | 16358429 | 16359073 | ENSMUSG00000022789 | Dnm1l | dynamin 1-like |
| chr16 | 16599421 | 16600473 | ENSMUSG00000022788 | Fgd4 | FYVE, RhoGEF and PH domain containing 4 |
| chr16 | 16983312 | 16984170 | ENSMUSG00000063358 | Mapk1 | mitogen-activated protein kinase 1 |
| chr16 | 17069773 | 17070522 | ENSMUSG00000022773 | Ypel1 | yippee like 1 |
| chr16 | 17233409 | 17234332 | ENSMUSG00000055692 | Tmem191c | transmembrane protein 191C |
| chr16 | 17276277 | 17276802 | ENSMUSG00000055692 | Tmem191c | transmembrane protein 191C |
| chr16 | 17451374 | 17452703 | ENSMUSG00000006134 | Crkl | v-crk avian sarcoma virus CT10 oncogene homolog-like |
| chr16 | 17530417 | 17531185 | ENSMUSG00000022760 | Thap7 | THAP domain containing 7 |
| chr16 | 17646296 | 17646755 | ENSMUSG00000041617 | Ccdc74a | coiled-coil domain containing 74A |
| chr16 | 17759548 | 17760347 | ENSMUSG00000022750 | Klhl22 | kelch-like 22 |
| chr16 | 17891112 | 17891846 | ENSMUSG00000003166 | Dgcr2 | DiGeorge syndrome critical region gene 2 |
| chr16 | 18127226 | 18128108 | ENSMUSG00000043811 | Rtn4r | reticulon 4 receptor |
| chr16 | 18234387 | 18235408 | ENSMUSG00000060166 | Zdhhc8 | zinc finger, DHHC domain containing 8 |
| chr16 | 18247977 | 18249963 | ENSMUSG00000022721 | Trmt2a | TRM2 tRNA methyltransferase 2A |
| chr16 | 18288735 | 18289617 | ENSMUSG00000022718 | Dgcr8 | DGCR8, microprocessor complex subunit |
| chr16 | 18347933 | 18348985 | ENSMUSG00000000325 | Arvcf | armadillo repeat gene deleted in velocardiofacial syndrome |
| chr16 | 18498346 | 18499319 | ENSMUSG00000000884 | Gnb1l | guanine nucleotide binding protein (G protein), beta polypeptide 1-like |
| chr16 | 18876335 | 18877928 | ENSMUSG00000022702 | Hira | histone cell cycle regulator |
| chr16 | 20516754 | 20517543 | ENSMUSG00000003233 | Dvl3 | dishevelled segment polarity protein 3 |
| chr16 | 21333097 | 21333530 | ENSMUSG00000116632 | Magef1 | MAGE family member F1 |
| chr16 | 21423000 | 21423749 | ENSMUSG00000033653 | Vps8 | VPS8 CORVET complex subunit |
| chr16 | 21946971 | 21947686 | ENSMUSG00000022856 | Tmem41a | transmembrane protein 41a |
| chr16 | 24393153 | 24394098 | ENSMUSG00000033306 | Lpp | LIM domain containing preferred translocation partner in lipoma |
| chr16 | 26105199 | 26106150 | ENSMUSG00000038168 | P3h2 | prolyl 3-hydroxylase 2 |
| chr16 | 26581321 | 26582050 | ENSMUSG00000022514 | Il1rap | interleukin 1 receptor accessory protein |
| chr16 | 27388679 | 27389424 | ENSMUSG00000038127 | Ccdc50 | coiled-coil domain containing 50 |
| chr16 | 29813762 | 29814357 | ENSMUSG00000075286 | Gm1968 | predicted gene 1968 |
| chr16 | 30008610 | 30009304 | ENSMUSG00000097184 | 4632428C04Rik | RIKEN cDNA 4632428C04 gene |
| chr16 | 30066025 | 30067466 | ENSMUSG00000022528 | Hes1 | hes family bHLH transcription factor 1 |
| chr16 | 30069939 | 30070887 | ENSMUSG00000022528 | Hes1 | hes family bHLH transcription factor 1 |
| chr16 | 30187880 | 30188327 | ENSMUSG00000023176 | Cpn2 | carboxypeptidase N, polypeptide 2 |
| chr16 | 30387954 | 30389038 | ENSMUSG00000022533 | Atp13a3 | ATPase type 13A3 |
| chr16 | 31080710 | 31081581 | ENSMUSG00000047434 | Xxylt1 | xyloside xylosyltransferase 1 |
| chr16 | 31274721 | 31275347 | ENSMUSG00000047714 | Ppp1r2 | protein phosphatase 1, regulatory inhibitor subunit 2 |
| chr16 | 31663280 | 31664645 | ENSMUSG00000022770 | Dlg1 | discs large MAGUK scaffold protein 1 |
| chr16 | 31948056 | 31949197 | ENSMUSG00000107002 | 0610012G03Rik | RIKEN cDNA 0610012G03 gene |
| chr16 | 32002690 | 32003379 | ENSMUSG00000022772 | Senp5 | SUMO/sentrin specific peptidase 5 |
| chr16 | 32078438 | 32079524 | ENSMUSG00000022781 | Pak2 | p21 (RAC1) activated kinase 2 |
| chr16 | 32099197 | 32100098 | ENSMUSG00000035790 | Cep19 | centrosomal protein 19 |
| chr16 | 32246283 | 32247529 | ENSMUSG00000035764 | Fbxo45 | F-box protein 45 |
| chr16 | 32430432 | 32431437 | ENSMUSG00000005615 | Pcyt1a | phosphate cytidylyltransferase 1, choline, alpha isoform |
| chr16 | 32608644 | 32609364 | ENSMUSG00000022797 | Tfrc | transferrin receptor |
| chr16 | 32877519 | 32878027 | ENSMUSG00000035629 | Rubcn | RUN domain and cysteine-rich domain containing, Beclin 1-interacting protein |
| chr16 | 33062049 | 33063090 | ENSMUSG00000022802 | Lmln | leishmanolysin-like (metallopeptidase M8 family) |
| chr16 | 33380473 | 33381722 | ENSMUSG00000097318 | 1700007L15Rik | RIKEN cDNA 1700007L15 gene |
| chr16 | 33684321 | 33684971 | ENSMUSG00000075254 | Heg1 | heart development protein with EGF-like domains 1 |
| chr16 | 33829592 | 33830478 | ENSMUSG00000022817 | Itgb5 | integrin beta 5 |
| chr16 | 34745070 | 34745575 | ENSMUSG00000022836 | Mylk | myosin, light polypeptide kinase |
| chr16 | 35490533 | 35491006 | ENSMUSG00000022844 | Pdia5 | protein disulfide isomerase associated 5 |
| chr16 | 35983147 | 35984130 | ENSMUSG00000022905 | Kpna1 | karyopherin (importin) alpha 1 |
| chr16 | 36071133 | 36072385 | ENSMUSG00000075229 | Ccdc58 | coiled-coil domain containing 58 |
| chr16 | 37867997 | 37869256 | ENSMUSG00000034158 | Lrrc58 | leucine rich repeat containing 58 |
| chr16 | 38088265 | 38090091 | ENSMUSG00000022812 | Gsk3b | glycogen synthase kinase 3 beta |
| chr16 | 38452414 | 38452939 | ENSMUSG00000002844 | Adprh | ADP-ribosylarginine hydrolase |
| chr16 | 38742148 | 38743062 | ENSMUSG00000022793 | B4galt4 | UDP-Gal:betaGlcNAc beta 1,4-galactosyltransferase, polypeptide 4 |
| chr16 | 38902177 | 38902598 | ENSMUSG00000022790 | Igsf11 | immunoglobulin superfamily, member 11 |
| chr16 | 44139653 | 44140718 | ENSMUSG00000052459 | Atp6v1a | ATPase, H+ transporting, lysosomal V1 subunit A |
| chr16 | 44347261 | 44347832 | ENSMUSG00000043065 | Spice1 | spindle and centriole associated protein 1 |
| chr16 | 49855005 | 49856486 | ENSMUSG00000055447 | Cd47 | CD47 antigen (Rh-related antigen, integrin-associated signal transducer) |
| chr16 | 50431421 | 50432657 | ENSMUSG00000022641 | Bbx | bobby sox HMG box containing |
| chr16 | 55821374 | 55822616 | ENSMUSG00000035356 | Nfkbiz | nuclear factor of kappa light polypeptide gene enhancer in B cells inhibitor, zeta |
| chr16 | 55895029 | 55895625 | ENSMUSG00000075033 | Nxpe3 | neurexophilin and PC-esterase domain family, member 3 |
| chr16 | 55973643 | 55974689 | ENSMUSG00000022601 | Zbtb11 | zinc finger and BTB domain containing 11 |
| chr16 | 56716800 | 56717441 | ENSMUSG00000022757 | Tfg | Trk-fused gene |
| chr16 | 57121342 | 57122454 | ENSMUSG00000022752 | Tomm70a | translocase of outer mitochondrial membrane 70A |
| chr16 | 57549001 | 57549585 | ENSMUSG00000043336 | Filip1l | filamin A interacting protein 1-like |
| chr16 | 58408229 | 58409366 | ENSMUSG00000035107 | Dcbld2 | discoidin, CUB and LCCL domain containing 2 |
| chr16 | 58522726 | 58523657 | ENSMUSG00000022747 | St3gal6 | ST3 beta-galactoside alpha-2,3-sialyltransferase 6 |
| chr16 | 58670154 | 58670869 | ENSMUSG00000022742 | Cpox | coproporphyrinogen oxidase |
| chr16 | 58727862 | 58728412 | ENSMUSG00000022744 | Cldnd1 | claudin domain containing 1 |
| chr16 | 59600543 | 59601302 | ENSMUSG00000022723 | Crybg3 | beta-gamma crystallin domain containing 3 |
| chr16 | 59639126 | 59639611 | ENSMUSG00000022722 | Arl6 | ADP-ribosylation factor-like 6 |
| chr16 | 64851061 | 64852915 | ENSMUSG00000054604 | Cggbp1 | CGG triplet repeat binding protein 1 |
| chr16 | 65814840 | 65816072 | ENSMUSG00000091243 | Vgll3 | vestigial like family member 3 |
| chr16 | 70313868 | 70314675 | ENSMUSG00000022707 | Gbe1 | glucan (1,4-alpha-), branching enzyme 1 |
| chr16 | 76372161 | 76373592 | ENSMUSG00000048490 | Nrip1 | nuclear receptor interacting protein 1 |
| chr16 | 77013613 | 77014389 | ENSMUSG00000022867 | Usp25 | ubiquitin specific peptidase 25 |
| chr16 | 78301230 | 78302452 | ENSMUSG00000022865 | Cxadr | coxsackie virus and adenovirus receptor |
| chr16 | 78375883 | 78377186 | ENSMUSG00000022863 | Btg3 | BTG anti-proliferation factor 3 |
| chr16 | 87454924 | 87455549 | ENSMUSG00000025616 | Usp16 | ubiquitin specific peptidase 16 |
| chr16 | 87698722 | 87700278 | ENSMUSG00000025612 | Bach1 | BTB and CNC homology 1, basic leucine zipper transcription factor 1 |
| chr16 | 90143206 | 90144450 | ENSMUSG00000002489 | Tiam1 | T cell lymphoma invasion and metastasis 1 |
| chr16 | 90220546 | 90221277 | ENSMUSG00000022982 | Sod1 | superoxide dismutase 1, soluble |
| chr16 | 90283516 | 90284747 | ENSMUSG00000022983 | Scaf4 | SR-related CTD-associated factor 4 |
| chr16 | 91010752 | 91011681 | ENSMUSG00000022973 | Synj1 | synaptojanin 1 |
| chr16 | 91043670 | 91044705 | ENSMUSG00000022974 | Paxbp1 | PAX3 and PAX7 binding protein 1 |
| chr16 | 91406116 | 91406690 | ENSMUSG00000022969 | Il10rb | interleukin 10 receptor, beta |
| chr16 | 91646704 | 91648083 | ENSMUSG00000022962 | Gart | phosphoribosylglycinamide formyltransferase |
| chr16 | 91688363 | 91689050 | ENSMUSG00000085169 | Gm10785 | predicted gene 10785 |
| chr16 | 91729147 | 91730214 | ENSMUSG00000022957 | Itsn1 | intersectin 1 (SH3 domain protein 1A) |
| chr16 | 92057811 | 92059007 | ENSMUSG00000039680 | Mrps6 | mitochondrial ribosomal protein S6 |
| chr16 | 92465712 | 92466404 | ENSMUSG00000022951 | Rcan1 | regulator of calcineurin 1 |
| chr16 | 92696697 | 92697918 | ENSMUSG00000022952 | Runx1 | runt related transcription factor 1 |
| chr16 | 92697918 | 92698798 | ENSMUSG00000022952 | Runx1 | runt related transcription factor 1 |
| chr16 | 93603475 | 93604322 | ENSMUSG00000022948 | Setd4 | SET domain containing 4 |
| chr16 | 93831978 | 93832887 | ENSMUSG00000039456 | Morc3 | microrchidia 3 |
| chr16 | 94289795 | 94290534 | ENSMUSG00000040820 | Hlcs | holocarboxylase synthetase (biotin- [propriony-Coenzyme A-carboxylase (ATP-hydrolysing)] ligase) |
| chr16 | 94313279 | 94313717 | ENSMUSG00000040820 | Hlcs | holocarboxylase synthetase (biotin- [propriony-Coenzyme A-carboxylase (ATP-hydrolysing)] ligase) |
| chr16 | 94370315 | 94371775 | ENSMUSG00000040785 | Ttc3 | tetratricopeptide repeat domain 3 |
| chr16 | 94526058 | 94527234 | ENSMUSG00000022898 | Vps26c | VPS26 endosomal protein sorting factor C |
| chr16 | 94569776 | 94571195 | ENSMUSG00000022897 | Dyrk1a | dual-specificity tyrosine-(Y)-phosphorylation regulated kinase 1a |
| chr16 | 95701945 | 95703010 | ENSMUSG00000022895 | Ets2 | E26 avian leukemia oncogene 2, 3' domain |
| chr16 | 96127108 | 96128260 | ENSMUSG00000040681 | Hmgn1 | high mobility group nucleosomal binding domain 1 |
| chr16 | 96145185 | 96145891 | ENSMUSG00000023147 | Get1 | guided entry of tail-anchored proteins factor 1 |
| chr16 | 97763184 | 97763904 | ENSMUSG00000005251 | Ripk4 | receptor-interacting serine-threonine kinase 4 |
| chr16 | 97851010 | 97852078 | ENSMUSG00000014039 | Prdm15 | PR domain containing 15 |
| chr16 | 97922009 | 97922857 | ENSMUSG00000045975 | C2cd2 | C2 calcium-dependent domain containing 2 |
| chr16 | 97961789 | 97962771 | ENSMUSG00000046962 | Zbtb21 | zinc finger and BTB domain containing 21 |
| chr17 | 3114293 | 3115871 | ENSMUSG00000046201 | Scaf8 | SR-related CTD-associated factor 8 |
| chr17 | 4908004 | 4908543 | ENSMUSG00000069729 | Arid1b | AT rich interactive domain 1B (SWI-like) |
| chr17 | 5189599 | 5190246 | ENSMUSG00000069729 | Arid1b | AT rich interactive domain 1B (SWI-like) |
| chr17 | 5840879 | 5842314 | ENSMUSG00000002365 | Snx9 | sorting nexin 9 |
| chr17 | 6782172 | 6783632 | ENSMUSG00000052397 | Ezr | ezrin |
| chr17 | 6827785 | 6828433 | ENSMUSG00000052397 | Ezr | ezrin |
| chr17 | 8283277 | 8284667 | ENSMUSG00000023861 | Mpc1 | mitochondrial pyruvate carrier 1 |
| chr17 | 12318053 | 12318998 | ENSMUSG00000097350 | 4732491K20Rik | RIKEN cDNA 4732491K20 gene |
| chr17 | 12991726 | 12992946 | ENSMUSG00000060475 | Wtap | Wilms tumour 1-associating protein |
| chr17 | 13007716 | 13008418 |  | Gm36117 | predicted gene, 36117 |
| chr17 | 14960677 | 14961281 | ENSMUSG00000023883 | Phf10 | PHD finger protein 10 |
| chr17 | 15374697 | 15375553 | ENSMUSG00000014773 | Dll1 | delta like canonical Notch ligand 1 |
| chr17 | 15526858 | 15527383 | ENSMUSG00000014771 | Pdcd2 | programmed cell death 2 |
| chr17 | 15704385 | 15706085 | ENSMUSG00000023852 | Chd1 | chromodomain helicase DNA binding protein 1 |
| chr17 | 15825691 | 15826839 | ENSMUSG00000048027 | Rgmb | repulsive guidance molecule family member B |
| chr17 | 15830530 | 15831337 | ENSMUSG00000048027 | Rgmb | repulsive guidance molecule family member B |
| chr17 | 23699366 | 23700371 | ENSMUSG00000066720 | Cldn9 | claudin 9 |
| chr17 | 24140977 | 24141816 | ENSMUSG00000024122 | Pdpk1 | 3-phosphoinositide dependent protein kinase 1 |
| chr17 | 24163198 | 24163899 | ENSMUSG00000036820 | Amdhd2 | amidohydrolase domain containing 2 |
| chr17 | 24250947 | 24251799 | ENSMUSG00000072082 | Ccnf | cyclin F |
| chr17 | 24426531 | 24427286 | ENSMUSG00000024132 | Eci1 | enoyl-Coenzyme A delta isomerase 1 |
| chr17 | 24469968 | 24471453 | ENSMUSG00000024137 | E4f1 | E4F transcription factor 1 |
| chr17 | 24549849 | 24550795 | ENSMUSG00000032855 | Pkd1 | polycystin 1, transient receptor poteintial channel interacting |
| chr17 | 24632369 | 24632969 | ENSMUSG00000041429 | Nthl1 | nth (endonuclease III)-like 1 (E.coli) |
| chr17 | 24649803 | 24650339 | ENSMUSG00000002504 | Slc9a3r2 | solute carrier family 9 (sodium/hydrogen exchanger), member 3 regulator 2 |
| chr17 | 24669503 | 24670219 | ENSMUSG00000041130 | Zfp598 | zinc finger protein 598 |
| chr17 | 24695723 | 24696762 | ENSMUSG00000040888 | Gfer | growth factor, augmenter of liver regeneration |
| chr17 | 24849913 | 24850994 | ENSMUSG00000024158 | Hagh | hydroxyacyl glutathione hydrolase |
| chr17 | 24960207 | 24961003 | ENSMUSG00000024165 | Jpt2 | Jupiter microtubule associated homolog 2 |
| chr17 | 25272469 | 25274344 | ENSMUSG00000015120 | Ube2i | ubiquitin-conjugating enzyme E2I |
| chr17 | 25868017 | 25868905 | ENSMUSG00000025732 | Mcrip2 | MAPK regulated corepressor interacting protein 2 |
| chr17 | 25985041 | 25985856 | ENSMUSG00000037326 | Capn15 | calpain 15 |
| chr17 | 26067703 | 26069676 | ENSMUSG00000037098 | Rab11fip3 | RAB11 family interacting protein 3 (class II) |
| chr17 | 26113228 | 26113806 | ENSMUSG00000024180 | Pgap6 | post-glycosylphosphatidylinositol attachment to proteins 6 |
| chr17 | 26138291 | 26139371 | ENSMUSG00000024182 | Axin1 | axin 1 |
| chr17 | 26507838 | 26509019 | ENSMUSG00000024190 | Dusp1 | dual specificity phosphatase 1 |
| chr17 | 26972761 | 26973443 | ENSMUSG00000079605 | Zbtb9 | zinc finger and BTB domain containing 9 |
| chr17 | 27056974 | 27057890 | ENSMUSG00000042644 | Itpr3 | inositol 1,4,5-triphosphate receptor 3 |
| chr17 | 27203244 | 27204686 | ENSMUSG00000044857 | Lemd2 | LEM domain containing 2 |
| chr17 | 27556183 | 27557572 | ENSMUSG00000078249 | Hmga1b | high mobility group AT-hook 1B |
| chr17 | 27557572 | 27558043 | ENSMUSG00000046711 | Hmga1 | high mobility group AT-hook 1 |
| chr17 | 27622638 | 27623919 | ENSMUSG00000024213 | Nudt3 | nudix (nucleotide diphosphate linked moiety X)-type motif 3 |
| chr17 | 27819847 | 27820680 | ENSMUSG00000056692 | Ilrun | inflammation and lipid regulator with UBA-like and NBR1-like domains |
| chr17 | 27856334 | 27857255 | ENSMUSG00000039512 | Uhrf1bp1 | UHRF1 (ICBP90) binding protein 1 |
| chr17 | 27908820 | 27910015 | ENSMUSG00000024219 | Anks1 | ankyrin repeat and SAM domain containing 1 |
| chr17 | 28328300 | 28329609 | ENSMUSG00000037805 | Rpl10a | ribosomal protein L10A |
| chr17 | 28349776 | 28351029 | ENSMUSG00000002249 | Tead3 | TEA domain family member 3 |
| chr17 | 28484792 | 28486395 | ENSMUSG00000024222 | Fkbp5 | FK506 binding protein 5 |
| chr17 | 28517880 | 28518726 | ENSMUSG00000024222 | Fkbp5 | FK506 binding protein 5 |
| chr17 | 28522939 | 28523446 | ENSMUSG00000024222 | Fkbp5 | FK506 binding protein 5 |
| chr17 | 28621597 | 28622640 | ENSMUSG00000004865 | Srpk1 | serine/arginine-rich protein specific kinase 1 |
| chr17 | 28690928 | 28692533 | ENSMUSG00000053436 | Mapk14 | mitogen-activated protein kinase 14 |
| chr17 | 28800768 | 28802368 | ENSMUSG00000063952 | Brpf3 | bromodomain and PHD finger containing, 3 |
| chr17 | 28952222 | 28952841 | ENSMUSG00000005936 | Kctd20 | potassium channel tetramerisation domain containing 20 |
| chr17 | 29032054 | 29033111 | ENSMUSG00000071172 | Srsf3 | serine and arginine-rich splicing factor 3 |
| chr17 | 29268823 | 29269299 | ENSMUSG00000052712 | BC004004 | cDNA sequence BC004004 |
| chr17 | 29347195 | 29348112 | ENSMUSG00000024012 | Mtch1 | mitochondrial carrier 1 |
| chr17 | 29614581 | 29615401 |  | Gm28043 | predicted gene, 28043 |
| chr17 | 29660506 | 29661153 | ENSMUSG00000024019 | Cmtr1 | cap methyltransferase 1 |
| chr17 | 30004281 | 30005649 | ENSMUSG00000044477 | Zfand3 | zinc finger, AN1-type domain 3 |
| chr17 | 30575817 | 30576509 | ENSMUSG00000062202 | Btbd9 | BTB (POZ) domain containing 9 |
| chr17 | 31511946 | 31512435 | ENSMUSG00000024037 | Wdr4 | WD repeat domain 4 |
| chr17 | 31564322 | 31565868 | ENSMUSG00000006705 | Pknox1 | Pbx/knotted 1 homeobox |
| chr17 | 31658110 | 31659078 | ENSMUSG00000061613 | U2af1 | U2 small nuclear ribonucleoprotein auxiliary factor (U2AF) 1 |
| chr17 | 31854951 | 31856062 | ENSMUSG00000024042 | Sik1 | salt inducible kinase 1 |
| chr17 | 32036041 | 32036752 | ENSMUSG00000058392 | Rrp1b | ribosomal RNA processing 1B |
| chr17 | 32283643 | 32284977 | ENSMUSG00000024002 | Brd4 | bromodomain containing 4 |
| chr17 | 33817202 | 33818004 | ENSMUSG00000042099 | Kank3 | KN motif and ankyrin repeat domains 3 |
| chr17 | 34027458 | 34027989 | ENSMUSG00000073422 | H2-Ke6 | H2-K region expressed gene 6 |
| chr17 | 34587297 | 34587926 | ENSMUSG00000015468 | Notch4 | notch 4 |
| chr17 | 34981704 | 34982370 | ENSMUSG00000007050 | Lsm2 | LSM2 homolog, U6 small nuclear RNA and mRNA degradation associated |
| chr17 | 35000704 | 35001892 | ENSMUSG00000007029 | Vars | valyl-tRNA synthetase |
| chr17 | 35089036 | 35089714 | ENSMUSG00000007036 | Abhd16a | abhydrolase domain containing 16A |
| chr17 | 35699816 | 35700874 | ENSMUSG00000003534 | Ddr1 | discoidin domain receptor family, member 1 |
| chr17 | 35700874 | 35701489 | ENSMUSG00000003534 | Ddr1 | discoidin domain receptor family, member 1 |
| chr17 | 35896677 | 35897483 | ENSMUSG00000050705 | 2310061I04Rik | RIKEN cDNA 2310061I04 gene |
| chr17 | 35969287 | 35970023 | ENSMUSG00000038762 | Abcf1 | ATP-binding cassette, sub-family F (GCN20), member 1 |
| chr17 | 35978889 | 35980698 | ENSMUSG00000024429 | Gnl1 | guanine nucleotide binding protein-like 1 |
| chr17 | 36270916 | 36271920 | ENSMUSG00000045409 | Trim39 | tripartite motif-containing 39 |
| chr17 | 36837050 | 36837606 | ENSMUSG00000024457 | Trim26 | tripartite motif-containing 26 |
| chr17 | 36942331 | 36943510 | ENSMUSG00000036492 | Rnf39 | ring finger protein 39 |
| chr17 | 42875833 | 42876903 | ENSMUSG00000061665 | Cd2ap | CD2-associated protein |
| chr17 | 46032417 | 46032916 | ENSMUSG00000023951 | Vegfa | vascular endothelial growth factor A |
| chr17 | 46202304 | 46203122 | ENSMUSG00000067150 | Xpo5 | exportin 5 |
| chr17 | 46383356 | 46384473 | ENSMUSG00000015597 | Zfp318 | zinc finger protein 318 |
| chr17 | 46555116 | 46556393 | ENSMUSG00000015605 | Srf | serum response factor |
| chr17 | 46890556 | 46891442 | ENSMUSG00000036430 | Tbcc | tubulin-specific chaperone C |
| chr17 | 47436325 | 47437380 | ENSMUSG00000034382 | AI661453 | expressed sequence AI661453 |
| chr17 | 47736820 | 47738149 | ENSMUSG00000023990 | Tfeb | transcription factor EB |
| chr17 | 47923989 | 47925277 | ENSMUSG00000023991 | Foxp4 | forkhead box P4 |
| chr17 | 51178513 | 51179590 | ENSMUSG00000023923 | Tbc1d5 | TBC1 domain family, member 5 |
| chr17 | 56218338 | 56219137 | ENSMUSG00000001229 | Dpp9 | dipeptidylpeptidase 9 |
| chr17 | 56325761 | 56326417 | ENSMUSG00000024201 | Kdm4b | lysine (K)-specific demethylase 4B |
| chr17 | 56475983 | 56476733 | ENSMUSG00000013236 | Ptprs | protein tyrosine phosphatase, receptor type, S |
| chr17 | 56584312 | 56585496 | ENSMUSG00000042625 | Safb2 | scaffold attachment factor B2 |
| chr17 | 56756975 | 56757785 | ENSMUSG00000039481 | Nrtn | neurturin |
| chr17 | 56935012 | 56935779 | ENSMUSG00000024212 | Mllt1 | myeloid/lymphoid or mixed-lineage leukemia; translocated to, 1 |
| chr17 | 62882757 | 62884418 | ENSMUSG00000048915 | Efna5 | ephrin A5 |
| chr17 | 63498916 | 63500657 | ENSMUSG00000023965 | Fbxl17 | F-box and leucine-rich repeat protein 17 |
| chr17 | 63863340 | 63864647 | ENSMUSG00000000127 | Fer | fer (fms/fps related) protein kinase |
| chr17 | 64331225 | 64332165 | ENSMUSG00000024083 | Pja2 | praja ring finger ubiquitin ligase 2 |
| chr17 | 64600541 | 64602198 | ENSMUSG00000024085 | Man2a1 | mannosidase 2, alpha 1 |
| chr17 | 65772141 | 65772756 | ENSMUSG00000056515 | Rab31 | RAB31, member RAS oncogene family |
| chr17 | 65782551 | 65784135 | ENSMUSG00000061950 | Ppp4r1 | protein phosphatase 4, regulatory subunit 1 |
| chr17 | 65884084 | 65885609 | ENSMUSG00000024096 | Ralbp1 | ralA binding protein 1 |
| chr17 | 65950660 | 65951370 | ENSMUSG00000024098 | Twsg1 | twisted gastrulation BMP signaling modulator 1 |
| chr17 | 66076249 | 66077558 | ENSMUSG00000034647 | Ankrd12 | ankyrin repeat domain 12 |
| chr17 | 67353304 | 67355169 | ENSMUSG00000033278 | Ptprm | protein tyrosine phosphatase, receptor type, M |
| chr17 | 69156566 | 69157541 | ENSMUSG00000024044 | Epb41l3 | erythrocyte membrane protein band 4.1 like 3 |
| chr17 | 69382373 | 69384366 | ENSMUSG00000049672 | Zbtb14 | zinc finger and BTB domain containing 14 |
| chr17 | 70990100 | 70990985 | ENSMUSG00000034868 | Myl12b | myosin, light chain 12B, regulatory |
| chr17 | 71183324 | 71184220 | ENSMUSG00000024052 | Lpin2 | lipin 2 |
| chr17 | 71474611 | 71476083 | ENSMUSG00000024054 | Smchd1 | SMC hinge domain containing 1 |
| chr17 | 71551794 | 71552472 | ENSMUSG00000052525 | Spdya | speedy/RINGO cell cycle regulator family, member A |
| chr17 | 71616091 | 71616956 | ENSMUSG00000041057 | Wdr43 | WD repeat domain 43 |
| chr17 | 72836903 | 72837841 | ENSMUSG00000039770 | Ypel5 | yippee like 5 |
| chr17 | 74294263 | 74295671 | ENSMUSG00000058704 | Memo1 | mediator of cell motility 1 |
| chr17 | 74315933 | 74316882 | ENSMUSG00000024067 | Dpy30 | dpy-30, histone methyltransferase complex regulatory subunit |
| chr17 | 74338611 | 74339750 | ENSMUSG00000024068 | Spast | spastin |
| chr17 | 74489477 | 74490084 | ENSMUSG00000024072 | Yipf4 | Yip1 domain family, member 4 |
| chr17 | 74527729 | 74529421 | ENSMUSG00000024073 | Birc6 | baculoviral IAP repeat-containing 6 |
| chr17 | 78199797 | 78201596 | ENSMUSG00000024074 | Crim1 | cysteine rich transmembrane BMP regulator 1 (chordin like) |
| chr17 | 78736161 | 78737017 | ENSMUSG00000024077 | Strn | striatin, calmodulin binding protein |
| chr17 | 79354407 | 79355835 | ENSMUSG00000036533 | Cdc42ep3 | CDC42 effector protein (Rho GTPase binding) 3 |
| chr17 | 79894971 | 79896344 | ENSMUSG00000059811 | Atl2 | atlastin GTPase 2 |
| chr17 | 80061121 | 80062570 | ENSMUSG00000024095 | Hnrnpll | heterogeneous nuclear ribonucleoprotein L-like |
| chr17 | 80206239 | 80207439 | ENSMUSG00000024097 | Srsf7 | serine and arginine-rich splicing factor 7 |
| chr17 | 80479484 | 80480521 | ENSMUSG00000024241 | Sos1 | SOS Ras/Rac guanine nucleotide exchange factor 1 |
| chr17 | 80727416 | 80728526 | ENSMUSG00000024242 | Map4k3 | mitogen-activated protein kinase kinase kinase kinase 3 |
| chr17 | 80728526 | 80728777 | ENSMUSG00000024242 | Map4k3 | mitogen-activated protein kinase kinase kinase kinase 3 |
| chr17 | 83350536 | 83351582 | ENSMUSG00000032624 | Eml4 | echinoderm microtubule associated protein like 4 |
| chr17 | 84187760 | 84188912 | ENSMUSG00000045817 | Zfp36l2 | zinc finger protein 36, C3H type-like 2 |
| chr17 | 84958053 | 84958752 | ENSMUSG00000061130 | Ppm1b | protein phosphatase 1B, magnesium dependent, beta isoform |
| chr17 | 86166394 | 86168177 | ENSMUSG00000045038 | Prkce | protein kinase C, epsilon |
| chr17 | 86962468 | 86963676 | ENSMUSG00000024143 | Rhoq | ras homolog family member Q |
| chr17 | 87107310 | 87108000 | ENSMUSG00000037104 | Socs5 | suppressor of cytokine signaling 5 |
| chr17 | 87265519 | 87266226 | ENSMUSG00000024150 | Mcfd2 | multiple coagulation factor deficiency 2 |
| chr17 | 87282326 | 87283543 | ENSMUSG00000036918 | Ttc7 | tetratricopeptide repeat domain 7 |
| chr17 | 87445866 | 87447480 | ENSMUSG00000036438 | Calm2 | calmodulin 2 |
| chr17 | 87636286 | 87637314 | ENSMUSG00000045394 | Epcam | epithelial cell adhesion molecule |
| chr17 | 87672426 | 87673097 | ENSMUSG00000024151 | Msh2 | mutS homolog 2 |
| chr17 | 87974602 | 87975747 | ENSMUSG00000005370 | Msh6 | mutS homolog 6 |
| chr17 | 88064655 | 88066591 | ENSMUSG00000005371 | Fbxo11 | F-box protein 11 |
| chr17 | 88440183 | 88441916 | ENSMUSG00000034998 | Foxn2 | forkhead box N2 |
| chr18 | 3336668 | 3338003 | ENSMUSG00000063889 | Crem | cAMP responsive element modulator |
| chr18 | 6489900 | 6491306 | ENSMUSG00000084514 | Mir1893 | microRNA 1893 |
| chr18 | 6515847 | 6516604 | ENSMUSG00000024240 | Epc1 | enhancer of polycomb homolog 1 |
| chr18 | 7868497 | 7870095 | ENSMUSG00000024283 | Wac | WW domain containing adaptor with coiled-coil |
| chr18 | 9212309 | 9213744 | ENSMUSG00000036904 | Fzd8 | frizzled class receptor 8 |
| chr18 | 9449191 | 9450512 | ENSMUSG00000024286 | Ccny | cyclin Y |
| chr18 | 10181290 | 10182487 | ENSMUSG00000024290 | Rock1 | Rho-associated coiled-coil containing protein kinase 1 |
| chr18 | 10609479 | 10610517 | ENSMUSG00000024293 | Esco1 | establishment of sister chromatid cohesion N-acetyltransferase 1 |
| chr18 | 10725485 | 10726494 | ENSMUSG00000024294 | Mib1 | mindbomb E3 ubiquitin protein ligase 1 |
| chr18 | 14682101 | 14683345 | ENSMUSG00000037013 | Ss18 | SS18, nBAF chromatin remodeling complex subunit |
| chr18 | 14782917 | 14783970 | ENSMUSG00000054321 | Taf4b | TATA-box binding protein associated factor 4b |
| chr18 | 20745563 | 20746530 | ENSMUSG00000056124 | B4galt6 | UDP-Gal:betaGlcNAc beta 1,4-galactosyltransferase, polypeptide 6 |
| chr18 | 21001242 | 21002373 | ENSMUSG00000024317 | Rnf138 | ring finger protein 138 |
| chr18 | 21299681 | 21300671 | ENSMUSG00000042680 | Garem1 | GRB2 associated regulator of MAPK1 subtype 1 |
| chr18 | 24205132 | 24206378 | ENSMUSG00000000420 | Galnt1 | polypeptide N-acetylgalactosaminyltransferase 1 |
| chr18 | 24709067 | 24710166 | ENSMUSG00000034295 | Fhod3 | formin homology 2 domain containing 3 |
| chr18 | 31910384 | 31911343 | ENSMUSG00000044982 | Sft2d3 | SFT2 domain containing 3 |
| chr18 | 34220873 | 34221556 | ENSMUSG00000005871 | Apc | APC, WNT signaling pathway regulator |
| chr18 | 34860387 | 34861418 | ENSMUSG00000038418 | Egr1 | early growth response 1 |
| chr18 | 35498550 | 35499067 | ENSMUSG00000024357 | Sil1 | endoplasmic reticulum chaperone SIL1 homolog (S. cerevisiae) |
| chr18 | 35562034 | 35562888 | ENSMUSG00000037236 | Matr3 | matrin 3 |
| chr18 | 35829581 | 35831172 | ENSMUSG00000046668 | Cxxc5 | CXXC finger 5 |
| chr18 | 36196526 | 36197564 | ENSMUSG00000060275 | Nrg2 | neuregulin 2 |
| chr18 | 36280744 | 36281660 | ENSMUSG00000043991 | Pura | purine rich element binding protein A |
| chr18 | 36286616 | 36287814 | ENSMUSG00000043991 | Pura | purine rich element binding protein A |
| chr18 | 36515185 | 36516043 | ENSMUSG00000024486 | Hbegf | heparin-binding EGF-like growth factor |
| chr18 | 38418855 | 38419412 | ENSMUSG00000024425 | Ndfip1 | Nedd4 family interacting protein 1 |
| chr18 | 39488808 | 39490728 | ENSMUSG00000024431 | Nr3c1 | nuclear receptor subfamily 3, group C, member 1 |
| chr18 | 42511501 | 42512198 | ENSMUSG00000024498 | Tcerg1 | transcription elongation regulator 1 (CA150) |
| chr18 | 46198467 | 46198982 | ENSMUSG00000033949 | Trim36 | tripartite motif-containing 36 |
| chr18 | 46525103 | 46526058 | ENSMUSG00000033319 | Fem1c | fem 1 homolog c |
| chr18 | 46741968 | 46742522 | ENSMUSG00000024480 | Ap3s1 | adaptor-related protein complex 3, sigma 1 subunit |
| chr18 | 49755236 | 49755877 | ENSMUSG00000024505 | Dtwd2 | DTW domain containing 2 |
| chr18 | 49832552 | 49833271 | ENSMUSG00000037416 | Dmxl1 | Dmx-like 1 |
| chr18 | 53245417 | 53246104 | ENSMUSG00000024535 | Snx24 | sorting nexing 24 |
| chr18 | 56707446 | 56709149 | ENSMUSG00000024590 | Lmnb1 | lamin B1 |
| chr18 | 57877914 | 57879760 | ENSMUSG00000024597 | Slc12a2 | solute carrier family 12, member 2 |
| chr18 | 60747838 | 60749022 | ENSMUSG00000054008 | Ndst1 | N-deacetylase/N-sulfotransferase (heparan glucosaminyl) 1 |
| chr18 | 61399363 | 61400556 | ENSMUSG00000033871 | Ppargc1b | peroxisome proliferative activated receptor, gamma, coactivator 1 beta |
| chr18 | 63691802 | 63692643 | ENSMUSG00000024583 | Txnl1 | thioredoxin-like 1 |
| chr18 | 67723907 | 67724842 | ENSMUSG00000024539 | Ptpn2 | protein tyrosine phosphatase, non-receptor type 2 |
| chr18 | 67774710 | 67775623 | ENSMUSG00000079614 | Seh1l | SEH1-like (S. cerevisiae |
| chr18 | 70529939 | 70530462 | ENSMUSG00000038425 | Poli | polymerase (DNA directed), iota |
| chr18 | 73572020 | 73573849 | ENSMUSG00000037253 | Mex3c | mex3 RNA binding family member C |
| chr18 | 73754080 | 73754657 | ENSMUSG00000036941 | Elac1 | elaC ribonuclease Z 1 |
| chr18 | 73815118 | 73815922 | ENSMUSG00000024556 | Me2 | malic enzyme 2, NAD(+)-dependent, mitochondrial |
| chr18 | 75342521 | 75343301 |  | Gm20544 | predicted gene 20544 |
| chr18 | 75366426 | 75367169 |  | Gm20544 | predicted gene 20544 |
| chr18 | 75367169 | 75368013 | ENSMUSG00000025880 | Smad7 | SMAD family member 7 |
| chr18 | 75368483 | 75369636 | ENSMUSG00000025880 | Smad7 | SMAD family member 7 |
| chr18 | 76241131 | 76242504 | ENSMUSG00000024563 | Smad2 | SMAD family member 2 |
| chr18 | 77065153 | 77066001 | ENSMUSG00000025423 | Pias2 | protein inhibitor of activated STAT 2 |
| chr18 | 80150731 | 80151579 | ENSMUSG00000053950 | Adnp2 | ADNP homeobox 2 |
| chr18 | 84085076 | 84086154 | ENSMUSG00000046982 | Tshz1 | teashirt zinc finger family member 1 |
| chr18 | 84087305 | 84088534 | ENSMUSG00000046982 | Tshz1 | teashirt zinc finger family member 1 |
| chr19 | 3767476 | 3769031 | ENSMUSG00000045098 | Kmt5b | lysine methyltransferase 5B |
| chr19 | 3851386 | 3852689 | ENSMUSG00000024843 | Chka | choline kinase alpha |
| chr19 | 4169356 | 4170084 | ENSMUSG00000075289 | Carns1 | carnosine synthase 1 |
| chr19 | 4214256 | 4215674 |  | Gm45928 | predicted gene, 45928 |
| chr19 | 4305566 | 4306441 | ENSMUSG00000024858 | Grk2 | G protein-coupled receptor kinase 2 |
| chr19 | 4396398 | 4397527 | ENSMUSG00000054611 | Kdm2a | lysine (K)-specific demethylase 2A |
| chr19 | 4439091 | 4439842 | ENSMUSG00000041845 | Rhod | ras homolog family member D |
| chr19 | 4476688 | 4477249 | ENSMUSG00000049303 | Syt12 | synaptotagmin XII |
| chr19 | 4614894 | 4616030 | ENSMUSG00000045045 | Lrfn4 | leucine rich repeat and fibronectin type III domain containing 4 |
| chr19 | 4625270 | 4626181 | ENSMUSG00000024889 | Rce1 | Ras converting CAAX endopeptidase 1 |
| chr19 | 5272527 | 5273244 | ENSMUSG00000024855 | Pacs1 | phosphofurin acidic cluster sorting protein 1 |
| chr19 | 5456583 | 5457703 | ENSMUSG00000095098 | Ccdc85b | coiled-coil domain containing 85B |
| chr19 | 5567145 | 5569081 | ENSMUSG00000049562 | Ap5b1 | adaptor-related protein complex 5, beta 1 subunit |
| chr19 | 5637017 | 5637960 | ENSMUSG00000024927 | Rela | v-rel reticuloendotheliosis viral oncogene homolog A (avian) |
| chr19 | 5741656 | 5742590 | ENSMUSG00000024940 | Ltbp3 | latent transforming growth factor beta binding protein 3 |
| chr19 | 6117733 | 6118628 | ENSMUSG00000024790 | Sac3d1 | SAC3 domain containing 1 |
| chr19 | 6276349 | 6278083 | ENSMUSG00000024772 | Ehd1 | EH-domain containing 1 |
| chr19 | 6306240 | 6306987 | ENSMUSG00000024769 | Cdc42bpg | CDC42 binding protein kinase gamma (DMPK-like) |
| chr19 | 6921293 | 6922076 | ENSMUSG00000024955 | Esrra | estrogen related receptor, alpha |
| chr19 | 6987012 | 6987890 | ENSMUSG00000024962 | Vegfb | vascular endothelial growth factor B |
| chr19 | 7056533 | 7057220 | ENSMUSG00000036278 | Macrod1 | mono-ADP ribosylhydrolase 1 |
| chr19 | 7057220 | 7057806 | ENSMUSG00000036278 | Macrod1 | mono-ADP ribosylhydrolase 1 |
| chr19 | 7340937 | 7342172 | ENSMUSG00000024969 | Mark2 | MAP/microtubule affinity regulating kinase 2 |
| chr19 | 7416828 | 7418389 | ENSMUSG00000053080 | 2700081O15Rik | RIKEN cDNA 2700081O15 gene |
| chr19 | 7494166 | 7495023 | ENSMUSG00000024759 | Atl3 | atlastin GTPase 3 |
| chr19 | 8774247 | 8775094 | ENSMUSG00000118346 | Tmem179b | transmembrane protein 179B |
| chr19 | 8819741 | 8820861 | ENSMUSG00000071659 | Hnrnpul2 | heterogeneous nuclear ribonucleoprotein U-like 2 |
| chr19 | 10100954 | 10102071 | ENSMUSG00000024665 | Fads2 | fatty acid desaturase 2 |
| chr19 | 10182828 | 10183454 | ENSMUSG00000010663 | Fads1 | fatty acid desaturase 1 |
| chr19 | 11818567 | 11819104 | ENSMUSG00000041488 | Stx3 | syntaxin 3 |
| chr19 | 11912017 | 11913192 | ENSMUSG00000046139 | Patl1 | protein associated with topoisomerase II homolog 1 (yeast) |
| chr19 | 12475434 | 12475774 | ENSMUSG00000046805 | Mpeg1 | macrophage expressed gene 1 |
| chr19 | 12500857 | 12501991 | ENSMUSG00000039982 | Dtx4 | deltex 4, E3 ubiquitin ligase |
| chr19 | 14597049 | 14598256 | ENSMUSG00000024642 | Tle4 | transducin-like enhancer of split 4 |
| chr19 | 15924212 | 15925575 | ENSMUSG00000024640 | Psat1 | phosphoserine aminotransferase 1 |
| chr19 | 15984504 | 15985198 | ENSMUSG00000097557 | C130060C02Rik | RIKEN cDNA C130060C02 gene |
| chr19 | 16132173 | 16134038 | ENSMUSG00000024639 | Gnaq | guanine nucleotide binding protein, alpha q polypeptide |
| chr19 | 16780186 | 16781202 | ENSMUSG00000046230 | Vps13a | vacuolar protein sorting 13A |
| chr19 | 18631101 | 18632128 | ENSMUSG00000024725 | Ostf1 | osteoclast stimulating factor 1 |
| chr19 | 18670605 | 18671748 | ENSMUSG00000024726 | Carnmt1 | carnosine N-methyltransferase 1 |
| chr19 | 18712821 | 18713795 | ENSMUSG00000047044 | D030056L22Rik | RIKEN cDNA D030056L22 gene |
| chr19 | 21271697 | 21273245 | ENSMUSG00000024750 | Zfand5 | zinc finger, AN1-type domain 5 |
| chr19 | 21652838 | 21653896 | ENSMUSG00000047368 | Abhd17b | abhydrolase domain containing 17B |
| chr19 | 21778104 | 21778970 | ENSMUSG00000024754 | Cemip2 | cell migration inducing hyaluronidase 2 |
| chr19 | 23140696 | 23142383 | ENSMUSG00000033863 | Klf9 | Kruppel-like factor 9 |
| chr19 | 23687141 | 23687967 | ENSMUSG00000074925 | Ptar1 | protein prenyltransferase alpha subunit repeat containing 1 |
| chr19 | 24173765 | 24174641 | ENSMUSG00000024812 | Tjp2 | tight junction protein 2 |
| chr19 | 24999095 | 25000042 | ENSMUSG00000052085 | Dock8 | dedicator of cytokinesis 8 |
| chr19 | 25236492 | 25237876 | ENSMUSG00000032702 | Kank1 | KN motif and ankyrin repeat domains 1 |
| chr19 | 26605845 | 26607770 | ENSMUSG00000024921 | Smarca2 | SWI/SNF related, matrix associated, actin dependent regulator of chromatin, subfamily a, member 2 |
| chr19 | 28677950 | 28679514 | ENSMUSG00000052942 | Glis3 | GLIS family zinc finger 3 |
| chr19 | 29251065 | 29252226 | ENSMUSG00000024789 | Jak2 | Janus kinase 2 |
| chr19 | 29521837 | 29523605 | ENSMUSG00000038658 | Ric1 | RAB6A GEF complex partner 1 |
| chr19 | 29647726 | 29648475 | ENSMUSG00000046324 | Ermp1 | endoplasmic reticulum metallopeptidase 1 |
| chr19 | 29805068 | 29806473 | ENSMUSG00000046138 | 9930021J03Rik | RIKEN cDNA 9930021J03 gene |
| chr19 | 32387697 | 32389056 | ENSMUSG00000040451 | Sgms1 | sphingomyelin synthase 1 |
| chr19 | 32389056 | 32389761 | ENSMUSG00000040451 | Sgms1 | sphingomyelin synthase 1 |
| chr19 | 32485620 | 32486684 | ENSMUSG00000024896 | Minpp1 | multiple inositol polyphosphate histidine phosphatase 1 |
| chr19 | 32711728 | 32712636 | ENSMUSG00000013662 | Atad1 | ATPase family, AAA domain containing 1 |
| chr19 | 32756989 | 32758385 | ENSMUSG00000013663 | Pten | phosphatase and tensin homolog |
| chr19 | 34192311 | 34192968 | ENSMUSG00000024776 | Stambpl1 | STAM binding protein like 1 |
| chr19 | 34878095 | 34879331 | ENSMUSG00000033610 | Pank1 | pantothenate kinase 1 |
| chr19 | 36347997 | 36349652 | ENSMUSG00000024805 | Pcgf5 | polycomb group ring finger 5 |
| chr19 | 36409409 | 36410377 | ENSMUSG00000024805 | Pcgf5 | polycomb group ring finger 5 |
| chr19 | 36834032 | 36834996 | ENSMUSG00000024811 | Tnks2 | tankyrase, TRF1-interacting ankyrin-related ADP-ribose polymerase 2 |
| chr19 | 36918358 | 36919345 | ENSMUSG00000047632 | Fgfbp3 | fibroblast growth factor binding protein 3 |
| chr19 | 36925721 | 36926753 | ENSMUSG00000040565 | Btaf1 | B-TFIID TATA-box binding protein associated factor 1 |
| chr19 | 37174165 | 37174886 | ENSMUSG00000039652 | Cpeb3 | cytoplasmic polyadenylation element binding protein 3 |
| chr19 | 37206951 | 37208052 | ENSMUSG00000039652 | Cpeb3 | cytoplasmic polyadenylation element binding protein 3 |
| chr19 | 37208052 | 37208525 | ENSMUSG00000039652 | Cpeb3 | cytoplasmic polyadenylation element binding protein 3 |
| chr19 | 37550328 | 37551080 | ENSMUSG00000053799 | Exoc6 | exocyst complex component 6 |
| chr19 | 38395694 | 38396370 | ENSMUSG00000044026 | Slc35g1 | solute carrier family 35, member G1 |
| chr19 | 38836428 | 38837573 | ENSMUSG00000048720 | Tbc1d12 | TBC1D12: TBC1 domain family, member 12 |
| chr19 | 38930724 | 38931685 | ENSMUSG00000025001 | Hells | helicase, lymphoid specific |
| chr19 | 40513484 | 40514083 | ENSMUSG00000025006 | Sorbs1 | sorbin and SH3 domain containing 1 |
| chr19 | 40831028 | 40832166 | ENSMUSG00000025010 | Ccnj | cyclin J |
| chr19 | 41263039 | 41264387 | ENSMUSG00000025016 | Tm9sf3 | transmembrane 9 superfamily member 3 |
| chr19 | 41482119 | 41483906 | ENSMUSG00000025019 | Lcor | ligand dependent nuclear receptor corepressor |
| chr19 | 41829765 | 41830700 | ENSMUSG00000067199 | Frat1 | frequently rearranged in advanced T cell lymphomas |
| chr19 | 41847067 | 41848547 | ENSMUSG00000047604 | Frat2 | frequently rearranged in advanced T cell lymphomas 2 |
| chr19 | 41911684 | 41912671 | ENSMUSG00000011752 | Pgam1 | phosphoglycerate mutase 1 |
| chr19 | 41980612 | 41982424 | ENSMUSG00000025171 | Ubtd1 | ubiquitin domain containing 1 |
| chr19 | 42090217 | 42090963 | ENSMUSG00000025178 | Pi4k2a | phosphatidylinositol 4-kinase type 2 alpha |
| chr19 | 42147217 | 42148643 | ENSMUSG00000044345 | Marveld1 | MARVEL (membrane-associating) domain containing 1 |
| chr19 | 43674316 | 43675989 | ENSMUSG00000117698 | BC037704 | cDNA sequence BC037704 |
| chr19 | 43689512 | 43690182 | ENSMUSG00000025192 | Entpd7 | ectonucleoside triphosphate diphosphohydrolase 7 |
| chr19 | 43752762 | 43753441 | ENSMUSG00000040018 | Cox15 | cytochrome c oxidase assembly protein 15 |
| chr19 | 43939751 | 43940315 | ENSMUSG00000025195 | Dnmbp | dynamin binding protein |
| chr19 | 44069324 | 44069961 | ENSMUSG00000025198 | Erlin1 | ER lipid raft associated 1 |
| chr19 | 44720114 | 44720602 | ENSMUSG00000004231 | Pax2 | paired box 2 |
| chr19 | 44746533 | 44747021 | ENSMUSG00000004231 | Pax2 | paired box 2 |
| chr19 | 45659399 | 45660719 | ENSMUSG00000040913 | Fbxw4 | F-box and WD-40 domain protein 4 |
| chr19 | 46003406 | 46003943 | ENSMUSG00000074811 | Hps6 | HPS6, biogenesis of lysosomal organelles complex 2 subunit 3 |
| chr19 | 46044750 | 46045735 | ENSMUSG00000025223 | Ldb1 | LIM domain binding 1 |
| chr19 | 46056506 | 46057466 | ENSMUSG00000055491 | Pprc1 | peroxisome proliferative activated receptor, gamma, coactivator-related 1 |
| chr19 | 46135862 | 46136783 | ENSMUSG00000025229 | Pitx3 | paired-like homeodomain transcription factor 3 |
| chr19 | 46136783 | 46137143 | ENSMUSG00000025229 | Pitx3 | paired-like homeodomain transcription factor 3 |
| chr19 | 46304291 | 46305791 | ENSMUSG00000118087 | 4833438C02Rik | RIKEN cDNA 4833438C02 gene |
| chr19 | 46328159 | 46329652 | ENSMUSG00000025226 | Fbxl15 | F-box and leucine-rich repeat protein 15 |
| chr19 | 46329652 | 46330383 | ENSMUSG00000025226 | Fbxl15 | F-box and leucine-rich repeat protein 15 |
| chr19 | 46348512 | 46349507 | ENSMUSG00000070127 | Mir146b | microRNA 146b |
| chr19 | 46573104 | 46573729 | ENSMUSG00000025035 | Arl3 | ADP-ribosylation factor-like 3 |
| chr19 | 46961503 | 46962608 | ENSMUSG00000025041 | Nt5c2 | 5'-nucleotidase, cytosolic II |
| chr19 | 47050249 | 47050868 | ENSMUSG00000025050 | Pcgf6 | polycomb group ring finger 6 |
| chr19 | 47090139 | 47091008 | ENSMUSG00000025047 | Pdcd11 | programmed cell death 11 |
| chr19 | 47578871 | 47580674 | ENSMUSG00000025060 | Slk | STE20-like kinase |
| chr19 | 47731084 | 47732014 | ENSMUSG00000025066 | Sfr1 | SWI5 dependent recombination repair 1 |
| chr19 | 53038176 | 53038720 | ENSMUSG00000025027 | Xpnpep1 | X-prolyl aminopeptidase (aminopeptidase P) 1, soluble |
| chr19 | 53312657 | 53314352 | ENSMUSG00000025025 | Mxi1 | MAX interactor 1, dimerization protein |
| chr19 | 53329342 | 53330476 | ENSMUSG00000025025 | Mxi1 | MAX interactor 1, dimerization protein |
| chr19 | 53528678 | 53530296 | ENSMUSG00000034765 | Dusp5 | dual specificity phosphatase 5 |
| chr19 | 53600027 | 53601386 | ENSMUSG00000024974 | Smc3 | structural maintenance of chromosomes 3 |
| chr19 | 53902886 | 53904157 | ENSMUSG00000024975 | Pdcd4 | programmed cell death 4 |
| chr19 | 53943789 | 53944708 | ENSMUSG00000084957 | Bbip1 | BBSome interacting protein 1 |
| chr19 | 53944708 | 53945578 | ENSMUSG00000024976 | Shoc2 | Shoc2, leucine rich repeat scaffold protein |
| chr19 | 55098743 | 55099739 | ENSMUSG00000024978 | Gpam | glycerol-3-phosphate acyltransferase, mitochondrial |
| chr19 | 55253586 | 55254429 | ENSMUSG00000024981 | Acsl5 | acyl-CoA synthetase long-chain family member 5 |
| chr19 | 55741427 | 55741790 | ENSMUSG00000024985 | Tcf7l2 | transcription factor 7 like 2, T cell specific, HMG box |
| chr19 | 55742412 | 55743977 | ENSMUSG00000024985 | Tcf7l2 | transcription factor 7 like 2, T cell specific, HMG box |
| chr19 | 56547826 | 56549124 | ENSMUSG00000025077 | Dclre1a | DNA cross-link repair 1A |
| chr19 | 57314401 | 57315091 | ENSMUSG00000025085 | Ablim1 | actin-binding LIM protein 1 |
| chr19 | 57360570 | 57361790 | ENSMUSG00000033478 | Fam160b1 | family with sequence similarity 160, member B1 |
| chr19 | 59074891 | 59076052 | ENSMUSG00000041362 | Shtn1 | shootin 1 |
| chr19 | 59459122 | 59460061 | ENSMUSG00000043969 | Emx2 | empty spiracles homeobox 2 |
| chr19 | 59943156 | 59943949 | ENSMUSG00000040022 | Rab11fip2 | RAB11 family interacting protein 2 (class I) |
| chr19 | 60755628 | 60757267 | ENSMUSG00000072437 | Nanos1 | nanos C2HC-type zinc finger 1 |
| chr19 | 60789924 | 60790921 | ENSMUSG00000024991 | Eif3a | eukaryotic translation initiation factor 3, subunit A |
| chr19 | 60874099 | 60874585 | ENSMUSG00000024997 | Prdx3 | peroxiredoxin 3 |
| chr19 | 60889667 | 60890827 | ENSMUSG00000003228 | Grk5 | G protein-coupled receptor kinase 5 |
| chr2 | 3118279 | 3118942 | ENSMUSG00000050530 | Fam171a1 | family with sequence similarity 171, member A1 |
| chr2 | 3283659 | 3285154 | ENSMUSG00000026643 | Nmt2 | N-myristoyltransferase 2 |
| chr2 | 4881478 | 4882729 | ENSMUSG00000026662 | Sephs1 | selenophosphate synthetase 1 |
| chr2 | 6129619 | 6130338 | ENSMUSG00000045319 | Proser2 | proline and serine rich 2 |
| chr2 | 6322467 | 6323542 | ENSMUSG00000039046 | Usp6nl | USP6 N-terminal like |
| chr2 | 6871867 | 6872935 | ENSMUSG00000002107 | Celf2 | CUGBP, Elav-like family member 2 |
| chr2 | 11501387 | 11502324 | ENSMUSG00000026773 | Pfkfb3 | 6-phosphofructo-2-kinase/fructose-2,6-biphosphatase 3 |
| chr2 | 18047292 | 18048228 | ENSMUSG00000026743 | Mllt10 | myeloid/lymphoid or mixed-lineage leukemia; translocated to, 10 |
| chr2 | 18056334 | 18057934 | ENSMUSG00000099050 | Mir7655 | microRNA 7655 |
| chr2 | 18064164 | 18065830 | ENSMUSG00000026743 | Mllt10 | myeloid/lymphoid or mixed-lineage leukemia; translocated to, 10 |
| chr2 | 18392102 | 18392938 | ENSMUSG00000026740 | Dnajc1 | DnaJ heat shock protein family (Hsp40) member C1 |
| chr2 | 18671967 | 18672808 | ENSMUSG00000051154 | Commd3 | COMM domain containing 3 |
| chr2 | 18676172 | 18677892 | ENSMUSG00000026739 | Bmi1 | Bmi1 polycomb ring finger oncogene |
| chr2 | 18693964 | 18694747 | ENSMUSG00000037708 | Spag6 | sperm associated antigen 6 |
| chr2 | 18997414 | 18998418 | ENSMUSG00000026737 | Pip4k2a | phosphatidylinositol-5-phosphate 4-kinase, type II, alpha |
| chr2 | 19657408 | 19659260 | ENSMUSG00000043415 | Otud1 | OTU domain containing 1 |
| chr2 | 20967782 | 20969757 |  | Gm13375 | predicted gene 13375 |
| chr2 | 22895443 | 22895994 | ENSMUSG00000026784 | Pdss1 | prenyl (solanesyl) diphosphate synthase, subunit 1 |
| chr2 | 24918835 | 24919857 | ENSMUSG00000036893 | Ehmt1 | euchromatic histone methyltransferase 1 |
| chr2 | 24949513 | 24950436 | ENSMUSG00000026974 | Zmynd19 | zinc finger, MYND domain containing 19 |
| chr2 | 25180675 | 25182080 | ENSMUSG00000078202 | Nrarp | Notch-regulated ankyrin repeat protein |
| chr2 | 25262349 | 25263924 | ENSMUSG00000048707 | Tprn | taperin |
| chr2 | 25365185 | 25365714 | ENSMUSG00000026956 | Uap1l1 | UDP-N-acteylglucosamine pyrophosphorylase 1-like 1 |
| chr2 | 25372200 | 25373103 | ENSMUSG00000026955 | Sapcd2 | suppressor APC domain containing 2 |
| chr2 | 25546487 | 25547103 | ENSMUSG00000026942 | Traf2 | TNF receptor-associated factor 2 |
| chr2 | 25557354 | 25558210 | ENSMUSG00000015092 | Edf1 | endothelial differentiation-related factor 1 |
| chr2 | 25982117 | 25983827 | ENSMUSG00000026933 | Camsap1 | calmodulin regulated spectrin-associated protein 1 |
| chr2 | 26122344 | 26122988 | ENSMUSG00000026932 | Nacc2 | nucleus accumbens associated 2, BEN and BTB (POZ) domain containing |
| chr2 | 26139701 | 26140687 | ENSMUSG00000087679 | Tmem250-ps | transmembrane protein 250, pseudogene |
| chr2 | 26315386 | 26316467 | ENSMUSG00000026930 | Gpsm1 | G-protein signalling modulator 1 (AGS3-like, C. elegans) |
| chr2 | 26502199 | 26503986 | ENSMUSG00000026923 | Notch1 | notch 1 |
| chr2 | 26603691 | 26604680 | ENSMUSG00000026922 | Agpat2 | 1-acylglycerol-3-phosphate O-acyltransferase 2 (lysophosphatidic acid acyltransferase, beta) |
| chr2 | 26628329 | 26628768 | ENSMUSG00000036186 | Dipk1b | divergent protein kinase domain 1B |
| chr2 | 26932858 | 26933873 | ENSMUSG00000014867 | Surf4 | surfeit gene 4 |
| chr2 | 27425346 | 27427203 | ENSMUSG00000009621 | Vav2 | vav 2 oncogene |
| chr2 | 27474901 | 27476147 | ENSMUSG00000026918 | Brd3 | bromodomain containing 3 |
| chr2 | 27514996 | 27515725 | ENSMUSG00000026917 | Wdr5 | WD repeat domain 5 |
| chr2 | 29124954 | 29125513 | ENSMUSG00000043535 | Setx | senataxin |
| chr2 | 29965174 | 29966216 | ENSMUSG00000057738 | Sptan1 | spectrin alpha, non-erythrocytic 1 |
| chr2 | 30065974 | 30067471 | ENSMUSG00000054766 | Set | SET nuclear oncogene |
| chr2 | 30237502 | 30238283 | ENSMUSG00000007476 | Lrrc8a | leucine rich repeat containing 8A VRAC subunit A |
| chr2 | 30358839 | 30359438 | ENSMUSG00000026860 | Sh3glb2 | SH3-domain GRB2-like endophilin B2 |
| chr2 | 30707745 | 30708459 | ENSMUSG00000093091 | Mir3089 | microRNA 3089 |
| chr2 | 30711603 | 30712390 | ENSMUSG00000093091 | Mir3089 | microRNA 3089 |
| chr2 | 31950102 | 31950892 | ENSMUSG00000001864 | Aif1l | allograft inflammatory factor 1-like |
| chr2 | 32381578 | 32381998 | ENSMUSG00000039195 | 1110008P14Rik | RIKEN cDNA 1110008P14 gene |
| chr2 | 32450881 | 32451608 | ENSMUSG00000026819 | Slc25a25 | solute carrier family 25 (mitochondrial carrier, phosphate carrier), member 25 |
| chr2 | 32535236 | 32536023 | ENSMUSG00000039157 | Fam102a | family with sequence similarity 102, member A |
| chr2 | 32693771 | 32694292 | ENSMUSG00000009566 | Fpgs | folylpolyglutamyl synthetase |
| chr2 | 34753790 | 34754670 | ENSMUSG00000026867 | Gapvd1 | GTPase activating protein and VPS9 domains 1 |
| chr2 | 35109363 | 35109974 | ENSMUSG00000057110 | Cntrl | centriolin |
| chr2 | 35462186 | 35462686 | ENSMUSG00000035778 | Ggta1 | glycoprotein galactosyltransferase alpha 1, 3 |
| chr2 | 35581454 | 35582925 | ENSMUSG00000026883 | Dab2ip | disabled 2 interacting protein |
| chr2 | 35661451 | 35662384 | ENSMUSG00000026883 | Dab2ip | disabled 2 interacting protein |
| chr2 | 35979139 | 35979968 | ENSMUSG00000026885 | Ttll11 | tubulin tyrosine ligase-like family, member 11 |
| chr2 | 37702588 | 37703919 | ENSMUSG00000026915 | Strbp | spermatid perinuclear RNA binding protein |
| chr2 | 38286904 | 38287565 | ENSMUSG00000035392 | Dennd1a | DENN/MADD domain containing 1A |
| chr2 | 38511710 | 38512198 | ENSMUSG00000026749 | Nek6 | NIMA (never in mitosis gene a)-related expressed kinase 6 |
| chr2 | 44926684 | 44927715 | ENSMUSG00000036890 | Gtdc1 | glycosyltransferase-like domain containing 1 |
| chr2 | 48813496 | 48814909 | ENSMUSG00000052155 | Acvr2a | activin receptor IIA |
| chr2 | 52072612 | 52073596 | ENSMUSG00000036202 | Rif1 | replication timing regulatory factor 1 |
| chr2 | 52424293 | 52425079 | ENSMUSG00000036093 | Arl5a | ADP-ribosylation factor-like 5A |
| chr2 | 52857556 | 52859018 | ENSMUSG00000036053 | Fmnl2 | formin-like 2 |
| chr2 | 57123428 | 57124363 | ENSMUSG00000026826 | Nr4a2 | nuclear receptor subfamily 4, group A, member 2 |
| chr2 | 57238019 | 57238766 | ENSMUSG00000026827 | Gpd2 | glycerol phosphate dehydrogenase 2, mitochondrial |
| chr2 | 58566539 | 58567691 | ENSMUSG00000026836 | Acvr1 | activin A receptor, type 1 |
| chr2 | 59160645 | 59162108 | ENSMUSG00000036641 | Ccdc148 | coiled-coil domain containing 148 |
| chr2 | 59611717 | 59613236 | ENSMUSG00000035168 | Tanc1 | tetratricopeptide repeat, ankyrin repeat and coiled-coil containing 1 |
| chr2 | 60209343 | 60210472 | ENSMUSG00000026987 | Baz2b | bromodomain adjacent to zinc finger domain, 2B |
| chr2 | 60880823 | 60881817 | ENSMUSG00000026970 | Rbms1 | RNA binding motif, single stranded interacting protein 1 |
| chr2 | 65237731 | 65239381 | ENSMUSG00000034903 | Cobll1 | Cobl-like 1 |
| chr2 | 66123894 | 66125048 | ENSMUSG00000026994 | Galnt3 | polypeptide N-acetylgalactosaminyltransferase 3 |
| chr2 | 68470769 | 68472479 | ENSMUSG00000027030 | Stk39 | serine/threonine kinase 39 |
| chr2 | 68861234 | 68862377 | ENSMUSG00000027035 | Cers6 | ceramide synthase 6 |
| chr2 | 69822752 | 69823394 | ENSMUSG00000042155 | Klhl23 | kelch-like 23 |
| chr2 | 70661364 | 70662138 | ENSMUSG00000014959 | Gorasp2 | golgi reassembly stacking protein 2 |
| chr2 | 70824855 | 70826073 | ENSMUSG00000041997 | Tlk1 | tousled-like kinase 1 |
| chr2 | 71528951 | 71529821 | ENSMUSG00000041911 | Dlx1 | distal-less homeobox 1 |
| chr2 | 71786730 | 71787672 | ENSMUSG00000027111 | Itga6 | integrin alpha 6 |
| chr2 | 71872991 | 71873852 | ENSMUSG00000006494 | Pdk1 | pyruvate dehydrogenase kinase, isoenzyme 1 |
| chr2 | 72285447 | 72286350 | ENSMUSG00000004085 | Map3k20 | mitogen-activated protein kinase kinase kinase 20 |
| chr2 | 72476122 | 72476963 | ENSMUSG00000055612 | Cdca7 | cell division cycle associated 7 |
| chr2 | 74697743 | 74698800 | ENSMUSG00000043342 | Hoxd9 | homeobox D9 |
| chr2 | 74704614 | 74706090 | ENSMUSG00000027102 | Hoxd8 | homeobox D8 |
| chr2 | 74711895 | 74712476 | ENSMUSG00000079277 | Hoxd3 | homeobox D3 |
| chr2 | 75703421 | 75705394 | ENSMUSG00000015839 | Nfe2l2 | nuclear factor, erythroid derived 2, like 2 |
| chr2 | 75831975 | 75832769 | ENSMUSG00000042410 | Agps | alkylglycerone phosphate synthase |
| chr2 | 76674983 | 76675762 | ENSMUSG00000002733 | Plekha3 | pleckstrin homology domain-containing, family A (phosphoinositide binding specific) member 3 |
| chr2 | 77280104 | 77280983 | ENSMUSG00000042272 | Sestd1 | SEC14 and spectrin domains 1 |
| chr2 | 78868610 | 78869733 | ENSMUSG00000027011 | Ube2e3 | ubiquitin-conjugating enzyme E2E 3 |
| chr2 | 79635115 | 79636143 | ENSMUSG00000027007 | Itprid2 | ITPR interacting domain containing 2 |
| chr2 | 80580437 | 80581757 | ENSMUSG00000027002 | Nckap1 | NCK-associated protein 1 |
| chr2 | 83724092 | 83725173 | ENSMUSG00000027087 | Itgav | integrin alpha V |
| chr2 | 84714454 | 84715541 | ENSMUSG00000034075 | Zdhhc5 | zinc finger, DHHC domain containing 5 |
| chr2 | 90579468 | 90581191 | ENSMUSG00000025314 | Ptprj | protein tyrosine phosphatase, receptor type, J |
| chr2 | 90846949 | 90847611 | ENSMUSG00000027282 | Mtch2 | mitochondrial carrier 2 |
| chr2 | 90917178 | 90918068 | ENSMUSG00000063235 | Ptpmt1 | protein tyrosine phosphatase, mitochondrial 1 |
| chr2 | 90940259 | 90941256 | ENSMUSG00000005506 | Celf1 | CUGBP, Elav-like family member 1 |
| chr2 | 91963118 | 91963911 | ENSMUSG00000040479 | Dgkz | diacylglycerol kinase zeta |
| chr2 | 92433611 | 92434374 | ENSMUSG00000068742 | Cry2 | cryptochrome 2 (photolyase-like) |
| chr2 | 93822136 | 93822675 | ENSMUSG00000027198 | Ext2 | exostosin glycosyltransferase 2 |
| chr2 | 102400424 | 102401330 | ENSMUSG00000027189 | Trim44 | tripartite motif-containing 44 |
| chr2 | 103566333 | 103567439 | ENSMUSG00000032724 | Abtb2 | ankyrin repeat and BTB (POZ) domain containing 2 |
| chr2 | 103796219 | 103797761 | ENSMUSG00000027184 | Caprin1 | cell cycle associated protein 1 |
| chr2 | 104493015 | 104494776 | ENSMUSG00000027177 | Hipk3 | homeodomain interacting protein kinase 3 |
| chr2 | 104742427 | 104743063 | ENSMUSG00000027173 | Depdc7 | DEP domain containing 7 |
| chr2 | 104816329 | 104817360 | ENSMUSG00000074994 | Qser1 | glutamine and serine rich 1 |
| chr2 | 104849218 | 104849932 | ENSMUSG00000027171 | Prrg4 | proline rich Gla (G-carboxyglutamic acid) 4 (transmembrane) |
| chr2 | 109917578 | 109918336 | ENSMUSG00000050199 | Lgr4 | leucine-rich repeat-containing G protein-coupled receptor 4 |
| chr2 | 112239182 | 112240417 | ENSMUSG00000027134 | Lpcat4 | lysophosphatidylcholine acyltransferase 4 |
| chr2 | 112265412 | 112266134 | ENSMUSG00000027130 | Slc12a6 | solute carrier family 12, member 6 |
| chr2 | 112492789 | 112493525 | ENSMUSG00000003604 | Aven | apoptosis, caspase activation inhibitor |
| chr2 | 117120364 | 117121945 | ENSMUSG00000027351 | Spred1 | sprouty protein with EVH-1 domain 1, related sequence |
| chr2 | 118544318 | 118544644 | ENSMUSG00000040093 | Bmf | BCL2 modifying factor |
| chr2 | 118901174 | 118902888 | ENSMUSG00000040007 | Bahd1 | bromo adjacent homology domain containing 1 |
| chr2 | 119236864 | 119238416 | ENSMUSG00000027315 | Spint1 | serine protease inhibitor, Kunitz type 1 |
| chr2 | 119757848 | 119758549 | ENSMUSG00000027297 | Ltk | leukocyte tyrosine kinase |
| chr2 | 119758549 | 119759020 | ENSMUSG00000027297 | Ltk | leukocyte tyrosine kinase |
| chr2 | 119799238 | 119800099 | ENSMUSG00000027298 | Tyro3 | TYRO3 protein tyrosine kinase 3 |
| chr2 | 120154152 | 120154790 | ENSMUSG00000027293 | Ehd4 | EH-domain containing 4 |
| chr2 | 120730861 | 120731662 | ENSMUSG00000027284 | Cdan1 | congenital dyserythropoietic anemia, type I (human) |
| chr2 | 120976997 | 120977465 | ENSMUSG00000054484 | Tmem62 | transmembrane protein 62 |
| chr2 | 121170756 | 121171701 | ENSMUSG00000027263 | Tubgcp4 | tubulin, gamma complex associated protein 4 |
| chr2 | 121413455 | 121414670 | ENSMUSG00000027248 | Pdia3 | protein disulfide isomerase associated 3 |
| chr2 | 121448973 | 121450073 | ENSMUSG00000074884 | Serf2 | small EDRK-rich factor 2 |
| chr2 | 122028238 | 122029214 | ENSMUSG00000043424 | Eif3j2 | eukaryotic translation initiation factor 3, subunit J2 |
| chr2 | 122348529 | 122349048 | ENSMUSG00000033256 | Shf | Src homology 2 domain containing F |
| chr2 | 122630508 | 122631800 |  | Spata5l1 | spermatogenesis associated 5-like 1 |
| chr2 | 122702102 | 122702882 | ENSMUSG00000005802 | Slc30a4 | solute carrier family 30 (zinc transporter), member 4 |
| chr2 | 125247130 | 125248143 | ENSMUSG00000027203 | Dut | deoxyuridine triphosphatase |
| chr2 | 125672931 | 125673587 | ENSMUSG00000035109 | Shc4 | SHC (Src homology 2 domain containing) family, member 4 |
| chr2 | 126875564 | 126876511 | ENSMUSG00000027365 | Trpm7 | transient receptor potential cation channel, subfamily M, member 7 |
| chr2 | 127269652 | 127270893 | ENSMUSG00000027367 | Stard7 | START domain containing 7 |
| chr2 | 127336023 | 127336653 | ENSMUSG00000027368 | Dusp2 | dual specificity phosphatase 2 |
| chr2 | 128125588 | 128127010 | ENSMUSG00000027381 | Bcl2l11 | BCL2-like 11 (apoptosis facilitator) |
| chr2 | 128127010 | 128128342 | ENSMUSG00000027381 | Bcl2l11 | BCL2-like 11 (apoptosis facilitator) |
| chr2 | 129065812 | 129066253 | ENSMUSG00000027394 | Ttl | tubulin tyrosine ligase |
| chr2 | 130563549 | 130564318 | ENSMUSG00000060029 | 4930473A02Rik | RIKEN cDNA 4930473A02 gene |
| chr2 | 130839475 | 130840528 | ENSMUSG00000027309 | 4930402H24Rik | RIKEN cDNA 4930402H24 gene |
| chr2 | 130906367 | 130907071 | ENSMUSG00000027312 | Atrn | attractin |
| chr2 | 131178994 | 131180354 | ENSMUSG00000068267 | Cenpb | centromere protein B |
| chr2 | 131262283 | 131262961 | ENSMUSG00000037514 | Pank2 | pantothenate kinase 2 |
| chr2 | 131491683 | 131492569 | ENSMUSG00000027333 | Smox | spermine oxidase |
| chr2 | 132262727 | 132264050 | ENSMUSG00000058793 | Cds2 | CDP-diacylglycerol synthase (phosphatidate cytidylyltransferase) 2 |
| chr2 | 132577650 | 132578398 | ENSMUSG00000027346 | Gpcpd1 | glycerophosphocholine phosphodiesterase 1 |
| chr2 | 132815744 | 132816417 | ENSMUSG00000027353 | Mcm8 | minichromosome maintenance 8 homologous recombination repair factor |
| chr2 | 132846580 | 132847315 | ENSMUSG00000027357 | Crls1 | cardiolipin synthase 1 |
| chr2 | 134643525 | 134644235 | ENSMUSG00000034723 | Tmx4 | thioredoxin-related transmembrane protein 4 |
| chr2 | 137116067 | 137117539 | ENSMUSG00000027276 | Jag1 | jagged 1 |
| chr2 | 138256468 | 138257269 | ENSMUSG00000062098 | Btbd3 | BTB (POZ) domain containing 3 |
| chr2 | 140395058 | 140396159 | ENSMUSG00000068205 | Macrod2 | mono-ADP ribosylhydrolase 2 |
| chr2 | 142901111 | 142901602 | ENSMUSG00000038844 | Kif16b | kinesin family member 16B |
| chr2 | 143914808 | 143915937 | ENSMUSG00000015932 | Dstn | destrin |
| chr2 | 144009881 | 144011388 | ENSMUSG00000027422 | Rrbp1 | ribosome binding protein 1 |
| chr2 | 144270146 | 144271234 | ENSMUSG00000027423 | Snx5 | sorting nexin 5 |
| chr2 | 144368784 | 144369333 | ENSMUSG00000027425 | Kat14 | lysine acetyltransferase 14 |
| chr2 | 145903138 | 145903821 | ENSMUSG00000002728 | Naa20 | N(alpha)-acetyltransferase 20, NatB catalytic subunit |
| chr2 | 146511648 | 146512315 | ENSMUSG00000037110 | Ralgapa2 | Ral GTPase activating protein, alpha subunit 2 (catalytic) |
| chr2 | 150904174 | 150904879 | ENSMUSG00000032046 | Abhd12 | abhydrolase domain containing 12 |
| chr2 | 151542312 | 151543197 | ENSMUSG00000032966 | Fkbp1a | FK506 binding protein 1a |
| chr2 | 152332071 | 152332707 | ENSMUSG00000027466 | Rbck1 | RanBP-type and C3HC4-type zinc finger containing 1 |
| chr2 | 152414031 | 152415125 | ENSMUSG00000074682 | Zcchc3 | zinc finger, CCHC domain containing 3 |
| chr2 | 153241091 | 153241850 | ENSMUSG00000046020 | Pofut1 | protein O-fucosyltransferase 1 |
| chr2 | 153345530 | 153346889 | ENSMUSG00000042548 | Asxl1 | additional sex combs like 1 |
| chr2 | 153632392 | 153632885 | ENSMUSG00000056941 | Commd7 | COMM domain containing 7 |
| chr2 | 154407475 | 154408217 | ENSMUSG00000027488 | Snta1 | syntrophin, acidic 1 |
| chr2 | 154436032 | 154437187 | ENSMUSG00000038533 | Cbfa2t2 | CBFA2/RUNX1 translocation partner 2 |
| chr2 | 154568733 | 154570310 | ENSMUSG00000027490 | E2f1 | E2F transcription factor 1 |
| chr2 | 154613262 | 154613960 | ENSMUSG00000059842 | Zfp341 | zinc finger protein 341 |
| chr2 | 154656825 | 154657523 | ENSMUSG00000038467 | Chmp4b | charged multivesicular body protein 4B |
| chr2 | 154790526 | 154792385 | ENSMUSG00000027593 | Raly | hnRNP-associated with lethal yellow |
| chr2 | 155133451 | 155134279 | ENSMUSG00000027598 | Itch | itchy, E3 ubiquitin protein ligase |
| chr2 | 155276263 | 155277057 | ENSMUSG00000027602 | Map1lc3a | microtubule-associated protein 1 light chain 3 alpha |
| chr2 | 155473104 | 155474008 | ENSMUSG00000038369 | Ncoa6 | nuclear receptor coactivator 6 |
| chr2 | 155517928 | 155518441 | ENSMUSG00000027603 | Ggt7 | gamma-glutamyltransferase 7 |
| chr2 | 155592276 | 155592883 | ENSMUSG00000027610 | Gss | glutathione synthetase |
| chr2 | 155691831 | 155692525 | ENSMUSG00000038324 | Trpc4ap | transient receptor potential cation channel, subfamily C, member 4 associated protein |
| chr2 | 155818767 | 155819478 | ENSMUSG00000074649 | BC029722 | cDNA sequence BC029722 |
| chr2 | 155819478 | 155820206 | ENSMUSG00000074649 | BC029722 | cDNA sequence BC029722 |
| chr2 | 156065284 | 156066070 | ENSMUSG00000038180 | Spag4 | sperm associated antigen 4 |
| chr2 | 156144031 | 156144728 | ENSMUSG00000027618 | Nfs1 | nitrogen fixation gene 1 (S. cerevisiae) |
| chr2 | 156196494 | 156197351 | ENSMUSG00000038116 | Phf20 | PHD finger protein 20 |
| chr2 | 156311859 | 156312837 | ENSMUSG00000046229 | Scand1 | SCAN domain-containing 1 |
| chr2 | 156420547 | 156421923 | ENSMUSG00000027624 | Epb41l1 | erythrocyte membrane protein band 4.1 like 1 |
| chr2 | 156474988 | 156476037 | ENSMUSG00000027624 | Epb41l1 | erythrocyte membrane protein band 4.1 like 1 |
| chr2 | 156720617 | 156721948 | ENSMUSG00000061689 | Dlgap4 | DLG associated protein 4 |
| chr2 | 156862643 | 156863673 | ENSMUSG00000027637 | Rab5if | RAB5 interacting factor |
| chr2 | 156991670 | 156992213 | ENSMUSG00000027634 | Ndrg3 | N-myc downstream regulated gene 3 |
| chr2 | 157006596 | 157007202 | ENSMUSG00000027635 | Dsn1 | DSN1 homolog, MIS12 kinetochore complex component |
| chr2 | 157077628 | 157079627 | ENSMUSG00000055485 | Soga1 | suppressor of glucose, autophagy associated 1 |
| chr2 | 157134499 | 157135551 | ENSMUSG00000027639 | Samhd1 | SAM domain and HD domain, 1 |
| chr2 | 157203958 | 157204945 | ENSMUSG00000027641 | Rbl1 | RB transcriptional corepressor like 1 |
| chr2 | 157278836 | 157279779 | ENSMUSG00000027642 | Rpn2 | ribophorin II |
| chr2 | 157422865 | 157424703 | ENSMUSG00000027646 | Src | Rous sarcoma oncogene |
| chr2 | 158409824 | 158410654 | ENSMUSG00000027652 | Ralgapb | Ral GTPase activating protein, beta subunit (non-catalytic) |
| chr2 | 158624963 | 158625645 | ENSMUSG00000037761 | Actr5 | ARP5 actin-related protein 5 |
| chr2 | 158767959 | 158768724 | ENSMUSG00000027654 | Fam83d | family with sequence similarity 83, member D |
| chr2 | 160645549 | 160646930 | ENSMUSG00000070544 | Top1 | topoisomerase (DNA) I |
| chr2 | 160730903 | 160732260 | ENSMUSG00000016933 | Plcg1 | phospholipase C, gamma 1 |
| chr2 | 160872380 | 160873204 | ENSMUSG00000035877 | Zhx3 | zinc fingers and homeoboxes 3 |
| chr2 | 161108277 | 161109653 | ENSMUSG00000057133 | Chd6 | chromodomain helicase DNA binding protein 6 |
| chr2 | 162930752 | 162932456 | ENSMUSG00000016921 | Srsf6 | serine and arginine-rich splicing factor 6 |
| chr2 | 163054417 | 163055286 | ENSMUSG00000017861 | Mybl2 | myeloblastosis oncogene-like 2 |
| chr2 | 163418825 | 163419941 | ENSMUSG00000099632 | 2900093K20Rik | RIKEN cDNA 2900093K20 gene |
| chr2 | 163472180 | 163472743 | ENSMUSG00000048486 | Fitm2 | fat storage-inducing transmembrane protein 2 |
| chr2 | 163602161 | 163603078 | ENSMUSG00000017679 | Ttpal | tocopherol (alpha) transfer protein-like |
| chr2 | 163644554 | 163645318 | ENSMUSG00000017707 | Serinc3 | serine incorporator 3 |
| chr2 | 163658213 | 163658675 | ENSMUSG00000035268 | Pkig | protein kinase inhibitor, gamma |
| chr2 | 163994833 | 163995765 | ENSMUSG00000018326 | Ywhab | tyrosine 3-monooxygenase/tryptophan 5-monooxygenase activation protein, beta polypeptide |
| chr2 | 164432618 | 164433350 | ENSMUSG00000017009 | Sdc4 | syndecan 4 |
| chr2 | 164442599 | 164443466 | ENSMUSG00000017009 | Sdc4 | syndecan 4 |
| chr2 | 164485965 | 164487327 | ENSMUSG00000017734 | Dbndd2 | dysbindin (dystrobrevin binding protein 1) domain containing 2 |
| chr2 | 164497252 | 164497780 | ENSMUSG00000017721 | Pigt | phosphatidylinositol glycan anchor biosynthesis, class T |
| chr2 | 164745756 | 164746619 | ENSMUSG00000076434 | Wfdc3 | WAP four-disulfide core domain 3 |
| chr2 | 164785490 | 164786358 | ENSMUSG00000050373 | Snx21 | sorting nexin family member 21 |
| chr2 | 164878509 | 164879337 | ENSMUSG00000039849 | Pcif1 | phosphorylated CTD interacting factor 1 |
| chr2 | 164879337 | 164880139 | ENSMUSG00000039849 | Pcif1 | phosphorylated CTD interacting factor 1 |
| chr2 | 165034043 | 165035057 | ENSMUSG00000039804 | Ncoa5 | nuclear receptor coactivator 5 |
| chr2 | 165492815 | 165493601 | ENSMUSG00000039725 | Trp53rka | transformation related protein 53 regulating kinase A |
| chr2 | 165992236 | 165993496 | ENSMUSG00000027678 | Ncoa3 | nuclear receptor coactivator 3 |
| chr2 | 166805443 | 166806267 | ENSMUSG00000074582 | Arfgef2 | ADP-ribosylation factor guanine nucleotide-exchange factor 2 (brefeldin A-inhibited) |
| chr2 | 166905841 | 166906622 | ENSMUSG00000002718 | Cse1l | chromosome segregation 1-like (S. cerevisiae) |
| chr2 | 166995650 | 166996503 | ENSMUSG00000039536 | Stau1 | staufen double-stranded RNA binding protein 1 |
| chr2 | 167062408 | 167063279 | ENSMUSG00000074578 | Zfas1 | zinc finger, NFX1-type containing 1, antisense RNA 1 |
| chr2 | 167348456 | 167349981 | ENSMUSG00000017929 | B4galt5 | UDP-Gal:betaGlcNAc beta 1,4-galactosyltransferase, polypeptide 5 |
| chr2 | 167503266 | 167504409 | ENSMUSG00000006418 | Rnf114 | ring finger protein 114 |
| chr2 | 167631180 | 167632402 | ENSMUSG00000078923 | Ube2v1 | ubiquitin-conjugating enzyme E2 variant 1 |
| chr2 | 167661126 | 167662002 | ENSMUSG00000090213 | Tmem189 | transmembrane protein 189 |
| chr2 | 167777540 | 167778406 | ENSMUSG00000006462 | A530013C23Rik | RIKEN cDNA A530013C23 gene |
| chr2 | 167931881 | 167932968 | ENSMUSG00000027540 | Ptpn1 | protein tyrosine phosphatase, non-receptor type 1 |
| chr2 | 168003367 | 168004110 | ENSMUSG00000074577 | Ripor3 | RIPOR family member 3 |
| chr2 | 168080480 | 168081954 | ENSMUSG00000044641 | Pard6b | par-6 family cell polarity regulator beta |
| chr2 | 168206322 | 168207092 | ENSMUSG00000051149 | Adnp | activity-dependent neuroprotective protein |
| chr2 | 168741292 | 168742506 | ENSMUSG00000027546 | Atp9a | ATPase, class II, type 9A |
| chr2 | 170130300 | 170131747 | ENSMUSG00000052056 | Zfp217 | zinc finger protein 217 |
| chr2 | 173000026 | 173000803 | ENSMUSG00000027509 | Rae1 | ribonucleic acid export 1 |
| chr2 | 173263619 | 173264691 | ENSMUSG00000038400 | Pmepa1 | prostate transmembrane protein, androgen induced 1 |
| chr2 | 173737371 | 173738004 | ENSMUSG00000054455 | Vapb | vesicle-associated membrane protein, associated protein B and C |
| chr2 | 174109998 | 174111079 | ENSMUSG00000039263 | Npepl1 | aminopeptidase-like 1 |
| chr2 | 174329551 | 174331262 | ENSMUSG00000027523 | Gnas | GNAS (guanine nucleotide binding protein, alpha stimulating) complex locus |
| chr2 | 174442529 | 174442879 | ENSMUSG00000016256 | Ctsz | cathepsin Z |
| chr2 | 180118810 | 180119959 | ENSMUSG00000039050 | Osbpl2 | oxysterol binding protein-like 2 |
| chr3 | 5575728 | 5576735 | ENSMUSG00000040374 | Pex2 | peroxisomal biogenesis factor 2 |
| chr3 | 7503308 | 7503851 | ENSMUSG00000043542 | Zc2hc1a | zinc finger, C2HC-type containing 1A |
| chr3 | 8666924 | 8668174 | ENSMUSG00000040289 | Hey1 | hairy/enhancer-of-split related with YRPW motif 1 |
| chr3 | 8923145 | 8923936 | ENSMUSG00000040269 | Mrps28 | mitochondrial ribosomal protein S28 |
| chr3 | 9004071 | 9004766 | ENSMUSG00000027506 | Tpd52 | tumor protein D52 |
| chr3 | 10439422 | 10440154 | ENSMUSG00000027534 | Snx16 | sorting nexin 16 |
| chr3 | 14578520 | 14579333 | ENSMUSG00000027552 | E2f5 | E2F transcription factor 5 |
| chr3 | 14886053 | 14887036 | ENSMUSG00000027562 | Car2 | carbonic anhydrase 2 |
| chr3 | 16182994 | 16183866 | ENSMUSG00000047213 | Ythdf3 | YTH N6-methyladenosine RNA binding protein 3 |
| chr3 | 19187766 | 19188602 | ENSMUSG00000027601 | Mtfr1 | mitochondrial fission regulator 1 |
| chr3 | 19310572 | 19312005 | ENSMUSG00000069094 | Pde7a | phosphodiesterase 7A |
| chr3 | 22076593 | 22077427 | ENSMUSG00000027630 | Tbl1xr1 | transducin (beta)-like 1X-linked receptor 1 |
| chr3 | 27709896 | 27711330 | ENSMUSG00000039286 | Fndc3b | fibronectin type III domain containing 3B |
| chr3 | 28805300 | 28805855 | ENSMUSG00000039221 | Rpl22l1 | ribosomal protein L22 like 1 |
| chr3 | 30792917 | 30794189 | ENSMUSG00000027706 | Sec62 | SEC62 homolog (S. cerevisiae) |
| chr3 | 30855979 | 30856497 | ENSMUSG00000037661 | Gpr160 | G protein-coupled receptor 160 |
| chr3 | 30995646 | 30996227 | ENSMUSG00000037643 | Prkci | protein kinase C, iota |
| chr3 | 32396806 | 32397853 | ENSMUSG00000027665 | Pik3ca | phosphatidylinositol-4,5-bisphosphate 3-kinase catalytic subunit alpha |
| chr3 | 32510342 | 32511448 | ENSMUSG00000027667 | Zfp639 | zinc finger protein 639 |
| chr3 | 32529427 | 32529977 | ENSMUSG00000027668 | Mfn1 | mitofusin 1 |
| chr3 | 33799833 | 33800695 | ENSMUSG00000027677 | Ttc14 | tetratricopeptide repeat domain 14 |
| chr3 | 34019524 | 34020888 | ENSMUSG00000027680 | Fxr1 | fragile X mental retardation gene 1, autosomal homolog |
| chr3 | 35753816 | 35754879 | ENSMUSG00000037400 | Atp11b | ATPase, class VI, type 11B |
| chr3 | 35932164 | 35933202 | ENSMUSG00000027708 | Dcun1d1 | DCN1, defective in cullin neddylation 1, domain containing 1 (S. cerevisiae) |
| chr3 | 36065921 | 36066439 | ENSMUSG00000027710 | Acad9 | acyl-Coenzyme A dehydrogenase family, member 9 |
| chr3 | 36571274 | 36572424 | ENSMUSG00000027715 | Ccna2 | cyclin A2 |
| chr3 | 37639750 | 37641235 | ENSMUSG00000037211 | Spry1 | sprouty RTK signaling antagonist 1 |
| chr3 | 38484260 | 38485322 | ENSMUSG00000044864 | Ankrd50 | ankyrin repeat domain 50 |
| chr3 | 40744835 | 40746021 | ENSMUSG00000025757 | Hspa4l | heat shock protein 4 like |
| chr3 | 40846471 | 40847348 | ENSMUSG00000107741 | Gm2011 | predicted gene 2011 |
| chr3 | 41082185 | 41083853 | ENSMUSG00000049940 | Pgrmc2 | progesterone receptor membrane component 2 |
| chr3 | 41563268 | 41563863 | ENSMUSG00000025764 | Jade1 | jade family PHD finger 1 |
| chr3 | 41564443 | 41566363 | ENSMUSG00000025764 | Jade1 | jade family PHD finger 1 |
| chr3 | 41741837 | 41742821 | ENSMUSG00000059834 | Sclt1 | sodium channel and clathrin linker 1 |
| chr3 | 51224353 | 51225649 | ENSMUSG00000023087 | Noct | nocturnin |
| chr3 | 51275927 | 51277912 | ENSMUSG00000087440 | 4930577N17Rik | RIKEN cDNA 4930577N17 gene |
| chr3 | 51415712 | 51416860 | ENSMUSG00000063273 | Naa15 | N(alpha)-acetyltransferase 15, NatA auxiliary subunit |
| chr3 | 51483659 | 51484765 | ENSMUSG00000027739 | Rab33b | RAB33B, member RAS oncogene family |
| chr3 | 51559791 | 51561340 |  | 5031434O11Rik | RIKEN cDNA 5031434O11 gene |
| chr3 | 53462910 | 53464055 | ENSMUSG00000042997 | Nhlrc3 | NHL repeat containing 3 |
| chr3 | 54734367 | 54735832 | ENSMUSG00000036632 | Alg5 | asparagine-linked glycosylation 5 (dolichyl-phosphate beta-glucosyltransferase) |
| chr3 | 54735832 | 54736494 | ENSMUSG00000036632 | Alg5 | asparagine-linked glycosylation 5 (dolichyl-phosphate beta-glucosyltransferase) |
| chr3 | 54807038 | 54808173 | ENSMUSG00000036615 | Rfxap | regulatory factor X-associated protein |
| chr3 | 56182781 | 56184135 | ENSMUSG00000027799 | Nbea | neurobeachin |
| chr3 | 57574770 | 57576676 | ENSMUSG00000027803 | Wwtr1 | WW domain containing transcription regulator 1 |
| chr3 | 58525183 | 58526169 | ENSMUSG00000027810 | Eif2a | eukaryotic translation initiation factor 2A |
| chr3 | 58576700 | 58577336 | ENSMUSG00000075700 | Selenot | selenoprotein T |
| chr3 | 58691452 | 58692523 | ENSMUSG00000036432 | Siah2 | siah E3 ubiquitin protein ligase 2 |
| chr3 | 60500812 | 60501935 | ENSMUSG00000027763 | Mbnl1 | muscleblind like splicing factor 1 |
| chr3 | 61364195 | 61365779 | ENSMUSG00000036894 | Rap2b | RAP2B, member of RAS oncogene family |
| chr3 | 63295355 | 63296225 | ENSMUSG00000027820 | Mme | membrane metallo endopeptidase |
| chr3 | 63929159 | 63929482 | ENSMUSG00000048581 | E130311K13Rik | RIKEN cDNA E130311K13 gene |
| chr3 | 65658739 | 65659777 | ENSMUSG00000098944 | Mir8120 | microRNA 8120 |
| chr3 | 65957423 | 65958661 | ENSMUSG00000027829 | Ccnl1 | cyclin L1 |
| chr3 | 66981037 | 66981492 | ENSMUSG00000027833 | Shox2 | short stature homeobox 2 |
| chr3 | 69005071 | 69006541 | ENSMUSG00000034349 | Smc4 | structural maintenance of chromosomes 4 |
| chr3 | 69044140 | 69045442 | ENSMUSG00000034317 | Trim59 | tripartite motif-containing 59 |
| chr3 | 69126293 | 69127098 | ENSMUSG00000027782 | Kpna4 | karyopherin (importin) alpha 4 |
| chr3 | 69316552 | 69317565 | ENSMUSG00000027784 | Ppm1l | protein phosphatase 1 (formerly 2C)-like |
| chr3 | 69721975 | 69722710 | ENSMUSG00000027787 | Nmd3 | NMD3 ribosome export adaptor |
| chr3 | 79286380 | 79287710 | ENSMUSG00000062232 | Rapgef2 | Rap guanine nucleotide exchange factor (GEF) 2 |
| chr3 | 79567300 | 79568192 | ENSMUSG00000061175 | Fnip2 | folliculin interacting protein 2 |
| chr3 | 84039051 | 84040508 | ENSMUSG00000033767 | Tmem131l | transmembrane 131 like |
| chr3 | 84480630 | 84481447 | ENSMUSG00000041842 | Fhdc1 | FH2 domain containing 1 |
| chr3 | 84813967 | 84815769 | ENSMUSG00000028086 | Fbxw7 | F-box and WD-40 domain protein 7 |
| chr3 | 85970912 | 85971921 | ENSMUSG00000028082 | Sh3d19 | SH3 domain protein D19 |
| chr3 | 86141889 | 86142820 | ENSMUSG00000028081 | Rps3a1 | ribosomal protein S3A1 |
| chr3 | 86223596 | 86225488 | ENSMUSG00000028080 | Lrba | LPS-responsive beige-like anchor |
| chr3 | 87617268 | 87617853 | ENSMUSG00000041977 | Arhgef11 | Rho guanine nucleotide exchange factor (GEF) 11 |
| chr3 | 87905866 | 87907693 | ENSMUSG00000004897 | Hdgf | heparin binding growth factor |
| chr3 | 87929664 | 87930802 | ENSMUSG00000004896 | Rrnad1 | ribosomal RNA adenine dimethylase domain containing 1 |
| chr3 | 87971003 | 87972057 | ENSMUSG00000004891 | Nes | nestin |
| chr3 | 88057951 | 88058751 | ENSMUSG00000028070 | Naxe | NAD(P)HX epimerase |
| chr3 | 88142012 | 88143267 | ENSMUSG00000001419 | Mef2d | myocyte enhancer factor 2D |
| chr3 | 88177533 | 88178137 | ENSMUSG00000096935 | 1700113A16Rik | RIKEN cDNA 1700113A16 gene |
| chr3 | 88409801 | 88410286 | ENSMUSG00000028066 | Pmf1 | polyamine-modulated factor 1 |
| chr3 | 88509707 | 88510250 | ENSMUSG00000028063 | Lmna | lamin A |
| chr3 | 88621734 | 88622711 | ENSMUSG00000028059 | Arhgef2 | rho/rac guanine nucleotide exchange factor (GEF) 2 |
| chr3 | 88685616 | 88686578 | ENSMUSG00000028060 | Khdc4 | KH domain containing 4, pre-mRNA splicing factor |
| chr3 | 88949659 | 88951253 | ENSMUSG00000028053 | Ash1l | ASH1 like histone lysine methyltransferase |
| chr3 | 89349667 | 89350544 | ENSMUSG00000028041 | Adam15 | a disintegrin and metallopeptidase domain 15 (metargidin) |
| chr3 | 89773026 | 89774147 | ENSMUSG00000042572 | Ube2q1 | ubiquitin-conjugating enzyme E2Q family member 1 |
| chr3 | 89912658 | 89913326 | ENSMUSG00000027947 | Il6ra | interleukin 6 receptor, alpha |
| chr3 | 90265156 | 90266060 | ENSMUSG00000042404 | Dennd4b | DENN/MADD domain containing 4B |
| chr3 | 90508864 | 90509399 | ENSMUSG00000001017 | Chtop | chromatin target of PRMT1 |
| chr3 | 94837990 | 94838438 | ENSMUSG00000038902 | Pogz | pogo transposable element with ZNF domain |
| chr3 | 95228088 | 95229094 | ENSMUSG00000053192 | Mllt11 | myeloid/lymphoid or mixed-lineage leukemia; translocated to, 11 |
| chr3 | 95229094 | 95229877 | ENSMUSG00000046722 | Cdc42se1 | CDC42 small effector 1 |
| chr3 | 95315070 | 95315937 | ENSMUSG00000015714 | Cers2 | ceramide synthase 2 |
| chr3 | 95658448 | 95659671 | ENSMUSG00000038612 | Mcl1 | myeloid cell leukemia sequence 1 |
| chr3 | 95659671 | 95659919 | ENSMUSG00000038612 | Mcl1 | myeloid cell leukemia sequence 1 |
| chr3 | 95818210 | 95819932 | ENSMUSG00000028106 | Rprd2 | regulation of nuclear pre-mRNA domain containing 2 |
| chr3 | 95881848 | 95882959 | ENSMUSG00000038550 | Ciart | circadian associated repressor of transcription |
| chr3 | 95929130 | 95929756 | ENSMUSG00000015749 | Anp32e | acidic (leucine-rich) nuclear phosphoprotein 32 family, member E |
| chr3 | 95929756 | 95930389 | ENSMUSG00000015749 | Anp32e | acidic (leucine-rich) nuclear phosphoprotein 32 family, member E |
| chr3 | 96104255 | 96104996 | ENSMUSG00000038495 | Otud7b | OTU domain containing 7B |
| chr3 | 96219800 | 96221732 | ENSMUSG00000068855 | H2ac20 | H2A clustered histone 20 |
| chr3 | 96245098 | 96245482 | ENSMUSG00000064220 | H2ac18 | H2A clustered histone 18 |
| chr3 | 96635238 | 96635844 | ENSMUSG00000028102 | Pex11b | peroxisomal biogenesis factor 11 beta |
| chr3 | 96696811 | 96697520 | ENSMUSG00000028101 | Pias3 | protein inhibitor of activated STAT 3 |
| chr3 | 96727360 | 96728092 | ENSMUSG00000028098 | Rnf115 | ring finger protein 115 |
| chr3 | 97609740 | 97610367 | ENSMUSG00000028089 | Chd1l | chromodomain helicase DNA binding protein 1-like |
| chr3 | 97658280 | 97658819 | ENSMUSG00000038205 | Prkab2 | protein kinase, AMP-activated, beta 2 non-catalytic subunit |
| chr3 | 98013301 | 98014053 | ENSMUSG00000027878 | Notch2 | notch 2 |
| chr3 | 98338669 | 98339845 | ENSMUSG00000053398 | Phgdh | 3-phosphoglycerate dehydrogenase |
| chr3 | 100684871 | 100686438 | ENSMUSG00000008763 | Man1a2 | mannosidase, alpha, class 1A, member 2 |
| chr3 | 101109705 | 101110559 | ENSMUSG00000027864 | Ptgfrn | prostaglandin F2 receptor negative regulator |
| chr3 | 101376937 | 101377741 | ENSMUSG00000042035 | Igsf3 | immunoglobulin superfamily, member 3 |
| chr3 | 101603938 | 101604981 | ENSMUSG00000033161 | Atp1a1 | ATPase, Na+/K+ transporting, alpha 1 polypeptide |
| chr3 | 101923848 | 101924638 | ENSMUSG00000033147 | Slc22a15 | solute carrier family 22 (organic anion/cation transporter), member 15 |
| chr3 | 102203817 | 102204747 | ENSMUSG00000027860 | Vangl1 | VANGL planar cell polarity 1 |
| chr3 | 102469855 | 102470683 | ENSMUSG00000027859 | Ngf | nerve growth factor |
| chr3 | 103020323 | 103021102 | ENSMUSG00000068823 | Csde1 | cold shock domain containing E1, RNA binding |
| chr3 | 103102063 | 103103033 | ENSMUSG00000007379 | Dennd2c | DENN/MADD domain containing 2C |
| chr3 | 103279125 | 103280661 | ENSMUSG00000033014 | Trim33 | tripartite motif-containing 33 |
| chr3 | 103790591 | 103791484 | ENSMUSG00000008730 | Hipk1 | homeodomain interacting protein kinase 1 |
| chr3 | 103967968 | 103968970 | ENSMUSG00000089998 | Phtf1os | putative homeodomain transcription factor 1, opposite strand |
| chr3 | 104219457 | 104220573 | ENSMUSG00000052539 | Magi3 | membrane associated guanylate kinase, WW and PDZ domain containing 3 |
| chr3 | 104511329 | 104512270 | ENSMUSG00000032913 | Lrig2 | leucine-rich repeats and immunoglobulin-like domains 2 |
| chr3 | 104637999 | 104639510 | ENSMUSG00000032902 | Slc16a1 | solute carrier family 16 (monocarboxylic acid transporters), member 1 |
| chr3 | 105686886 | 105687927 | ENSMUSG00000027905 | Ddx20 | DEAD box helicase 20 |
| chr3 | 105705093 | 105705629 | ENSMUSG00000048458 | Inka2 | inka box actin regulator 2 |
| chr3 | 105800757 | 105801445 | ENSMUSG00000068798 | Rap1a | RAS-related protein 1a |
| chr3 | 106547200 | 106548246 | ENSMUSG00000027900 | Dram2 | DNA-damage regulated autophagy modulator 2 |
| chr3 | 107517160 | 107518224 | ENSMUSG00000027894 | Slc6a17 | solute carrier family 6 (neurotransmitter transporter), member 17 |
| chr3 | 107695907 | 107696767 | ENSMUSG00000027893 | Ahcyl1 | S-adenosylhomocysteine hydrolase-like 1 |
| chr3 | 107759752 | 107760511 | ENSMUSG00000014599 | Csf1 | colony stimulating factor 1 (macrophage) |
| chr3 | 108401620 | 108402470 | ENSMUSG00000068740 | Celsr2 | cadherin, EGF LAG seven-pass G-type receptor 2 |
| chr3 | 108591197 | 108592248 | ENSMUSG00000040389 | Wdr47 | WD repeat domain 47 |
| chr3 | 108911011 | 108911770 | ENSMUSG00000027881 | Prpf38b | PRP38 pre-mRNA processing factor 38 (yeast) domain containing B |
| chr3 | 109026469 | 109027979 | ENSMUSG00000040339 | Fam102b | family with sequence similarity 102, member B |
| chr3 | 109340525 | 109341166 | ENSMUSG00000033721 | Vav3 | vav 3 oncogene |
| chr3 | 116661795 | 116662873 | ENSMUSG00000089911 | Mfsd14a | major facilitator superfamily domain containing 14A |
| chr3 | 116711850 | 116712506 | ENSMUSG00000027957 | Slc35a3 | solute carrier family 35 (UDP-N-acetylglucosamine (UDP-GlcNAc) transporter), member 3 |
| chr3 | 117868628 | 117869186 | ENSMUSG00000028007 | Snx7 | sorting nexin 7 |
| chr3 | 119782472 | 119783672 | ENSMUSG00000028134 | Ptbp2 | polypyrimidine tract binding protein 2 |
| chr3 | 121262511 | 121263257 | ENSMUSG00000028132 | Tlcd4 | TLC domain containing 4 |
| chr3 | 121426185 | 121427335 | ENSMUSG00000053931 | Cnn3 | calponin 3, acidic |
| chr3 | 121427335 | 121428050 | ENSMUSG00000053931 | Cnn3 | calponin 3, acidic |
| chr3 | 121722889 | 121723863 | ENSMUSG00000028128 | F3 | coagulation factor III |
| chr3 | 121814615 | 121815550 | ENSMUSG00000028127 | Abcd3 | ATP-binding cassette, sub-family D (ALD), member 3 |
| chr3 | 121953061 | 121954013 | ENSMUSG00000039831 | Arhgap29 | Rho GTPase activating protein 29 |
| chr3 | 122245244 | 122246372 | ENSMUSG00000028124 | Gclm | glutamate-cysteine ligase, modifier subunit |
| chr3 | 122292953 | 122294179 | ENSMUSG00000028121 | Bcar3 | breast cancer anti-estrogen resistance 3 |
| chr3 | 122419542 | 122420969 | ENSMUSG00000028121 | Bcar3 | breast cancer anti-estrogen resistance 3 |
| chr3 | 122983864 | 122985029 | ENSMUSG00000039701 | Usp53 | ubiquitin specific peptidase 53 |
| chr3 | 123267067 | 123268192 | ENSMUSG00000039234 | Sec24d | Sec24 related gene family, member D (S. cerevisiae) |
| chr3 | 123690633 | 123691080 | ENSMUSG00000027977 | Ndst3 | N-deacetylase/N-sulfotransferase (heparan glucosaminyl) 3 |
| chr3 | 126596225 | 126597813 | ENSMUSG00000053819 | Camk2d | calcium/calmodulin-dependent protein kinase II, delta |
| chr3 | 127633008 | 127634628 | ENSMUSG00000027967 | Neurog2 | neurogenin 2 |
| chr3 | 127836908 | 127837916 | ENSMUSG00000074238 | Ap1ar | adaptor-related protein complex 1 associated regulatory protein |
| chr3 | 127895685 | 127896466 | ENSMUSG00000050549 | Fam241a | family with sequence similarity 241, member A |
| chr3 | 129533483 | 129534606 | ENSMUSG00000041220 | Elovl6 | ELOVL family member 6, elongation of long chain fatty acids (yeast) |
| chr3 | 129830681 | 129831627 | ENSMUSG00000028010 | Gar1 | GAR1 ribonucleoprotein |
| chr3 | 129878510 | 129878912 | ENSMUSG00000027999 | Pla2g12a | phospholipase A2, group XIIA |
| chr3 | 130060039 | 130061074 | ENSMUSG00000001052 | Sec24b | Sec24 related gene family, member B (S. cerevisiae) |
| chr3 | 131302406 | 131303349 | ENSMUSG00000027983 | Cyp2u1 | cytochrome P450, family 2, subfamily u, polypeptide 1 |
| chr3 | 131489590 | 131491019 | ENSMUSG00000050931 | Sgms2 | sphingomyelin synthase 2 |
| chr3 | 131564475 | 131565274 | ENSMUSG00000028032 | Papss1 | 3'-phosphoadenosine 5'-phosphosulfate synthase 1 |
| chr3 | 132949657 | 132950440 | ENSMUSG00000040998 | Npnt | nephronectin |
| chr3 | 133309966 | 133310664 | ENSMUSG00000028013 | Ppa2 | pyrophosphatase (inorganic) 2 |
| chr3 | 133544197 | 133545006 | ENSMUSG00000040943 | Tet2 | tet methylcytosine dioxygenase 2 |
| chr3 | 135437945 | 135439690 | ENSMUSG00000097032 | 4930539J05Rik | RIKEN cDNA 4930539J05 gene |
| chr3 | 135690682 | 135691743 | ENSMUSG00000028163 | Nfkb1 | nuclear factor of kappa light polypeptide gene enhancer in B cells 1, p105 |
| chr3 | 137863997 | 137865900 | ENSMUSG00000037894 | H2az1 | H2A.Z variant histone 1 |
| chr3 | 137867307 | 137868248 | ENSMUSG00000074212 | Dnajb14 | DnaJ heat shock protein family (Hsp40) member B14 |
| chr3 | 138143383 | 138144058 | ENSMUSG00000004127 | Trmt10a | tRNA methyltransferase 10A |
| chr3 | 138488813 | 138489849 | ENSMUSG00000005813 | Metap1 | methionyl aminopeptidase 1 |
| chr3 | 138526378 | 138527261 | ENSMUSG00000088373 | Mir1956 | microRNA 1956 |
| chr3 | 138527261 | 138528242 | ENSMUSG00000028156 | Eif4e | eukaryotic translation initiation factor 4E |
| chr3 | 138741942 | 138743608 | ENSMUSG00000028152 | Tspan5 | tetraspanin 5 |
| chr3 | 139074429 | 139075949 | ENSMUSG00000028149 | Rap1gds1 | RAP1, GTP-GDP dissociation stimulator 1 |
| chr3 | 142394654 | 142395949 | ENSMUSG00000028273 | Pdlim5 | PDZ and LIM domain 5 |
| chr3 | 142764908 | 142765527 | ENSMUSG00000028271 | Gtf2b | general transcription factor IIB |
| chr3 | 142881075 | 142882548 | ENSMUSG00000004591 | Pkn2 | protein kinase N2 |
| chr3 | 144198496 | 144198856 | ENSMUSG00000028266 | Lmo4 | LIM domain only 4 |
| chr3 | 144201705 | 144202615 | ENSMUSG00000028266 | Lmo4 | LIM domain only 4 |
| chr3 | 144204863 | 144205871 | ENSMUSG00000028266 | Lmo4 | LIM domain only 4 |
| chr3 | 144569584 | 144571077 | ENSMUSG00000040151 | Hs2st1 | heparan sulfate 2-O-sulfotransferase 1 |
| chr3 | 144719263 | 144720545 | ENSMUSG00000037062 | Sh3glb1 | SH3-domain GRB2-like B1 (endophilin) |
| chr3 | 145576090 | 145576772 | ENSMUSG00000074182 | Znhit6 | zinc finger, HIT type 6 |
| chr3 | 145649117 | 145650154 | ENSMUSG00000028195 | Ccn1 | cellular communication network factor 1 |
| chr3 | 145653128 | 145653784 | ENSMUSG00000028195 | Ccn1 | cellular communication network factor 1 |
| chr3 | 145758435 | 145759358 | ENSMUSG00000028194 | Ddah1 | dimethylarginine dimethylaminohydrolase 1 |
| chr3 | 145924167 | 145925032 | ENSMUSG00000028191 | Bcl10 | B cell leukemia/lymphoma 10 |
| chr3 | 145987485 | 145989179 | ENSMUSG00000036863 | Syde2 | synapse defective 1, Rho GTPase, homolog 2 (C. elegans) |
| chr3 | 146404498 | 146405102 | ENSMUSG00000036825 | Ssx2ip | synovial sarcoma, X 2 interacting protein |
| chr3 | 146499346 | 146500388 | ENSMUSG00000068523 | Gng5 | guanine nucleotide binding protein (G protein), gamma 5 |
| chr3 | 152165748 | 152166214 | ENSMUSG00000039131 | Gipc2 | GIPC PDZ domain containing family, member 2 |
| chr3 | 152209716 | 152211245 | ENSMUSG00000028034 | Fubp1 | far upstream element (FUSE) binding protein 1 |
| chr3 | 152395373 | 152397219 | ENSMUSG00000039068 | Zzz3 | zinc finger, ZZ domain containing 3 |
| chr3 | 153912608 | 153913200 | ENSMUSG00000038975 | Rabggtb | Rab geranylgeranyl transferase, b subunit |
| chr3 | 157946880 | 157948246 | ENSMUSG00000039988 | Ankrd13c | ankyrin repeat domain 13c |
| chr3 | 158031237 | 158031992 | ENSMUSG00000055436 | Srsf11 | serine and arginine-rich splicing factor 11 |
| chr4 | 6453514 | 6454352 | ENSMUSG00000028245 | Nsmaf | neutral sphingomyelinase (N-SMase) activation associated factor |
| chr4 | 8647037 | 8647662 | ENSMUSG00000041235 | Chd7 | chromodomain helicase DNA binding protein 7 |
| chr4 | 11006933 | 11008104 | ENSMUSG00000049969 | Plekhf2 | pleckstrin homology domain containing, family F (with FYVE domain) member 2 |
| chr4 | 11190471 | 11192312 | ENSMUSG00000028212 | Ccne2 | cyclin E2 |
| chr4 | 11965687 | 11966992 |  | 1700123M08Rik | RIKEN cDNA 1700123M08 gene |
| chr4 | 14864023 | 14864760 | ENSMUSG00000028221 | Pip4p2 | phosphatidylinositol-4,5-bisphosphate 4-phosphatase 2 |
| chr4 | 15265516 | 15266646 | ENSMUSG00000043252 | Tmem64 | transmembrane protein 64 |
| chr4 | 21930987 | 21931739 | ENSMUSG00000028246 | Faxc | failed axon connections homolog |
| chr4 | 24966229 | 24967052 | ENSMUSG00000040372 | Gpr63 | G protein-coupled receptor 63 |
| chr4 | 25281609 | 25282000 | ENSMUSG00000040359 | Ufl1 | UFM1 specific ligase 1 |
| chr4 | 32238493 | 32240007 | ENSMUSG00000040270 | Bach2 | BTB and CNC homology, basic leucine zipper transcription factor 2 |
| chr4 | 33031014 | 33032037 | ENSMUSG00000085037 | 4933421O10Rik | RIKEN cDNA 4933421O10 gene |
| chr4 | 33247421 | 33248742 | ENSMUSG00000040128 | Pnrc1 | proline-rich nuclear receptor coactivator 1 |
| chr4 | 34686873 | 34687519 | ENSMUSG00000028293 | Slc35a1 | solute carrier family 35 (CMP-sialic acid transporter), member 1 |
| chr4 | 34882411 | 34883174 | ENSMUSG00000039967 | Zfp292 | zinc finger protein 292 |
| chr4 | 34886582 | 34887111 | ENSMUSG00000039967 | Zfp292 | zinc finger protein 292 |
| chr4 | 40722173 | 40723245 | ENSMUSG00000028410 | Dnaja1 | DnaJ heat shock protein family (Hsp40) member A1 |
| chr4 | 40853398 | 40854277 | ENSMUSG00000028413 | B4galt1 | UDP-Gal:betaGlcNAc beta 1,4- galactosyltransferase, polypeptide 1 |
| chr4 | 43381719 | 43382306 | ENSMUSG00000035969 | Rusc2 | RUN and SH3 domain containing 2 |
| chr4 | 43492761 | 43493804 | ENSMUSG00000088088 | Rmrp | RNA component of mitochondrial RNAase P |
| chr4 | 44072314 | 44072966 | ENSMUSG00000028479 | Gne | glucosamine (UDP-N-acetyl)-2-epimerase/N-acetylmannosamine kinase |
| chr4 | 44167324 | 44168500 | ENSMUSG00000035696 | Rnf38 | ring finger protein 38 |
| chr4 | 45529561 | 45531232 | ENSMUSG00000044813 | Shb | src homology 2 domain-containing transforming protein B |
| chr4 | 45972224 | 45972943 | ENSMUSG00000035517 | Tdrd7 | tudor domain containing 7 |
| chr4 | 47352817 | 47353953 | ENSMUSG00000007613 | Tgfbr1 | transforming growth factor, beta receptor I |
| chr4 | 48044492 | 48045550 | ENSMUSG00000028341 | Nr4a3 | nuclear receptor subfamily 4, group A, member 3 |
| chr4 | 53439892 | 53441323 | ENSMUSG00000028412 | Slc44a1 | solute carrier family 44, member 1 |
| chr4 | 56946596 | 56947518 | ENSMUSG00000055296 | Tmem245 | transmembrane protein 245 |
| chr4 | 57143339 | 57143898 | ENSMUSG00000028434 | Epb41l4b | erythrocyte membrane protein band 4.1 like 4b |
| chr4 | 58911493 | 58912860 | ENSMUSG00000050812 | Ecpas | Ecm29 proteasome adaptor and scaffold |
| chr4 | 59189069 | 59189742 | ENSMUSG00000028381 | Ugcg | UDP-glucose ceramide glucosyltransferase |
| chr4 | 59438071 | 59438748 | ENSMUSG00000038578 | Susd1 | sushi domain containing 1 |
| chr4 | 59548560 | 59549735 | ENSMUSG00000028382 | Ptbp3 | polypyrimidine tract binding protein 3 |
| chr4 | 59625892 | 59626851 | ENSMUSG00000045071 | E130308A19Rik | RIKEN cDNA E130308A19 gene |
| chr4 | 62519656 | 62520283 | ENSMUSG00000028393 | Alad | aminolevulinate, delta-, dehydratase |
| chr4 | 63558393 | 63558964 | ENSMUSG00000045917 | Tmem268 | transmembrane protein 268 |
| chr4 | 72199754 | 72201406 | ENSMUSG00000008305 | Tle1 | transducin-like enhancer of split 1 |
| chr4 | 74251601 | 74252474 | ENSMUSG00000028397 | Kdm4c | lysine (K)-specific demethylase 4C |
| chr4 | 81442498 | 81443029 | ENSMUSG00000028402 | Mpdz | multiple PDZ domain crumbs cell polarity complex component |
| chr4 | 82506119 | 82506981 | ENSMUSG00000008575 | Nfib | nuclear factor I/B |
| chr4 | 82512995 | 82513508 | ENSMUSG00000087413 | Gm11266 | predicted gene 11266 |
| chr4 | 82859366 | 82859888 | ENSMUSG00000028403 | Zdhhc21 | zinc finger, DHHC domain containing 21 |
| chr4 | 83323484 | 83324459 | ENSMUSG00000038172 | Ttc39b | tetratricopeptide repeat domain 39B |
| chr4 | 83485723 | 83486629 | ENSMUSG00000028484 | Psip1 | PC4 and SFRS1 interacting protein 1 |
| chr4 | 83525188 | 83525950 | ENSMUSG00000052407 | Ccdc171 | coiled-coil domain containing 171 |
| chr4 | 84545326 | 84545795 | ENSMUSG00000028487 | Bnc2 | basonuclin 2 |
| chr4 | 88094400 | 88095101 | ENSMUSG00000038368 | Focad | focadhesin |
| chr4 | 88937726 | 88938863 |  | Gm49890 | predicted gene, 49890 |
| chr4 | 89294201 | 89295300 | ENSMUSG00000044303 | Cdkn2a | cyclin dependent kinase inhibitor 2A |
| chr4 | 95967104 | 95967779 | ENSMUSG00000044125 | 9530080O11Rik | RIKEN cDNA 9530080O11 gene |
| chr4 | 97777254 | 97778614 | ENSMUSG00000028565 | Nfia | nuclear factor I/A |
| chr4 | 98395779 | 98396265 | ENSMUSG00000061859 | Patj | PATJ, crumbs cell polarity complex component |
| chr4 | 98923472 | 98924405 | ENSMUSG00000028560 | Usp1 | ubiquitin specific peptidase 1 |
| chr4 | 99120292 | 99121235 | ENSMUSG00000028556 | Dock7 | dedicator of cytokinesis 7 |
| chr4 | 103119206 | 103120022 | ENSMUSG00000028522 | Mier1 | MEIR1 treanscription regulator |
| chr4 | 105109373 | 105110125 | ENSMUSG00000028518 | Prkaa2 | protein kinase, AMP-activated, alpha 2 catalytic subunit |
| chr4 | 106315983 | 106316860 | ENSMUSG00000028514 | Usp24 | ubiquitin specific peptidase 24 |
| chr4 | 106560917 | 106561626 | ENSMUSG00000034926 | Dhcr24 | 24-dehydrocholesterol reductase |
| chr4 | 106910990 | 106912308 | ENSMUSG00000061887 | Ssbp3 | single-stranded DNA binding protein 3 |
| chr4 | 108619559 | 108620423 | ENSMUSG00000028582 | Cc2d1b | coiled-coil and C2 domain containing 1B |
| chr4 | 108779895 | 108781034 | ENSMUSG00000034557 | Zfyve9 | zinc finger, FYVE domain containing 9 |
| chr4 | 109406697 | 109407371 | ENSMUSG00000028555 | Ttc39a | tetratricopeptide repeat domain 39A |
| chr4 | 109676197 | 109677195 | ENSMUSG00000010517 | Faf1 | Fas-associated factor 1 |
| chr4 | 116463726 | 116464449 | ENSMUSG00000003810 | Mast2 | microtubule associated serine/threonine kinase 2 |
| chr4 | 116876989 | 116877666 | ENSMUSG00000033948 | Zswim5 | zinc finger SWIM-type containing 5 |
| chr4 | 117155551 | 117156204 | ENSMUSG00000092680 | Snord55 | small nucleolar RNA, C/D box 55 |
| chr4 | 117835533 | 117836703 | ENSMUSG00000028542 | Slc6a9 | solute carrier family 6 (neurotransmitter transporter, glycine), member 9 |
| chr4 | 119539226 | 119540398 | ENSMUSG00000032998 | Foxj3 | forkhead box J3 |
| chr4 | 120148680 | 120149108 | ENSMUSG00000028635 | Edn2 | endothelin 2 |
| chr4 | 120405149 | 120405838 | ENSMUSG00000000085 | Scmh1 | sex comb on midleg homolog 1 |
| chr4 | 120569582 | 120570633 | ENSMUSG00000028633 | Ctps | cytidine 5'-triphosphate synthase |
| chr4 | 123411321 | 123412222 | ENSMUSG00000028649 | Macf1 | microtubule-actin crosslinking factor 1 |
| chr4 | 123664067 | 123665227 | ENSMUSG00000028649 | Macf1 | microtubule-actin crosslinking factor 1 |
| chr4 | 123904734 | 123905558 | ENSMUSG00000028647 | Mycbp | MYC binding protein |
| chr4 | 123917102 | 123918125 | ENSMUSG00000028646 | Rragc | Ras-related GTP binding C |
| chr4 | 124850132 | 124851489 | ENSMUSG00000028889 | Yrdc | yrdC domain containing (E.coli) |
| chr4 | 126321071 | 126321767 | ENSMUSG00000042558 | Adprhl2 | ADP-ribosylhydrolase like 2 |
| chr4 | 128727483 | 128728089 | ENSMUSG00000028796 | Phc2 | polyhomeotic 2 |
| chr4 | 129057965 | 129059396 | ENSMUSG00000028793 | Rnf19b | ring finger protein 19B |
| chr4 | 129334679 | 129336150 | ENSMUSG00000057236 | Rbbp4 | retinoblastoma binding protein 4, chromatin remodeling factor |
| chr4 | 129377011 | 129377896 | ENSMUSG00000028807 | Zbtb8a | zinc finger and BTB domain containing 8a |
| chr4 | 129819702 | 129821056 | ENSMUSG00000028788 | Ptp4a2 | protein tyrosine phosphatase 4a2 |
| chr4 | 132048625 | 132049361 | ENSMUSG00000028906 | Epb41 | erythrocyte membrane protein band 4.1 |
| chr4 | 132211199 | 132212357 | ENSMUSG00000040025 | Ythdf2 | YTH N6-methyladenosine RNA binding protein 2 |
| chr4 | 133011333 | 133012357 | ENSMUSG00000037692 | Ahdc1 | AT hook, DNA binding motif, containing 1 |
| chr4 | 133338890 | 133339498 | ENSMUSG00000037622 | Wdtc1 | WD and tetratricopeptide repeats 1 |
| chr4 | 134286711 | 134287719 | ENSMUSG00000050890 | Pdik1l | PDLIM1 interacting kinase 1 like |
| chr4 | 135272125 | 135273275 | ENSMUSG00000037242 | Clic4 | chloride intracellular channel 4 (mitochondrial) |
| chr4 | 135855638 | 135856729 | ENSMUSG00000028676 | Srsf10 | serine and arginine-rich splicing factor 10 |
| chr4 | 136462393 | 136462893 | ENSMUSG00000001089 | Luzp1 | leucine zipper protein 1 |
| chr4 | 136469736 | 136470771 | ENSMUSG00000001089 | Luzp1 | leucine zipper protein 1 |
| chr4 | 136602014 | 136602842 | ENSMUSG00000036940 | Kdm1a | lysine (K)-specific demethylase 1A |
| chr4 | 137593934 | 137594645 | ENSMUSG00000043411 | Usp48 | ubiquitin specific peptidase 48 |
| chr4 | 138216436 | 138217696 | ENSMUSG00000028759 | Hp1bp3 | heterochromatin protein 1, binding protein 3 |
| chr4 | 138454744 | 138455198 | ENSMUSG00000046447 | Camk2n1 | calcium/calmodulin-dependent protein kinase II inhibitor 1 |
| chr4 | 140701146 | 140702908 | ENSMUSG00000040945 | Rcc2 | regulator of chromosome condensation 2 |
| chr4 | 141537611 | 141538930 | ENSMUSG00000040761 | Spen | spen family transcription repressor |
| chr4 | 141722803 | 141723662 | ENSMUSG00000078515 | Ddi2 | DNA-damage inducible protein 2 |
| chr4 | 142238755 | 142239539 | ENSMUSG00000040606 | Kazn | kazrin, periplakin interacting protein |
| chr4 | 143161598 | 143162455 | ENSMUSG00000106597 | Mir7021 | microRNA 7021 |
| chr4 | 147940860 | 147941802 | ENSMUSG00000044496 | 2510039O18Rik | RIKEN cDNA 2510039O18 gene |
| chr4 | 148900526 | 148901601 | ENSMUSG00000028977 | Casz1 | castor zinc finger 1 |
| chr4 | 149737490 | 149738198 | ENSMUSG00000044700 | Tmem201 | transmembrane protein 201 |
| chr4 | 149773454 | 149774528 | ENSMUSG00000028982 | Slc25a33 | solute carrier family 25, member 33 |
| chr4 | 149954711 | 149955497 | ENSMUSG00000039911 | Spsb1 | splA/ryanodine receptor domain and SOCS box containing 1 |
| chr4 | 150008435 | 150009205 | ENSMUSG00000028980 | H6pd | hexose-6-phosphate dehydrogenase (glucose 1-dehydrogenase) |
| chr4 | 150371657 | 150372343 | ENSMUSG00000039852 | Rere | arginine glutamic acid dipeptide (RE) repeats |
| chr4 | 150569224 | 150570040 | ENSMUSG00000039852 | Rere | arginine glutamic acid dipeptide (RE) repeats |
| chr4 | 150854889 | 150855602 | ENSMUSG00000078492 | 1700045H11Rik | RIKEN cDNA 1700045H11 gene |
| chr4 | 150909392 | 150909959 | ENSMUSG00000028964 | Park7 | Parkinson disease (autosomal recessive, early onset) 7 |
| chr4 | 151057422 | 151058136 | ENSMUSG00000028955 | Vamp3 | vesicle-associated membrane protein 3 |
| chr4 | 152008665 | 152010192 | ENSMUSG00000073700 | Klhl21 | kelch-like 21 |
| chr4 | 152010192 | 152010740 | ENSMUSG00000073700 | Klhl21 | kelch-like 21 |
| chr4 | 152039202 | 152040040 | ENSMUSG00000028948 | Nol9 | nucleolar protein 9 |
| chr4 | 152177590 | 152178736 | ENSMUSG00000028937 | Acot7 | acyl-CoA thioesterase 7 |
| chr4 | 153956957 | 153957671 | ENSMUSG00000047613 | A430005L14Rik | RIKEN cDNA A430005L14 gene |
| chr4 | 154160171 | 154160778 | ENSMUSG00000029030 | Tprgl | transformation related protein 63 regulated like |
| chr4 | 155221712 | 155222635 | ENSMUSG00000029050 | Ski | ski sarcoma viral oncogene homolog (avian) |
| chr4 | 155563671 | 155564468 | ENSMUSG00000029063 | Nadk | NAD kinase |
| chr4 | 155694387 | 155694901 | ENSMUSG00000074738 | Fndc10 | fibronectin type III domain containing 10 |
| chr4 | 155891086 | 155892846 | ENSMUSG00000029033 | Acap3 | ArfGAP with coiled-coil, ankyrin repeat and PH domains 3 |
| chr4 | 155992172 | 155993060 | ENSMUSG00000050796 | B3galt6 | UDP-Gal:betaGal beta 1,3-galactosyltransferase, polypeptide 6 |
| chr4 | 156235888 | 156236365 | ENSMUSG00000095567 | Noc2l | NOC2 like nucleolar associated transcriptional repressor |
| chr5 | 3343527 | 3344813 | ENSMUSG00000040274 | Cdk6 | cyclin-dependent kinase 6 |
| chr5 | 3543654 | 3544277 | ENSMUSG00000058503 | Fam133b | family with sequence similarity 133, member B |
| chr5 | 3647178 | 3647973 | ENSMUSG00000007415 | Gatad1 | GATA zinc finger domain containing 1 |
| chr5 | 3802327 | 3803718 | ENSMUSG00000040351 | Ankib1 | ankyrin repeat and IBR domain containing 1 |
| chr5 | 3928026 | 3929004 | ENSMUSG00000040407 | Akap9 | A kinase (PRKA) anchor protein (yotiao) 9 |
| chr5 | 4104018 | 4104928 | ENSMUSG00000001467 | Cyp51 | cytochrome P450, family 51 |
| chr5 | 4756697 | 4758357 | ENSMUSG00000044674 | Fzd1 | frizzled class receptor 1 |
| chr5 | 5379647 | 5380957 | ENSMUSG00000028926 | Cdk14 | cyclin-dependent kinase 14 |
| chr5 | 5514212 | 5514939 | ENSMUSG00000046798 | Cldn12 | claudin 12 |
| chr5 | 5693525 | 5694404 | ENSMUSG00000015653 | Steap2 | six transmembrane epithelial antigen of prostate 2 |
| chr5 | 5749005 | 5749637 | ENSMUSG00000015652 | Steap1 | six transmembrane epithelial antigen of the prostate 1 |
| chr5 | 8056348 | 8056842 | ENSMUSG00000003161 | Sri | sorcin |
| chr5 | 8422063 | 8423242 | ENSMUSG00000002297 | Dbf4 | DBF4 zinc finger |
| chr5 | 8622057 | 8623414 | ENSMUSG00000040570 | Rundc3b | RUN domain containing 3B |
| chr5 | 9100538 | 9101495 | ENSMUSG00000079659 | Tmem243 | transmembrane protein 243, mitochondrial |
| chr5 | 9160946 | 9162022 | ENSMUSG00000042508 | Dmtf1 | cyclin D binding myb-like transcription factor 1 |
| chr5 | 9265871 | 9266615 | ENSMUSG00000056004 | Elapor2 | endosome-lysosome associated apoptosis and autophagy regulator family member 2 |
| chr5 | 14514711 | 14515544 | ENSMUSG00000061601 | Pclo | piccolo (presynaptic cytomatrix protein) |
| chr5 | 17574252 | 17575052 | ENSMUSG00000028780 | Sema3c | sema domain, immunoglobulin domain (Ig), short basic domain, secreted, (semaphorin) 3C |
| chr5 | 20701808 | 20702922 | ENSMUSG00000040003 | Magi2 | membrane associated guanylate kinase, WW and PDZ domain containing 2 |
| chr5 | 20881296 | 20882748 | ENSMUSG00000045435 | Tmem60 | transmembrane protein 60 |
| chr5 | 20950859 | 20951697 | ENSMUSG00000101013 | A630072M18Rik | RIKEN cDNA A630072M18 gene |
| chr5 | 21055005 | 21056586 | ENSMUSG00000028771 | Ptpn12 | protein tyrosine phosphatase, non-receptor type 12 |
| chr5 | 21424982 | 21425587 | ENSMUSG00000047221 | Fam185a | family with sequence similarity 185, member A |
| chr5 | 21645885 | 21646512 | ENSMUSG00000038525 | Armc10 | armadillo repeat containing 10 |
| chr5 | 21701215 | 21701906 | ENSMUSG00000044968 | Napepld | N-acyl phosphatidylethanolamine phospholipase D |
| chr5 | 21784297 | 21785439 | ENSMUSG00000028932 | Psmc2 | proteasome (prosome, macropain) 26S subunit, ATPase 2 |
| chr5 | 23418305 | 23418677 | ENSMUSG00000073147 | 5031425E22Rik | RIKEN cDNA 5031425E22 gene |
| chr5 | 23433712 | 23434897 | ENSMUSG00000073147 | 5031425E22Rik | RIKEN cDNA 5031425E22 gene |
| chr5 | 23674860 | 23676222 | ENSMUSG00000062604 | Srpk2 | serine/arginine-rich protein specific kinase 2 |
| chr5 | 23783238 | 23783913 | ENSMUSG00000057541 | Pus7 | pseudouridylate synthase 7 |
| chr5 | 24164847 | 24165383 | ENSMUSG00000048439 | Nupl2 | nucleoporin like 2 |
| chr5 | 24451600 | 24453519 | ENSMUSG00000023353 | Agap3 | ArfGAP with GTPase domain, ankyrin repeat and PH domain 3 |
| chr5 | 24601612 | 24602833 | ENSMUSG00000028949 | Smarcd3 | SWI/SNF related, matrix associated, actin dependent regulator of chromatin, subfamily d, member 3 |
| chr5 | 24604484 | 24605120 | ENSMUSG00000028949 | Smarcd3 | SWI/SNF related, matrix associated, actin dependent regulator of chromatin, subfamily d, member 3 |
| chr5 | 24685588 | 24686368 | ENSMUSG00000028954 | Nub1 | negative regulator of ubiquitin-like proteins 1 |
| chr5 | 24841779 | 24843349 | ENSMUSG00000028945 | Rheb | Ras homolog enriched in brain |
| chr5 | 24907836 | 24909170 | ENSMUSG00000028944 | Prkag2 | protein kinase, AMP-activated, gamma 2 non-catalytic subunit |
| chr5 | 25222716 | 25223256 | ENSMUSG00000038072 | Galnt11 | polypeptide N-acetylgalactosaminyltransferase 11 |
| chr5 | 25529558 | 25530410 | ENSMUSG00000101856 | 1700096K18Rik | RIKEN cDNA 1700096K18 gene |
| chr5 | 27790812 | 27791916 | ENSMUSG00000002221 | Paxip1 | PAX interacting (with transcription-activation domain) protein 1 |
| chr5 | 28070682 | 28072227 | ENSMUSG00000045294 | Insig1 | insulin induced gene 1 |
| chr5 | 28316491 | 28317699 | ENSMUSG00000048271 | Rbm33 | RNA binding motif protein 33 |
| chr5 | 29568796 | 29570248 | ENSMUSG00000039000 | Ube3c | ubiquitin protein ligase E3C |
| chr5 | 30232196 | 30233418 | ENSMUSG00000075703 | Selenoi | selenoprotein I |
| chr5 | 30888787 | 30889468 | ENSMUSG00000029165 | Agbl5 | ATP/GTP binding protein-like 5 |
| chr5 | 30907277 | 30908184 | ENSMUSG00000038803 | Ost4 | oligosaccharyltransferase complex subunit 4 (non-catalytic) |
| chr5 | 31047896 | 31049012 | ENSMUSG00000006641 | Slc5a6 | solute carrier family 5 (sodium-dependent vitamin transporter), member 6 |
| chr5 | 31054602 | 31055303 | ENSMUSG00000013629 | Cad | carbamoyl-phosphate synthetase 2, aspartate transcarbamylase, and dihydroorotase |
| chr5 | 31192507 | 31193729 | ENSMUSG00000029145 | Eif2b4 | eukaryotic translation initiation factor 2B, subunit 4 delta |
| chr5 | 31219902 | 31220843 | ENSMUSG00000029147 | Ppm1g | protein phosphatase 1G (formerly 2C), magnesium-dependent, gamma isoform |
| chr5 | 31240681 | 31241732 | ENSMUSG00000029148 | Nrbp1 | nuclear receptor binding protein 1 |
| chr5 | 31697454 | 31698536 | ENSMUSG00000052139 | Babam2 | BRISC and BRCA1 A complex member 2 |
| chr5 | 32458548 | 32459789 | ENSMUSG00000014956 | Ppp1cb | protein phosphatase 1 catalytic subunit beta |
| chr5 | 32610541 | 32612014 | ENSMUSG00000014932 | Yes1 | YES proto-oncogene 1, Src family tyrosine kinase |
| chr5 | 32785174 | 32785928 | ENSMUSG00000023452 | Pisd | phosphatidylserine decarboxylase |
| chr5 | 33018222 | 33019513 | ENSMUSG00000018965 | Ywhah | tyrosine 3-monooxygenase/tryptophan 5-monooxygenase activation protein, eta polypeptide |
| chr5 | 33273925 | 33275480 | ENSMUSG00000037373 | Ctbp1 | C-terminal binding protein 1 |
| chr5 | 33335206 | 33336340 | ENSMUSG00000079562 | Maea | macrophage erythroblast attacher |
| chr5 | 33628921 | 33630034 | ENSMUSG00000037339 | Fam53a | family with sequence similarity 53, member A |
| chr5 | 33782149 | 33782938 | ENSMUSG00000005299 | Letm1 | leucine zipper-EF-hand containing transmembrane protein 1 |
| chr5 | 33819708 | 33821652 | ENSMUSG00000057406 | Nsd2 | nuclear receptor binding SET domain protein 2 |
| chr5 | 34186921 | 34188438 | ENSMUSG00000037235 | Mxd4 | Max dimerization protein 4 |
| chr5 | 34288059 | 34288717 | ENSMUSG00000109572 | Cfap99 | cilia and flagella associated protein 99 |
| chr5 | 34336097 | 34337383 | ENSMUSG00000029110 | Rnf4 | ring finger protein 4 |
| chr5 | 34369346 | 34370353 | ENSMUSG00000037210 | Fam193a | family with sequence homology 193, member A |
| chr5 | 34573411 | 34574349 | ENSMUSG00000099082 | Mir7036b | microRNA 7036b |
| chr5 | 34659797 | 34661230 | ENSMUSG00000052783 | Grk4 | G protein-coupled receptor kinase 4 |
| chr5 | 36483561 | 36484648 | ENSMUSG00000029196 | Tada2b | transcriptional adaptor 2B |
| chr5 | 36695074 | 36696330 | ENSMUSG00000029190 | D5Ertd579e | DNA segment, Chr 5, ERATO Doi 579, expressed |
| chr5 | 36795911 | 36796958 | ENSMUSG00000055302 | Mrfap1 | Morf4 family associated protein 1 |
| chr5 | 36830128 | 36830853 | ENSMUSG00000029119 | Man2b2 | mannosidase 2, alpha B2 |
| chr5 | 38219975 | 38220896 | ENSMUSG00000067367 | Lyar | Ly1 antibody reactive clone |
| chr5 | 38560606 | 38562118 | ENSMUSG00000005103 | Wdr1 | WD repeat domain 1 |
| chr5 | 39644300 | 39645284 | ENSMUSG00000051022 | Hs3st1 | heparan sulfate (glucosamine) 3-O-sulfotransferase 1 |
| chr5 | 41707016 | 41708384 | ENSMUSG00000029128 | Rab28 | RAB28, member RAS oncogene family |
| chr5 | 41843823 | 41844568 | ENSMUSG00000061755 | Bod1l | biorientation of chromosomes in cell division 1-like |
| chr5 | 43781341 | 43782563 | ENSMUSG00000039753 | Fbxl5 | F-box and leucine-rich repeat protein 5 |
| chr5 | 44225957 | 44227493 | ENSMUSG00000046985 | Tapt1 | transmembrane anterior posterior transformation 1 |
| chr5 | 45449708 | 45450628 | ENSMUSG00000015806 | Qdpr | quinoid dihydropteridine reductase |
| chr5 | 45493140 | 45493780 | ENSMUSG00000039682 | Lap3 | leucine aminopeptidase 3 |
| chr5 | 45856533 | 45857960 | ENSMUSG00000015882 | Lcorl | ligand dependent nuclear receptor corepressor-like |
| chr5 | 50058399 | 50059221 | ENSMUSG00000029090 | Adgra3 | adhesion G protein-coupled receptor A3 |
| chr5 | 52189680 | 52190860 | ENSMUSG00000029169 | Dhx15 | DEAH (Asp-Glu-Ala-His) box polypeptide 15 |
| chr5 | 52833714 | 52834400 | ENSMUSG00000029176 | Anapc4 | anaphase promoting complex subunit 4 |
| chr5 | 53995471 | 53995725 | ENSMUSG00000039156 | Stim2 | stromal interaction molecule 2 |
| chr5 | 53998041 | 53999449 | ENSMUSG00000039156 | Stim2 | stromal interaction molecule 2 |
| chr5 | 62765656 | 62766575 | ENSMUSG00000037999 | Arap2 | ArfGAP with RhoGAP domain, ankyrin repeat and PH domain 2 |
| chr5 | 63968271 | 63969183 | ENSMUSG00000047881 | Rell1 | RELT-like 1 |
| chr5 | 64159678 | 64160908 | ENSMUSG00000029174 | Tbc1d1 | TBC1 domain family, member 1 |
| chr5 | 64230180 | 64230831 | ENSMUSG00000029174 | Tbc1d1 | TBC1 domain family, member 1 |
| chr5 | 64803344 | 64804459 | ENSMUSG00000029178 | Klf3 | Kruppel-like factor 3 (basic) |
| chr5 | 65335238 | 65335894 | ENSMUSG00000029191 | Rfc1 | replication factor C (activator 1) 1 |
| chr5 | 65435515 | 65436560 | ENSMUSG00000029201 | Ugdh | UDP-glucose dehydrogenase |
| chr5 | 65492355 | 65493413 | ENSMUSG00000037822 | Smim14 | small integral membrane protein 14 |
| chr5 | 65536755 | 65538189 | ENSMUSG00000029203 | Ube2k | ubiquitin-conjugating enzyme E2K |
| chr5 | 65696738 | 65698593 | ENSMUSG00000029202 | Pds5a | PDS5 cohesin associated factor A |
| chr5 | 66150674 | 66151720 | ENSMUSG00000070780 | Rbm47 | RNA binding motif protein 47 |
| chr5 | 66617923 | 66619199 | ENSMUSG00000029207 | Apbb2 | amyloid beta (A4) precursor protein-binding, family B, member 2 |
| chr5 | 66620161 | 66620628 | ENSMUSG00000029207 | Apbb2 | amyloid beta (A4) precursor protein-binding, family B, member 2 |
| chr5 | 67306773 | 67307524 | ENSMUSG00000029221 | Slc30a9 | solute carrier family 30 (zinc transporter), member 9 |
| chr5 | 67847027 | 67847915 | ENSMUSG00000037685 | Atp8a1 | ATPase, aminophospholipid transporter (APLT), class I, type 8A, member 1 |
| chr5 | 72913802 | 72915175 | ENSMUSG00000036087 | Slain2 | SLAIN motif family, member 2 |
| chr5 | 73255392 | 73257070 | ENSMUSG00000070733 | Fryl | FRY like transcription coactivator |
| chr5 | 74068016 | 74069067 | ENSMUSG00000054814 | Usp46 | ubiquitin specific peptidase 46 |
| chr5 | 75044331 | 75044887 | ENSMUSG00000029229 | Chic2 | cysteine-rich hydrophobic domain 2 |
| chr5 | 76140125 | 76140794 | ENSMUSG00000029233 | Srd5a3 | steroid 5 alpha-reductase 3 |
| chr5 | 76183557 | 76184658 | ENSMUSG00000029234 | Tmem165 | transmembrane protein 165 |
| chr5 | 76303616 | 76305352 | ENSMUSG00000029238 | Clock | circadian locomotor output cycles kaput |
| chr5 | 76588573 | 76589294 | ENSMUSG00000036403 | Cep135 | centrosomal protein 135 |
| chr5 | 76905250 | 76905761 | ENSMUSG00000055923 | Aasdh | aminoadipate-semialdehyde dehydrogenase |
| chr5 | 77265205 | 77266898 | ENSMUSG00000029249 | Rest | RE1-silencing transcription factor |
| chr5 | 77309059 | 77310947 | ENSMUSG00000029250 | Polr2b | polymerase (RNA) II (DNA directed) polypeptide B |
| chr5 | 77407352 | 77408331 | ENSMUSG00000036256 | Igfbp7 | insulin-like growth factor binding protein 7 |
| chr5 | 86172220 | 86172932 | ENSMUSG00000035898 | Uba6 | ubiquitin-like modifier activating enzyme 6 |
| chr5 | 86804038 | 86804908 | ENSMUSG00000035851 | Ythdc1 | YTH domain containing 1 |
| chr5 | 88564634 | 88566019 | ENSMUSG00000029291 | Rufy3 | RUN and FYVE domain containing 3 |
| chr5 | 88675051 | 88676432 | ENSMUSG00000044221 | Grsf1 | G-rich RNA sequence binding factor 1 |
| chr5 | 88764915 | 88765565 | ENSMUSG00000029366 | Dck | deoxycytidine kinase |
| chr5 | 88886503 | 88888067 | ENSMUSG00000060961 | Slc4a4 | solute carrier family 4 (anion exchanger), member 4 |
| chr5 | 90223543 | 90224319 | ENSMUSG00000035505 | Cox18 | cytochrome c oxidase assembly protein 18 |
| chr5 | 90365276 | 90367031 | ENSMUSG00000054945 | Gm9958 | predicted gene 9958 |
| chr5 | 90930983 | 90931904 | ENSMUSG00000029376 | Mthfd2l | methylenetetrahydrofolate dehydrogenase (NADP+ dependent) 2-like |
| chr5 | 92082857 | 92084057 | ENSMUSG00000029405 | G3bp2 | GTPase activating protein (SH3 domain) binding protein 2 |
| chr5 | 92137589 | 92138570 | ENSMUSG00000029407 | Uso1 | USO1 vesicle docking factor |
| chr5 | 92434472 | 92435212 | ENSMUSG00000034826 | Nup54 | nucleoporin 54 |
| chr5 | 92505182 | 92505862 | ENSMUSG00000029426 | Scarb2 | scavenger receptor class B, member 2 |
| chr5 | 92897652 | 92898516 | ENSMUSG00000029381 | Shroom3 | shroom family member 3 |
| chr5 | 93044884 | 93045502 | ENSMUSG00000045314 | Sowahb | sosondowah ankyrin repeat domain family member B |
| chr5 | 93092791 | 93094611 | ENSMUSG00000058013 | Septin11 | septin 11 |
| chr5 | 96997157 | 96998426 | ENSMUSG00000034663 | Bmp2k | BMP2 inducible kinase |
| chr5 | 98030420 | 98031409 | ENSMUSG00000029338 | Antxr2 | anthrax toxin receptor 2 |
| chr5 | 98185248 | 98186239 |  | A730035I17Rik | RIKEN cDNA A730035I17 gene |
| chr5 | 99252327 | 99253300 | ENSMUSG00000089809 | Rasgef1b | RasGEF domain family, member 1B |
| chr5 | 99978011 | 99979570 | ENSMUSG00000000568 | Hnrnpd | heterogeneous nuclear ribonucleoprotein D |
| chr5 | 100038304 | 100040212 | ENSMUSG00000029328 | Hnrnpdl | heterogeneous nuclear ribonucleoprotein D-like |
| chr5 | 100040212 | 100040692 | ENSMUSG00000029326 | Enoph1 | enolase-phosphatase 1 |
| chr5 | 100415863 | 100416627 | ENSMUSG00000035325 | Sec31a | Sec31 homolog A (S. cerevisiae) |
| chr5 | 100673877 | 100674491 | ENSMUSG00000029319 | Coq2 | coenzyme Q2 4-hydroxybenzoate polyprenyltransferase |
| chr5 | 100820415 | 100821116 | ENSMUSG00000035234 | Abraxas1 | BRCA1 A complex subunit |
| chr5 | 100845778 | 100846878 | ENSMUSG00000029314 | Gpat3 | glycerol-3-phosphate acyltransferase 3 |
| chr5 | 101764687 | 101765500 | ENSMUSG00000029330 | Cds1 | CDP-diacylglycerol synthase 1 |
| chr5 | 102068994 | 102070340 | ENSMUSG00000043940 | Wdfy3 | WD repeat and FYVE domain containing 3 |
| chr5 | 103424805 | 103425896 | ENSMUSG00000034573 | Ptpn13 | protein tyrosine phosphatase, non-receptor type 13 |
| chr5 | 103692091 | 103692941 | ENSMUSG00000029313 | Aff1 | AF4/FMR2 family, member 1 |
| chr5 | 103753610 | 103754421 | ENSMUSG00000029313 | Aff1 | AF4/FMR2 family, member 1 |
| chr5 | 103910669 | 103911405 | ENSMUSG00000029312 | Klhl8 | kelch-like 8 |
| chr5 | 104046414 | 104047257 | ENSMUSG00000029310 | Nudt9 | nudix (nucleoside diphosphate linked moiety X)-type motif 9 |
| chr5 | 104459440 | 104460426 | ENSMUSG00000034462 | Pkd2 | polycystin 2, transient receptor potential cation channel |
| chr5 | 105415194 | 105416167 | ENSMUSG00000070639 | Lrrc8b | leucine rich repeat containing 8 family, member B |
| chr5 | 105519094 | 105519867 | ENSMUSG00000054720 | Lrrc8c | leucine rich repeat containing 8 family, member C |
| chr5 | 105699499 | 105701235 | ENSMUSG00000046079 | Lrrc8d | leucine rich repeat containing 8D |
| chr5 | 105731776 | 105732698 | ENSMUSG00000046079 | Lrrc8d | leucine rich repeat containing 8D |
| chr5 | 105876265 | 105877322 | ENSMUSG00000029290 | Zfp326 | zinc finger protein 326 |
| chr5 | 106695679 | 106696966 | ENSMUSG00000049606 | Zfp644 | zinc finger protein 644 |
| chr5 | 107288697 | 107289779 | ENSMUSG00000029287 | Tgfbr3 | transforming growth factor, beta receptor III |
| chr5 | 107597259 | 107598101 | ENSMUSG00000029276 | Glmn | glomulin, FKBP associated protein |
| chr5 | 107868280 | 107869405 | ENSMUSG00000011831 | Evi5 | ecotropic viral integration site 5 |
| chr5 | 107986561 | 107987529 | ENSMUSG00000029270 | Dipk1a | divergent protein kinase domain 1A |
| chr5 | 108132119 | 108133128 | ENSMUSG00000063406 | Tmed5 | transmembrane p24 trafficking protein 5 |
| chr5 | 108268582 | 108269395 | ENSMUSG00000029265 | Dr1 | down-regulator of transcription 1 |
| chr5 | 108629203 | 108630134 | ENSMUSG00000062234 | Gak | cyclin G associated kinase |
| chr5 | 108660191 | 108660839 | ENSMUSG00000004815 | Dgkq | diacylglycerol kinase, theta |
| chr5 | 110106918 | 110108336 | ENSMUSG00000033434 | Gtpbp6 | GTP binding protein 6 (putative) |
| chr5 | 110230666 | 110231584 | ENSMUSG00000029501 | Ankle2 | ankyrin repeat and LEM domain containing 2 |
| chr5 | 110269435 | 110270006 | ENSMUSG00000029500 | Pgam5 | phosphoglycerate mutase family member 5 |
| chr5 | 110285765 | 110286679 | ENSMUSG00000007080 | Pole | polymerase (DNA directed), epsilon |
| chr5 | 110769948 | 110771136 | ENSMUSG00000086401 | Gm15559 | predicted gene 15559 |
| chr5 | 110839355 | 110840074 | ENSMUSG00000043510 | Hscb | HscB iron-sulfur cluster co-chaperone |
| chr5 | 111330386 | 111331308 | ENSMUSG00000050017 | Pitpnb | phosphatidylinositol transfer protein, beta |
| chr5 | 112276266 | 112277110 | ENSMUSG00000029344 | Tpst2 | protein-tyrosine sulfotransferase 2 |
| chr5 | 113137649 | 113139078 | ENSMUSG00000051339 | 2900026A02Rik | RIKEN cDNA 2900026A02 gene |
| chr5 | 113220533 | 113221290 | ENSMUSG00000051339 | 2900026A02Rik | RIKEN cDNA 2900026A02 gene |
| chr5 | 113771145 | 113771989 | ENSMUSG00000018974 | Sart3 | squamous cell carcinoma antigen recognized by T cells 3 |
| chr5 | 113772577 | 113773119 | ENSMUSG00000025825 | Iscu | iron-sulfur cluster assembly enzyme |
| chr5 | 113907945 | 113908867 | ENSMUSG00000004530 | Coro1c | coronin, actin binding protein 1C |
| chr5 | 113993139 | 113994121 | ENSMUSG00000042121 | Ssh1 | slingshot protein phosphatase 1 |
| chr5 | 114130288 | 114131682 | ENSMUSG00000029591 | Ung | uracil DNA glycosylase |
| chr5 | 114567143 | 114568145 | ENSMUSG00000041930 | Fam222a | family with sequence similarity 222, member A |
| chr5 | 114689768 | 114690662 | ENSMUSG00000011884 | Gltp | glycolipid transfer protein |
| chr5 | 114772855 | 114773620 | ENSMUSG00000041890 | Git2 | GIT ArfGAP 2 |
| chr5 | 114774393 | 114775424 | ENSMUSG00000092183 | 4930515G01Rik | RIKEN cDNA 4930515G01 gene |
| chr5 | 115010765 | 115012255 | ENSMUSG00000029550 | Sppl3 | signal peptide peptidase 3 |
| chr5 | 115119115 | 115119519 | ENSMUSG00000029545 | Acads | acyl-Coenzyme A dehydrogenase, short chain |
| chr5 | 115237762 | 115238513 | ENSMUSG00000060152 | Pop5 | processing of precursor 5, ribonuclease P/MRP family (S. cerevisiae) |
| chr5 | 115300480 | 115301355 |  | Gm13830 | predicted gene 13830 |
| chr5 | 115326825 | 115327944 | ENSMUSG00000029538 | Srsf9 | serine and arginine-rich splicing factor 9 |
| chr5 | 115491912 | 115492331 | ENSMUSG00000029524 | Sirt4 | sirtuin 4 |
| chr5 | 115565035 | 115565714 | ENSMUSG00000041638 | Gcn1 | GCN1 activator of EIF2AK4 |
| chr5 | 115631276 | 115632559 | ENSMUSG00000107121 | 1110006O24Rik | RIKEN cDNA 1110006O24 gene |
| chr5 | 115731154 | 115732823 | ENSMUSG00000041609 | Bicdl1 | BICD family like cargo adaptor 1 |
| chr5 | 117287088 | 117287790 | ENSMUSG00000032959 | Pebp1 | phosphatidylethanolamine binding protein 1 |
| chr5 | 117318927 | 117319728 | ENSMUSG00000066894 | Vsig10 | V-set and immunoglobulin domain containing 10 |
| chr5 | 117357179 | 117358173 | ENSMUSG00000029364 | Wsb2 | WD repeat and SOCS box-containing 2 |
| chr5 | 117363430 | 117363824 | ENSMUSG00000029364 | Wsb2 | WD repeat and SOCS box-containing 2 |
| chr5 | 117413616 | 117414329 | ENSMUSG00000061578 | Ksr2 | kinase suppressor of ras 2 |
| chr5 | 117976563 | 117977394 | ENSMUSG00000032898 | Fbxo21 | F-box protein 21 |
| chr5 | 118559803 | 118561445 | ENSMUSG00000018076 | Med13l | mediator complex subunit 13-like |
| chr5 | 120503100 | 120503729 | ENSMUSG00000029598 | Plbd2 | phospholipase B domain containing 2 |
| chr5 | 121190837 | 121191568 | ENSMUSG00000043733 | Ptpn11 | protein tyrosine phosphatase, non-receptor type 11 |
| chr5 | 121220126 | 121221006 | ENSMUSG00000042744 | Hectd4 | HECT domain E3 ubiquitin protein ligase 4 |
| chr5 | 121451930 | 121453015 | ENSMUSG00000029616 | Erp29 | endoplasmic reticulum protein 29 |
| chr5 | 121545049 | 121546262 | ENSMUSG00000029454 | Mapkapk5 | MAP kinase-activated protein kinase 5 |
| chr5 | 121660329 | 121661089 | ENSMUSG00000029456 | Acad10 | acyl-Coenzyme A dehydrogenase family, member 10 |
| chr5 | 122049676 | 122050625 | ENSMUSG00000042589 | Cux2 | cut-like homeobox 2 |
| chr5 | 122158080 | 122159120 | ENSMUSG00000004455 | Ppp1cc | protein phosphatase 1 catalytic subunit gamma |
| chr5 | 122284333 | 122285461 | ENSMUSG00000038582 | Pptc7 | PTC7 protein phosphatase homolog |
| chr5 | 122354016 | 122354940 | ENSMUSG00000038569 | Rad9b | RAD9 checkpoint clamp component B |
| chr5 | 122371839 | 122372860 | ENSMUSG00000029464 | Gpn3 | GPN-loop GTPase 3 |
| chr5 | 122501079 | 122502808 | ENSMUSG00000029467 | Atp2a2 | ATPase, Ca++ transporting, cardiac muscle, slow twitch 2 |
| chr5 | 122707357 | 122707967 | ENSMUSG00000029470 | P2rx4 | purinergic receptor P2X, ligand-gated ion channel 4 |
| chr5 | 122778590 | 122779718 | ENSMUSG00000029471 | Camkk2 | calcium/calmodulin-dependent protein kinase kinase 2, beta |
| chr5 | 122820698 | 122821520 | ENSMUSG00000029472 | Anapc5 | anaphase-promoting complex subunit 5 |
| chr5 | 122899477 | 122900788 | ENSMUSG00000029475 | Kdm2b | lysine (K)-specific demethylase 2B |
| chr5 | 123014768 | 123015887 | ENSMUSG00000049686 | Orai1 | ORAI calcium release-activated calcium modulator 1 |
| chr5 | 123057767 | 123058250 | ENSMUSG00000054434 | Tmem120b | transmembrane protein 120B |
| chr5 | 123076046 | 123076630 | ENSMUSG00000054434 | Tmem120b | transmembrane protein 120B |
| chr5 | 123394392 | 123395395 | ENSMUSG00000038342 | Mlxip | MLX interacting protein |
| chr5 | 123523637 | 123524608 | ENSMUSG00000029433 | Diablo | diablo, IAP-binding mitochondrial protein |
| chr5 | 123683466 | 123684678 | ENSMUSG00000049550 | Clip1 | CAP-GLY domain containing linker protein 1 |
| chr5 | 123973337 | 123974448 | ENSMUSG00000000915 | Hip1r | huntingtin interacting protein 1 related |
| chr5 | 124006734 | 124007704 | ENSMUSG00000066278 | Vps37b | vacuolar protein sorting 37B |
| chr5 | 124111750 | 124113218 | ENSMUSG00000023707 | Ogfod2 | 2-oxoglutarate and iron-dependent oxygenase domain containing 2 |
| chr5 | 124249099 | 124250376 | ENSMUSG00000029406 | Pitpnm2 | phosphatidylinositol transfer protein, membrane-associated 2 |
| chr5 | 124327479 | 124328240 | ENSMUSG00000038126 | Mphosph9 | M-phase phosphoprotein 9 |
| chr5 | 124353376 | 124354927 | ENSMUSG00000029394 | Cdk2ap1 | CDK2 (cyclin-dependent kinase 2)-associated protein 1 |
| chr5 | 124424665 | 124425737 | ENSMUSG00000038095 | Sbno1 | strawberry notch 1 |
| chr5 | 124483147 | 124483708 | ENSMUSG00000029402 | Snrnp35 | small nuclear ribonucleoprotein 35 (U11/U12) |
| chr5 | 124540452 | 124541590 | ENSMUSG00000029390 | Tmed2 | transmembrane p24 trafficking protein 2 |
| chr5 | 124552513 | 124553249 | ENSMUSG00000029389 | Ddx55 | DEAD box helicase 55 |
| chr5 | 125140785 | 125141720 | ENSMUSG00000029478 | Ncor2 | nuclear receptor co-repressor 2 |
| chr5 | 125177622 | 125179423 | ENSMUSG00000029478 | Ncor2 | nuclear receptor co-repressor 2 |
| chr5 | 125433650 | 125434199 | ENSMUSG00000029480 | Dhx37 | DEAH (Asp-Glu-Ala-His) box polypeptide 37 |
| chr5 | 125441291 | 125442149 | ENSMUSG00000037905 | Bri3bp | Bri3 binding protein |
| chr5 | 125465627 | 125466119 | ENSMUSG00000029482 | Aacs | acetoacetyl-CoA synthetase |
| chr5 | 125475722 | 125476214 | ENSMUSG00000029482 | Aacs | acetoacetyl-CoA synthetase |
| chr5 | 127616136 | 127617610 | ENSMUSG00000029416 | Slc15a4 | solute carrier family 15, member 4 |
| chr5 | 129008122 | 129008789 | ENSMUSG00000029428 | Stx2 | syntaxin 2 |
| chr5 | 129019556 | 129021255 | ENSMUSG00000029430 | Ran | RAN, member RAS oncogene family |
| chr5 | 129724950 | 129725580 | ENSMUSG00000029432 | Nipsnap2 | nipsnap homolog 2 |
| chr5 | 129908391 | 129909103 | ENSMUSG00000095789 | Nupr1l | nuclear protein transcriptional regulator 1 like |
| chr5 | 129941541 | 129942656 | ENSMUSG00000066735 | Vkorc1l1 | vitamin K epoxide reductase complex, subunit 1-like 1 |
| chr5 | 130079243 | 130079897 | ENSMUSG00000034118 | Tpst1 | protein-tyrosine sulfotransferase 1 |
| chr5 | 130144669 | 130145340 | ENSMUSG00000034110 | Kctd7 | potassium channel tetramerisation domain containing 7 |
| chr5 | 130171460 | 130172650 | ENSMUSG00000025340 | Rabgef1 | RAB guanine nucleotide exchange factor (GEF) 1 |
| chr5 | 130222142 | 130222816 | ENSMUSG00000053094 | Tmem248 | transmembrane protein 248 |
| chr5 | 134183852 | 134184628 | ENSMUSG00000015942 | Gtf2ird2 | GTF2I repeat domain containing 2 |
| chr5 | 134313486 | 134315058 | ENSMUSG00000060261 | Gtf2i | general transcription factor II I |
| chr5 | 134638521 | 134639749 | ENSMUSG00000040731 | Eif4h | eukaryotic translation initiation factor 4H |
| chr5 | 134688384 | 134689143 | ENSMUSG00000029674 | Limk1 | LIM-domain containing, protein kinase |
| chr5 | 135168127 | 135168659 | ENSMUSG00000029681 | Bcl7b | B cell CLL/lymphoma 7B |
| chr5 | 135186666 | 135188319 | ENSMUSG00000002748 | Baz1b | bromodomain adjacent to zinc finger domain, 1B |
| chr5 | 135393825 | 135394860 | ENSMUSG00000053293 | Pom121 | nuclear pore membrane protein 121 |
| chr5 | 135688506 | 135689712 | ENSMUSG00000005514 | Por | P450 (cytochrome) oxidoreductase |
| chr5 | 135743631 | 135744403 | ENSMUSG00000039886 | Tmem120a | transmembrane protein 120A |
| chr5 | 135962549 | 135963378 | ENSMUSG00000029699 | Ssc4d | scavenger receptor cysteine rich family, 4 domains |
| chr5 | 137031361 | 137033039 | ENSMUSG00000037428 | Vgf | VGF nerve growth factor inducible |
| chr5 | 137741385 | 137741909 | ENSMUSG00000045348 | Nyap1 | neuronal tyrosine-phosphorylated phosphoinositide 3-kinase adaptor 1 |
| chr5 | 137778855 | 137779865 | ENSMUSG00000029725 | Ppp1r35 | protein phosphatase 1, regulatory subunit 35 |
| chr5 | 139149387 | 139150985 | ENSMUSG00000025855 | Prkar1b | protein kinase, cAMP dependent regulatory, type I beta |
| chr5 | 139200252 | 139201293 | ENSMUSG00000036817 | Sun1 | Sad1 and UNC84 domain containing 1 |
| chr5 | 139252134 | 139253823 | ENSMUSG00000025858 | Get4 | golgi to ER traffic protein 4 |
| chr5 | 139324775 | 139325786 | ENSMUSG00000056413 | Adap1 | ArfGAP with dual PH domains 1 |
| chr5 | 139371979 | 139372460 | ENSMUSG00000065593 | Mir339 | microRNA 339 |
| chr5 | 139380269 | 139381043 | ENSMUSG00000044197 | Gpr146 | G protein-coupled receptor 146 |
| chr5 | 139460353 | 139460784 | ENSMUSG00000053553 | 3110082I17Rik | RIKEN cDNA 3110082I17 gene |
| chr5 | 140388681 | 140389548 | ENSMUSG00000029560 | Snx8 | sorting nexin 8 |
| chr5 | 140419220 | 140420074 | ENSMUSG00000056076 | Eif3b | eukaryotic translation initiation factor 3, subunit B |
| chr5 | 140607128 | 140608143 | ENSMUSG00000029570 | Lfng | LFNG O-fucosylpeptide 3-beta-N-acetylglucosaminyltransferase |
| chr5 | 140610376 | 140611716 | ENSMUSG00000029570 | Lfng | LFNG O-fucosylpeptide 3-beta-N-acetylglucosaminyltransferase |
| chr5 | 140701872 | 140702497 | ENSMUSG00000036555 | Iqce | IQ motif containing E |
| chr5 | 140829829 | 140830660 | ENSMUSG00000000149 | Gna12 | guanine nucleotide binding protein, alpha 12 |
| chr5 | 142401373 | 142402152 | ENSMUSG00000056493 | Foxk1 | forkhead box K1 |
| chr5 | 142905093 | 142906943 | ENSMUSG00000029580 | Actb | actin, beta |
| chr5 | 143112221 | 143113180 | ENSMUSG00000045078 | Rnf216 | ring finger protein 216 |
| chr5 | 143314226 | 143315510 | ENSMUSG00000039244 | E130309D02Rik | RIKEN cDNA E130309D02 gene |
| chr5 | 143403598 | 143404541 | ENSMUSG00000079111 | Kdelr2 | KDEL (Lys-Asp-Glu-Leu) endoplasmic reticulum protein retention receptor 2 |
| chr5 | 143527344 | 143528317 | ENSMUSG00000001847 | Rac1 | Rac family small GTPase 1 |
| chr5 | 143548465 | 143549185 | ENSMUSG00000083012 | Fam220a | family with sequence similarity 220, member A |
| chr5 | 143622266 | 143623061 | ENSMUSG00000018001 | Cyth3 | cytohesin 3 |
| chr5 | 143731433 | 143732861 | ENSMUSG00000051306 | Usp42 | ubiquitin specific peptidase 42 |
| chr5 | 143817701 | 143818269 | ENSMUSG00000029613 | Eif2ak1 | eukaryotic translation initiation factor 2 alpha kinase 1 |
| chr5 | 144100301 | 144101186 | ENSMUSG00000038970 | Lmtk2 | lemur tyrosine kinase 2 |
| chr5 | 144255124 | 144256159 | ENSMUSG00000047843 | Bri3 | brain protein I3 |
| chr5 | 144256159 | 144256737 | ENSMUSG00000047843 | Bri3 | brain protein I3 |
| chr5 | 144357537 | 144358722 | ENSMUSG00000104600 | Dmrt1i | Dmrt1 interacting ncRNA |
| chr5 | 144767878 | 144768834 | ENSMUSG00000045482 | Trrap | transformation/transcription domain-associated protein |
| chr5 | 144965453 | 144966164 | ENSMUSG00000038780 | Smurf1 | SMAD specific E3 ubiquitin protein ligase 1 |
| chr5 | 145231367 | 145232422 | ENSMUSG00000007812 | Zfp655 | zinc finger protein 655 |
| chr5 | 146230899 | 146232047 | ENSMUSG00000029635 | Cdk8 | cyclin-dependent kinase 8 |
| chr5 | 146793946 | 146795272 | ENSMUSG00000029640 | Usp12 | ubiquitin specific peptidase 12 |
| chr5 | 146832934 | 146833587 | ENSMUSG00000041453 | Rpl21 | ribosomal protein L21 |
| chr5 | 146844984 | 146845339 | ENSMUSG00000029641 | Rasl11a | RAS-like, family 11, member A |
| chr5 | 146948637 | 146949097 | ENSMUSG00000016503 | Gtf3a | general transcription factor III A |
| chr5 | 147075981 | 147077947 | ENSMUSG00000016520 | Lnx2 | ligand of numb-protein X 2 |
| chr5 | 147429820 | 147431813 | ENSMUSG00000029647 | Pan3 | PAN3 poly(A) specific ribonuclease subunit |
| chr5 | 148398816 | 148400177 | ENSMUSG00000041313 | Slc7a1 | solute carrier family 7 (cationic amino acid transporter, y+ system), member 1 |
| chr5 | 148551368 | 148552139 | ENSMUSG00000001687 | Ubl3 | ubiquitin-like 3 |
| chr5 | 148552139 | 148552887 | ENSMUSG00000001687 | Ubl3 | ubiquitin-like 3 |
| chr5 | 149678094 | 149678846 | ENSMUSG00000051950 | B3glct | beta-3-glucosyltransferase |
| chr5 | 150522457 | 150522948 | ENSMUSG00000041147 | Brca2 | breast cancer 2, early onset |
| chr5 | 150673409 | 150674638 | ENSMUSG00000034021 | Pds5b | PDS5 cohesin associated factor B |
| chr5 | 151233365 | 151233992 | ENSMUSG00000016128 | Stard13 | StAR-related lipid transfer (START) domain containing 13 |
| chr5 | 151368323 | 151368932 | ENSMUSG00000097321 | 1700028E10Rik | RIKEN cDNA 1700028E10 gene |
| chr6 | 4902980 | 4903896 | ENSMUSG00000032827 | Ppp1r9a | protein phosphatase 1, regulatory subunit 9A |
| chr6 | 5298138 | 5298467 | ENSMUSG00000032667 | Pon2 | paraoxonase 2 |
| chr6 | 7692622 | 7693390 | ENSMUSG00000029752 | Asns | asparagine synthetase |
| chr6 | 7844613 | 7845456 | ENSMUSG00000042460 | C1galt1 | core 1 synthase, glycoprotein-N-acetylgalactosamine 3-beta-galactosyltransferase, 1 |
| chr6 | 8209060 | 8209923 | ENSMUSG00000042447 | Mios | meiosis regulator for oocyte development |
| chr6 | 8509017 | 8510439 | ENSMUSG00000029638 | Glcci1 | glucocorticoid induced transcript 1 |
| chr6 | 11907149 | 11908130 | ENSMUSG00000029632 | Ndufa4 | Ndufa4, mitochondrial complex associated |
| chr6 | 13677075 | 13678190 | ENSMUSG00000042742 | Bmt2 | base methyltransferase of 25S rRNA 2 |
| chr6 | 17463285 | 17464714 | ENSMUSG00000009376 | Met | met proto-oncogene |
| chr6 | 17636776 | 17637642 | ENSMUSG00000015733 | Capza2 | capping protein (actin filament) muscle Z-line, alpha 2 |
| chr6 | 17693860 | 17694912 | ENSMUSG00000029534 | St7 | suppression of tumorigenicity 7 |
| chr6 | 21851419 | 21852182 | ENSMUSG00000029669 | Tspan12 | tetraspanin 12 |
| chr6 | 22355590 | 22356471 | ENSMUSG00000029672 | Fam3c | family with sequence similarity 3, member C |
| chr6 | 24664103 | 24665500 | ENSMUSG00000029684 | Wasl | WASP like actin nucleation promoting factor |
| chr6 | 28260801 | 28262599 | ENSMUSG00000039841 | Zfp800 | zinc finger protein 800 |
| chr6 | 28479942 | 28480797 | ENSMUSG00000001424 | Snd1 | staphylococcal nuclease and tudor domain containing 1 |
| chr6 | 29211856 | 29212460 | ENSMUSG00000003500 | Impdh1 | inosine monophosphate dehydrogenase 1 |
| chr6 | 29272327 | 29272915 | ENSMUSG00000043421 | Hilpda | hypoxia inducible lipid droplet associated |
| chr6 | 29735426 | 29736649 | ENSMUSG00000001761 | Smo | smoothened, frizzled class receptor |
| chr6 | 29768206 | 29769227 | ENSMUSG00000029772 | Ahcyl2 | S-adenosylhomocysteine hydrolase-like 2 |
| chr6 | 30047620 | 30048891 | ENSMUSG00000058440 | Nrf1 | nuclear respiratory factor 1 |
| chr6 | 30303801 | 30304578 | ENSMUSG00000039159 | Ube2h | ubiquitin-conjugating enzyme E2H |
| chr6 | 31216301 | 31217027 | ENSMUSG00000044471 | Lncpint | long non-protein coding RNA, Trp53 induced transcript |
| chr6 | 31398608 | 31399355 | ENSMUSG00000086212 | Mkln1os | muskelin 1, intracellular mediator containing kelch motifs, opposite strand |
| chr6 | 31563452 | 31564286 | ENSMUSG00000025608 | Podxl | podocalyxin-like |
| chr6 | 34780361 | 34781028 | ENSMUSG00000038836 | Agbl3 | ATP/GTP binding protein-like 3 |
| chr6 | 35177399 | 35178231 | ENSMUSG00000038759 | Nup205 | nucleoporin 205 |
| chr6 | 37870545 | 37871731 | ENSMUSG00000029833 | Trim24 | tripartite motif-containing 24 |
| chr6 | 38253727 | 38254635 | ENSMUSG00000063455 | D630045J12Rik | RIKEN cDNA D630045J12 gene |
| chr6 | 38298709 | 38299365 | ENSMUSG00000047749 | Zc3hav1l | zinc finger CCCH-type, antiviral 1-like |
| chr6 | 38433265 | 38434981 | ENSMUSG00000038538 | Ubn2 | ubinuclein 2 |
| chr6 | 38635996 | 38637410 | ENSMUSG00000071537 | Klrg2 | killer cell lectin-like receptor subfamily G, member 2 |
| chr6 | 38875713 | 38876840 | ENSMUSG00000061436 | Hipk2 | homeodomain interacting protein kinase 2 |
| chr6 | 39117763 | 39118590 | ENSMUSG00000038507 | Parp12 | poly (ADP-ribose) polymerase family, member 12 |
| chr6 | 40471328 | 40471824 | ENSMUSG00000029911 | Ssbp1 | single-stranded DNA binding protein 1 |
| chr6 | 42264764 | 42265423 | ENSMUSG00000029863 | Casp2 | caspase 2 |
| chr6 | 42349397 | 42350459 | ENSMUSG00000029860 | Zyx | zyxin |
| chr6 | 47453799 | 47455311 | ENSMUSG00000029686 | Cul1 | cullin 1 |
| chr6 | 47594084 | 47595728 | ENSMUSG00000029687 | Ezh2 | enhancer of zeste 2 polycomb repressive complex 2 subunit |
| chr6 | 47835540 | 47836560 | ENSMUSG00000062519 | Zfp398 | zinc finger protein 398 |
| chr6 | 48047276 | 48049127 | ENSMUSG00000071477 | Zfp777 | zinc finger protein 777 |
| chr6 | 48085838 | 48086824 | ENSMUSG00000057691 | Zfp746 | zinc finger protein 746 |
| chr6 | 50109924 | 50111025 | ENSMUSG00000038388 | Mpp6 | membrane protein, palmitoylated 6 (MAGUK p55 subfamily member 6) |
| chr6 | 51523832 | 51524234 | ENSMUSG00000038301 | Snx10 | sorting nexin 10 |
| chr6 | 54039385 | 54040498 | ENSMUSG00000004633 | Chn2 | chimerin 2 |
| chr6 | 54594845 | 54595701 | ENSMUSG00000005225 | Plekha8 | pleckstrin homology domain containing, family A (phosphoinositide binding specific) member 8 |
| chr6 | 54816468 | 54817738 | ENSMUSG00000058446 | Znrf2 | zinc and ring finger 2 |
| chr6 | 55037692 | 55038467 | ENSMUSG00000029777 | Gars | glycyl-tRNA synthetase |
| chr6 | 56714730 | 56715481 | ENSMUSG00000029787 | Avl9 | AVL9 cell migration associated |
| chr6 | 56796871 | 56798267 | ENSMUSG00000059486 | Kbtbd2 | kelch repeat and BTB (POZ) domain containing 2 |
| chr6 | 56831891 | 56832477 | ENSMUSG00000029781 | Fkbp9 | FK506 binding protein 9 |
| chr6 | 57824557 | 57825216 | ENSMUSG00000037788 | Vopp1 | vesicular, overexpressed in cancer, prosurvival protein 1 |
| chr6 | 58831329 | 58832069 | ENSMUSG00000029804 | Herc3 | hect domain and RLD 3 |
| chr6 | 61180038 | 61181120 |  | A730020E08Rik | RIKEN cDNA A730020E08 gene |
| chr6 | 65042515 | 65043181 | ENSMUSG00000029920 | Smarcad1 | SWI/SNF-related, matrix-associated actin-dependent regulator of chromatin, subfamily a, containing DEAD/H box 1 |
| chr6 | 66896180 | 66896952 | ENSMUSG00000036402 | Gng12 | guanine nucleotide binding protein (G protein), gamma 12 |
| chr6 | 67036408 | 67037586 | ENSMUSG00000036390 | Gadd45a | growth arrest and DNA-damage-inducible 45 alpha |
| chr6 | 67266547 | 67268275 | ENSMUSG00000036371 | Serbp1 | serpine1 mRNA binding protein 1 |
| chr6 | 70791717 | 70792293 | ENSMUSG00000053604 | Rpia | ribose 5-phosphate isomerase A |
| chr6 | 70844366 | 70845306 | ENSMUSG00000031668 | Eif2ak3 | eukaryotic translation initiation factor 2 alpha kinase 3 |
| chr6 | 71271419 | 71272533 | ENSMUSG00000098623 | Mir8112 | microRNA 8112 |
| chr6 | 71439769 | 71441448 | ENSMUSG00000002222 | Rmnd5a | required for meiotic nuclear division 5 homolog A |
| chr6 | 71493738 | 71494886 | ENSMUSG00000052656 | Rnf103 | ring finger protein 103 |
| chr6 | 72344723 | 72345232 | ENSMUSG00000056305 | Usp39 | ubiquitin specific peptidase 39 |
| chr6 | 72347235 | 72347913 | ENSMUSG00000058706 | 0610030E20Rik | RIKEN cDNA 0610030E20 gene |
| chr6 | 72438732 | 72440110 | ENSMUSG00000053907 | Mat2a | methionine adenosyltransferase II, alpha |
| chr6 | 72898651 | 72900552 | ENSMUSG00000055239 | Kcmf1 | potassium channel modulatory factor 1 |
| chr6 | 83101468 | 83102659 | ENSMUSG00000107499 | Ccdc142 | coiled-coil domain containing 142 |
| chr6 | 83115279 | 83116166 | ENSMUSG00000030036 | Mogs | mannosyl-oligosaccharide glucosidase |
| chr6 | 83317023 | 83317715 | ENSMUSG00000005667 | Mthfd2 | methylenetetrahydrofolate dehydrogenase (NAD+ dependent), methenyltetrahydrofolate cyclohydrolase |
| chr6 | 83441312 | 83441958 | ENSMUSG00000034832 | Tet3 | tet methylcytosine dioxygenase 3 |
| chr6 | 83455821 | 83457236 | ENSMUSG00000034832 | Tet3 | tet methylcytosine dioxygenase 3 |
| chr6 | 85374140 | 85375019 | ENSMUSG00000051343 | Rab11fip5 | RAB11 family interacting protein 5 (class I) |
| chr6 | 85450955 | 85452570 | ENSMUSG00000030007 | Cct7 | chaperonin containing Tcp1, subunit 7 (eta) |
| chr6 | 86194927 | 86195876 | ENSMUSG00000029999 | Tgfa | transforming growth factor alpha |
| chr6 | 86471988 | 86472536 | ENSMUSG00000046679 | C87436 | expressed sequence C87436 |
| chr6 | 86732600 | 86733489 | ENSMUSG00000001157 | Gmcl1 | germ cell-less, spermatogenesis associated 1 |
| chr6 | 87042263 | 87043180 | ENSMUSG00000029992 | Gfpt1 | glutamine fructose-6-phosphate transaminase 1 |
| chr6 | 87810987 | 87811961 | ENSMUSG00000030055 | Rab43 | RAB43, member RAS oncogene family |
| chr6 | 87850312 | 87851297 | ENSMUSG00000030057 | Cnbp | cellular nucleic acid binding protein |
| chr6 | 87913888 | 87914384 | ENSMUSG00000030060 | Hmces | 5-hydroxymethylcytosine (hmC) binding, ES cell specific |
| chr6 | 88446045 | 88446723 | ENSMUSG00000033216 | Eefsec | eukaryotic elongation factor, selenocysteine-tRNA-specific |
| chr6 | 89643901 | 89644544 | ENSMUSG00000000811 | Txnrd3 | thioredoxin reductase 3 |
| chr6 | 90369381 | 90370660 | ENSMUSG00000034430 | Zxdc | ZXD family zinc finger C |
| chr6 | 90736122 | 90737156 | ENSMUSG00000034312 | Iqsec1 | IQ motif and Sec7 domain 1 |
| chr6 | 91116287 | 91117141 | ENSMUSG00000030091 | Nup210 | nucleoporin 210 |
| chr6 | 91410825 | 91411648 | ENSMUSG00000030093 | Wnt7a | wingless-type MMTV integration site family, member 7A |
| chr6 | 91683975 | 91685302 | ENSMUSG00000030096 | Slc6a6 | solute carrier family 6 (neurotransmitter transporter, taurine), member 6 |
| chr6 | 92090975 | 92092661 | ENSMUSG00000005893 | Nr2c2 | nuclear receptor subfamily 2, group C, member 2 |
| chr6 | 94282692 | 94284504 | ENSMUSG00000045095 | Magi1 | membrane associated guanylate kinase, WW and PDZ domain containing 1 |
| chr6 | 94698992 | 94700603 | ENSMUSG00000030029 | Lrig1 | leucine-rich repeats and immunoglobulin-like domains 1 |
| chr6 | 95718130 | 95718806 | ENSMUSG00000061838 | Suclg2 | succinate-Coenzyme A ligase, GDP-forming, beta subunit |
| chr6 | 97178599 | 97179488 | ENSMUSG00000030059 | Tmf1 | TATA element modulatory factor 1 |
| chr6 | 99521259 | 99522970 | ENSMUSG00000030067 | Foxp1 | forkhead box P1 |
| chr6 | 100286648 | 100288052 | ENSMUSG00000072872 | Rybp | RING1 and YY1 binding protein |
| chr6 | 100833318 | 100834469 | ENSMUSG00000052144 | Ppp4r2 | protein phosphatase 4, regulatory subunit 2 |
| chr6 | 108659216 | 108660927 | ENSMUSG00000030103 | Bhlhe40 | basic helix-loop-helix family, member e40 |
| chr6 | 108828338 | 108829304 | ENSMUSG00000030104 | Edem1 | ER degradation enhancer, mannosidase alpha-like 1 |
| chr6 | 113076058 | 113077825 | ENSMUSG00000086429 | Gt(ROSA)26Sor | gene trap ROSA 26, Philippe Soriano |
| chr6 | 113623919 | 113624500 | ENSMUSG00000033933 | Vhl | von Hippel-Lindau tumor suppressor |
| chr6 | 113696817 | 113698101 | ENSMUSG00000056952 | Tatdn2 | TatD DNase domain containing 2 |
| chr6 | 114968641 | 114970189 | ENSMUSG00000030315 | Vgll4 | vestigial like family member 4 |
| chr6 | 115675685 | 115676871 | ENSMUSG00000000441 | Raf1 | v-raf-leukemia viral oncogene 1 |
| chr6 | 116192869 | 116193733 | ENSMUSG00000030126 | Tmcc1 | transmembrane and coiled coil domains 1 |
| chr6 | 116337786 | 116338520 | ENSMUSG00000025702 | Marchf8 | membrane associated ring-CH-type finger 8 |
| chr6 | 117879165 | 117880005 | ENSMUSG00000042097 | Zfp239 | zinc finger protein 239 |
| chr6 | 117906747 | 117908037 | ENSMUSG00000042079 | Hnrnpf | heterogeneous nuclear ribonucleoprotein F |
| chr6 | 117916723 | 117917901 | ENSMUSG00000042079 | Hnrnpf | heterogeneous nuclear ribonucleoprotein F |
| chr6 | 118138810 | 118139323 | ENSMUSG00000042042 | Csgalnact2 | chondroitin sulfate N-acetylgalactosaminyltransferase 2 |
| chr6 | 119388194 | 119388907 | ENSMUSG00000030168 | Adipor2 | adiponectin receptor 2 |
| chr6 | 119847853 | 119848685 | ENSMUSG00000030172 | Erc1 | ELKS/RAB6-interacting/CAST family member 1 |
| chr6 | 120036927 | 120038558 | ENSMUSG00000045962 | Wnk1 | WNK lysine deficient protein kinase 1 |
| chr6 | 120463087 | 120463957 | ENSMUSG00000002897 | Il17ra | interleukin 17 receptor A |
| chr6 | 121109714 | 121110298 | ENSMUSG00000051586 | Mical3 | microtubule associated monooxygenase, calponin and LIM domain containing 3 |
| chr6 | 122819705 | 122820656 | ENSMUSG00000003154 | Foxj2 | forkhead box J2 |
| chr6 | 124745421 | 124746114 | ENSMUSG00000004263 | Atn1 | atrophin 1 |
| chr6 | 124917100 | 124918437 | ENSMUSG00000030122 | Ptms | parathymosin |
| chr6 | 125009033 | 125009614 | ENSMUSG00000038346 | Zfp384 | zinc finger protein 384 |
| chr6 | 125313366 | 125314352 | ENSMUSG00000030339 | Ltbr | lymphotoxin B receptor |
| chr6 | 127330163 | 127330605 | ENSMUSG00000107153 | Gm38404 | predicted gene, 38404 |
| chr6 | 128299747 | 128300968 | ENSMUSG00000030353 | Tead4 | TEA domain family member 4 |
| chr6 | 128437467 | 128438962 | ENSMUSG00000030357 | Fkbp4 | FK506 binding protein 4 |
| chr6 | 131387201 | 131388714 | ENSMUSG00000030189 | Ybx3 | Y box protein 3 |
| chr6 | 134034881 | 134036233 | ENSMUSG00000030199 | Etv6 | ets variant 6 |
| chr6 | 134566296 | 134567414 | ENSMUSG00000030201 | Lrp6 | low density lipoprotein receptor-related protein 6 |
| chr6 | 134830034 | 134830642 | ENSMUSG00000032652 | Crebl2 | cAMP responsive element binding protein-like 2 |
| chr6 | 134919803 | 134921443 | ENSMUSG00000003031 | Cdkn1b | cyclin-dependent kinase inhibitor 1B |
| chr6 | 134926700 | 134927574 | ENSMUSG00000098318 | Lockd | lncRNA downstream of Cdkn1b |
| chr6 | 134928152 | 134928848 | ENSMUSG00000098318 | Lockd | lncRNA downstream of Cdkn1b |
| chr6 | 134983557 | 134984029 | ENSMUSG00000090698 | Apold1 | apolipoprotein L domain containing 1 |
| chr6 | 136808170 | 136809043 | ENSMUSG00000060032 | H2aj | H2J.A histone |
| chr6 | 140423708 | 140425078 | ENSMUSG00000030231 | Plekha5 | pleckstrin homology domain containing, family A member 5 |
| chr6 | 140623263 | 140624605 | ENSMUSG00000030232 | Aebp2 | AE binding protein 2 |
| chr6 | 143099333 | 143100310 | ENSMUSG00000030279 | C2cd5 | C2 calcium-dependent domain containing 5 |
| chr6 | 143166997 | 143167891 | ENSMUSG00000030275 | Etnk1 | ethanolamine kinase 1 |
| chr6 | 145249320 | 145250643 | ENSMUSG00000086013 | Gm15706 | predicted gene 15706 |
| chr6 | 145746573 | 145747428 | ENSMUSG00000030259 | Rassf8 | Ras association (RalGDS/AF-6) domain family (N-terminal) member 8 |
| chr6 | 145862794 | 145864099 | ENSMUSG00000030256 | Bhlhe41 | basic helix-loop-helix family, member e41 |
| chr6 | 145865118 | 145866892 | ENSMUSG00000030256 | Bhlhe41 | basic helix-loop-helix family, member e41 |
| chr6 | 146634263 | 146634788 | ENSMUSG00000040234 | Tm7sf3 | transmembrane 7 superfamily member 3 |
| chr6 | 146724806 | 146725571 | ENSMUSG00000001630 | Stk38l | serine/threonine kinase 38 like |
| chr6 | 146888138 | 146888871 | ENSMUSG00000016487 | Ppfibp1 | PTPRF interacting protein, binding protein 1 (liprin beta 1) |
| chr6 | 147091184 | 147092509 | ENSMUSG00000040102 | Klhl42 | kelch-like 42 |
| chr6 | 149100698 | 149101776 | ENSMUSG00000030313 | Dennd5b | DENN/MADD domain containing 5B |
| chr6 | 149408904 | 149409361 | ENSMUSG00000003452 | Bicd1 | BICD cargo adaptor 1 |
| chr7 | 4788896 | 4789714 | ENSMUSG00000030431 | Tmem238 | transmembrane protein 238 |
| chr7 | 7298407 | 7298924 | ENSMUSG00000093599 | Mir5620 | microRNA 5620 |
| chr7 | 13024054 | 13025174 | ENSMUSG00000005566 | Trim28 | tripartite motif-containing 28 |
| chr7 | 16309225 | 16310969 | ENSMUSG00000002083 | Bbc3 | BCL2 binding component 3 |
| chr7 | 16313522 | 16314286 | ENSMUSG00000002083 | Bbc3 | BCL2 binding component 3 |
| chr7 | 16614334 | 16615516 | ENSMUSG00000058230 | Arhgap35 | Rho GTPase activating protein 35 |
| chr7 | 16923743 | 16924349 | ENSMUSG00000019370 | Calm3 | calmodulin 3 |
| chr7 | 19094607 | 19095465 | ENSMUSG00000040841 | Six5 | sine oculis-related homeobox 5 |
| chr7 | 24507122 | 24507727 | ENSMUSG00000064264 | Zfp428 | zinc finger protein 428 |
| chr7 | 25281747 | 25282943 | ENSMUSG00000005442 | Cic | capicua transcriptional repressor |
| chr7 | 28961781 | 28962559 | ENSMUSG00000054808 | Actn4 | actinin alpha 4 |
| chr7 | 29281249 | 29282032 | ENSMUSG00000074227 | Spint2 | serine protease inhibitor, Kunitz type 2 |
| chr7 | 30955779 | 30957698 | ENSMUSG00000058239 | Usf2 | upstream transcription factor 2 |
| chr7 | 34312970 | 34313681 | ENSMUSG00000066571 | Garre1 | granule associated Rac and RHOG effector 1 |
| chr7 | 35555118 | 35555926 | ENSMUSG00000034875 | Nudt19 | nudix (nucleoside diphosphate linked moiety X)-type motif 19 |
| chr7 | 38018848 | 38019636 | ENSMUSG00000030421 | Uri1 | URI1, prefoldin-like chaperone |
| chr7 | 38106901 | 38107935 | ENSMUSG00000002068 | Ccne1 | cyclin E1 |
| chr7 | 45628419 | 45629109 | ENSMUSG00000044562 | Rasip1 | Ras interacting protein 1 |
| chr7 | 45866069 | 45866690 | ENSMUSG00000002771 | Grin2d | glutamate receptor, ionotropic, NMDA2D (epsilon 4) |
| chr7 | 46845481 | 46846417 | ENSMUSG00000063229 | Ldha | lactate dehydrogenase A |
| chr7 | 49245773 | 49247040 | ENSMUSG00000052512 | Nav2 | neuron navigator 2 |
| chr7 | 49778307 | 49778848 | ENSMUSG00000030505 | Prmt3 | protein arginine N-methyltransferase 3 |
| chr7 | 51861681 | 51862367 | ENSMUSG00000092118 | Fancf | Fanconi anemia, complementation group F |
| chr7 | 59228292 | 59229724 | ENSMUSG00000025326 | Ube3a | ubiquitin protein ligase E3A |
| chr7 | 65370366 | 65371370 | ENSMUSG00000030516 | Tjp1 | tight junction protein 1 |
| chr7 | 65644715 | 65645354 | ENSMUSG00000030515 | Tarsl2 | threonyl-tRNA synthetase-like 2 |
| chr7 | 66109027 | 66110319 | ENSMUSG00000032640 | Chsy1 | chondroitin sulfate synthase 1 |
| chr7 | 70359770 | 70360896 | ENSMUSG00000030551 | Nr2f2 | nuclear receptor subfamily 2, group F, member 2 |
| chr7 | 70360896 | 70362190 | ENSMUSG00000030551 | Nr2f2 | nuclear receptor subfamily 2, group F, member 2 |
| chr7 | 73557718 | 73558654 | ENSMUSG00000101970 | Chaserr | CHD2 adjacent suppressive regulatory RNA |
| chr7 | 73618124 | 73618891 | ENSMUSG00000101970 | Chaserr | CHD2 adjacent suppressive regulatory RNA |
| chr7 | 74554067 | 74555189 | ENSMUSG00000025790 | Slco3a1 | solute carrier organic anion transporter family, member 3a1 |
| chr7 | 79465707 | 79466668 | ENSMUSG00000039176 | Polg | polymerase (DNA directed), gamma |
| chr7 | 80114534 | 80115430 | ENSMUSG00000030541 | Idh2 | isocitrate dehydrogenase 2 (NADP+), mitochondrial |
| chr7 | 80198178 | 80198926 | ENSMUSG00000030539 | Sema4b | sema domain, immunoglobulin domain (Ig), transmembrane domain (TM) and short cytoplasmic domain, (semaphorin) 4B |
| chr7 | 80370634 | 80371541 | ENSMUSG00000038886 | Man2a2 | mannosidase 2, alpha 2 |
| chr7 | 80687896 | 80689138 | ENSMUSG00000030527 | Crtc3 | CREB regulated transcription coactivator 3 |
| chr7 | 80993075 | 80994171 | ENSMUSG00000005621 | Zfp592 | zinc finger protein 592 |
| chr7 | 81571177 | 81572122 | ENSMUSG00000045795 | Whamm | WAS protein homolog associated with actin, golgi membranes and microtubules |
| chr7 | 81789164 | 81789648 | ENSMUSG00000025102 | 3110040N11Rik | RIKEN cDNA 3110040N11 gene |
| chr7 | 81828753 | 81829762 | ENSMUSG00000025103 | Btbd1 | BTB (POZ) domain containing 1 |
| chr7 | 81933461 | 81934600 | ENSMUSG00000025104 | Hdgfl3 | HDGF like 3 |
| chr7 | 83882795 | 83884678 | ENSMUSG00000038503 | Mesd | mesoderm development LRP chaperone |
| chr7 | 84151072 | 84152166 | ENSMUSG00000038459 | Abhd17c | abhydrolase domain containing 17C |
| chr7 | 84689491 | 84690075 | ENSMUSG00000030629 | Zfand6 | zinc finger, AN1-type domain 6 |
| chr7 | 89517300 | 89517837 | ENSMUSG00000039405 | Prss23 | protease, serine 23 |
| chr7 | 90442717 | 90444081 | ENSMUSG00000051451 | Crebzf | CREB/ATF bZIP transcription factor |
| chr7 | 98177789 | 98178382 | ENSMUSG00000035547 | Capn5 | calpain 5 |
| chr7 | 100159198 | 100159875 | ENSMUSG00000030725 | Lipt2 | lipoyl(octanoyl) transferase 2 (putative) |
| chr7 | 100607244 | 100608536 | ENSMUSG00000030704 | Rab6a | RAB6A, member RAS oncogene family |
| chr7 | 100931000 | 100932173 | ENSMUSG00000032875 | Arhgef17 | Rho guanine nucleotide exchange factor (GEF) 17 |
| chr7 | 101011697 | 101012290 | ENSMUSG00000032860 | P2ry2 | purinergic receptor P2Y, G-protein coupled 2 |
| chr7 | 101969642 | 101970538 | ENSMUSG00000066306 | Numa1 | nuclear mitotic apparatus protein 1 |
| chr7 | 105633244 | 105633726 | ENSMUSG00000036989 | Trim3 | tripartite motif-containing 3 |
| chr7 | 110121428 | 110123162 | ENSMUSG00000031016 | Wee1 | WEE 1 homolog 1 (S. pombe) |
| chr7 | 110614167 | 110615143 | ENSMUSG00000038371 | Sbf2 | SET binding factor 2 |
| chr7 | 110615143 | 110615304 | ENSMUSG00000038371 | Sbf2 | SET binding factor 2 |
| chr7 | 110768385 | 110769068 | ENSMUSG00000005686 | Ampd3 | adenosine monophosphate deaminase 3 |
| chr7 | 111081914 | 111083545 | ENSMUSG00000005610 | Eif4g2 | eukaryotic translation initiation factor 4, gamma 2 |
| chr7 | 112023208 | 112024175 | ENSMUSG00000059263 | Usp47 | ubiquitin specific peptidase 47 |
| chr7 | 112679165 | 112680139 | ENSMUSG00000055320 | Tead1 | TEA domain family member 1 |
| chr7 | 114116843 | 114117807 | ENSMUSG00000055723 | Rras2 | related RAS viral (r-ras) oncogene 2 |
| chr7 | 116307217 | 116308380 | ENSMUSG00000045659 | Plekha7 | pleckstrin homology domain containing, family A member 7 |
| chr7 | 118242866 | 118243794 | ENSMUSG00000030655 | Smg1 | SMG1 homolog, phosphatidylinositol 3-kinase-related kinase (C. elegans) |
| chr7 | 118855483 | 118856198 | ENSMUSG00000073856 | Iqck | IQ motif containing K |
| chr7 | 120842656 | 120843489 | ENSMUSG00000035064 | Eef2k | eukaryotic elongation factor-2 kinase |
| chr7 | 120917809 | 120918301 | ENSMUSG00000030880 | Polr3e | polymerase (RNA) III (DNA directed) polypeptide E |
| chr7 | 121034466 | 121035383 | ENSMUSG00000030876 | Mettl9 | methyltransferase like 9 |
| chr7 | 126272294 | 126272739 | ENSMUSG00000042978 | Sbk1 | SH3-binding kinase 1 |
| chr7 | 126502876 | 126503764 | ENSMUSG00000032637 | Atxn2l | ataxin 2-like |
| chr7 | 127232786 | 127233584 | ENSMUSG00000000486 | Gm4532 | predicted gene 4532 |
| chr7 | 127253715 | 127254798 | ENSMUSG00000042462 | Dctpp1 | dCTP pyrophosphatase 1 |
| chr7 | 127512083 | 127513052 | ENSMUSG00000053877 | Srcap | Snf2-related CREBBP activator protein |
| chr7 | 127708304 | 127709045 | ENSMUSG00000030814 | Bcl7c | B cell CLL/lymphoma 7C |
| chr7 | 127776960 | 127777976 | ENSMUSG00000042308 | Setd1a | SET domain containing 1A |
| chr7 | 128739696 | 128740636 | ENSMUSG00000048170 | Mcmbp | minichromosome maintenance complex binding protein |
| chr7 | 130265745 | 130266553 | ENSMUSG00000030849 | Fgfr2 | fibroblast growth factor receptor 2 |
| chr7 | 130865611 | 130866564 | ENSMUSG00000040268 | Plekha1 | pleckstrin homology domain containing, family A (phosphoinositide binding specific) member 1 |
| chr7 | 132812902 | 132814116 | ENSMUSG00000030956 | Fam53b | family with sequence similarity 53, member B |
| chr7 | 132930860 | 132931856 | ENSMUSG00000030967 | Zranb1 | zinc finger, RAN-binding domain containing 1 |
| chr7 | 134670397 | 134671094 | ENSMUSG00000058325 | Dock1 | dedicator of cytokinesis 1 |
| chr7 | 138909364 | 138909928 | ENSMUSG00000078566 | Bnip3 | BCL2/adenovirus E1B interacting protein 3 |
| chr7 | 139248171 | 139248945 | ENSMUSG00000060260 | Pwwp2b | PWWP domain containing 2B |
| chr7 | 141193265 | 141194453 | ENSMUSG00000025499 | Hras | Harvey rat sarcoma virus oncogene |
| chr7 | 141561915 | 141562877 | ENSMUSG00000002957 | Ap2a2 | adaptor-related protein complex 2, alpha 2 subunit |
| chr7 | 142094688 | 142095597 | ENSMUSG00000037887 | Dusp8 | dual specificity phosphatase 8 |
| chr8 | 3500210 | 3500880 | ENSMUSG00000004567 | Mcoln1 | mucolipin 1 |
| chr8 | 3587575 | 3588142 | ENSMUSG00000044433 | Camsap3 | calmodulin regulated spectrin-associated protein family, member 3 |
| chr8 | 4324590 | 4325475 | ENSMUSG00000040028 | Elavl1 | ELAV (embryonic lethal, abnormal vision)-like 1 (Hu antigen R) |
| chr8 | 8658976 | 8660511 | ENSMUSG00000001300 | Efnb2 | ephrin B2 |
| chr8 | 8660511 | 8661902 | ENSMUSG00000001300 | Efnb2 | ephrin B2 |
| chr8 | 8689579 | 8690754 | ENSMUSG00000098917 | Mir7654 | microRNA 7654 |
| chr8 | 9977671 | 9978333 | ENSMUSG00000049717 | Lig4 | ligase IV, DNA, ATP-dependent |
| chr8 | 11550340 | 11550805 | ENSMUSG00000056228 | Cars2 | cysteinyl-tRNA synthetase 2 (mitochondrial)(putative) |
| chr8 | 11634820 | 11636769 | ENSMUSG00000031508 | Ankrd10 | ankyrin repeat domain 10 |
| chr8 | 12756785 | 12757872 | ENSMUSG00000031441 | Atp11a | ATPase, class VI, type 11A |
| chr8 | 13159086 | 13159820 | ENSMUSG00000031447 | Lamp1 | lysosomal-associated membrane protein 1 |
| chr8 | 13338423 | 13339544 | ENSMUSG00000038482 | Tfdp1 | transcription factor Dp 1 |
| chr8 | 13339544 | 13339876 | ENSMUSG00000038482 | Tfdp1 | transcription factor Dp 1 |
| chr8 | 13889904 | 13890506 | ENSMUSG00000031458 | Coprs | coordinator of PRMT5, differentiation stimulator |
| chr8 | 18594954 | 18595766 | ENSMUSG00000039842 | Mcph1 | microcephaly, primary autosomal recessive 1 |
| chr8 | 18845841 | 18847085 | ENSMUSG00000031467 | Agpat5 | 1-acylglycerol-3-phosphate O-acyltransferase 5 (lysophosphatidic acid acyltransferase, epsilon) |
| chr8 | 22192663 | 22193258 | ENSMUSG00000031479 | Vps36 | vacuolar protein sorting 36 |
| chr8 | 22398029 | 22398697 | ENSMUSG00000031482 | Slc25a15 | solute carrier family 25 (mitochondrial carrier ornithine transporter), member 15 |
| chr8 | 22476342 | 22477304 | ENSMUSG00000031534 | Smim19 | small integral membrane protein 19 |
| chr8 | 22593242 | 22594237 | ENSMUSG00000008892 | Vdac3 | voltage-dependent anion channel 3 |
| chr8 | 22706218 | 22706697 | ENSMUSG00000031537 | Ikbkb | inhibitor of kappaB kinase beta |
| chr8 | 22858970 | 22860333 | ENSMUSG00000031540 | Kat6a | K(lysine) acetyltransferase 6A |
| chr8 | 23207776 | 23208579 | ENSMUSG00000031545 | Gpat4 | glycerol-3-phosphate acyltransferase 4 |
| chr8 | 25016377 | 25017665 | ENSMUSG00000031555 | Adam9 | a disintegrin and metallopeptidase domain 9 (meltrin gamma) |
| chr8 | 25596981 | 25597727 | ENSMUSG00000037363 | Letm2 | leucine zipper-EF-hand containing transmembrane protein 2 |
| chr8 | 25601289 | 25602938 | ENSMUSG00000054823 | Nsd3 | nuclear receptor binding SET domain protein 3 |
| chr8 | 25753766 | 25754833 | ENSMUSG00000091514 | Gm17484 | predicted gene, 17484 |
| chr8 | 25784890 | 25785786 | ENSMUSG00000037296 | Lsm1 | LSM1 homolog, mRNA degradation associated |
| chr8 | 26118765 | 26119901 | ENSMUSG00000037234 | Hook3 | hook microtubule tethering protein 3 |
| chr8 | 26158375 | 26159021 | ENSMUSG00000037214 | Thap1 | THAP domain containing, apoptosis associated protein 1 |
| chr8 | 32008416 | 32009839 | ENSMUSG00000062991 | Nrg1 | neuregulin 1 |
| chr8 | 33385423 | 33386251 | ENSMUSG00000031583 | Wrn | Werner syndrome RecQ like helicase |
| chr8 | 33599484 | 33600334 | ENSMUSG00000009630 | Ppp2cb | protein phosphatase 2 (formerly 2A), catalytic subunit, beta isoform |
| chr8 | 33653308 | 33654270 | ENSMUSG00000031584 | Gsr | glutathione reductase |
| chr8 | 33731658 | 33732713 | ENSMUSG00000031585 | Gtf2e2 | general transcription factor II E, polypeptide 2 (beta subunit) |
| chr8 | 33929111 | 33930469 | ENSMUSG00000031586 | Rbpms | RNA binding protein gene with multiple splicing |
| chr8 | 34146302 | 34147091 | ENSMUSG00000031513 | Leprotl1 | leptin receptor overlapping transcript-like 1 |
| chr8 | 34806826 | 34808773 | ENSMUSG00000031530 | Dusp4 | dual specificity phosphatase 4 |
| chr8 | 34808773 | 34810398 | ENSMUSG00000031530 | Dusp4 | dual specificity phosphatase 4 |
| chr8 | 35494893 | 35495689 | ENSMUSG00000031527 | Eri1 | exoribonuclease 1 |
| chr8 | 36247870 | 36249744 | ENSMUSG00000039633 | Lonrf1 | LON peptidase N-terminal domain and ring finger 1 |
| chr8 | 39005765 | 39006535 | ENSMUSG00000039530 | Tusc3 | tumor suppressor candidate 3 |
| chr8 | 40423271 | 40424482 | ENSMUSG00000039470 | Zdhhc2 | zinc finger, DHHC domain containing 2 |
| chr8 | 40510891 | 40512408 | ENSMUSG00000031601 | Cnot7 | CCR4-NOT transcription complex, subunit 7 |
| chr8 | 41239396 | 41240295 | ENSMUSG00000031592 | Pcm1 | pericentriolar material 1 |
| chr8 | 41374271 | 41375181 | ENSMUSG00000031591 | Asah1 | N-acylsphingosine amidohydrolase 1 |
| chr8 | 44936980 | 44937748 | ENSMUSG00000070047 | Fat1 | FAT atypical cadherin 1 |
| chr8 | 44937748 | 44938686 | ENSMUSG00000070047 | Fat1 | FAT atypical cadherin 1 |
| chr8 | 45999612 | 46000269 | ENSMUSG00000050914 | Ankrd37 | ankyrin repeat domain 37 |
| chr8 | 46151541 | 46152585 | ENSMUSG00000038291 | Snx25 | sorting nexin 25 |
| chr8 | 46163236 | 46163957 | ENSMUSG00000031631 | Cfap97 | cilia and flagella associated protein 97 |
| chr8 | 46210331 | 46211247 | ENSMUSG00000031633 | Slc25a4 | solute carrier family 25 (mitochondrial carrier, adenine nucleotide translocator), member 4 |
| chr8 | 46470923 | 46471719 | ENSMUSG00000018796 | Acsl1 | acyl-CoA synthetase long-chain family member 1 |
| chr8 | 46740429 | 46741654 | ENSMUSG00000031627 | Irf2 | interferon regulatory factor 2 |
| chr8 | 47713153 | 47714618 | ENSMUSG00000038069 | Cdkn2aip | CDKN2A interacting protein |
| chr8 | 47989447 | 47991372 | ENSMUSG00000031563 | Wwc2 | WW, C2 and coiled-coil domain containing 2 |
| chr8 | 48109958 | 48110748 | ENSMUSG00000031562 | Dctd | dCMP deaminase |
| chr8 | 54529361 | 54530165 | ENSMUSG00000031519 | Asb5 | ankyrin repeat and SOCs box-containing 5 |
| chr8 | 57651666 | 57653421 |  | AW046200 | expressed sequence AW046200 |
| chr8 | 60632392 | 60633451 | ENSMUSG00000031647 | Mfap3l | microfibrillar-associated protein 3-like |
| chr8 | 60982675 | 60983954 | ENSMUSG00000004319 | Clcn3 | chloride channel, voltage-sensitive 3 |
| chr8 | 61223754 | 61225049 | ENSMUSG00000031642 | Sh3rf1 | SH3 domain containing ring finger 1 |
| chr8 | 64692551 | 64693301 | ENSMUSG00000037852 | Cpe | carboxypeptidase E |
| chr8 | 64733275 | 64733948 | ENSMUSG00000031604 | Msmo1 | methylsterol monoxygenase 1 |
| chr8 | 64849206 | 64850344 | ENSMUSG00000031605 | Klhl2 | kelch-like 2, Mayven |
| chr8 | 64947045 | 64947806 | ENSMUSG00000025521 | Tmem192 | transmembrane protein 192 |
| chr8 | 66860116 | 66860514 | ENSMUSG00000014907 | Naf1 | nuclear assembly factor 1 ribonucleoprotein |
| chr8 | 66860911 | 66861265 | ENSMUSG00000014907 | Naf1 | nuclear assembly factor 1 ribonucleoprotein |
| chr8 | 69973711 | 69974691 | ENSMUSG00000036180 | Gatad2a | GATA zinc finger domain containing 2A |
| chr8 | 69995810 | 69996880 | ENSMUSG00000036180 | Gatad2a | GATA zinc finger domain containing 2A |
| chr8 | 70233874 | 70235033 | ENSMUSG00000036054 | Sugp2 | SURP and G patch domain containing 2 |
| chr8 | 70352618 | 70353510 | ENSMUSG00000058301 | Upf1 | UPF1 regulator of nonsense transcripts homolog (yeast) |
| chr8 | 70439188 | 70440034 | ENSMUSG00000003575 | Crtc1 | CREB regulated transcription coactivator 1 |
| chr8 | 70510056 | 70510692 | ENSMUSG00000090137 | Uba52 | ubiquitin A-52 residue ribosomal protein fusion product 1 |
| chr8 | 70594011 | 70595215 | ENSMUSG00000019139 | Isyna1 | myo-inositol 1-phosphate synthase A1 |
| chr8 | 70609136 | 70609915 | ENSMUSG00000070003 | Ssbp4 | single stranded DNA binding protein 4 |
| chr8 | 70698761 | 70700241 | ENSMUSG00000071076 | Jund | jun D proto-oncogene |
| chr8 | 70760359 | 70760998 | ENSMUSG00000035559 | Mpv17l2 | MPV17 mitochondrial membrane protein-like 2 |
| chr8 | 71380894 | 71382527 | ENSMUSG00000002393 | Nr2f6 | nuclear receptor subfamily 2, group F, member 6 |
| chr8 | 71476065 | 71477095 | ENSMUSG00000074247 | Dda1 | DET1 and DDB1 associated 1 |
| chr8 | 71591734 | 71592829 | ENSMUSG00000031807 | Pgls | 6-phosphogluconolactonase |
| chr8 | 71610860 | 71611705 | ENSMUSG00000034807 | Colgalt1 | collagen beta(1-O)galactosyltransferase 1 |
| chr8 | 71676292 | 71676612 | ENSMUSG00000031805 | Jak3 | Janus kinase 3 |
| chr8 | 71725438 | 71725872 | ENSMUSG00000070000 | Fcho1 | FCH domain only 1 |
| chr8 | 72135002 | 72136564 | ENSMUSG00000031799 | Tpm4 | tropomyosin 4 |
| chr8 | 72491554 | 72493075 | ENSMUSG00000019731 | Slc35e1 | solute carrier family 35, member E1 |
| chr8 | 73352581 | 73353975 | ENSMUSG00000004383 | Large1 | LARGE xylosyl- and glucuronyltransferase 1 |
| chr8 | 77517275 | 77518431 | ENSMUSG00000037148 | Arhgap10 | Rho GTPase activating protein 10 |
| chr8 | 78508408 | 78509752 | ENSMUSG00000037070 | Rbmxl1 | RNA binding motif protein, X-linked like-1 |
| chr8 | 78820640 | 78821215 | ENSMUSG00000031683 | Lsm6 | LSM6 homolog, U6 small nuclear RNA and mRNA degradation associated |
| chr8 | 79294609 | 79295475 | ENSMUSG00000037022 | Mmaa | methylmalonic aciduria (cobalamin deficiency) type A |
| chr8 | 79398373 | 79399794 | ENSMUSG00000031681 | Smad1 | SMAD family member 1 |
| chr8 | 79638703 | 79640585 | ENSMUSG00000036990 | Otud4 | OTU domain containing 4 |
| chr8 | 79711198 | 79712047 | ENSMUSG00000058355 | Abce1 | ATP-binding cassette, sub-family E (OABP), member 1 |
| chr8 | 80738359 | 80739443 | ENSMUSG00000031715 | Smarca5 | SWI/SNF related, matrix associated, actin dependent regulator of chromatin, subfamily a, member 5 |
| chr8 | 80879261 | 80880795 | ENSMUSG00000031714 | Gab1 | growth factor receptor bound protein 2-associated protein 1 |
| chr8 | 83441987 | 83442679 | ENSMUSG00000063253 | Scoc | short coiled-coil protein |
| chr8 | 83698309 | 83699474 | ENSMUSG00000057672 | Pkn1 | protein kinase N1 |
| chr8 | 83899938 | 83901388 | ENSMUSG00000013033 | Adgrl1 | adhesion G protein-coupled receptor L1 |
| chr8 | 84011067 | 84012872 | ENSMUSG00000080665 | Mir1199 | microRNA 1199 |
| chr8 | 84638792 | 84639895 | ENSMUSG00000034656 | Cacna1a | calcium channel, voltage-dependent, P/Q type, alpha 1A subunit |
| chr8 | 84661538 | 84663011 | ENSMUSG00000053560 | Ier2 | immediate early response 2 |
| chr8 | 84687401 | 84688064 | ENSMUSG00000001910 | Nacc1 | nucleus accumbens associated 1, BEN and BTB (POZ) domain containing |
| chr8 | 84799054 | 84800738 | ENSMUSG00000001911 | Nfix | nuclear factor I/X |
| chr8 | 84810961 | 84811270 | ENSMUSG00000001911 | Nfix | nuclear factor I/X |
| chr8 | 84977161 | 84979001 | ENSMUSG00000052837 | Junb | jun B proto-oncogene |
| chr8 | 85026698 | 85027481 | ENSMUSG00000041203 | Trir | telomerase RNA component interacting RNase |
| chr8 | 85059955 | 85060662 | ENSMUSG00000087026 | A230103J11Rik | RIKEN cDNA A230103J11 gene |
| chr8 | 85432211 | 85433074 | ENSMUSG00000036934 | 4921524J17Rik | RIKEN cDNA 4921524J17 gene |
| chr8 | 85554598 | 85555704 | ENSMUSG00000031701 | Dnaja2 | DnaJ heat shock protein family (Hsp40) member A2 |
| chr8 | 85840423 | 85841259 | ENSMUSG00000036879 | Phkb | phosphorylase kinase beta |
| chr8 | 86623888 | 86624671 | ENSMUSG00000047866 | Lonp2 | lon peptidase 2, peroxisomal |
| chr8 | 86745303 | 86746351 | ENSMUSG00000036840 | Siah1a | siah E3 ubiquitin protein ligase 1A |
| chr8 | 86884472 | 86885481 | ENSMUSG00000031652 | N4bp1 | NEDD4 binding protein 1 |
| chr8 | 88137764 | 88138574 | ENSMUSG00000099080 | Mir7071 | microRNA 7071 |
| chr8 | 88198835 | 88200625 | ENSMUSG00000036779 | Tent4b | terminal nucleotidyltransferase 4B |
| chr8 | 88361089 | 88362381 | ENSMUSG00000031660 | Brd7 | bromodomain containing 7 |
| chr8 | 90907889 | 90909458 | ENSMUSG00000056608 | Chd9 | chromodomain helicase DNA binding protein 9 |
| chr8 | 91133482 | 91134375 | ENSMUSG00000031667 | Aktip | thymoma viral proto-oncogene 1 interacting protein |
| chr8 | 94011622 | 94013039 | ENSMUSG00000031751 | Amfr | autocrine motility factor receptor |
| chr8 | 94214256 | 94215006 | ENSMUSG00000032939 | Nup93 | nucleoporin 93 |
| chr8 | 94386108 | 94387166 | ENSMUSG00000031770 | Herpud1 | homocysteine-inducible, endoplasmic reticulum stress-inducible, ubiquitin-like domain member 1 |
| chr8 | 94387166 | 94387659 | ENSMUSG00000031770 | Herpud1 | homocysteine-inducible, endoplasmic reticulum stress-inducible, ubiquitin-like domain member 1 |
| chr8 | 94532776 | 94533272 | ENSMUSG00000034361 | Cpne2 | copine II |
| chr8 | 94666518 | 94667212 | ENSMUSG00000031776 | Arl2bp | ADP-ribosylation factor-like 2 binding protein |
| chr8 | 95142316 | 95143168 | ENSMUSG00000031788 | Kifc3 | kinesin family member C3 |
| chr8 | 95331370 | 95332350 | ENSMUSG00000031792 | Usb1 | U6 snRNA biogenesis 1 |
| chr8 | 95488019 | 95489067 | ENSMUSG00000046707 | Csnk2a2 | casein kinase 2, alpha prime polypeptide |
| chr8 | 95715749 | 95716523 | ENSMUSG00000031671 | Setd6 | SET domain containing 6 |
| chr8 | 104394905 | 104395961 | ENSMUSG00000096188 | Cmtm4 | CKLF-like MARVEL transmembrane domain containing 4 |
| chr8 | 105170366 | 105171885 | ENSMUSG00000031885 | Cbfb | core binding factor beta |
| chr8 | 105496425 | 105497161 | ENSMUSG00000039199 | Zdhhc1 | zinc finger, DHHC domain containing 1 |
| chr8 | 105518697 | 105519267 | ENSMUSG00000031891 | Hsd11b2 | hydroxysteroid 11-beta dehydrogenase 2 |
| chr8 | 105605141 | 105606157 | ENSMUSG00000038604 | Ripor1 | RHO family interacting cell polarization regulator 1 |
| chr8 | 105606157 | 105606919 | ENSMUSG00000038604 | Ripor1 | RHO family interacting cell polarization regulator 1 |
| chr8 | 105636068 | 105637394 | ENSMUSG00000005705 | Agrp | agouti related neuropeptide |
| chr8 | 105758201 | 105758838 | ENSMUSG00000013150 | Gfod2 | glucose-fructose oxidoreductase domain containing 2 |
| chr8 | 105860132 | 105861304 | ENSMUSG00000008450 | Nutf2 | nuclear transport factor 2 |
| chr8 | 105900296 | 105900914 | ENSMUSG00000048310 | Pskh1 | protein serine kinase H1 |
| chr8 | 105965444 | 105966224 | ENSMUSG00000017765 | Slc12a4 | solute carrier family 12, member 4 |
| chr8 | 106059397 | 106060280 | ENSMUSG00000031902 | Nfatc3 | nuclear factor of activated T cells, cytoplasmic, calcineurin dependent 3 |
| chr8 | 106168627 | 106169621 | ENSMUSG00000031904 | Slc7a6 | solute carrier family 7 (cationic amino acid transporter, y+ system), member 6 |
| chr8 | 106210528 | 106211303 | ENSMUSG00000060098 | Prmt7 | protein arginine N-methyltransferase 7 |
| chr8 | 106892780 | 106894045 | ENSMUSG00000046691 | Chtf8 | CTF8, chromosome transmission fidelity factor 8 |
| chr8 | 106935868 | 106936713 | ENSMUSG00000041308 | Sntb2 | syntrophin, basic 2 |
| chr8 | 107047943 | 107048986 | ENSMUSG00000078931 | Pdf | peptide deformylase (mitochondrial) |
| chr8 | 107096060 | 107096609 | ENSMUSG00000098671 | Mir7075 | microRNA 7075 |
| chr8 | 108703762 | 108704930 | ENSMUSG00000038872 | Zfhx3 | zinc finger homeobox 3 |
| chr8 | 108714549 | 108715518 | ENSMUSG00000038872 | Zfhx3 | zinc finger homeobox 3 |
| chr8 | 109608342 | 109608727 | ENSMUSG00000031730 | Dhodh | dihydroorotate dehydrogenase |
| chr8 | 109705297 | 109706398 | ENSMUSG00000031728 | Zfp821 | zinc finger protein 821 |
| chr8 | 109868321 | 109869416 | ENSMUSG00000031732 | Phlpp2 | PH domain and leucine rich repeat protein phosphatase 2 |
| chr8 | 110618378 | 110619391 | ENSMUSG00000010936 | Vac14 | Vac14 homolog (S. cerevisiae) |
| chr8 | 111056230 | 111057276 | ENSMUSG00000109941 | Exosc6 | exosome component 6 |
| chr8 | 111094443 | 111095195 | ENSMUSG00000033624 | Pdpr | pyruvate dehydrogenase phosphatase regulatory subunit |
| chr8 | 111144975 | 111146022 |  | 9430091E24Rik | RIKEN cDNA 9430091E24 gene |
| chr8 | 111258482 | 111259441 | ENSMUSG00000003316 | Glg1 | golgi apparatus protein 1 |
| chr8 | 111536251 | 111537597 | ENSMUSG00000033545 | Znrf1 | zinc and ring finger 1 |
| chr8 | 111643527 | 111644007 | ENSMUSG00000055835 | Zfp1 | zinc finger protein 1 |
| chr8 | 115706874 | 115707985 | ENSMUSG00000055435 | Maf | avian musculoaponeurotic fibrosarcoma oncogene homolog |
| chr8 | 117157751 | 117158554 | ENSMUSG00000052557 | Gan | giant axonal neuropathy |
| chr8 | 117256302 | 117257665 | ENSMUSG00000034390 | Cmip | c-Maf inducing protein |
| chr8 | 119394658 | 119395560 | ENSMUSG00000074064 | Mlycd | malonyl-CoA decarboxylase |
| chr8 | 119558401 | 119558743 | ENSMUSG00000031835 | Mbtps1 | membrane-bound transcription factor peptidase, site 1 |
| chr8 | 119574651 | 119575373 | ENSMUSG00000031831 | Dnaaf1 | dynein, axonemal assembly factor 1 |
| chr8 | 119910539 | 119911604 | ENSMUSG00000089704 | Galnt2 | polypeptide N-acetylgalactosaminyltransferase 2 |
| chr8 | 119918314 | 119918761 | ENSMUSG00000031826 | Usp10 | ubiquitin specific peptidase 10 |
| chr8 | 120113649 | 120114881 | ENSMUSG00000031824 | 6430548M08Rik | RIKEN cDNA 6430548M08 gene |
| chr8 | 120634147 | 120634748 | ENSMUSG00000043687 | 1190005I06Rik | RIKEN cDNA 1190005I06 gene |
| chr8 | 120667578 | 120668738 | ENSMUSG00000097919 | Gm27021 | predicted gene, 27021 |
| chr8 | 121578327 | 121578941 | ENSMUSG00000052934 | Fbxo31 | F-box protein 31 |
| chr8 | 121590257 | 121591002 | ENSMUSG00000031812 | Map1lc3b | microtubule-associated protein 1 light chain 3 beta |
| chr8 | 121651768 | 121653108 | ENSMUSG00000061410 | Zcchc14 | zinc finger, CCHC domain containing 14 |
| chr8 | 121906735 | 121907855 | ENSMUSG00000040010 | Slc7a5 | solute carrier family 7 (cationic amino acid transporter, y+ system), member 5 |
| chr8 | 121950405 | 121951421 | ENSMUSG00000025316 | Banp | BTG3 associated nuclear protein |
| chr8 | 122230129 | 122230830 | ENSMUSG00000043903 | Zfp469 | zinc finger protein 469 |
| chr8 | 122550388 | 122551840 | ENSMUSG00000014444 | Piezo1 | piezo-type mechanosensitive ion channel component 1 |
| chr8 | 122567362 | 122568358 | ENSMUSG00000006585 | Cdt1 | chromatin licensing and DNA replication factor 1 |
| chr8 | 122576220 | 122576869 | ENSMUSG00000006589 | Aprt | adenine phosphoribosyl transferase |
| chr8 | 122677899 | 122678773 | ENSMUSG00000006362 | Cbfa2t3 | CBFA2/RUNX1 translocation partner 3 |
| chr8 | 123041041 | 123042954 | ENSMUSG00000099881 | 2810013P06Rik | RIKEN cDNA 2810013P06 gene |
| chr8 | 123332459 | 123333384 | ENSMUSG00000010154 | Spire2 | spire type actin nucleation factor 2 |
| chr8 | 123373265 | 123374094 | ENSMUSG00000001472 | Tcf25 | transcription factor 25 (basic helix-loop-helix) |
| chr8 | 123477795 | 123478244 | ENSMUSG00000031967 | Afg3l1 | AFG3-like AAA ATPase 1 |
| chr8 | 123653641 | 123654542 | ENSMUSG00000039960 | Rhou | ras homolog family member U |
| chr8 | 123859007 | 123860083 | ENSMUSG00000031971 | Ccsap | centriole, cilia and spindle associated protein |
| chr8 | 123982292 | 123983312 | ENSMUSG00000031974 | Abcb10 | ATP-binding cassette, sub-family B (MDR/TAP), member 10 |
| chr8 | 124020695 | 124021693 | ENSMUSG00000038697 | Taf5l | TATA-box binding protein associated factor 5 like |
| chr8 | 124231217 | 124232243 | ENSMUSG00000089704 | Galnt2 | polypeptide N-acetylgalactosaminyltransferase 2 |
| chr8 | 124896944 | 124898569 | ENSMUSG00000074030 | Exoc8 | exocyst complex component 8 |
| chr8 | 124948139 | 124949419 | ENSMUSG00000031987 | Egln1 | egl-9 family hypoxia-inducible factor 1 |
| chr8 | 125012670 | 125013329 | ENSMUSG00000056820 | Tsnax | translin-associated factor X |
| chr8 | 125569454 | 125570374 | ENSMUSG00000001995 | Sipa1l2 | signal-induced proliferation-associated 1 like 2 |
| chr8 | 125909700 | 125910611 | ENSMUSG00000031853 | Map3k21 | mitogen-activated protein kinase kinase kinase 21 |
| chr8 | 125910611 | 125911585 | ENSMUSG00000031853 | Map3k21 | mitogen-activated protein kinase kinase kinase 21 |
| chr8 | 125995020 | 125996747 | ENSMUSG00000033998 | Kcnk1 | potassium channel, subfamily K, member 1 |
| chr8 | 126474070 | 126475243 | ENSMUSG00000090290 | Tarbp1 | TAR RNA binding protein 1 |
| chr8 | 126945142 | 126946069 | ENSMUSG00000093904 | Tomm20 | translocase of outer mitochondrial membrane 20 |
| chr8 | 127063576 | 127065079 | ENSMUSG00000025812 | Pard3 | par-3 family cell polarity regulator |
| chr8 | 128685344 | 128686440 | ENSMUSG00000025809 | Itgb1 | integrin beta 1 (fibronectin receptor beta) |
| chr9 | 7763866 | 7764610 | ENSMUSG00000050912 | Tmem123 | transmembrane protein 123 |
| chr9 | 7836387 | 7837410 | ENSMUSG00000057367 | Birc2 | baculoviral IAP repeat-containing 2 |
| chr9 | 8003440 | 8004946 | ENSMUSG00000053110 | Yap1 | yes-associated protein 1 |
| chr9 | 9238543 | 9239288 | ENSMUSG00000050730 | Arhgap42 | Rho GTPase activating protein 42 |
| chr9 | 13748930 | 13749649 | ENSMUSG00000031918 | Mtmr2 | myotubularin related protein 2 |
| chr9 | 14276047 | 14276906 | ENSMUSG00000032009 | Sesn3 | sestrin 3 |
| chr9 | 14380451 | 14381445 | ENSMUSG00000037419 | Endod1 | endonuclease domain containing 1 |
| chr9 | 15493201 | 15494127 | ENSMUSG00000058173 | Smco4 | single-pass membrane protein with coiled-coil domains 4 |
| chr9 | 20877927 | 20878656 | ENSMUSG00000038742 | Angptl6 | angiopoietin-like 6 |
| chr9 | 21238508 | 21239484 | ENSMUSG00000003308 | Keap1 | kelch-like ECH-associated protein 1 |
| chr9 | 21546142 | 21547517 | ENSMUSG00000032185 | Carm1 | coactivator-associated arginine methyltransferase 1 |
| chr9 | 21615943 | 21616595 | ENSMUSG00000032187 | Smarca4 | SWI/SNF related, matrix associated, actin dependent regulator of chromatin, subfamily a, member 4 |
| chr9 | 22070804 | 22071483 | ENSMUSG00000096981 | Gm16845 | predicted gene, 16845 |
| chr9 | 22467617 | 22468447 | ENSMUSG00000032239 | Rp9 | retinitis pigmentosa 9 (human) |
| chr9 | 24502660 | 24503681 | ENSMUSG00000043067 | Dpy19l1 | dpy-19-like 1 (C. elegans) |
| chr9 | 25150863 | 25152325 | ENSMUSG00000008429 | Herpud2 | HERPUD family member 2 |
| chr9 | 25481363 | 25482006 | ENSMUSG00000036611 | Eepd1 | endonuclease/exonuclease/phosphatase family domain containing 1 |
| chr9 | 31030123 | 31031261 | ENSMUSG00000047412 | Zbtb44 | zinc finger and BTB domain containing 44 |
| chr9 | 31131201 | 31132056 | ENSMUSG00000031995 | St14 | suppression of tumorigenicity 14 (colon carcinoma) |
| chr9 | 31211089 | 31211883 | ENSMUSG00000031996 | Aplp2 | amyloid beta (A4) precursor-like protein 2 |
| chr9 | 31280752 | 31281446 | ENSMUSG00000042496 | Prdm10 | PR domain containing 10 |
| chr9 | 32695579 | 32696730 | ENSMUSG00000032035 | Ets1 | E26 avian leukemia oncogene 1, 5' domain |
| chr9 | 35116425 | 35117326 | ENSMUSG00000032038 | St3gal4 | ST3 beta-galactoside alpha-2,3-sialyltransferase 4 |
| chr9 | 35421090 | 35422060 | ENSMUSG00000038119 | Cdon | cell adhesion molecule-related/down-regulated by oncogenes |
| chr9 | 36796903 | 36797403 | ENSMUSG00000062762 | Ei24 | etoposide induced 2.4 mRNA |
| chr9 | 42263800 | 42264562 | ENSMUSG00000032018 | Sc5d | sterol-C5-desaturase |
| chr9 | 43105323 | 43106309 | ENSMUSG00000059495 | Arhgef12 | Rho guanine nucleotide exchange factor (GEF) 12 |
| chr9 | 44233486 | 44234319 | ENSMUSG00000034342 | Cbl | Casitas B-lineage lymphoma |
| chr9 | 44334602 | 44335865 | ENSMUSG00000049932 | H2ax | H2A.X variant histone |
| chr9 | 44498334 | 44499131 | ENSMUSG00000063382 | Bcl9l | B cell CLL/lymphoma 9-like |
| chr9 | 44603953 | 44605538 | ENSMUSG00000032097 | Ddx6 | DEAD (Asp-Glu-Ala-Asp) box polypeptide 6 |
| chr9 | 47529915 | 47531832 | ENSMUSG00000032076 | Cadm1 | cell adhesion molecule 1 |
| chr9 | 48480241 | 48480806 | ENSMUSG00000032026 | Rexo2 | RNA exonuclease 2 |
| chr9 | 48984908 | 48985864 | ENSMUSG00000032267 | Usp28 | ubiquitin specific peptidase 28 |
| chr9 | 50528312 | 50528864 | ENSMUSG00000032067 | Pts | 6-pyruvoyl-tetrahydropterin synthase |
| chr9 | 50616703 | 50617482 | ENSMUSG00000059820 | Nkapd1 | NKAP domain containing 1 |
| chr9 | 51008077 | 51009423 | ENSMUSG00000037112 | Sik2 | salt inducible kinase 2 |
| chr9 | 51963044 | 51963709 | ENSMUSG00000032051 | Fdx1 | ferredoxin 1 |
| chr9 | 52047843 | 52049116 | ENSMUSG00000032050 | Rdx | radixin |
| chr9 | 52167879 | 52168986 | ENSMUSG00000035164 | Zc3h12c | zinc finger CCCH type containing 12C |
| chr9 | 53383816 | 53384612 | ENSMUSG00000034487 | Poglut3 | protein O-glucosyltransferase 3 |
| chr9 | 53771497 | 53772138 | ENSMUSG00000042195 | Slc35f2 | solute carrier family 35, member F2 |
| chr9 | 54501030 | 54502234 | ENSMUSG00000041268 | Dmxl2 | Dmx-like 2 |
| chr9 | 55149167 | 55150192 | ENSMUSG00000032307 | Ube2q2 | ubiquitin-conjugating enzyme E2Q family member 2 |
| chr9 | 56417531 | 56418271 | ENSMUSG00000074305 | Peak1 | pseudopodium-enriched atypical kinase 1 |
| chr9 | 56994822 | 56995929 | ENSMUSG00000032290 | Ptpn9 | protein tyrosine phosphatase, non-receptor type 9 |
| chr9 | 57261453 | 57262844 | ENSMUSG00000032300 | 1700017B05Rik | RIKEN cDNA 1700017B05 gene |
| chr9 | 57521006 | 57521883 | ENSMUSG00000000088 | Cox5a | cytochrome c oxidase subunit 5A |
| chr9 | 57644863 | 57645836 | ENSMUSG00000032312 | Csk | c-src tyrosine kinase |
| chr9 | 57764194 | 57766129 | ENSMUSG00000032316 | Clk3 | CDC-like kinase 3 |
| chr9 | 58487897 | 58489031 | ENSMUSG00000066607 | Insyn1 | inhibitory synaptic factor 1 |
| chr9 | 58554750 | 58555220 | ENSMUSG00000035914 | Cd276 | CD276 antigen |
| chr9 | 58581957 | 58582900 | ENSMUSG00000032336 | Nptn | neuroplastin |
| chr9 | 59035651 | 59037350 | ENSMUSG00000032340 | Neo1 | neogenin |
| chr9 | 59291405 | 59292106 | ENSMUSG00000025236 | Adpgk | ADP-dependent glucokinase |
| chr9 | 59485616 | 59486998 | ENSMUSG00000025234 | Arih1 | ariadne RBR E3 ubiquitin protein ligase 1 |
| chr9 | 59616972 | 59617819 | ENSMUSG00000025237 | Parp6 | poly (ADP-ribose) polymerase family, member 6 |
| chr9 | 59680022 | 59680617 | ENSMUSG00000074259 | Gramd2 | GRAM domain containing 2 |
| chr9 | 59750779 | 59751578 | ENSMUSG00000039585 | Myo9a | myosin IXa |
| chr9 | 62810965 | 62811990 | ENSMUSG00000032244 | Fem1b | fem 1 homolog b |
| chr9 | 63132675 | 63133189 | ENSMUSG00000022245 | Skor1 | SKI family transcriptional corepressor 1 |
| chr9 | 63398577 | 63399712 | ENSMUSG00000032403 | 2300009A05Rik | RIKEN cDNA 2300009A05 gene |
| chr9 | 63757084 | 63758458 | ENSMUSG00000032402 | Smad3 | SMAD family member 3 |
| chr9 | 64340411 | 64341429 | ENSMUSG00000032396 | Dis3l | DIS3 like exosome 3'-5' exoribonuclease |
| chr9 | 65346249 | 65347016 | ENSMUSG00000041837 | Pdcd7 | programmed cell death 7 |
| chr9 | 66059913 | 66060532 | ENSMUSG00000032383 | Ppib | peptidylprolyl isomerase B |
| chr9 | 66511402 | 66512174 | ENSMUSG00000050503 | Fbxl22 | F-box and leucine-rich repeat protein 22 |
| chr9 | 66592236 | 66593176 | ENSMUSG00000032376 | Usp3 | ubiquitin specific peptidase 3 |
| chr9 | 66945919 | 66946683 | ENSMUSG00000036781 | Rps27l | ribosomal protein S27-like |
| chr9 | 66974908 | 66975548 | ENSMUSG00000032370 | Lactb | lactamase, beta |
| chr9 | 67043364 | 67044118 | ENSMUSG00000032366 | Tpm1 | tropomyosin 1, alpha |
| chr9 | 70503105 | 70503915 | ENSMUSG00000032217 | Rnf111 | ring finger 111 |
| chr9 | 70542497 | 70543477 | ENSMUSG00000032212 | Sltm | SAFB-like, transcription modulator |
| chr9 | 70656490 | 70657454 | ENSMUSG00000042444 | Mindy2 | MINDY lysine 48 deubiquitinase 2 |
| chr9 | 70678715 | 70679820 | ENSMUSG00000054693 | Adam10 | a disintegrin and metallopeptidase domain 10 |
| chr9 | 72110951 | 72112353 | ENSMUSG00000032228 | Tcf12 | transcription factor 12 |
| chr9 | 72132786 | 72133536 | ENSMUSG00000032228 | Tcf12 | transcription factor 12 |
| chr9 | 72274687 | 72275552 | ENSMUSG00000038535 | Zfp280d | zinc finger protein 280D |
| chr9 | 72531744 | 72533037 | ENSMUSG00000037674 | Rfx7 | regulatory factor X, 7 |
| chr9 | 72661757 | 72663489 | ENSMUSG00000032216 | Nedd4 | neural precursor cell expressed, developmentally down-regulated 4 |
| chr9 | 74952727 | 74953622 | ENSMUSG00000034858 | Fam214a | family with sequence similarity 214, member A |
| chr9 | 75070900 | 75072037 | ENSMUSG00000034593 | Myo5a | myosin VA |
| chr9 | 75408962 | 75410469 | ENSMUSG00000042688 | Mapk6 | mitogen-activated protein kinase 6 |
| chr9 | 77543709 | 77545008 | ENSMUSG00000032352 | Lrrc1 | leucine rich repeat containing 1 |
| chr9 | 77750510 | 77750987 | ENSMUSG00000032350 | Gclc | glutamate-cysteine ligase, catalytic subunit |
| chr9 | 77754458 | 77755294 | ENSMUSG00000032350 | Gclc | glutamate-cysteine ligase, catalytic subunit |
| chr9 | 77910454 | 77911145 | ENSMUSG00000032349 | Elovl5 | ELOVL family member 5, elongation of long chain fatty acids (yeast) |
| chr9 | 77917264 | 77918004 | ENSMUSG00000032349 | Elovl5 | ELOVL family member 5, elongation of long chain fatty acids (yeast) |
| chr9 | 78480716 | 78482199 | ENSMUSG00000037742 | Eef1a1 | eukaryotic translation elongation factor 1 alpha 1 |
| chr9 | 80066742 | 80067929 | ENSMUSG00000034252 | Senp6 | SUMO/sentrin specific peptidase 6 |
| chr9 | 80164652 | 80165451 | ENSMUSG00000033577 | Myo6 | myosin VI |
| chr9 | 82974106 | 82975184 | ENSMUSG00000032253 | Phip | pleckstrin homology domain interacting protein |
| chr9 | 86464450 | 86465057 | ENSMUSG00000032415 | Ube2cbp | ubiquitin-conjugating enzyme E2C binding protein |
| chr9 | 86466981 | 86467722 | ENSMUSG00000034973 | Dop1a | DOP1 leucine zipper like protein A |
| chr9 | 86695352 | 86696147 | ENSMUSG00000032418 | Me1 | malic enzyme 1, NADP(+)-dependent, cytosolic |
| chr9 | 88438464 | 88439272 | ENSMUSG00000032422 | Snx14 | sorting nexin 14 |
| chr9 | 88522898 | 88523382 | ENSMUSG00000097195 | Snhg5 | small nucleolar RNA host gene 5 |
| chr9 | 92541861 | 92542976 | ENSMUSG00000032374 | Plod2 | procollagen lysine, 2-oxoglutarate 5-dioxygenase 2 |
| chr9 | 94537083 | 94538749 | ENSMUSG00000045414 | Dipk2a | divergent protein kinase domain 2A |
| chr9 | 95637534 | 95638042 | ENSMUSG00000015354 | Pcolce2 | procollagen C-endopeptidase enhancer 2 |
| chr9 | 95749534 | 95750419 | ENSMUSG00000032839 | Trpc1 | transient receptor potential cation channel, subfamily C, member 1 |
| chr9 | 96196092 | 96196887 | ENSMUSG00000032411 | Tfdp2 | transcription factor Dp 2 |
| chr9 | 96363502 | 96364792 | ENSMUSG00000032412 | Atp1b3 | ATPase, Na+/K+ transporting, beta 3 polypeptide |
| chr9 | 96630632 | 96631741 | ENSMUSG00000032413 | Rasa2 | RAS p21 protein activator 2 |
| chr9 | 96719609 | 96720063 | ENSMUSG00000040433 | Zbtb38 | zinc finger and BTB domain containing 38 |
| chr9 | 97018198 | 97019144 | ENSMUSG00000046997 | Spsb4 | splA/ryanodine receptor domain and SOCS box containing 4 |
| chr9 | 99139516 | 99140524 | ENSMUSG00000032462 | Pik3cb | phosphatidylinositol-4,5-bisphosphate 3-kinase catalytic subunit beta |
| chr9 | 99436337 | 99436961 | ENSMUSG00000032470 | Mras | muscle and microspikes RAS |
| chr9 | 99568087 | 99568960 | ENSMUSG00000032468 | Armc8 | armadillo repeat containing 8 |
| chr9 | 100545451 | 100546642 | ENSMUSG00000032475 | Nck1 | non-catalytic region of tyrosine kinase adaptor protein 1 |
| chr9 | 100643023 | 100644584 | ENSMUSG00000037286 | Stag1 | stromal antigen 1 |
| chr9 | 101251026 | 101251947 | ENSMUSG00000043154 | Ppp2r3a | protein phosphatase 2, regulatory subunit B'', alpha |
| chr9 | 102625675 | 102626411 | ENSMUSG00000032534 | Cep63 | centrosomal protein 63 |
| chr9 | 102717108 | 102718896 | ENSMUSG00000032531 | Amotl2 | angiomotin-like 2 |
| chr9 | 102834660 | 102835540 | ENSMUSG00000032547 | Ryk | receptor-like tyrosine kinase |
| chr9 | 103304898 | 103305728 | ENSMUSG00000032555 | Topbp1 | topoisomerase (DNA) II binding protein 1 |
| chr9 | 103364294 | 103366081 | ENSMUSG00000032803 | Cdv3 | carnitine deficiency-associated gene expressed in ventricle 3 |
| chr9 | 104262622 | 104263235 | ENSMUSG00000032560 | Dnajc13 | DnaJ heat shock protein family (Hsp40) member C13 |
| chr9 | 105494512 | 105495566 | ENSMUSG00000032570 | Atp2c1 | ATPase, Ca++-sequestering |
| chr9 | 106685027 | 106686100 | ENSMUSG00000040813 | Tex264 | testis expressed gene 264 |
| chr9 | 106885892 | 106887498 | ENSMUSG00000074102 | Rbm15b | RNA binding motif protein 15B |
| chr9 | 107554243 | 107555511 | ENSMUSG00000010067 | Rassf1 | Ras association (RalGDS/AF-6) domain family member 1 |
| chr9 | 107635064 | 107635608 | ENSMUSG00000032562 | Gnai2 | guanine nucleotide binding protein (G protein), alpha inhibiting 2 |
| chr9 | 107709267 | 107710542 | ENSMUSG00000034684 | Sema3f | sema domain, immunoglobulin domain (Ig), short basic domain, secreted, (semaphorin) 3F |
| chr9 | 107872198 | 107873027 | ENSMUSG00000032582 | Rbm6 | RNA binding motif protein 6 |
| chr9 | 108263180 | 108264389 | ENSMUSG00000039952 | Dag1 | dystroglycan 1 |
| chr9 | 108305860 | 108306970 | ENSMUSG00000039461 | Tcta | T cell leukemia translocation altered gene |
| chr9 | 108347502 | 108348195 | ENSMUSG00000032612 | Usp4 | ubiquitin specific peptidase 4 (proto-oncogene) |
| chr9 | 108516716 | 108518058 | ENSMUSG00000006673 | Qrich1 | glutamine-rich 1 |
| chr9 | 108691958 | 108692911 | ENSMUSG00000032601 | Prkar2a | protein kinase, cAMP dependent regulatory, type II alpha |
| chr9 | 108808206 | 108808764 | ENSMUSG00000032598 | Nckipsd | NCK interacting protein with SH3 domain |
| chr9 | 109931263 | 109932211 | ENSMUSG00000032479 | Map4 | microtubule-associated protein 4 |
| chr9 | 109932211 | 109932485 | ENSMUSG00000032479 | Map4 | microtubule-associated protein 4 |
| chr9 | 110131774 | 110132539 | ENSMUSG00000032481 | Smarcc1 | SWI/SNF related, matrix associated, actin dependent regulator of chromatin, subfamily c, member 1 |
| chr9 | 110531991 | 110533312 | ENSMUSG00000044791 | Setd2 | SET domain containing 2 |
| chr9 | 110653584 | 110654451 | ENSMUSG00000056724 | Nbeal2 | neurobeachin-like 2 |
| chr9 | 110879553 | 110880600 | ENSMUSG00000049555 | Tmie | transmembrane inner ear |
| chr9 | 111117761 | 111118859 | ENSMUSG00000032497 | Lrrfip2 | leucine rich repeat (in FLII) interacting protein 2 |
| chr9 | 111271064 | 111272508 | ENSMUSG00000032498 | Mlh1 | mutL homolog 1 |
| chr9 | 113741139 | 113742143 | ENSMUSG00000033392 | Clasp2 | CLIP associating protein 2 |
| chr9 | 113930814 | 113931911 | ENSMUSG00000009741 | Ubp1 | upstream binding protein 1 |
| chr9 | 114390089 | 114390767 | ENSMUSG00000032431 | Crtap | cartilage associated protein |
| chr9 | 114978314 | 114979165 | ENSMUSG00000040875 | Osbpl10 | oxysterol binding protein-like 10 |
| chr9 | 115309472 | 115311152 | ENSMUSG00000032437 | Stt3b | STT3, subunit of the oligosaccharyltransferase complex, homolog B (S. cerevisiae) |
| chr9 | 116174384 | 116175564 | ENSMUSG00000032440 | Tgfbr2 | transforming growth factor, beta receptor II |
| chr9 | 119321576 | 119323124 | ENSMUSG00000036737 | Oxsr1 | oxidative-stress responsive 1 |
| chr9 | 119401901 | 119403421 | ENSMUSG00000061393 | Acvr2b | activin receptor IIB |
| chr9 | 119982569 | 119983977 | ENSMUSG00000032515 | Csrnp1 | cysteine-serine-rich nuclear protein 1 |
| chr9 | 120011049 | 120011587 | ENSMUSG00000079243 | Xirp1 | xin actin-binding repeat containing 1 |
| chr9 | 120571356 | 120571907 | ENSMUSG00000025794 | Rpl14 | ribosomal protein L14 |
| chr9 | 120933206 | 120934439 | ENSMUSG00000006932 | Ctnnb1 | catenin (cadherin associated protein), beta 1 |
| chr9 | 121297406 | 121298154 | ENSMUSG00000032536 | Trak1 | trafficking protein, kinesin binding 1 |
| chr9 | 121856985 | 121858070 | ENSMUSG00000038412 | Higd1a | HIG1 domain family, member 1A |
| chr9 | 122117209 | 122117893 | ENSMUSG00000038145 | Snrk | SNF related kinase |
| chr9 | 122351345 | 122352246 | ENSMUSG00000032540 | Abhd5 | abhydrolase domain containing 5 |
| chr9 | 123020737 | 123021586 | ENSMUSG00000025787 | Tgm4 | transglutaminase 4 (prostate) |
| chr9 | 123112777 | 123113700 | ENSMUSG00000025785 | Exosc7 | exosome component 7 |
| chr9 | 123259698 | 123261019 | ENSMUSG00000054871 | Tmem158 | transmembrane protein 158 |
| chr9 | 123529524 | 123530330 | ENSMUSG00000025240 | Sacm1l | SAC1 suppressor of actin mutations 1-like (yeast) |
| chr9 | 123924566 | 123925123 | ENSMUSG00000025804 | Ccr1 | chemokine (C-C motif) receptor 1 |
| chrX | 12761308 | 12762475 | ENSMUSG00000064127 | Med14 | mediator complex subunit 14 |
| chrX | 13071706 | 13072942 | ENSMUSG00000031010 | Usp9x | ubiquitin specific peptidase 9, X chromosome |
| chrX | 13280282 | 13282255 | ENSMUSG00000000787 | Ddx3x | DEAD box helicase 3, X-linked |
| chrX | 18161961 | 18163680 | ENSMUSG00000037369 | Kdm6a | lysine (K)-specific demethylase 6A |
| chrX | 48452950 | 48454634 | ENSMUSG00000031103 | Elf4 | E74-like factor 4 (ets domain transcription factor) |
| chrX | 51205037 | 51206000 | ENSMUSG00000036109 | Mbnl3 | muscleblind like splicing factor 3 |
| chrX | 60403575 | 60404622 | ENSMUSG00000062949 | Atp11c | ATPase, class VI, type 11C |
| chrX | 71364559 | 71365219 | ENSMUSG00000015214 | Mtmr1 | myotubularin related protein 1 |
| chrX | 71555728 | 71557266 | ENSMUSG00000015217 | Hmgb3 | high mobility group box 3 |
| chrX | 73228171 | 73229352 | ENSMUSG00000078317 | F8a | factor 8-associated gene A |
| chrX | 94122435 | 94123496 | ENSMUSG00000079509 | Zfx | zinc finger protein X-linked |
| chrX | 140598389 | 140600312 | ENSMUSG00000031431 | Tsc22d3 | TSC22 domain family, member 3 |
| chrX | 152769114 | 152770230 | ENSMUSG00000045180 | Shroom2 | shroom family member 2 |
| chrX | 157491331 | 157492249 | ENSMUSG00000071708 | Sms | spermine synthase |
| chrX | 159372079 | 159372775 | ENSMUSG00000067194 | Eif1ax | eukaryotic translation initiation factor 1A, X-linked |
